# *Inventa*: a computational tool to discover chemical novelty in natural extracts libraries

**DOI:** 10.1101/2022.08.25.505324

**Authors:** Luis-Manuel Quiros-Guerrero, Louis-Félix Nothias, Arnaud Gaudry, Laurence Marcourt, Pierre-Marie Allard, Adriano Rutz, Bruno David, Emerson Ferreira Queiroz, Jean-Luc Wolfender

## Abstract

Collections of natural extracts hold potential for the discovery of novel natural products with original modes of action. The prioritization of extracts from collections remains challenging due to the lack of workflow that combines multiple-source information to facilitate the data interpretation. Results from different analysis techniques and literature reports need to be organized, processed, and interpreted to enable optimal decision-making for extracts prioritization. Here, we introduce *Inventa*, a computational tool that highlights the chemical novelty potential within extracts, considering untargeted mass spectrometry data, spectral annotation, and literature reports. Based on this information, *Inventa* calculates multiple scores that inform their chemical potential. Thus, *Inventa* has the potential to accelerate new natural products discovery. *Inventa* was applied to a set of plants from the Celastraceae family as a proof of concept. The *Pristimera indica* (Willd.) A.C.Sm roots extract was highlighted as a promising source of potentially novel compounds. Its phytochemical investigation resulted in the isolation and *de novo* characterization of thirteen new dihydro-*β*-agarofuran sesquiterpenes, five of them presenting a new 9-oxodihydro-*β*-agarofuran base scaffold.

## 1 Introduction

Natural products (NPs) are specialized metabolites from different biological sources like plants, fungi, bacteria, and marine organisms, have enormously contributed to and inspired the development of drugs (Newman and Cragg, 2020). These biodiverse sources often produce NPs with complex molecular structures displaying remarkable bioactivities and represent an unique source of novel scaffolds with unprecedented modes of action (Howes, 2018; Howes et al., 2020; Verma et al., 2020). In NPs research, the prioritization of extracts produced from these collections is a keystone for the continuous discovery of novel bioactive specialized metabolites (Wolfender et al., 2019).

After the 1980s, NPs researchers started facing the problem of re-isolating known chemical entities, resulting in a waste of time and resources, which continues until today. Dereplication structure-based approaches were designed to assist the classical bio-guided isolation workflow to reduce the reisolation problem. These approaches can obtain information on extracts based on the expressed and potential metabolism via compound dereplication, metabolomics, or genome mining (Henke and Kelleher, 2016; Louwen and van der Hooft, 2021; Singh et al., 2022). While genome mining strategies became central for studying microbial NPs, it is not presently fully applicable to plants (Pieters and Vlietinck, 2005; Henke and Kelleher, 2016; Medema et al., 2021).

Multiple strategies have been proposed to prioritize extracts and efficiently isolate compounds displaying interesting bioactivity and novel structural properties. For example, classical metabolomic studies combine mass spectrometry, a particular bioactivity test, and chemometrics to highlight extracts through statistics (Fiehn, 2002). The integration of genomic information has recently enhanced the capacity to point out extracts based on the potential of their phenotypic expression (Caesar et al., 2021). The introduction of Molecular Networking (MN) allowed visualization and interpretation of relatively large spectral/chemical spaces, easing the comparison of the extracts at the spectral level (Wang et al., 2016). MN can be combined with bioactivity test results and dereplication information to prioritize particular features [a peak with an *m/z* value at a given retention time (RT)] within an extract by novelty or biological activity potential (Olivon et al., 2017; Nothias et al., 2018; Fox Ramos et al., 2019; Wolfender et al., 2019).

Other published studies proposed mass-spectrometry-based workflows selecting extracts to accelerate the discovery of novel NPs, for example, utilizing liquid chromatography-mass spectrometry profiling and MS^1^ level (exact mass and molecular formula match) annotation rates against databases of NPs. This study was centered on the discovery of novel marine NPs. It classified the extracts based on the presence and proportion of features in the chromatogram with a particular set of scores based on their area and intensity. The scores tried to reflect each extract’s chemical complexity and structural novelty. The application of this workflow in a small set of marine sponges and tunicates extracts resulted in the isolation of two new eudistomin analogs and two new nucleosides (Tabudravu et al., 2019). Another study proposed using the CSCS metric (Sedio et al., 2018) to prioritize extracts according to their spectral uniqueness in a set of fungal extracts. It is based on the principle that dissimilar extracts would hold a particular chemistry, different from the ensemble of extracts. Recently, an application of this workflow led to the isolation of three new drimane-type sesquiterpenes (Pham et al., 2021). Finally, <u>FERMO</u> is a tool presently in development for the prioritization of relevant bioactive compounds (metabolites) within natural extracts based on chromatographic characteristics, bioactivity, and dereplication results. This tool aims to explore and suggest peaks of interest in a particular extract for isolation (Zdouc M., Medema M., van der Hooft J.).

With the increasing capacities of the analytical profiling techniques, and the broad applications of bioinformatics tools in the field of NPs chemistry, the quantity of analytical information obtained increased proportionally. The clear and concise analysis of the resulting massive datasets is challenging and reduces the efficiency of data-driven prioritization (Brejnrod et al., 2019; Caesar et al., 2021; Amara et al., 2022). This is partly due to the time-consuming efforts required for the manual exploration of the data, the compilation of literature reports for individual organisms, the interpretation of the spectral annotation results, and extract comparison techniques. Yet, even after carefully curating the data and the results, exploring and interpreting all this information is the main bottleneck to efficiently prioritized the extracts with the highest chemical potential within collections (Louwen and van der Hooft, 2021). The conception and implementation of comprehensive prioritization pipelines that combine results from several bioinformatic tools are imperative to speed up and rationalize extract selection.

Here, we introduce <u>*Inventa*</u>, a computational tool that highlights the chemical novelty potential of novel NPs within extracts, considering untargeted mass spectrometry data and literature reports for the organism’s taxa of interest. It was designed to accelerate mining data sets in a scalable manner. As a proof of concept, we applied it to a collection of taxonomically related extracts of the Celastraceae family. Plants from this family are characterized for producing a wide range of specialized bioactive metabolites from different chemical classes, like macrolide sesquiterpene pyridine alkaloids (Callies et al., 2017), maytansinoids (Kupchan et al., 1972), and quinone methide triterpenoids (Alvarenga and Ferro, 2006; Salminen et al., 2010). Most of them have important pharmacological importance (González et al., 2000; Moin et al., 2014; Lv et al., 2019), and some are considered chemotaxonomic markers for particular genera and the family (González et al., 1986; Rogers et al., 2000).

In this study, we present the application of *Inventa* for selecting extracts based on predicted chemical novelty. The data generated from the Celastraceae set was used to explore the effect of the various parameters which led to the prioritization of an extract from seventy-six and the subsequent isolation and structural identification of thirteen molecules.

## 2 Materials and Methods

### 2.1 Chemicals

HPLC grade methanol (MeOH) and ethyl acetate (EtOAc) were purchased from Fisher Chemicals, Reinach, Switzerland, LC-MS grade water, acetonitrile (ACN), and formic acid were purchased from Fisher Chemicals, Reinach, Switzerland, Dimethyl Sulfoxide (DMSO) molecular biology grade was purchased from Sigma, St Louis, USA.

### 2.2 General Experimental Procedures

NMR spectroscopic data were recorded on a Bruker Avance Neo 600 MHz spectrometer equipped with a QCI 5mm Cryoprobe and a sampleJet automated extract changer (Bruker BioSpin, Rheinstetten, Germany). Chemical shifts are reported in parts per million (ppm, δ), and coupling constants are reported in Hz (*J*). The residual CD_3_OD signals (δ_H_ 3.31, δ_C_ 49.8) were used as internal standards for ^1^H and ^13^C, respectively. Complete assignments were based on 2D-NMR spectroscopy: COSY, edited-HSQC, HMBC, and ROESY. The Electronic Circular Dichroism (ECD) was recorded on a JASCO J-815 spectrometer (Loveland, CO, United States) in acetonitrile using a 1 cm cell. The scan speed was 200 nm/min in continuous mode between 600 nm and 150 nm. The optical rotations were measured in acetonitrile on a JASCO P-1030 polarimeter (Loveland, CO, USA) in a 1 mL, 10 cm tube.

### 2.3 Plant Material, Small Scale Extraction, and extract Preparation for UHPLC-HRMS/MS Analysis of the Celastraceae set

#### 2.3.1 Plant Material

The set comprises seventy-six extracts from different plant parts (leaves, stems, roots, fruits, seeds, bark, and branches) of thirty-six species belonging to fourteen different genera. These plants belong to the Pierre-Fabre Laboratories (PFL) collection with over 17,000 unique samples collected worldwide. The PFL collection was registered at the European Commission under the accession number 03-FR-2020.This registration certifies that the collection meets the criteria set out in the EU ABS Regulation which implements at EU level the requirements of the Nagoya Protocol regarding access to genetic resources and the fair and equitable sharing of benefits arising from their utilization (https://ec.europa.eu/environment/nature/biodiversity/international/abs/pdf/Register%20of%20Collections.pdf). The PFL supplied all the vegetal material (grounded dry material). The collected samples have photographs, herbarium vouchers, and leaf extracts preserved in dry silica gel. Precise localization of the initial collection, unique ID and barcode, and GPS data are stored in the dedicated data management system. The plant material was dried for three days at 55 °C in an oven; then the material was grounded and stored in plastic pots at a controlled temperature and humidity in the Pierre-Fabre Laboratories facilities.

#### 2.3.2 Taxonomical Metadata

The taxonomic names were searched in the <u>Open Tree of Life</u> (OTL v13.4) (Rees and Cranston, 2017) to most recent ‘accepted’ name. The metadata added includes, if found, the taxon’s OTTid, rank, source, all available synonyms, and their corresponding references (NCBI, GBIF, IRMNG). When the species was not defined, the next genus was used in the search. The original genus and species names provided with the collection are kept in the respective columns.

#### 2.3.3 UHPLC-HRMS/MS Analysis

Analyses were performed with a Waters Acquity UPLC system equipped with a PDA detector coupled to a Q-Exactive Focus mass spectrometer (Thermo ScientificTM, Bremen, Germany), employing a heated electrospray ionization source (HESI-II) with the following parameters: spray voltage: + 3.5 kV; heater temperature: 220 ºC; capillary temperature: 350.00 ºC; S-lens RF: 45 (arb. units); sheath gas flow rate: 55 (arb. units) and auxiliary gas flow rate: 15.00 (arb. units). The mass analyzer was calibrated using a mixture of caffeine, methionine–arginine–phenylalanine–alanine–acetate (MRFA), sodium dodecyl sulfate, and sodium taurocholate, and Ultramark 1621 in an acetonitrile/methanol/water solution containing 1% formic acid by direct injection. The system was coupled to a Charged aerosol detector (CAD, Thermo ScientificTM, Bremen, Germany) kept at 40 ºC. The PDA wavelength range was from 210 nm to 400 nm with a resolution of 1.2 nm. Control of the instruments was done using Thermo Scientific Xcalibur 3.1 software.

For the centroid data-dependent MS^2^ (dd-MS^2^) experiments in positive ionization mode, full scans were acquired at a resolution of 35,000 FWHM (at *m/z* 200) and MS^2^ scans at 17,500 FWHM in the range 100 to 1500 *m/z*. The dd-MS^2^ scan acquisition events were performed in discovery mode with an isolation window of 1.5 Da and stepped normalized collision energy (NCE) of 15, 30, and 45 units. Additional parameters were set as follows: default mass charge: 1; Automatic gain control (AGC) target 2E^5^; Maximum IT: 119 ms; Loop count: 3; Min AGC target: 2.6E^4^; Intensity threshold: 1. Up to three dd-MS^2^ scans (Top 3) were acquired for the most abundant ions per scan in MS^1^, using the Apex trigger mode (2 to 7 s), dynamic exclusion (9.0 s), and automatic isotope exclusion. A specific exclusion list was created for the measurement using the solvent as a background extract with an IODA Mass Spec notebook (Zuo et al., 2021).

The chromatographic separation was done on a Waters BEH C18 column (50× 2.1 mm i.d., 1.7 µm, Waters, Milford, MA, USA) through a linear gradient of 5−100% B over 7 min and an isocratic step at 100% B for 1 min. The mobile phases were: (A) water with 0.1% formic acid and (B) acetonitrile with 0.1% formic acid. The flow rate was set to 600 µL/min, the injection volume was 2 µL, and the column was kept at 40 °C. The set of extracts was randomized before injection, including pooled QC extracts and blanks, repeated once every ten extracts.

#### 2.3.4 UHPLC-HRMS/MS Data Analysis

##### 2.3.4.1 Data Preprocessing

The data were converted from .RAW (Thermo) standard data format to an open .mzXML format employing the MS Convert software, part of the ProteoWizard package (Chambers et al., 2012). The converted files were processed with the <u>MZmine3</u> software (Pluskal et al., 2010). For mass detection at the MS^1^ level, the noise level was set to 1.0E^6^ for positive mode and 1.0E^5^ for negative mode. For MS^2^ detection, the noise level was set to 0.00 for both ionization modes. The ADAP chromatogram builder parameters were set as follows: minimum group size in # of scans, 4; Group intensity threshold, 1.0E^6^ (1.0E^5^ negative); Minimum highest intensity, 1.0E^6^ (1.0E^5^ negative) and Scan to scan accuracy (*m/z*) of 0.0020 or 10.0 ppm. The ADAP feature resolver algorithm was used for chromatogram deconvolution with the following parameters: S/N threshold, 30; minimum feature height, 1.0E^6^ (1.0E^5^ negative); coefficient area threshold, 110; peak duration range, 0.01 - 1.0 min; RT wavelet range, 0.01 - 0.08 min. Isotopes were detected using the 13C isotope filter with an m/z tolerance of 0.0050 or 8.0 ppm, an Retention Time tolerance of 0.03 min (absolute), the maximum charge set at 2, and the representative isotope used was the lowest m/z. Each file was filtered by RT (positive mode: 0.70 - 8.00 min, negative mode: 0.40 - 8.00 min), and only the ions with an associated MS^2^ spectrum were kept. Alignment was done with the join-aligner (m/z tolerance, 0.0050 or 8.0 ppm; RT tolerance, 0.05 min), and the align list was filtered to remove any duplicate (m/z tolerance, 8.0 ppm; RT tolerance, 0.10 min).

The resulting filtered list was subjected to Ion Identity Networking (Schmid et al., 2021) starting with the metaCorrelate module (RT tolerance, 0.10 min; minimum height, 1.0E^5^; Intensity correlation threshold 1.0E^5^ and the Correlation Grouping with the default parameters). Followed by the Ion identity networking (m/z tolerance, 8.0 ppm; check: one feature; minimum height: 1.0E^5^, annotation library [maximum charge, 2; maximum molecules/cluster, 2; Adducts ([M+H]^+^, [M+Na]^+^, [M+K]^+^, [M+NH_4_]^+^, [M+2H]^2+^), Modifications ([M-H_2_O], [M-2H_2_O], [M-CO_2_], [M+HFA], [M+ACN])], Annotation refinement (Delete small networks without major ion, yes; Delete networks without monomer, yes), Add ion identities networks (m/z tolerance, 8 ppm; minimum height, 1.0E^5^; Annotation refinement (Minimum size, 1; Delete small networks without major ion, yes; Delete small networks: Link threshold, 4; Delete networks without monomer, yes)) and Check all ion identities by MS/MS (m/z tolerance (MS^2^), 10 ppm; min-height (in MS^2^), 1.0E^3^; Check for multimers, yes; Check neutral losses (MS^1^ ->MS^2^), yes) modules. The resulting aligned peak list was exported as a .mgf file for further analysis.

##### 2.3.4.2 MS/MS Spectral Organization

A molecular network was constructed from the .*mgf* file exported from MZmine, using the feature-based molecular networking workflow (https://ccms-ucsd.github.io/GNPSDocumentation/) on the <u>GNPS</u> website (Nothias et al., 2020). The precursor ion mass tolerance was set to 0.02 Da with an MS/MS fragment ion tolerance of 0.02 Da. A network was created where edges were filtered to have a cosine score above 0.7 and more than six matched peaks. The spectra in the network were then searched against GNPS’ spectral libraries. All matches between network and library spectra were required to have a score above 0.6, and at least three matched peaks. Jobs links: https://gnps.ucsd.edu/ProteoSAFe/status.jsp?task=df71854c6e644b979228d96b521a490b (positive), https://gnps.ucsd.edu/ProteoSAFe/status.jsp?task=d477f360ddb344a593b935624782d8eb (negative).

##### 2.3.4.3 Taxonomically Informed Metabolite Annotation

The .*mgf* file exported from MZmine was also annotated by spectral matching against an *in-silico* database to obtain putative annotations (Allard et al., 2016). The resulting annotations were subjected to taxonomically informed metabolite scoring (Rutz et al., 2019) (https://taxonomicallyinformedannotation.github.io/tima-r/, v 2.4.0) and re-ranking from the chemotaxonomical information available on <u>LOTUS</u> (Rutz et al., 2022).The *in-silico* database used for this process includes the combined records of the Dictionary of Natural Products (DNP, v 30.2) and the <u>LOTUS</u> Initiative outputs (Rutz et al., 2022).

##### 2.3.4.4 SIRIUS Metabolite Annotation

The SIRIUS .mgf file exported from MZmine (using the SIRIUS export module) that contains MS1 and MS2 information was processed with SIRIUS (v 5.5.5) command-line tools on a Linux server (Dührkop et al., 2019). The molecular formula and metabolite database used for SIRIUS includes NPs from LOTUS (Rutz et al., 2022) and the Dictionary of Natural Products (DNP). The parameters were set as follows: *Possible ionizations*: [M+H]^+^, [M+NH_4_]^+^, [M-H_2_O+H]^+^, [M+K]^+^, [M+Na]^+^,[M-4H2O+H]^+^; *Instrument profile*: Orbitrap; *mass accuracy*: 5 ppm for MS^1^ and 7 ppm for MS^2^, database for molecular formulas and structures:BIO and custom databases (LOTUS, DNP), *maximum m/z to compute*: 1000. ZODIAC was used to improve molecular formula prediction using a threshold filter of 0.99 a (Ludwig et al., 2020). Metabolite structure prediction was made with CSI: FingerID (Dührkop et al., 2015) and significance computed with COSMIC (Hoffmann et al., 2021). The chemical class prediction was made with CANOPUS (Dührkop et al., 2020) using the NPClassifier ontology (Kim et al., 2021).

##### 2.3.4.5 Mass Spectrometry-based extract Vectorization <u>(MEMO)</u>

The MS^2^ spectra were processed with the memo_ms package (0.1.3). The parameters were set as follows: *min_rel_intensity*: 0.01, *max_relative_intensity*: 1, *min_peaks_required*: 10, *losses_from*: 10, l*osses_to*: 00, *n_decimal*: 2. All the Peak/loss present in the blanks were removed before the computation of the distance matrix (Gaudry et al., 2022).

### 2.4 Implementation of *Inventa*

All the previously described information was fed into a set of scripts called *Inventa* (https://luigiquiros.github.io/inventa/v1.0.0). These scripts are made available as a Jupyter notebook that can be deployed directly on the cloud using a <u>Binder</u> link(Project Jupyter et al., 2018). All the components were calculated, and the same weight (w =1) was given to each. For the cleaning-up of the GNPS annotations the following parameters were used, *max_ppm_error*: 5, *shared_peaks*: 10, *min_cosine*: 0.6, *ionisation_mode*: ‘pos’, *max_spec_charge*: 2. For calculation of the feature component the following parameters were used, *min_specificity*: 0.9, *min_score_final*: 0.3, *min_ZODIACScore*: 0.9, *min_ConfidenceScore*: 0.25, *annotation_preference*: 0. For the literature component calculations the max_comp_reported_sp, *max_comp_reported_g, max_comp_reported_f* were set to 20, 100, 500 respectively. For the class component, the following parameters were used: *min_class_confidence*: 0.8 and *min_recurrence*: 0.8. The results displayed in the manuscript were based on the MZmine3 Ion Identity Networking. A complete glossary for terms and default parameters can be found in the Supporting Information Table I.

### 2.5 Extraction and Isolation of Compounds from the *Pristimera indica* Roots

The dried ground roots of *Pristimera indica* (Willd.) A.C.Sm. (19.8 g) were extracted successively with hexane (3 × 200 mL), EtOAc (3 × 200 mL), and MeOH (3 × 200 mL), with constant agitation at room temperature for a 12h period each. The organic solvents were filtered and evaporated under reduced pressure to give 61.5 mg of hexane extract, 100.4 mg of ethyl acetate extract, and 728.3 mg of methanolic extract.

Separations were performed in a semi-preparative Shimadzu system equipped with a LC-20A module pumps, an SPD-20A UV/Vis, a 7725I Rheodyne® valve, and an FRC-10A fraction collector (Shimadzu, Kyoto, Japan). The HPLC conditions were as follows: X-Bridge C_18_ column (250 × 19 mm i.d., 5 *μ*m) equipped with a Waters C18 precolumn cartridge holder (10 × 19 mm i.d., 5 *μ*m); solvent system ACN (B) and H_2_O (A), both containing 0.1% FA. The separation was performed in gradient mode as follows: 5 to 40% B in 5 min, 40 to 55% B in 52 min, and 55 to 100% B in 25 min. The flow rate was fixed to 17.0 mL/min. The extract was injected by dry load according to a protocol developed in our laboratory (Queiroz et al., 2019). The collection was done based on the UV/Vis trace peaks at 254 nm.

From the ethyl acetate extract (59.2 mg) 13 fractions (corresponding to the HPLC-UV peaks) were collected to give pure compounds **1** (0.8 mg, t_R_ 21.5 min), **2** (0.7 mg, t_R_ 22.0 min), **3** (1.5 mg, t_R_ 22.5 min), **6** (1.2 mg, t_R_ 23.0 min), **7** (0.6 mg, t_R_ 36.0 min), **8** (0.8 mg, t_R_ 40.0 min), **12** (0.4 mg, t_R_ 41.0 min), **5** (0.6 mg, t_R_ 44.0 min), **13** (0.7 mg, t_R_ 48.5 min), **9** (0.9 mg, t_R_ 50.0 min), **4** (0.4 mg, t_R_ 63.0 min). The fraction collected at t_R_ 34.5 min (0.8 mg), was separated in a X-Bridge C_18_ column (250 × 10 mm i.d., 5 μm) equipped with a Waters C18 precolumn cartridge holder (5 × 10 mm i.d., 5 *μ*m); solvent system ACN (B) and H_2_O (A), both containing 0.1% FA, in an isocratic run 50% ACN, to give **8** (0.2 mg, t_R_ 18.0 min) and **10** (0.4 mg, t_R_ 15.0 min).

The methanolic extract (276.8 mg) was fractionated, in the same conditions as the ethyl acetate extract, to give compounds **1** (1.9 mg, t_R_ 21.5 min), **2** (1.3 mg, t_R_ 22.0 min), **3** (2.6 mg, t_R_ 22.5 min), **6** (0.3 mg, t_R_ 23.0 min), **7** (1.1 mg, t_R_ 36.0 min), **8** (1.3 mg, t_R_ 40.0 min), **5** (0.5 mg, t_R_ 44.0 min), **4** (0.6 mg, t_R_ 63.0 min), **10** (0.3 mg, t_R_ 31.5 min) and **11** (0.4 mg, t_R_ 22.0 min). Fractions collected at t_R_ 41.0 min (0.9 mg) and t_R_ 48.5 min (0.4 mg), were re-purified in a X-Bridge C_18_ column (250 × 10 mm i.d., 5 *μ*m) equipped with a Waters C18 pre-column cartridge holder (5 × 10 mm i.d., 5 *μ*m); solvent system ACN (B) and H_2_O (A), both containing 0.1% FA, in an isocratic run 50% ACN, to give **13** (0.2 mg, t_R_ 27.0 min).

#### 2.5.1 Description of the Isolated Compounds

Compound **1** ((1*R*,2*S*,4*R*,5*S*,6*R*,7*S*,8*S*,10*S*)-1*α*,6*β*-diacetoxy-15-*iso*-butanoyloxy-2*α*,8*β*-di-(5-carboxy-*N*-methyl-3-pyridoxy)-9-oxodihydro-*β*-agarofuran, Silviatine A). Amorphous white powder; 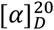 + 25 (ACN); UV (ACN) λ_max_ 193, 270 nm.

^1^H NMR (CD_3_OD, 600 MHz) δ 1.22 (3H, d, *J* = 7.6 Hz, H_3_-14), 1.24 (3H, d, *J* = 7.0 Hz, H_3_-15d), 1.27 (3H, d, *J* = 7.0 Hz, H_3_-15c), 1.42 (3H, s, H_3_-13), 1.49 (3H, s, H_3_-12), 1.95 (3H, s, H_3_-1b), 1.97 (1H, dd, *J* = 14.3, 3.0 Hz, H-3α), 2.19 (3H, s, H_3_-6b), 2.40 (2H, m, H-3β, H-4), 2.80 (1H, hept, *J* = 7.0 Hz, H-15b), 2.97 (1H, d, *J* = 3.4 Hz, H-7), 3.62 (3H, s, H_3_-8g), 3.70 (3H, s, H_3_-2g), 4.70 (1H, d, *J* = 12.2 Hz, H-15’’), 5.10 (1H, d, *J* = 12.2 Hz, H-15’), 5.54 (1H, q, *J* = 3.9, 3.0 Hz, H-2), 5.74 (1H, d, *J* = 3.9 Hz, H-1), 6.03 (1H, d, *J* = 3.4 Hz, H-8), 6.31 (1H, s, H-6), 6.56 (1H, d, *J* = 9.5 Hz, H-8e), 6.58 (1H, d, *J* = 9.5 Hz, H-2e), 7.95 (1H, dd, *J* = 9.5, 2.5 Hz, H-8f), 8.01 (1H, dd, *J* = 9.5, 2.6 Hz, H-2f), 8.44 (1H, d, *J* = 2.5 Hz, H-8c), 8.48 (1H, d, *J* = 2.6 Hz, H-2c); ^13^C NMR (CD_3_OD, 151 MHz) δ 17.5 (CH_3_-14), 19.2 (CH_3_-15c), 19.3 (CH_3_-15d), 20.9 (CH_3_-6b), 21.0 (CH_3_-1b), 27.2 (CH_3_-13), 30.2 (CH_3_-12), 31.6 (CH_2_-3), 34.1 (CH-4), 35.2 (CH-15b), 38.7 (CH_3_-2g, CH_3_-8g), 54.1 (CH-7), 60.2 (C-10), 63.9 (CH_2_-15), 71.1 (CH-1), 72.5 (CH-2), 77.6 (CH-6), 78.8 (CH-8), 85.7 (C-11), 91.9 (C-5), 110.7 (C-8b), 111.1 (C-2b), 119.6 (CH-8e), 119.9 (CH-2e), 140.5 (CH-8f), 140.6 (CH-2f), 145.7 (CH-2c), 146.3 (CH-8c), 165.2 (C-2d, C-8d), 171.4 (C-1a, C-6a), 177.8 (C-15a), 203.1 (C-9).For NMR spectra see **Supplementary Figures S1-S6**. HRESIMS *m/z* 741.2864 [M+H]^+^ (calculated for C_37_H_45_N_2_O_14_, error -0.13 ppm). MS/MS spectrum: <u>CCMSLIB00009919267</u>.

SMILES:CCC(C)C(=O)OC[C@@]12[C@@H](O)[C@H](C[C@@H](C)[C@]11OC(C)(C)[C@@H]([C@H]1OC(C)=O)[C@H](OC(=O)C1=CN(C)C(=O)C=C1)C2=O)OC(=O)C1=CN(C)C(=O)C=C1. InChIKey=HPZNCFSLZGFDST-SMRRRHQGNA-N.

Compound **2**: (1*R*,2*S*,4*R*,5*S*,6*R*,7*S*,8*S*,10*S*)-6*β*-acetoxy-2*α*,8*β*-di-(5-carboxy-*N*-methyl-3-pyridoxy)-1*α*-hydroxy-15-(2-methylbutanoyloxy)-9-oxodihydro-*β*-agarofuran, Silviatine B. Amorphous white powder, 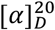 + 19 (ACN); UV (ACN) λ_max_ 200, 268 nm.

^1^H NMR (CD_3_OD, 600 MHz) δ 0.93 (3H, t, *J* = 7.5 Hz, H_3_-15d), 1.15 (3H, d, *J* = 7.7 Hz, H_3_-14), 1.27 (3H, d, *J* = 7.0 Hz, H_3_-15e), 1.43 (3H, s, H_3_-13), 1.48 (3H, s, H_3_-12), 1.58 (1H, m, H-15c’’), 1.77 (1H, m, H-15c’), 1.98 (1H, dd, *J* = 14.2, 3.5 Hz, H-3α), 2.18 (3H, s, H_3_-6b), 2.31 (1H, ddd, *J* = 15.2, 6.4, 3.8 Hz, H-3β), 2.36 (1H, m, H-4), 2.67 (1H, h, *J* = 7.0 Hz, H-15b), 2.97 (1H, d, *J* = 3.4 Hz, H-7), 3.63 (3H, s, H_3_-8g), 3.71 (3H, s, H_3_-2g), 4.61 (1H, d, *J* = 3.8 Hz, H-1), 4.77 (1H, d, *J* = 12.2 Hz, H-15’’), 5.05 (1H, d, *J* = 12.2 Hz, H-15’), 5.44 (1H, q, *J* = 3.8, 3.5 Hz, H-2), 6.10 (1H, d, *J* = 3.4 Hz, H-8), 6.24 (1H, s, H-6), 6.57 (2H, 2xd, *J* = 9.5 Hz, H-2e, H-8e), 7.97 (1H, dd, *J* = 9.5, 2.5 Hz, H-8f), 8.04 (1H, dd, *J* = 9.5, 2.6 Hz, H-2f), 8.47 (1H, d, *J* = 2.5 Hz, H-8c), 8.48 (1H, d, *J* = 2.6 Hz, H-2c); ^13^C NMR (CD_3_OD, 151 MHz) δ 11.6 (CH_3_-15d), 16.5 (CH_3_-15e), 17.4 (CH_3_-14), 20.6 (CH_3_-6b), 27.1 (CH_3_-13), 27.4 (CH_2_-15c), 29.9 (CH_3_-12), 31.3 (CH_2_-3), 34.1 (CH-4), 38.3 (CH_3_-2g), 38.4 (CH_3_-8g), 42.0 (CH-15b), 54.0 (CH-7), 61.4 (C-10), 63.6 (CH_2_-15), 70.0 (CH-1), 74.7 (CH-2), 77.4 (CH-6), 78.4 (CH-8), 85.0 (C-11), 91.3 (C-5), 110.5 (C-8b), 111.3 (C-2b), 119.4 (CH-2e, CH-8e), 140.1 (CH-8f), 140.5 (CH-2f), 145.1 (CH-2c), 145.9 (CH-8c), 163.6 (C-8a), 164.1 (C-2a), 165.0 (C-2d), 164.8 (C-8d), 171.1 (C-6a), 177.8 (C-15a), 206.8 (C-9). For NMR spectra see **Supplementary Figures S7-S11**. HRESIMS *m/z* 713.2323 [M+H]^+^ (calculated for C_36_H_45_N_2_O_13_, error -1.043 ppm);MS/MS spectrum: <u>CCMSLIB00009919268</u>.

SMILES:

O=C1[C@](OC(C2=C([H])N(C([H])([H])[H])C(C([H])=C2[H])=O)=O)([H])[C@](C3(C([H])([H])[H])C([H])([H])[H])([H])[C@](OC(C([H])([H])[H])=O)([H])[C@]4(O3)[C@@](C([H])([H])[H])([H])C([H])([H])C(OC(C(C([H])=C5[H])=C([H])N(C([H])([H])[H])C5=O)=O)([H])[C@@](O[H])([H])[C@]41C([H])([H])OC([C@@](C([H])([H])[H])([H])C([H])([H])C([H])([H])[H])=O. InChIKey=HPZNCFSLZGFDST-SWBINLJCSA-N.

Compound **4**: (1*R*,2*S*,4*R*,5*S*,6*R*,7*S*,8*S*,10*S*)-1α,6β-diacetoxy-2α,8β-di-(5-carboxy-*N*-methyl-3-pyridoxy)-15-(2-methylbutanoyloxy)-9-oxodihydro-*β*-agarofuran, Silviatine C. Amorphous white powder, 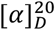 + 33 (ACN); UV (ACN) λ_max_ 194, 267 nm

^1^H NMR (CD_3_OD, 600 MHz) δ 0.92 (3H, t, *J* = 7.4 Hz, H_3_-15d), 1.22 (3H, d, *J* = 7.7 Hz, H_3_-14), 1.25 (3H, d, *J* = 7.0 Hz, H_3_-15e), 1.41 (3H, s, H_3_-13), 1.49 (3H, s, H_3_-12), 1.57 (1H, m, H-15c’’), 1.76 (1H, m, H-15c’), 1.95 (3H, s, H_3_-1b), 1.97 (1H, dd, 14.0, 2.8 Hz, H-3α), 2.19 (3H, s, H_3_-6b), 2.41 (2H, m, H-3β, H-4), 2.65 (1H, h, *J* = 7.0 Hz, H-15b), 2.97 (1H, dd, *J* = 3.4, 0.8 Hz, H-7), 3.62 (3H, s, H_3_-8g), 3.70 (3H, s, H_3_-2g), 4.65 (1H, d, *J* = 12.3 Hz, H-15’’), 5.12 (1H, d, *J* = 12.3 Hz, H-15’), 5.55 (1H, q, *J* = 3.9, 2.8 Hz, H-2), 5.75 (1H, d, *J* = 3.9 Hz, H-1), 6.01 (1H, d, *J* = 3.4 Hz, H-8), 6.30 (1H, d, *J* = 0.8 Hz, H-6), 6.56 (1H, d, *J* = 9.5 Hz, H-8e), 6.58 (1H, d, *J* = 9.5 Hz, H-2e), 7.95 (1H, dd, *J* = 9.5, 2.5 Hz, H-8f), 8.01 (1H, dd, *J* = 9.5, 2.5 Hz, H-2f), 8.44 (1H, d, *J* = 2.5 Hz, H-8c), 8.47 (1H, d, *J* = 2.5 Hz, H-2c); ^13^C NMR (CD_3_OD, 151 MHz) δ 11.9 (CH_3_-15d), 16.7 (CH_3_-15e), 17.5 (CH_3_-14), 21.0 (CH_3_-1b, CH_3_-6b), 27.2 (CH_3_-13), 27.8 (CH_2_-15c), 30.2 (CH_3_-12), 31.6 (CH_2_-3), 34.1 (CH-4), 38.7 (CH_3_-2g, CH_3_-8g), 42.3 (CH-15b), 54.1 (CH-7), 60.1 (C-10), 63.9 (CH_2_-15), 71.1 (CH-1), 72.5 (CH-2), 77.7 (CH-6), 78.8 (CH-8), 85.6 (C-11), 91.9 (C-5), 110.7 (C-8b), 111.1 (C-2b), 119.6 (CH-8e), 119.9 (CH-2e), 140.5 (CH-8f), 140.6 (CH-2f), 145.6 (CH-2c), 146.3 (CH-8c), 163.9 (C-8a), 164.8 (C-2a), 165.2 (C-2d, C-8d), 171.2 (C-1a), 171.3 (C-6a), 177.6 (C-15a), 203.1 (C-9). For NMR spectra see **Supplementary Figures S12-S17**. HRESIMS *m/z* 755.3017 [M+H]^+^ (calculated for C_38_H_47_N_2_O_14_, error -0.55 ppm);MS/MS spectrum: <u>CCMSLIB00009919270</u>.

SMILES:O=C(OC([C@@]1(C2=O)[C@@]([C@](OC(C3=C([H])N(C(C([H])=C3[H])=O)C([H])([H])[H])=O)([H])C([C@@](C14OC(C([H])([H])[H])([C@@](C2([H])OC(C5=C([H])N(C(C([H])=C5[H])=O)C([H])([H])[H])=O)([H])C4([H])OC(C([H])([H])[H])=O)C([H])([H])[H])([H])C([H])([H])[H])([H])[H])([H])OC(C([H])([H])[H])=O)([H])[H])[C@@](C([H])([H])[H])([H])C([H])([H])C([H])([H])[H]. InChIKey=ATASRQYZQYFECN-QRKLMBQASA-N.

Compound **4**: (1*R*,2*S*,4*R*,5*S*,6*R*,7*S*,8*S*,10*S*)-6*β*-acetoxy-8*β*-benzoyloxy-2*α*-(5-carboxy-*N*-methyl-3-pyridoxy)-1*α*-hydroxy-15-(2-methylbutanoyloxy)-9-oxodihydro-*β*-agarofuran, Silviatine D. Amorphous white powder, 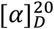 + 13 (ACN); UV (ACN) λ_max_ 196, 229, 273 nm.

^1^H NMR (CD_3_OD, 600 MHz) δ 0.95 (3H, t, *J* = 7.5 Hz, H_3_-15d), 1.15 (3H, d, *J* = 7.6 Hz, H_3_-14), 1.30 (3H, d, *J* = 7.0 Hz, H_3_-15e), 1.48 (3H, s, H_3_-13), 1.49 (3H, s, H_3_-12), 1.60 (1H, m, H-15c’’), 1.81 (1H, m, H-15c’), 1.99 (1H, dt, *J* = 15.4, 2.7, 0.9 Hz, H-3α), 2.18 (3H, s, H_3_-6b), 2.32 (1H, ddd, *J* = 15.4, 6.4, 3.6 Hz, H-3β), 2.37 (1H, m, H-4), 2.70 (1H, h, *J* = 7.0 Hz, H-15b), 3.00 (1H, d, *J* = 3.5 Hz, H-7), 3.71 (3H, s, H_3_-2g), 4.62 (1H, d, *J* = 3.8 Hz, H-1), 4.79 (1H, d, *J* = 12.3 Hz, H-15’’), 5.06 (1H, d, *J* = 12.3 Hz, H-15’), 5.45 (1H, q, *J* = 3.8, 2.7 Hz, H-2), 6.17 (1H, d, *J* = 3.5 Hz, H-8), 6.28 (1H, s, H-6), 6.58 (1H, d, *J* = 9.4 Hz, H-2e), 7.53 (2H, tt, *J* = 8.0, 1.3 Hz, H-8d, H-8f), 7.66 (1H, tt, *J* = 8.0, 1.3 Hz, H-8e), 8.04 (1H, dd, *J* = 9.4, 2.5 Hz, H-2f), 8.07 (2H, dd, *J* = 8.0, 1.3 Hz, H-8c, H-8g), 8.49 (1H, d, *J* = 2.5 Hz, H-2c); ^13^C NMR (CD_3_OD, 151 MHz) δ 11.9 (CH_3_-15d), 16.8 (CH_3_-15e), 17.7 (CH_3_-14), 21.0 (CH_3_-6b), 27.5 (CH_3_-13), 27.8 (CH_2_-15c), 30.2 (CH_3_-12), 31.7 (CH_3_-3), 34.4 (CH-4), 38.7 (CH_3_-2g), 42.4 (CH-15b), 54.4 (CH-7), 61.7 (C-10), 64.0 (CH_2_-15), 70.4 (CH-1), 75.1 (CH-2), 77.8 (CH-6), 78.8 (CH-8), 85.2 (C-11), 91.6 (C-5), 111.7 (C-2b), 119.8 (CH-2e), 129.8 (CH-8d, CH-8f), 130.6 (C-8b), 130.8 (CH-8c, CH-8g), 134.7 (CH-8e), 140.8 (CH-2f), 145.4 (CH-2c), 165.2 (C-2d), 166.3 (C-8a), 171.3 (C-6a), 178.1 (C-15a), 207.0 (C-9).. For NMR spectra see **Supplementary Figures S18-S23**. HRESIMS *m/z* 682.2850 [M+H]^+^ (calculated for C_36_H_44_NO_12_, error -1.132 ppm); MS/MS spectrum: <u>CCMSLIB00009919278</u>.

SMILES: O=C(C([H])([H])[H])O[C@@](C([C@@]([C@@]1([H])OC(C(C([H])([H])[H])([H])C([H])([H])[H])=O)([H])OC(C(C([H])=C2[H])=C([H])N(C([H])([H])[H])C2=O)=O)([H])C3(C([H])([H])[H])C([H])([H])[H])([H])[C@]4(O3)[C@@](C([H])([H])[H])([H])C([H])([H])C(OC(C(C([H])=C5[H])=C([H])N(C([H])([H])[H])C5=O)=O)([H])[C@@](OC(C([H])([H])[H])=O)([H])[C@]41C([H])([H])OC([C@@](C([H])([H])[H])([H])C([H])([H])C([H])([H])[H])=O.InChIKey=VKILZIVFMPURPQ-JIIDOJBKSA-N.

Compound **5**: (1*R*,2*S*,3*S*,4*R*,5*S*,6*R*,7*S*,8*S*,10*S*)-6*β*-acetoxy-8*β*-benzoyloxy-2*α*-(5-carboxy-*N*-methyl-3-pyridoxy)-1*α*,3*β*-dihydroxy-15-(2-methylbutanoyloxy)-9-oxodihydro-*β*-agarofuran, Silviatine E. Amorphous white powder. 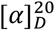 + 12 (ACN); UV (ACN) λ_max_ 200, 226, 267 nm.

^1^H NMR (CD_3_OD, 600 MHz) δ 0.96 (3H, t, *J* = 7.5 Hz, H_3_-15d), 1.16 (3H, d, *J* = 7.9 Hz, H_3_-14), 1.30 (3H, d, *J* = 7.0 Hz, H_3_-15e), 1.53 (3H, s, H_3_-13), 1.54 (3H, s, H_3_-12), 1.61 (1H, m, H-15c’’), 1.81 (1H, m, H-15c’), 2.20 (3H, s, H_3_-6b), 2.51 (1H, qt, *J* = 7.9, 1.8, 1.1 Hz, H-4), 2.71 (1H, h, *J* = 7.0 Hz, H-15b), 3.00 (1H, d, *J* = 3.5 Hz, H-7), 3.70 (3H, s, H_3_-2g), 3.92 (1H, dd, *J* = 3.3, 1.8 Hz, H-3), 4.75 (1H, d, *J* = 12.3 Hz, H-15’’), 4.88 (1H, d, *J* = 4.0 Hz, H-1), 4.99 (1H, d, *J* = 12.3 Hz, H-15’), 5.42 (1H, ddd, *J* = 4.0, 3.3, 1.1 Hz, H-2), 6.18 (1H, d, *J* = 3.5 Hz, H-8), 6.33 (1H, s, H-6), 6.58 (1H, d, *J* = 9.5 Hz, H-2e), 7.53 (2H, t, *J* = 8.0 Hz, H-8d, H-8f), 7.66 (1H, tt, *J* = 8.0, 1.2 Hz, H-8e), 8.04 (1H, dd, *J* = 9.5, 2.6 Hz, H-2f), 8.07 (2H, dd, *J* = 8.0, 1.2 Hz, H-8c, H-8g), 8.48 (1H, d, *J* = 2.6 Hz, H-2c); ^13^C NMR (CD_3_OD, 151 MHz) δ 11.9 (CH_3_-15d), 15.4 (CH_3_-14), 16.8 (CH_3_-15e), 21.0 (CH_3_-6b), 27.4 (CH_3_-13), 27.8 (CH_2_-15c), 30.1 (CH_3_-12), 38.7 (CH_3_-2g), 40.9 (CH-4), 42.4 (CH-15b), 53.2 (CH-7), 61.6 (C-10), 63.8 (CH_2_-15), 67.0 (CH-1), 73.1 (CH-3), 77.6 (CH-2), 78.3 (CH-6), 78.7 (CH-8), 87.1 (C-11), 92.9 (C-5), 111.2 (C-2b), 119.8 (CH-2e), 129.8 (CH-8d, CH-8f), 130.6 (C-8b), 130.8 (CH-8c, CH-8g), 134.8 (CH-8e), 140.8 (CH-2f), 145.6 (CH-2c), 164.6 (C-2a), 165.3 (C-2d), 166.3 (C-8a), 171.2 (C-6a), 178.0 (C-15a), 206.3 (C-9). For NMR spectra see **Supplementary Figures S24-S29**. HRESIMS *m/z* 698.2802 [M+H]^+^ (calculated for C_36_H_44_NO_13_, error -0.640 ppm); MS/MS spectrum: <u>CCMSLIB00009919275</u>.

SMILES: O=C(C([H])([H])[H])O[C@@](C([C@@]([C@@]1([H])OC(/C(C([H])([H])[H])=C([H])\C([H])([H])[H])=O)([H])OC(C([H])([H])[H])=O)([H])C2(C([H])([H])[H])C([H])([H])[H])([H])[C@]([C@@]1(C([H])([H])OC([C@@](C([H])([H])[H])([H])C([H])([H])C([H])([H])[H])=O)[C@@]3([H])OC(C([H])([H])[H])=O)(O2)[C@@](C([H])([H])[H])([H])[C@](OC(C([H])([H])[H])=O)([H])[C@]3([H])OC(C(C([H])=C4[H])=C([H])N(C([H])([H])[H])C4=O)=O.InChIKey=IEENCNNCPOLOQP-FFXLREQYSA-N.

Compound **6**: (1*R*,2*S*,4*R*,5*S*,6*R*,7*S*,8*R*,9*S*,10*S*)-6*β*-acetoxy-2*α*,8*α*-di-(5-carboxy-*N*-methyl-3-pyridoxy)-9*α*,15-di-(2-methylbutanoyloxy)-dihydro-*β*-agarofuran. Amorphous white powder, 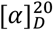 - 15 (ACN); UV (ACN) λ_max_ 194, 267 nm.

^1^H NMR (CD_3_OD, 600 MHz) δ 0.55 (3H, t, *J* = 7.4 Hz, H_3_-15d), 0.88 (3H, t, *J* = 7.5 Hz, H_3_-9d), 1.03 (3H, d, *J* = 7.0 Hz, H_3_-9e), 1.11 (3H, d, *J* = 7.7 Hz, H_3_-14), 1.13 (3H, d, *J* = 7.0 Hz, H_3_-15e), 1.24 (1H, m, H-15c’’), 1.37 (1H, m, H-9c’’), 1.46 (3H, s, H_3_-13), 1.48 (1H, m, H-15c’), 1.54 (3H, s, H_3_-12), 1.68 (1H, m, H-9c’), 1.90 (1H, d, *J* = 13.7 Hz, H-3α), 2.17 (3H, s, H_3_-6b), 2.24 (1H, q, *J* = 6.9 Hz, H-9b), 2.35 (1H, m, H-4), 2.37 (1H, m, H-15b), 2.41 (1H, m, H-3β), 2.62 (1H, d, *J* = 3.0 Hz, H-7), 3.70 (3H, s, H_3_-8g), 3.71 (3H, s, H_3_-2g), 4.37 (1H, d, *J* = 13.3 Hz, H-15’’), 4.39 (1H, d, *J* = 4.1 Hz, H-1), 5.37 (1H, td, *J* = 4.1, 2.2 Hz, H-2), 5.41 (1H, d, *J* = 13.3 Hz, H-15’), 5.64 (1H, d, *J* = 6.3 Hz, H-9), 5.66 (1H, d, *J* = 6.3 Hz, H-8), 6.56 (1H, d, *J* = 9.5 Hz, H-2e), 6.60 (1H, d, *J* = 9.4 Hz, H-8e), 6.74 (1H, s, H-6), 8.03 (1H, dd, *J* = 9.4, 2.5 Hz, H-8f), 8.07 (1H, dd, *J* = 9.5, 2.5 Hz, H-2f), 8.61 (1H, d, *J* = 2.5 Hz, H-2c), 8.89 (1H, d, *J* = 2.5 Hz, H-8c); ^13^C NMR (CD_3_OD, 151 MHz) δ 11.6 (CH_3_-15d), 12.0 (CH_3_-9d), 16.4 (CH_3_-9e), 17.5 (CH_3_-14), 17.8 (CH_3_-15e), 21.3 (CH_3_-6b), 24.9 (CH_3_-12), 27.3 (CH_2_-9c), 27.7 (CH_2_-15c), 30.5 (CH_3_-13), 32.4 (CH_2_-3), 33.9 (CH-4), 38.5 (CH_3_-2g), 38.8 (CH_3_-8g), 42.1 (CH-15b), 42.5 (CH-9b), 53.1 (C-10), 55.7 (CH-7), 63.0 (CH_2_-15), 72.8 (CH-8), 73.3 (CH-9), 75.1 (CH-2), 76.0 (CH-1), 76.9 (CH-6), 81.6 (C-11), 91.1 (C-5), 119.6 (CH-2e), 119.8 (CH-8e), 140.9 (CH-8f), 141.1 (CH-2f), 145.5 (CH-2c), 146.5 (CH-8c), 165.2 (C-2d, C-8d), 172.1 (C-6a), 176.6 (C-9a), 179.0 (C-15a). For NMR spectra see **Supplementary Figures S30-S35**. HRESIMS *m/z* 799.3649 [M+H]^+^ (calculated for C_41_H_55_N_2_O_14_, error 0.224 ppm); MS/MS spectrum: <u>CCMSLIB00009919271</u>.

SMILES: O=C(C([H])([H])[H])O[C@@](C([C@@]([C@@]1([H])OC([C@](C([H])([H])[H])([H])C([H])([H])C([H])([H])[H])=O)([H])OC(C2=C([H])N(C([H])([H])[H])C(C([H])=C2[H])=O)=O)([H])C3(C([H])([H])[H])C([H])([H])[H])([H])[C@]4(O3)[C@@](C([H])([H])[H])([H])C([H])([H])[C@](OC(C(C([H])=C5[H])=C([H])N(C([H])([H])[H])C5=O)=O)([H])[C@@](O[H])([H])[C@]41C([H])([H])OC([C@@](C([H])([H])[H])([H])C([H])([H])C([H])([H])[H])=O.InChIKey=LVFIUDAMNWXFMK-FSASPUCBSA-N.

Compound **7**: (1*R*,2*S*,4*R*,5*S*,6*R*,7*S*,8*R*,9*S*,10*S*)-1*α*,6*β*-diacetoxy-2*α*,8*α*-di-(5-carboxy-*N*-methyl-3-pyridoxy)-15-*iso*-butanoyloxy-9*α*-(2-methylbutanoyloxy)-dihydro-*β*-agarofuran. Amorphous white powder, 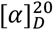 - 9 (ACN); UV (ACN) λ_max_ 195, 266 nm.

^1^H NMR (CD_3_OD, 600 MHz) δ 0.85 (3H, d, *J* = 6.9 Hz, H_3_-15d), 0.87 (3H, t, *J* = 7.5 Hz, H_3_-9d), 1.06 (3H, d, *J* = 7.1 Hz, H_3_-9e), 1.15 (3H, d, *J* = 6.9 Hz, H_3_-15c), 1.21 (3H, d, *J* = 7.7 Hz, H_3_-14), 1.31 (1H, m, H-9c’’), 1.49 (3H, s, H_3_-13), 1.53 (3H, s, H_3_-12), 1.69 (1H, m, H-9c’), 1.85 (3H, s, H_3_-1b), 1.91 (1H, d, *J* = 15.4 Hz, H-3α), 2.19 (3H, s, H_3_-6b), 2.26 (1H, m, H-9b), 2.43 (1H, p, *J* = 7.5 Hz, H-4), 2.52 (1H, ddd, *J* = 15.4, 6.9, 4.6 Hz, H-3β), 2.58 (1H, hept, *J* = 6.9 Hz, H-15b), 2.67 (1H, m, H-7), 3.68 (3H, s, H_3_-2g), 3.70 (3H, s, H_3_-8g), 4.24 (1H, d, *J* = 13.2 Hz, H-15’’), 5.46 (1H, d, *J* = 13.2 Hz, H-15’), 5.51 (1H, d, *J* = 6.3 Hz, H-9), 5.58 (1H, dd, *J* = 6.3, 3.9 Hz, H-8), 5.61 (1H, td, *J* = 4.0, 2.1 Hz, H-2), 5.70 (1H, d, *J* = 4.0 Hz, H-1), 6.56 (1H, d, *J* = 9.5 Hz, H-2e), 6.59 (1H, d, *J* = 9.4 Hz, H-8e), 6.74 (1H, d, *J* = 0.9 Hz, H-6), 8.01 (1H, dd, *J* = 9.5, 2.5 Hz, H-2f), 8.04 (1H, dd, *J* = 9.4, 2.5 Hz, H-8f), 8.52 (1H, d, *J* = 2.5 Hz, H-2c), 8.83 (1H, d, *J* = 2.5 Hz, H-8c); ^13^C NMR (CD_3_OD, 151 MHz) δ 12.1 (CH_3_-9d), 16.2 (CH_3_-9e), 17.6 (CH_3_-14), 19.3 (CH_3_-15d), 19.4 (CH_3_-15c), 21.1 (CH_3_-1b), 21.2 (CH_3_-6b), 24.8 (CH_3_-12), 26.9 (CH_2_-9c), 30.4 (CH_3_-13), 32.3 (CH_2_-3), 33.8 (CH-4), 35.1 (CH-15b), 38.6 (CH_3_-2g), 38.7 (CH_3_-8g), 42.0 (CH-9b), 52.7 (C-10), 55.3 (CH-7), 62.8 (CH_2_-15), 70.9 (CH-2), 72.2 (CH-9), 72.4 (CH-8), 76.5 (CH-6), 77.9 (CH-1), 82.0 (C-11), 91.2 (C-5), 111.0 (CH-2b), 111.9 (CH-8b), 119.7 (CH-2e), 119.8 (CH-8e), 140.7 (CH-2f), 140.8 (CH-8f), 145.7 (CH-2c), 146.4 (CH-8c), 164.8 (C-2a), 165.0 (C-8a), 165.1 (C-2d), 165.2 (C-8d), 171.3 (C-1a), 172.0 (C-6a), 175.9 (C-9a), 179.0 (C-15a). For NMR spectra see **Supplementary Figures S36-S41**. HRESIMS *m/z* 827.3598 [M+H]^+^ (calculated for C_42_H_55_N_2_O_15_, error 0.127 ppm); MS/MS spectrum: <u>CCMSLIB00009919272</u>.

SMILES: O=C(C([H])([H])[H])O[C@@]([C@]([C@@]([C@@]1([H])OC([C@](C([H])([H])[H])([H])C([H])([H])C([H])([H])[H])=O)([H])OC(C2=C([H])N(C([H])([H])[H])C(C([H])=C2[H])=O)=O)([H])C3(C([H])([H])[H])C([H])([H])[H])([H])[C@]4(O3)C(C([H])([H])[H])([H])C([H])([H])[C@@](OC(C(C([H])=C5[H])=C([H])N(C([H])([H])[H])C5=O)=O)([H])[C@@](OC(C([H])([H])[H])=O)([H])[C@]41C([H])([H])OC(C(C([H])([H])[H])([H])C([H])([H])[H])=O.InChIKey=HDNFKWOIOAMTET-UVPOIIEDSA-N.

Compound **6**: (1*R*,2*S*,4*R*,5*S*,6*R*,7*S*,8*R*,9*S*,10*S*)-1*α*,6*β*-diacetoxy-2*α*,8*α*-di-(5-carboxy-*N*-methyl-3-pyridoxy)-9*α*,15-di-(2-methylbutanoyloxy)-dihydro-*β*-agarofuran. Amorphous white powder, 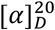 - 14 (ACN); UV (ACN) λ_max_ 204, 266 nm.

^1^H NMR (CD_3_OD, 600 MHz) δ 0.55 (3H, t, *J* = 7.4 Hz, H_3_-15d), 0.88 (3H, t, *J* = 7.4 Hz, H_3_-9d), 1.07 (3H, d, *J* = 7.1 Hz, H_3_-9e), 1.16 (3H, d, *J* = 7.0 Hz, H_3_-15e), 1.21 (3H, d, *J* = 7.6 Hz, H_3_-14), 1.25 (1H, m, H-15c’’), 1.29 (1H, m, H-9c’’), 1.49 (3H, s, H_3_-13), 1.50 (1H, m, H-15c’), 1.53 (3H, s, H_3_-12), 1.71 (1H, dqd, *J* = 13.2, 7.4, 5.5 Hz, H-9c’), 1.85 (3H, s, H_3_-1b), 1.91 (1H, dd, *J* = 15.7, 2.0 Hz, H-3α), 2.20 (3H, s, H_3_-6b), 2.24 (1H, m, H-9b), 2.39 (1H, m, H-15b), 2.44 (1H, m, H-4), 2.52 (1H, ddd, *J* = 15.7, 6.9, 4.6 Hz, H-3β), 2.66 (1H, m, H-7), 3.68 (3H, s, H_3_-2g), 3.71 (3H, s, H_3_-8g), 4.19 (1H, d, *J* = 13.2 Hz, H-15’’), 5.49 (1H, d, *J* = 13.2 Hz, H-15’), 5.52 (1H, d, *J* = 6.3 Hz, H-9), 5.58 (1H, dd, *J* = 6.3, 3.8 Hz, H-8), 5.61 (1H, td, *J* = 4.2, 2.2 Hz, H-2), 5.71 (1H, d, *J* = 4.2 Hz, H-1), 6.57 (1H, d, *J* = 9.5 Hz, H-2e), 6.60 (1H, d, *J* = 9.4 Hz, H-8e), 6.76 (1H, d, *J* = 1.0 Hz, H-6), 8.02 (1H, dd, *J* = 9.5, 2.5 Hz, H-2f), 8.03 (1H, dd, *J* = 9.4, 2.5 Hz, H-8f), 8.54 (1H, d, *J* = 2.5 Hz, H-2c), 8.88 (1H, d, *J* = 2.5 Hz, H-8c); ^13^C NMR (CD_3_OD, 151 MHz) δ 11.6 (CH_3_-15d), 12.2 (CH_3_-9d), 16.2 (CH_3_-9e), 17.6 (CH_3_-14), 17.7 (CH_3_-15e), 21.1 (CH_3_-1b), 21.2 (CH_3_-6b), 24.9 (CH_3_-12), 26.8 (CH_2_-9c), 27.7 (CH_2_-15c), 30.4 (CH_3_-13), 32.3 (CH_2_-3), 33.8 (CH-4), 38.6 (CH_3_-2g), 38.8 (CH_3_-8g), 42.0 (CH-9b), 42.1 (CH-15b), 52.7 (C-10), 55.5 (CH-7), 62.6 (CH_2_-15), 70.9 (CH-2), 72.0 (CH-9), 72.4 (CH-8), 76.6 (CH-6), 77.8 (CH-1), 82.0 (C-11), 91.1 (C-5), 111.0 (C-2b), 112.1 (C-8b), 119.8 (CH-2e, CH-8e), 140.8 (CH-2f), 140.9 (CH-8f), 145.7 (CH-2c), 146.6 (CH-8c), 164.8 (C-2a), 165.0 (C-8d), 165.2 (C-8a), 165.2 (C-2d), 171.3 (C-1a), 172.0 (C-6a), 175.9 (C-9a), 178.6 (C-15a). For NMR spectra see **Supplementary Figures S42-S47**. HRESIMS *m/z* 841.3736 [M+H]^+^ (calculated for C_43_H_57_N_2_O_15_, error -2.00 ppm); MS/MS spectrum: <u>CCMSLIB00009919274</u>.

SMILES: O=C(C([H])([H])[H])O[C@@]([C@]([C@@]([C@@]1([H])OC(/C(C([H])([H])[H])=C([H])/C([H])([H])[H])=O)([H])OC(C([H])([H])[H])=O)([H])C2(C([H])([H])[H])C([H])([H])[H])([H])[C@]([C@@]1(C([H])([H])OC(C(C([H])([H])[H])([H])C([H])([H])[H])=O)[C@@]3([H])OC(C([H])([H])[H])=O)(O2)C(C([H])([H])[H])([H])[C@](OC(C([H])([H])[H])=O)([H])[C@]3([H])OC(C(C([H])=C4[H])=C([H])N(C([H])([H])[H])C4=O)=O.InChIKey=WUSPTHMFTBWJDO-PEKKODFFSA-N.

Compound **8**: (1*R*,2*S*,4*R*,5*S*,6*R*,7*S*,8*R*,9*S*,10*S*)-1*α*,6*β*-diacetoxy-2*α*-(5-carboxy-*N*-methyl-3-pyridoxy)-9*α*,15-di-(2-methylbutanoyloxy)-8*α*-nicotinoyloxydihydro-*β*-agarofuran. Amorphous white powder, 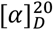 - 15 (ACN); UV (ACN) λ_max_ 194, 267 nm.

^1^H NMR (CD_3_OD, 600 MHz) δ 0.35 (3H, t, *J* = 7.5 Hz, H_3_-15d), 0.85 (3H, t, *J* = 7.5 Hz, H_3_-9d), 1.06 (3H, d, *J* = 7.2 Hz, H_3_-9e), 1.10 (1H, m, H-15c’’), 1.12 (3H, d, *J* = 7.0 Hz, H_3_-15e), 1.21 (3H, d, *J* = 7.7 Hz, H_3_-14), 1.29 (1H, m, H-9c’’), 1.35 (1H, m, H-15c’), 1.51 (3H, s, H_3_-13), 1.56 (3H, s, H_3_-12), 1.68 (1H, m, H-9c’), 1.85 (3H, s, H_3_-1b), 1.91 (1H, d, *J* = 15.3 Hz, H-3α), 2.18 (3H, s, H_3_-6b), 2.24 (1H, m, H-9b), 2.29 (1H, m, H-15b), 2.45 (1H, m, H-4), 2.52 (1H, m, H-3β), 2.75 (1H, d, *J* = 3.8 Hz, H-7), 3.68 (3H, s, H_3_-2g), 4.15 (1H, d, *J* = 13.2 Hz, H-15’’), 5.51 (1H, d, *J* = 13.2 Hz, H-15’), 5.57 (1H, d, *J* = 6.5 Hz, H-9), 5.62 (1H, td, *J* = 4.0, 2.2 Hz, H-2), 5.72 (1H, d, *J* = 4.0 Hz, H-1), 5.76 (1H, dd, *J* = 6.5, 3.8 Hz, H-8), 6.56 (1H, d, *J* = 9.5 Hz, H-2e), 6.78 (1H, s, H-6), 7.65 (1H, dd, *J* = 7.9, 5.0 Hz, H-8e), 8.02 (1H, dd, *J* = 9.5, 2.6 Hz, H-2f), 8.55 (1H, d, *J* = 2.6 Hz, H-2c), 8.57 (1H, dt, *J* = 7.9, 1.9 Hz, H-8f), 8.82 (1H, dd, *J* = 5.0, 1.9 Hz, H-8d), 9.40 (1H, d, *J* = 1.9 Hz, H-8c); ^13^C NMR (CD_3_OD, 151 MHz) δ 11.1 (CH_3_-15d), 11.8 (CH_3_-9d), 15.8 (CH_3_-9e), 17.2 (CH_3_-14, CH_3_-15e), 20.7 (CH_3_-1b, CH_3_-6b), 24.6 (CH_3_-12), 26.5 (CH_2_-9c), 27.2 (CH_2_-15c), 30.2 (CH_3_-13), 31.9 (CH_2_-3), 33.4 (CH-4), 38.2 (CH_3_-2g), 41.4 (CH-15b), 41.8 (CH-9b), 54.8 (CH-7), 62.1 (CH_2_-15), 70.5 (CH-2), 71.7 (CH-9), 72.3 (CH-8), 76.1 (CH-6), 77.5 (CH-1), 81.8 (C-11), 91.0 (C-5), 119.4 (CH-2e), 125.0 (CH-8e), 139.1 (CH-8f), 140.5 (CH-2f), 145.2 (CH-2c), 151.4 (CH-8c), 154.2 (CH-8d), 164.9 (C-2d), 171.0 (C-1a), 171.2 (C-6a), 175.5 (C-9a), 178.6 (C-15a). For NMR spectra see **Supplementary Figures S48-S52**. HRESIMS *m/z* 811.3668 [M+H]^+^ (calculated for C_42_H_53_N_2_O_11_, error 2.58 ppm); MS/MS spectrum: <u>CCMSLIB00009919277</u>.

SMILES: O=C1[C@](OC(C2=C([H])C([H])=C([H])C([H])=C2[H])=O)([H])[C@](C3(C([H])([H])[H])C([H])([H])[H])([H])[C@](OC(C([H])([H])[H])=O)([H])[C@]4(O3)[C@@](C([H])([H])[H])([H])C([H])([H])[C@@](OC(C(C([H])=C5[H])=C([H])N(C([H])([H])[H])C5=O)=O)([H])[C@@](O[H])([H])[C@]41C([H])([H])OC([C@@](C([H])([H])[H])([H])C([H])([H])C([H])([H])[H])=O. InChIKey=TXTJGSCAWSRSFC-GCFOXSEASA-N.

Compound **9**: (1*R*,2*S*,4*R*,5*S*,6*R*,7*S*,8*R*,9*S*,10*S*)-1*α*,6*β*-diacetoxy-9*α*-*iso*-butanoyloxy-2*α*,8*α*-di-(5-carboxy-*N*-methyl-3-pyridoxy)-15-methylbutanoyloxydihydro-*β*-agarofuran. Amorphous white powder, 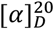 - 6 (ACN); UV (ACN) λ_max_ 206, 269 nm.

^1^H NMR (CD_3_OD, 600 MHz) δ 0.54 (3H, t, *J* = 7.4 Hz, H_3_-15d), 1.07 (3H, d, *J* = 7.0 Hz, H_3_-9d), 1.09 (3H, d, *J* = 7.1 Hz, H_3_-9c), 1.15 (3H, d, *J* = 7.0 Hz, H_3_-15e), 1.21 (3H, d, *J* = 7.6 Hz, H_3_-14), 1.25 (1H, m, H-15c’’), 1.47 (1H, m, H-15c’), 1.49 (3H, s, H_3_-13), 1.52 (3H, s, H_3_-12), 1.85 (3H, s, H_3_-1b), 1.92 (1H, m, H-3α), 2.21 (3H, d, *J* = 1.1 Hz, H_3_-6b), 2.38 (1H, m, H-15b), 2.44 (1H, m, H-4), 2.45 (1H, m, H-9b), 2.52 (1H, ddd, *J* = 15.5, 6.9, 4.6 Hz, H-3β), 2.67 (1H, dd, *J* = 3.8, 0.9 Hz, H-7), 3.68 (3H, s, H_3_-2g), 3.70 (3H, s, H_3_-8g), 4.18 (1H, d, *J* = 13.1 Hz, H-15’’), 5.49 (1H, d, *J* = 13.1 Hz, H-15’), 5.52 (1H, d, *J* = 6.3 Hz, H-9), 5.56 (1H, dd, *J* = 6.3, 3.8 Hz, H-8), 5.61 (1H, dt, *J* = 4.0, 2.1 Hz, H-2), 5.70 (1H, d, *J* = 4.0 Hz, H-1), 6.56 (1H, d, *J* = 9.5 Hz, H-2e), 6.60 (1H, d, *J* = 9.4 Hz, H-8e), 6.77 (1H, d, *J* = 0.9 Hz, H-6), 8.02 (1H, dd, *J* = 9.5, 2.6 Hz, H-2f), 8.04 (1H, dd, *J* = 9.4, 2.5 Hz, H-8f), 8.54 (1H, d, *J* = 2.6 Hz, H-2c), 8.88 (1H, d, *J* = 2.5 Hz, H-8c); ^13^C NMR (CD_3_OD, 151 MHz) δ 11.6 (CH_3_-15d), 17.5 (CH_3_-14), 17.7 (CH_3_-15e), 18.8 (CH_3_-9d), 19.0 (CH_3_-9c), 21.0 (CH_3_-1b), 21.2 (CH_3_-6b), 24.9 (CH_3_-12), 27.7 (CH_2_-15c), 30.4 (CH_3_-13), 32.3 (CH_2_-3), 33.8 (CH-4), 35.3 (CH-9b), 38.6 (CH_3_-2g), 38.8 (CH_3_-8g), 42.1 (CH-15b), 52.7 (C-10), 55.5 (CH-7), 62.6 (CH_2_-15), 70.9 (CH-2), 71.9 (CH-9), 72.4 (CH-8), 76.6 (CH-6), 77.8 (CH-1), 81.9 (C-11), 91.1 (C-5), 111.0 (C-2b), 112.0 (C-8b), 119.8 (CH-2e, CH-8e), 140.8 (CH-2f), 140.9 (CH-8f), 145.7 (CH-2c), 146.6 (CH-8c), 164.8 (C-2d, C-8d), 171.3 (C-1a), 172.0 (C-6a), 176.3 (C-9a), 178.7 (C-15a). For NMR spectra see **Supplementary Figures S53-S58**. HRESIMS *m/z* 827.3595 [M+H]^+^ (calculated for C_42_H_55_N_2_O_15_, error -0.16 ppm); MS/MS spectrum: <u>CCMSLIB00009919279</u>.

SMILES: O=C(C([H])([H])[H])O[C@@](C([C@@]([C@@]1([H])OC(C(C([H])([H])[H])([H])C([H])([H])[H])=O)([H])OC(C(C([H])=C2[H])=C([H])N(C([H])([H])[H])C2=O)=O)([H])C3(C([H])([H])[H])C([H])([H])[H])([H])[C@]4(O3)[C@@](C([H])([H])[H])([H])C([H])([H]). InChIKey=KXKFNEWNZKWNFD-DCBBRINESA-N.

Compound **10**: (1*R*,2*S*,4*R*,5*S*,6*R*,7*S*,8*R*,9*S*,10*S*)-6*β*-diacetoxy-9*α*-*iso*-butanoyloxy-2*α*,8*α*-di-(5-carboxy-*N*-methyl-3-pyridoxy)-1*α*-hydroxy-15-methylbutanoyloxydihydro-*β*-agarofuran. Amorphous white powder, 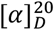 - 30 (ACN); UV (ACN) λ_max_ 195, 266 nm.

^1^H NMR (CD_3_OD, 600 MHz) δ 0.54 (3H, t, *J* = 7.5 Hz, H_3_-15d), 1.07 (3H, d, *J* = 7.0 Hz, H_3_-9d), 1.09 (3H, d, *J* = 7.0 Hz, H_3_-9c), 1.11 (3H, d, *J* = 7.9 Hz, H_3_-14), 1.13 (3H, d, *J* = 7.3 Hz, H_3_-15e), 1.22 (1H, m, H-15c’), 1.46 (1H, m, H-15c’’), 1.46 (3H, s, H_3_-13), 1.54 (3H, s, H_3_-12), 1.90 (1H, d, *J* = 15.3 Hz, H-3α), 2.18 (3H, s, H_3_-6b), 2.36 (2H, m, H-4, H-15b), 2.40 (1H, m, H-3β), 2.43 (1H, hept, *J* = 7.0 Hz, H-9b), 2.63 (1H, d, *J* = 3.9 Hz, H-7), 3.70 (6H, 2xs, H_3_-2g, H_3_-8g), 4.36 (1H, d, *J* = 13.2 Hz, H-15’’), 4.38 (1H, d, *J* = 4.1 Hz, H-1), 5.37 (1H, m, H-2), 5.41 (1H, d, *J* = 13.2 Hz, H-15’), 5.62 (1H, dd, *J* = 6.1, 3.9 Hz, H-8), 5.65 (1H, d, *J* = 6.1 Hz, H-9), 6.56 (1H, d, *J* = 9.4 Hz, H-2e), 6.60 (1H, d, *J* = 9.5 Hz, H-8e), 6.75 (1H, s, H-6), 8.05 (1H, dd, *J* = 9.5, 2.5 Hz, H-8f), 8.07 (1H, dd, *J* = 9.4, 2.5 Hz, H-2f), 8.61 (1H, d, *J* = 2.5 Hz, H-2c), 8.90 (1H, d, *J* = 2.5 Hz, H-8c); ^13^C NMR (CD_3_OD, 151 MHz) δ 11.3 (CH_3_-15d), 17.2 (CH_3_-14), 17.4 (CH_3_-15e), 18.6 (CH_3_-9c, CH_3_-9d), 20.9 (CH_3_-6b), 24.6 (CH_3_-12), 27.5 (CH_2_-15c), 30.2 (CH_3_-13), 32.2 (CH_2_-3), 33.5 (CH-4), 35.3 (CH-9b), 38.6 (CH_3_-2g), 38.3 (CH_3_-2g, CH_3_-8g), 41.9 (CH-15b), 55.4 (CH-7), 67.9 (CH_2_-15), 72.6 (CH-8), 73.0 (CH-9), 74.7 (CH-2), 75.6 (CH-1), 76.7 (CH-6), 81.2 (C-11), 90.8 (C-5), 111.5 (C-2b), 112.0 (C-8b), 119.2 (CH-2e), 119.4 (CH-8e), 140.7 (CH-2f, CH-8f), 145.2 (CH-2c), 146.3 (CH-8c), 165.0 (C-2d, C-8d), 171.7 (C-6a), 176.7 (C-9a), 178.7 (C-15a).. For NMR spectra see **Supplementary Figures S59-S63**. HRESIMS *m/z* 785.3511 [M+H]^+^ (calculated for C_40_H_53_N_2_O_14_, error -2.55 ppm);MS/MS spectrum: <u>CCMSLIB00009919269</u>.

SMILES: O=C(C([H])([H])[H])O[C@@](C([C@@]([C@@]1([H])OC([C@](C([H])([H])[H])([H])C([H])([H])C([H])([H])[H])=O)([H])OC(C2=C([H])N(C([H])([H])[H])C(C([H])=C2[H])=O)=O)([H])C3(C([H])([H])[H])C([H])([H])[H])([H])[C@]4(O3)[C@@](C([H])([H])[H])([H])C([H])([H])C(OC(C(C([H])=C5[H])=C([H])N(C([H])([H])[H])C5=O)=O)([H])[C@@](OC(C([H])([H])[H])=O)([H])[C@]41C([H])([H])OC([C@@](C([H])([H])[H])([H])C([H])([H])C([H])([H])[H])=O. InChIKey=IJMXFBHJNXUVLI-VPQZVAQISA-N.

Compound **11**: (1*R*,2*S*,3*S*,4*R*,5*S*,6*R*,7*S*,8*R*,9*S*,10*S*)-1*α*,3*β*,6*β*,8*α*-tetraacetoxy-15-*iso*-butanoyloxy-2*α*-(5-carboxy-*N*-methyl-3-pyridoxy)-9*α*-tigloyloxydihydro-*β*-agarofuran. Amorphous white powder, 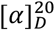 - 2 (ACN); UV (ACN) λ_max_ 207, 268 nm.

^1^H NMR (CD_3_OD, 600 MHz) δ 1.21 (3H, d, *J* = 7.9 Hz, H_3_-14), 1.24 (3H, d, *J* = 6.9 Hz, H_3_-15d), 1.26 (3H, d, *J* = 6.9 Hz, H_3_-15c), 1.44 (3H, s, H_3_-13), 1.53 (3H, s, H_3_-12), 1.75 (3H, s, H_3_-1b), 1.77 (3H, p, *J* = 1.3 Hz, H_3_-9e), 1.78 (3H, dq, *J* = 6.9, 1.3 Hz, H_3_-9d), 2.11 (3H, s, H_3_-8b), 2.12 (3H, s, H_3_-6b), 2.12 (3H, s, H_3_-3b), 2.50 (1H, dd, *J* = 3.8, 1.0 Hz, H-7), 2.56 (1H, m, H-4), 2.90 (1H, hept, *J* = 6.9 Hz, H-15b), 3.71 (3H, d, *J* = 1.7 Hz, H_3_-2g), 4.28 (1H, d, *J* = 13.2 Hz, H-15’’), 4.87 (1H, overlapped, H-3), 5.36 (1H, d, *J* = 13.2 Hz, H-15’), 5.49 (1H, d, *J* = 6.5 Hz, H-9), 5.52 (2H, m, H-2, H-8), 5.91 (1H, d, *J* = 4.2 Hz, H-1), 6.58 (1H, d, *J* = 1.1 Hz, H-6), 6.58 (1H, d, *J* = 9.5 Hz, H-2e), 6.83 (1H, qq, *J* = 6.9, 1.3 Hz, H-9c), 8.00 (1H, dd, *J* = 9.5, 2.6 Hz, H-2f), 8.55 (2H, d, *J* = 2.6 Hz, H-2c); ^13^C NMR (CD_3_OD, 151 MHz) δ 11.6 (CH_3_-9e), 14.1 (CH_3_-9d), 14.9 (CH_3_-14), 19.3 (CH_3_-15c, CH_3_-15d), 20.2 (CH_3_-1b), 20.7 (CH_3_-3b, CH_3_-6b,, CH_3_-8b), 24.6 (CH_3_-12), 30.2 (CH_3_-13), 34.8 (CH-15b), 38.0 (CH-4), 38.3 (CH_3_-2g), 52.0 (C-10), 53.3 (CH-7), 61.9 (CH_2_-15), 70.5 (CH-8), 71.3 (CH-2), 71.9 (CH-9), 74.9 (CH-, 75.8 (CH-3), 76.5 (CH-6), 82.5 (C-11), 90.3 (C-5), 109.9 (C-2b), 119.6 (CH-2e), 128.9 (C-9b), 140.0 (CH-9c), 140.4 (CH-2f), 145.8 (CH-2c), 163.9 (C-2a), 165.0 (C-2d), 167.0 (C-9a), 170.8 (C-1a), 171.1 (C-6a), 171.3 (C-3a), 171.5 (C-8a), 178.9 (C-15a). For NMR spectra see **Supplementary Figures S64-S68**. HRESIMS *m/z* 790.3289 [M+H]^+^ (calculated for C_39_H_52_NO_16_, error 1.16 ppm); MS/MS spectrum: <u>CCMSLIB00009919273</u>.

SMILES: O=C1[C@@](OC(C2=C([H])C([H])=C([H])C([H])=C2[H])=O)([H])[C@](C3(C([H])([H])[H])C([H])([H])[H])([H])C(OC(C([H])([H])[H])=O)([H])[C@]([C@](C([H])([H])OC([C@@](C([H])([H])[H])([H])C([H])([H])C([H])([H])[H])=O)1C4([H])O[H])(O3)[C@@](C([H])([H])[H])([H])[C@](O[H])([H])[C@]4([H])OC(C(C([H])=C5[H])=C([H])N(C([H])([H])[H])C5=O)=O. InChIKey=ZSYJSJVZJMUAMB-VWMXXGJYSA-N.

Compound **12**: (1*R*,2*S*,3*S*,4*R*,5*S*,6*R*,7*S*,8*R*,9*S*,10*S*)-1*α*,3*β*,6*β*,8*α*-tetraacetoxy-2*α*-(5-carboxy-*N*-methyl-3-pyridoxy)-15-(2-methylbutanoyloxy)-9*α*-tigloyloxydihydro-*β*-agarofuran. Amorphous white powder, 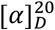 - 15 (ACN); UV (ACN) λ_max_ 194, 270 nm.

^1^H NMR (CD_3_OD, 600 MHz) δ 0.97 (3H, t, *J* = 7.4 Hz, H_3_-15d), 1.20 (3H, d, *J* = 7.9 Hz, H_3_-14), 1.25 (3H, d, *J* = 7.0 Hz, H_3_-15e), 1.43 (3H, s, H_3_-13), 1.53 (3H, s, H_3_-12), 1.58 (1H, ddd, *J* = 13.8, 7.5, 6.4 Hz, H-15c’’), 1.75 (3H, s, H_3_-1b), 1.78 (6H, m, H_3_-9d, H_3_-9e), 1.80 (1H, m, H-15c’), 2.12 (3H, s, H_3_-6b), 2.12 (3H, s, H_3_-3b), 2.13 (3H, s, H_3_-8b), 2.50 (1H, dd, *J* = 3.7, 1.0 Hz, H-7), 2.56 (1H, qt, *J* = 8.0, 1.0 Hz, H-4), 2.74 (1H, h, *J* = 7.0 Hz, H-15b), 3.71 (3H, s, H_3_-2g), 4.22 (1H, d, *J* = 13.3 Hz, H-15’’), 4.87 (1H, overlapped, H-3), 5.44 (1H, d, *J* = 13.3 Hz, H-15’), 5.49 (1H, d, *J* = 6.6 Hz, H-9), 5.53 (2H, m, H-2, H-8), 5.92 (1H, d, *J* = 4.3 Hz, H-1), 6.55 (1H, d, *J* = 1.0 Hz, H-6), 6.58 (1H, d, *J* = 9.5 Hz, H-2e), 6.83 (1H, qq, *J* = 7.3, 1.6 Hz, H-9c), 8.02 (1H, dd, *J* = 9.5, 2.5 Hz, H-2f), 8.58 (1H, d, *J* = 2.5 Hz, H-2c); ^13^C NMR (CD_3_OD, 151 MHz) δ 12.1 (CH_3_-9e, CH_3_-15d), 14.4 (CH_3_-9d), 15.2 (CH_3_-14), 17.4 (CH_3_-15e), 20.5 (CH_3_-1b), 21.1 (CH_3_-6b), 21.2 (CH_3_-3b), 21.4 (CH_3_-8b), 24.9 (CH_3_-12), 27.9 (CH_2_-15c), 30.6 (CH_3_-13), 38.2 (CH-4), 38.6 (CH_3_-2g), 42.3 (CH-15b), 52.1 (C-10), 53.6 (CH-7), 62.1 (CH_2_-15), 70.8 (CH-8), 71.6 (CH-2), 72.1 (CH-9), 75.2 (CH-1), 76.1 (CH-3), 76.9 (CH-6), 82.7 (C-11), 90.6 (C-5), 110.3 (C-2b), 119.9 (CH-2e), 129.2 (C-9b), 140.2 (CH-9c), 140.7 (CH-2f), 146.1 (CH-2c), 164.1 (C-2a), 165.3 (C-2d), 167.3 (C-9a), 171.1 (C-1a), 171.2 (C-6a), 171.5 (C-3a), 171.7 (C-8a), 178.9 (C-15a). For NMR spectra see **Supplementary Figures S69-S74**. HRESIMS *m/z* 804.3445 [M+H]^+^ (calculated for C_40_H_54_NO_16_, error 1.03 ppm); MS/MS spectrum: <u>CCMSLIB00009919276</u>.

SMILES: O=C(C([H])([H])[H])O[C@@]([C@]([C@@]([C@@]1([H])OC([C@](C([H])([H])[H])([H])C([H])([H])C([H])([H])[H])=O)([H])OC(C2=C([H])N=C([H])C([H])=C2[H])=O)([H])C3(C([H])([H])[H])C([H])([H])[H])([H])[C@]4(O3)C(C([H])([H])[H])([H])C([H])([H])C(OC(C(C([H])=C5[H])=C([H])N(C([H])([H])[H])C5=O)=O)([H])[C@@](O/C(C([H])([H])[H])=C([H])\[H])([H])[C@]41C([H])([H])OC([C@@](C([H])([H])[H])([H])C([H])([H])C([H])([H])[H])=O. InChIKey=QQDABXSQXJXVOM-DEEWNFLNSA-N.

#### 2.5.2 Electronic Circular Dichroism (ECD) Calculations

The absolute configuration of all compounds was assigned according to the comparison of the calculated and experimental ECD. Based on their relative configuration proposed by NMR 2D ROESY experiments, the structures were employed for the random conformational search using MMFF94s force field by Spartan Student v7 (Wavefunction, Irvine, CA, USA). From the results, the 20 isomers with lower energy were subjected to further successive PM3 and B3LYP/6-31G(d,p) optimizations in Gaussian 16 software (© 2015-2022, Gaussian Inc., Wallingford, CT, USA) using the CPCM model in acetonitrile (Nugroho and Morita, 2014; Mándi and Kurtán, 2019). All optimized conformers in each step were checked to avoid imaginary frequencies. After a cut-off of 4 kcal/mol in energy, conformers were submitted to Gaussian16 software for ECD calculations, using TD-DFT B3LYP/def2svp as a basis set with the CPCM model in acetonitrile. The calculated ECD spectrum was generated in SpecVis1.71 software (Berlin, Germany) based on the Boltzmann weighting average. Results are shown in **Supplementary Figure S75**). The ECD calculations on Gaussian 16 (© 2015–2022, Gaussian inc.) were performed at the University of Geneva on the HPC <u>Baobab cluster</u>.

## 3 Results and Discussion

The prioritization of a particular natural extract for the search for NPs with novel structural characteristics is linked to the availability of literature reports and the dereplication results. The first one allows visualizing the extension of the knowledge for a particular taxon and deciding if it is worthy of further studies. The second one will help putatively highlight a particular extract’s composition at the analytical level. A combination of both aspects could indicate where to focus the isolation efforts.

<u>*Inventa*</u> automatically calculates multiple scores that estimate each extract’s chemical novelty from previous literature reports and MS-based metabolomics analysis. The scores consider the compounds reported in the literature for the taxon, the occurrence of specific features in the mass spectrometry profiles of all extracts, and the MS^2^ annotations obtained with a combination of advanced computational annotation methods. *Inventa*’s scores are related to four different components. The individual calculations and the user’s tunable parameters are described below.

### 3.1 The conception of the priority score

*Inventa* focuses on the discovery of novel NPs in a series of extracts by giving a rank of prioritization for the extracts before being subject to phytochemical studies. Additional information on potentially putative new compounds within such extract is available for precise localization of the features of interest for targeted isolation.

*Inventa* takes a Feature Based Molecular Network (FBMN) job as minimum input. This workflow is preferred over the classical MN since it incorporates mass spectrometry (MS^1^ and MS^2^), and semi quantitative chromatographic information (retention time, intensity/area) specific for each feature (Nothias et al., 2020). FBMN was considered since it became a widely used workflow for data comparison, spectral space visualization, and automated annotation against experimental databases. From these results, *Inventa* will use as input the feature table, the annotation results, and the taxonomic information of the extracts. The specificity for each feature will be assigned according to the aligned feature table (generated initially by MZmine). Other software can be used, if compatible with GNPS, the user can recover the table from the MN results. Their annotation status is based on the GNPS annotation results. To guarantee a minimum quality of the putative identities, the GNPS annotations are automatically cleaned and filtered (cosine, error in ppm, number of shared peaks, polarity, etc.) before the calculations (https://github.com/lfnothias/gnps_postprocessing). Additional feature dereplications results using *in-silico* databases and reponderation strategies to improve the putative annotation (Allard et al., 2016, 2017; Dührkop et al., 2019; Rutz et al., 2019) can be included in the pipeline. If so, the annotation status of the features considered them as well. Finally, the metadata table should indicate the characteristics of the extracts, like the filename and the species, genus, and family, for searching reports in the literature.

If the data treatment is performed with a version of <u>MZmine</u> supporting IIN (custom 2.53 version or MZmine 3), the user can leverage the grouping of multiple ion forms identified for a given molecule and reduce the total number of features. The species generated from the same molecules (adducts, in source fragments, etc.) are collapsed into a single feature group (ion identity networks, IIN) through an MS^1^ feature chromatographic shape correlation. *Inventa* will perform the calculations related to FC based on the new MS^1^-based group features and MS^2^ spectral similarity cosine comparison (Schmid et al., 2021). The area/height used will correspond to the maximum value found within each IIN (most representative ion-adduct). Using IIN will necessarily facilitate the extract selection by deconvolving the mass spectrometry data into several molecules present in each extract.

*Inventa* considers the information at two levels to rank the extracts: individual features within each extract and the extract itself by considering the overall pool of MS^2^ data. The specificity and annotations (structure, molecular formula, and chemical classes) are pondered at the features level to express each extract’s measurable unknown chemical richness. At the extract level, each extract’s available spectral space is compared to each other to spot dissimilarities using a dissimilarity matrix based on the MEMO vectors (Gaudry et al., 2022). A combination of both levels and the literature reports for the taxon will highlight the extracts with an unknown specialized metabolism.

The priority score comes from the addition of four individual components: Feature component (FC), Literature component (LC), Class component (CC), and Similarity component (SC) (**Figure 1**). Each component is normalized from 0 to 1. *Inventa* implements a modulating factor (*w*_*n*_ in **Figure 1**) to give the appropriate weight to each component according to the type of study and the user’s preferences. A full glossary with terms and default values is available in the **Supplementary Table S1**.

**Figure 1.**
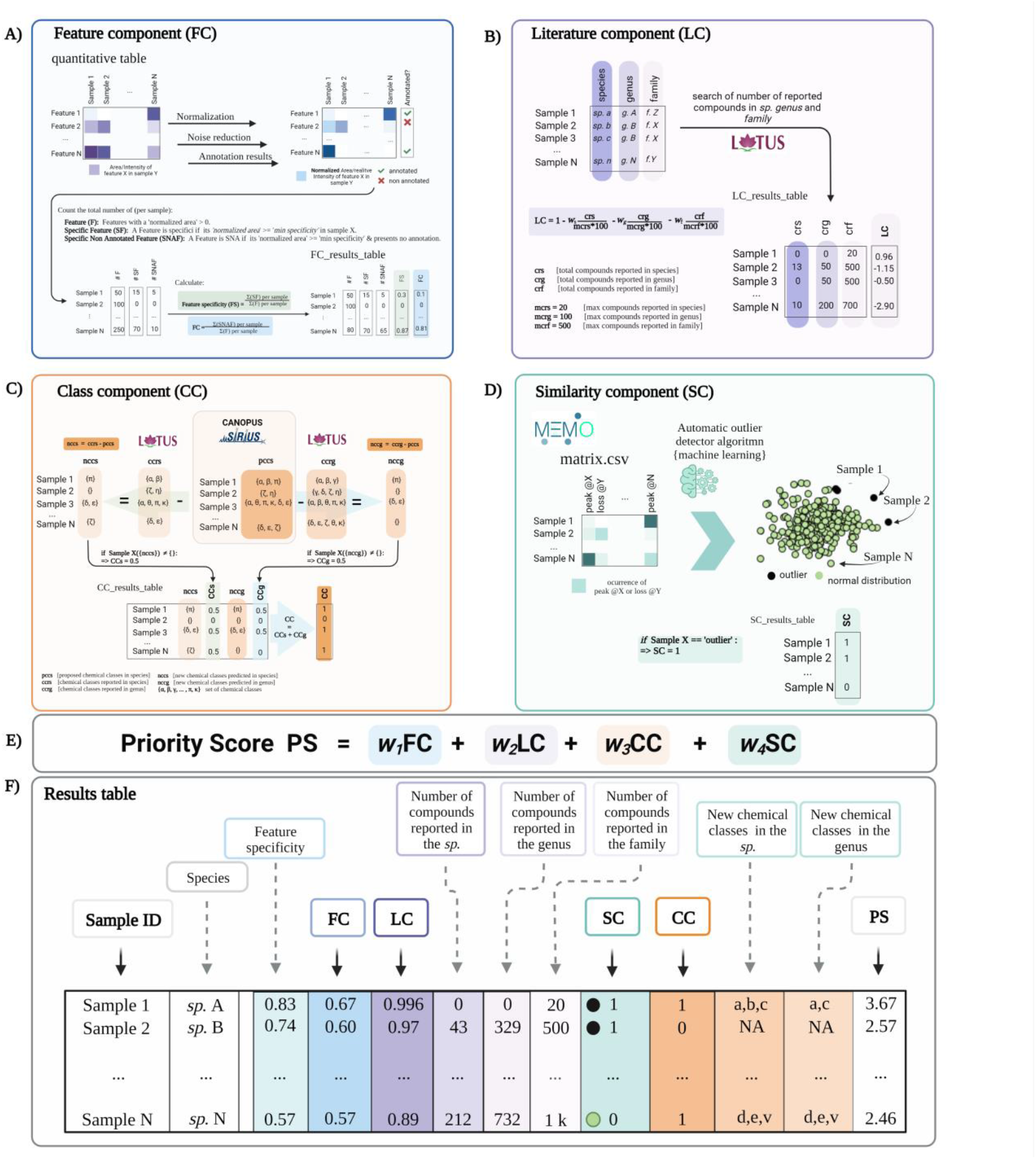
A conceptual overview of *Inventa*’s priority score and its components. **A) Feature Component (FC)**: is a ratio of the number of specific and unannotated features over the total number of features by extract. **B) Literature Component (LC)**: is a score based on the number of compounds reported in the literature for the taxon of a given extract. It is independent of the spectral data. **C) Class Component (CC)**: indicates if an unreported chemical class is detected in each extract compared to those reported in the species and the genus. **D) Similarity Component (SC)**: is a complementary score that compares extracts based on their general MS2 spectral information independently from the feature alignment used in FC, using MEMO. **E)** The **Priority Score (PS)** is the addition of the four components. A modulating factor (*w*^*n*^) gives each component a relative weight according to the user’s preferences. The higher the value, the higher the rank of the extract. **F) Results Table** is a resume of individual calculation components and results.

The ***Feature Component*** (**FC, Figure 1.A**) is the ratio for the number of specific and unannotated features over the total number of features of each extract. For example, an FC of ‘0.6’ implies that 60% of the total features in each extract are specific within the set and do not present structural annotations. For the calculation of this ratio, the aligned feature table is normalized row-wise (each row corresponding to a feature). Based on this normalized table, a feature is considered specific in each extract, compared to the whole extract set, if its normalized area is higher than the *minimum specificity* value. By default, a feature is considered specific if at least 90% of the normalized peak area is detected in each extract (*minimum specificity* set to 0.90; this parameter can be modified by the user). Then, the annotation status (annotated or unannotated) is checked based on the dereplication results used as input. Finally, the total number of specific unannotated features in each extract is calculated and divided by the total number of detected features in the same extract. The evaluation of the specificity of the features (without information on their annotation status) a given extract within the set can be done based on the ‘Feature Specificity’ (FS) value (is computed similarly to FC without considering the annotations). **Supplementary Figure S76** shows the detailed calculations performed for obtaining the FC score.

Usually, collections of natural extracts include extracts of the same species but with distinct characteristics, such as organs (flowers, leaves, stems, fruits), collection sites, culture media (in the case of micro-organisms) or extraction solvents, among others. As explained above, the FS and FC consider a feature specific if its relative intensity is higher than the ‘*minimum specificity*’ defined by the user. When multiple extracts with the same species are present, even if a feature is specific at the species level, its relative intensity may be spread over its various extracts. Consequently, that feature will not be considered specific and will be ignored in the calculations. To address this limitation, the user can define the maximum occurrence of the species allowing the script to consider a feature as ‘specific’ based on a shared specificity within multiple extracts (detailed calculations are shown in **Supplementary Figure S77**). **Supplementary Figure S78** shows what happens on FS and FC calculation when a plant within a set is analyzed based on four independent organs (one extract per organ). For example, for Catha edulis four extracts corresponding to its aerial parts, leaves, roots, and stems, were profiled. If the ‘*maximum occurrence* (N)’ is 1, many features will be not considered specific because they are shared between the plant parts. If for the data set the ‘*maximum occurrence* (N)’ is set to 4, the number of specific features increased. This immediately raised the FS and FC in general, and the common tissue parts (aerial parts, leaves, and stems) gained up to 4 fold the FC’s original value.

The ***Literature Component*** (**LC**, See **Figure 1.B**) is a score based on the number of compounds reported in the literature for the taxon of a given extract. It is independent of the spectral data. For example, an LC value of 1 indicates no reported compounds for the considered taxon. From this initial value (‘1’), fractions (ratio of reported compounds over the user-defined maximum value of reported compounds) are subtracted. The first fraction is related to compounds found in the species, the second one to those found in the genus, and the third one in the family (see the formula in **Figure 1.B**). By default, the weight of each fraction is equal; it can be pondered by the user depending on the needs. For the calculation of this value, the clean taxonomic information (based on the <u>Open Tree of Life</u>, OTL) is retrieved from the metadata table and used to query the NPs occurrences reported in the <u>LOTUS</u> initiative (Rutz et al., 2022). The LC represents a rough estimation of the literature knowledge on a given extract in terms of reported compounds. It does not replace an extensive literature search but allows to rapidly visualize the species that have been heavily studied or not in a set. **Supplementary Figure S79** shows the detailed calculations performed for the LC score.

The first evaluation of both FC and LC components provides an excellent way to highlight extracts containing an important proportion of specific unannotated features that have not been the topic of extensive phytochemical studies. Regarding this calculation, it is essential to recall that the reported chemistry is not specified to a plant-organ level in the databases. Thus, no part-specific relation can be constructed relative to the tissue involved. For example, a specific plant part extract could have a high FC due to a specific profile with no annotation and a bad LC score because the taxon presents a high number of the reported compounds, not necessarily in the same organ. Reports in the genus and family are considered for prioritizing a particular lack of annotation but belonging to an extensive phytochemically studied genus or family.

The ***Class Component*** (**CC, Figure 1.C**) indicates if an unreported chemical class is detected in each extract compared to those reported in the species and the genus. A CC value of 1 implies that the chemical class is new to both the species (CCs 0.5) and the genus (CCg 0.5). The CC calculation is derived from the CANOPUS sub-tool integrated in SIRIUS and that is used to propose a chemical class directly from the MS^2^ spectral fingerprint of the features without the need for a formal structural annotation (Dührkop et al., 2019, 2020). The chemical taxonomy classification is based on the standardized NPClassifier chemical taxonomy (Kim et al., 2021). This chemical class annotation provides a partial but systematic annotation for the detected features, even for novel molecules. The NPClassifier chemical classes have unique standardized names that can be compared computationally as text strings. *Inventa* compares the predicted chemical classes in each extract to those reported in the species in LOTUS, which also uses the NPClassifier ontology. The comparison is performed by string set subtraction. If one or several unreported classes were annotated in the extract compared to the literature, the CC value at the species level (CCs) is set to 0.5. The same calculation is performed for comparing the reports at the genus level, and similarly, a value of CCg (value at the genus level) is set to 0.5 if at least one unreported class is found. Both values are added to give the final CC value. To avoid inconsistent proposed chemical classes throughout a given extract, a ‘*minimum recurrence filter*’ is used to verify that at least more than ‘*n*’ features are annotated with a given NPClassifier class (the user can modify this value). **Supplementary Figure S80** shows the detailed calculations performed for obtaining the CC score.

The ***Similarity Component*** (**SC, Figure 1.D**) is a complementary score that compares extracts based on their general MS^2^ spectral information independently from the feature alignment used in FC, using MEMO (Gaudry et al., 2022). This metric generates a matrix containing all the MS^2^ information in the form of peaks and neutral losses without annotations. The matrix is mined through multiple outlier detection machine learning algorithms to highlight spectrally dissimilar extracts (outliers). An SC value of ‘1’ implies the extract is classified as an *outlier* within the extract set studied. This score highlights spectrally dissimilar extracts. Such information may be linked to spectral fingerprints that are likely related to singular chemistry. This score can be compared to the FC, and since it is independent of alignment and annotation might help to evaluate the specificity of the extract from an orthogonal perspective. For its calculation, the dissimilarity matrix created is subjected to three different unsupervised algorithms: Local outlier factor (LOF, distance-based method) (Breunig et al., 2000), One-Class Support Vector Machine (OCSVM, domain-based method) (Wang et al., 2004), and Isolation Forest (IF, isolation-based method) (Liu et al., 2008). In general, IF and OCSVM are reported to achieve the best outlier detection results for large data sets. LOF has an average performance for different multivariate set sizes. They all stand out for their robustness when noise is introduced into the dataset (Domingues et al., 2018). If an extract is considered an outlier in at least one algorithm, an SC value of ‘1’ is given; otherwise, ‘0’. **Supplementary Figure S81** shows the detailed calculations performed for obtaining the SC score.

### 3.2 Combination of the results and formatting

To globally visualize the various scores and additional information produced for each extract in the set, *Inventa* combines and organizes the results as an interactive table (Gratzl et al., 2013; Furmanova et al., 2020) with the same format as shown in **Figure 1.E/F**. It can be sorted by the priority score (final score) or by each component, depending on the user’s needs. This interactive table allows a straightforward evaluation of the scoring parameters based on modifications of the parameters that the user can tune according to the type of study (see Glossary in Supporting Information, #userdefined tag).

### 3.3 Implementation of *Inventa* to prioritize extracts in a collection of plants from the Celastraceae family

According to LOTUS (Rutz et al., 2022) and the <u>Dictionary of Natural Products</u>, 4,800 unique NPs have been reported for the Celastraceae family (0.98% of the total entries for the Archaeplastida), involving around 38 genera and 168 species. These NPs present 130 different chemical classes (NPClassifier (Kim et al., 2021)), covering approximately 20% of the known chemical classes of the Archaeplastida.

The <u>set</u> of plants from the Celastraceae family considered in this study consists of 36 species and 14 different genera. Several plant parts were considered, depending on the availability, yielding 76 extracts in total. To improve the detection of medium polarity specialized metabolites, only ethyl acetate extracts were prepared. Extensive metabolite profiling of all extracts was performed by UHPLC-HRMS/MS operating in Data Dependent Acquisition mode. A careful comparison of the Base Peak Intensity (BPI) traces for both positive and negative ionization modes with the semiquantitative Charged Aerosol Detector trace (CAD) indicated that the positive mode was the most representative of the composition of the extracts. Thus, for this study, only the positive ionization data was considered. The data were processed with MZmine3 ((Pluskal et al., 2010), producing a list of 16,139 features. After the application of the MS^1^ Ion identity feature grouping, these features were grouped into 14,554 IIN, where 3,610 features were identified with their adducts. The resulting tables and spectral data were uploaded to the GNPS website to generate a Feature Based Molecular Network (Wang et al., 2016; Nothias et al., 2020). The resulting MN was composed of 16,139 nodes (5,922 singletons) and 22,656 edges. As a result of the annotation process against the GNPS open databases, 2494 nodes (ca 15%) were annotated, wherefrom 1751 nodes (ca 11%) were considered valid after cleaning and filtering. This was followed by extensive spectral matching against *in-silico* predicted MS^2^ NPs databases from ISDB-DNP and computational annotation with SIRIUS (Allard et al., 2016; Dührkop et al., 2019; Rutz et al., 2022). After these processes, a total of 11,370 nodes were annotated (ca 70%). The overall combined structural annotation rate for the MN was around 68 %.

The set of Celastraceae extracts was used to test the capacity of *Inventa* to prioritize extracts with a chemical novelty potential. The main results obtained with default parameters are shown in **Table 1** (full results **Supplementary Table S2**).

**Table 1.** Top and lowest five results from the application of Inventa on the Celastraceae collection.

| Rank | Genus | Species | Organ | FS | FC | LC | rcg | rcf | CC | nccs | nccg | SC | PR |
| --- | --- | --- | --- | --- | --- | --- | --- | --- | --- | --- | --- | --- | --- |
| 1 | *Pristimera* | *Pristimera indica* | Roots | 0.37 | 0.36 | 0.87 | 2 | 8 | 1 | Agarofuran<br>sesquiterpenoid | Agarofuran<br>sesquiterpenoid | 1 | 3.23 |
| 2 | *Euonymus* | *Euonymus sanguineus* | Roots | 0.26 | 0.24 | 0.78 | 1 | 440 | 1 | Cinnamic acids<br>and derivatives,<br>etc. | Cinnamic acids<br>and derivatives,<br>etc. | 1 | 3.02 |
| 3 | *Celastrus* | *Celastrus paniculatus* | Seeds | 0.37 | 0.34 | 0.66 | 71 | 732 | 1 | Cholestane<br>steroids,<br>Agarofuran<br>sesquiterpenoids | Cholestane<br>steroids | 1 | 3.00 |
| 4 | *Salacia* | *Salacia letestuana* | Fruits | 0.44 | 0.44 | 0.77 | 0 | 514 | 0 |  |  | 1 | 2.18 |
| 5 | *Euonymus* | *Euonymus*<br>*cochinchinensis* | Leaves | 0.37 | 0.37 | 0.79 | 0 | 440 | 0 |  |  | 1 | 2.13 |
| // | // | // | // | // | // | // | // | // | // | // | // | // | // |
| 72 | *Euonymus* | *Euonymus myrianthus* | Stems | 0.1 | 0.08 | 0.79 | 0 | 440 | 0 |  |  | 0 | 0.87 |
| 73 | *Salacia* | *Salacia cochinchinensis* | Branches | 0.1 | 0.09 | 0.77 | 0 | 514 | 0 |  |  | 0 | 0.86 |
| 74 | *Euonymus* | *Euonymus dielsianus* | Roots | 0.06 | 0.06 | 0.79 | 0 | 440 | 0 |  |  | 0 | 0.85 |
| 75 | *Tripterygium* | *Tripterygium wilfordii* | Roots | 0.24 | 0.22 | -0.40 | 1011 | 1353 | 1 | Open-chain<br>polyketides | Open-chain<br>polyketides | 0 | 0.81 |
| 76 | *Tripterygium* | *Tripterygium wilfordii* | Stems | 0.19 | 0.17 | -0.40 | 1011 | 1353 | 1 | Other<br>Octadecanoids | Other<br>Octadecanoids | 0 | 0.76 |

Inventa’s results table contains all the components scoring and overall priority score (sum of FC, LC, CC, and SC). The plant extracts shown were ranked based on the PS value. The *Pristimera indica* roots extract was ranked first with a PR value of 3.23. It presents an FS of 0.37, indicating that 37% of its features are specific, with at least 90% of the normalized peak area in this extract. Among these specific features, only 1% was annotated as reflected by the FC 0.36, which indicates that 36% of the ions are specific and unannotated. At this stage, evaluation of these two values indicates that such features are very specific at the data set’s level, and the absence of annotations possibly reflects the presence of novel or unreported molecules.

This extract presents an LC of 0.87. For this study, the score was considered if less than 10 compounds were found in the species (crs), less than fifty in the genus (crg), and less than five hundred in the family (crf); these correspond to user-defined parameters. In the case of this extract, only two compounds were reported in the species and 8 in the genus (Chang et al., 2003; Gao et al., 2007; Ramos et al.). Application of these values in the formula shown in **Figure 1.B**, lower the maximum LC value of 1 by 0.13 only, highlighting poorly studied plant species. In our case, the values of reported compounds in the family (6,064) affected equally all extracts since they belong to the same family. In our set, evaluation of this component is important since there is a substantial number of reports for certain genera like Celastrus and Salacia, with 732 and 514 compounds, respectively. For example, the extract ranked three has the same FC as first rank. However, the high number of compounds reported in both species and genus (LC 0.66) suggested a lower possibility of finding new compounds.

The CC value of 1, addition of CCs 0.5 and CCg 0.5, implied that at least one chemical class proposed by SIRIUS-CANOPUS had not been reported in species or the genus. CANOPUS proposed the chemical class dihydro-*β*-agarofuran sesquiterpenoids for the major peaks in the extract according to the BPI (see **Figure 2.A**, zone highlighted in green). Finally, the SC value of 1 indicated that the extract was considered dissimilar within the data set based on its total spectral pattern (MEMO vector), implying a particular composition. A detailed evaluation of the annotation results for *Pristimera indic*a roots extract revealed that the only few annotated features were dihydro-*β*-agarofuran previously reported in *Celastrus angulatus* (an ISDB-DNP spectral match) and two friedelane triterpenoids, pristimerin, and maytenin (GNPS matches), both previously reported in the Celastraceae family (See **Supplementary Table S3**). Considering these annotation results and these chemotaxonomic considerations, we interpreted that several of the most intense ions annotated as dihydro-*β*-agarofurans for by CANOPUS, as shown in **Figure 2.A** (zone highlighted in green), were new derivatives. **Figure 2.B** shows an ion map of all detected features of *Pristimera indic*a roots extract (unfiltered normalized area intensity) is displayed. In this map, a color coding represents the category for the features: specific unannotated (blue, worthy of isolation), specific annotated (green), and not interesting (yellow, nonspecific annotated). Such visualization helps localize inside the extract of interest the TIC peaks and their features, potentially corresponding to novel NPs.

**Figure 2.**
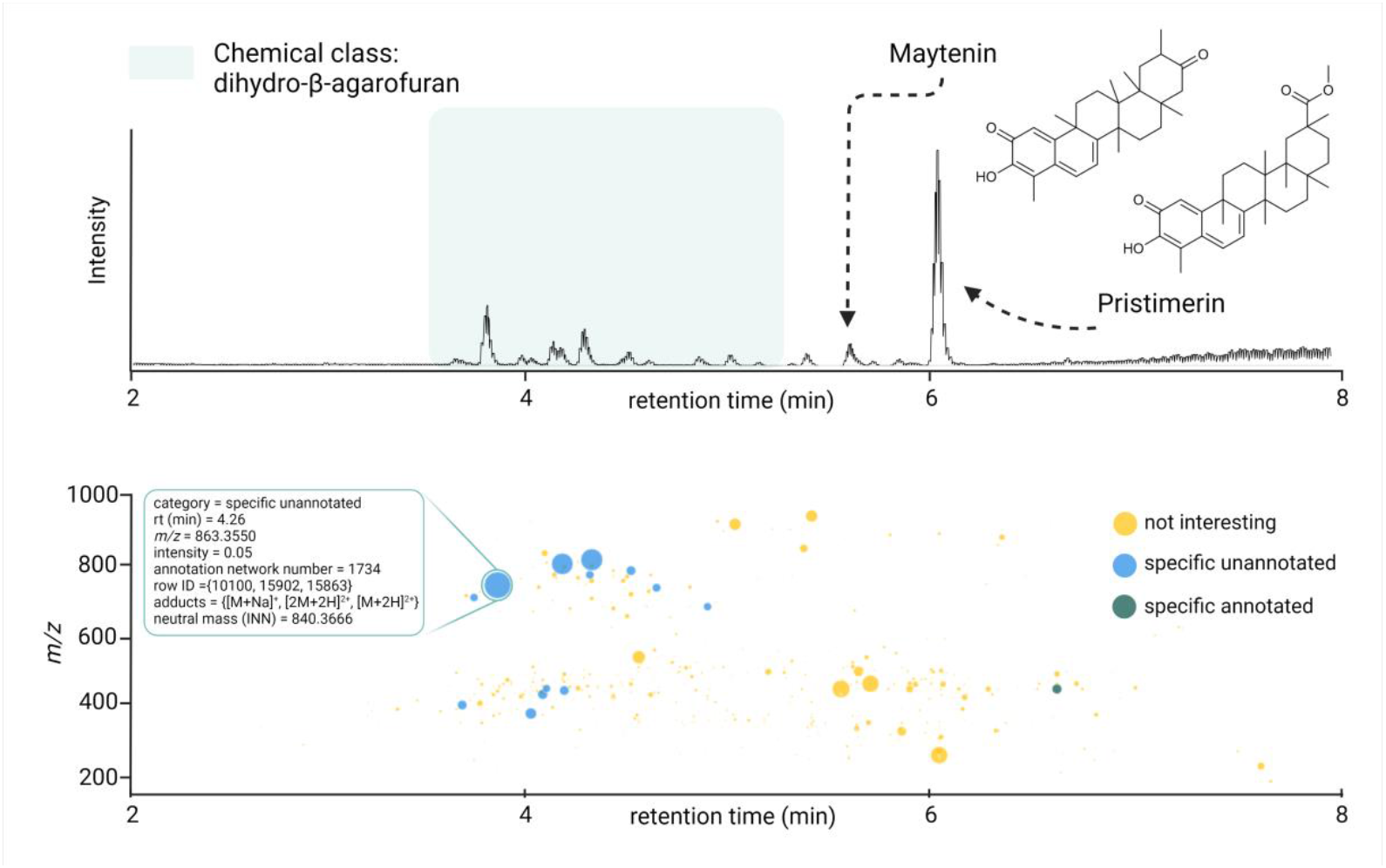
**A)** UHPLC-HRMS chromatogram (BPI positive ion mode) showing the region where the dihydro-*β*-agarofuran sesquiterpenoids derivatives are suspected and displaying the only two compounds annotated for *P. indica* roots (plant with the highest PS). **B)** Ion identity networking-based interactive ion <u>map</u> showing the combined results of the FC and CC for the IIN. In such display all features of a single neutral molecule are grouped under a single spot. The *IIN* are displayed according to their status (specific unannotated (blue), specific annotated (green), and non-specific unannotated - not interesting-(yellow)). Complementary information (adducts, row id, chemical class, etc.) are displayed interactively for each IIN if available, as shown in the zoom sections for the ion identity network 1734. The intensities in both cases (bar’s height and bubble’s size) are proportional to the original quantification table (before any filtering step). The scatter plot shows the *m/z* ratio of each feature (or ion network identity) on the y-axis. The feature-based <u>ion map</u> can be found in **Supplementary Figure S82**.

**Figure 3.**
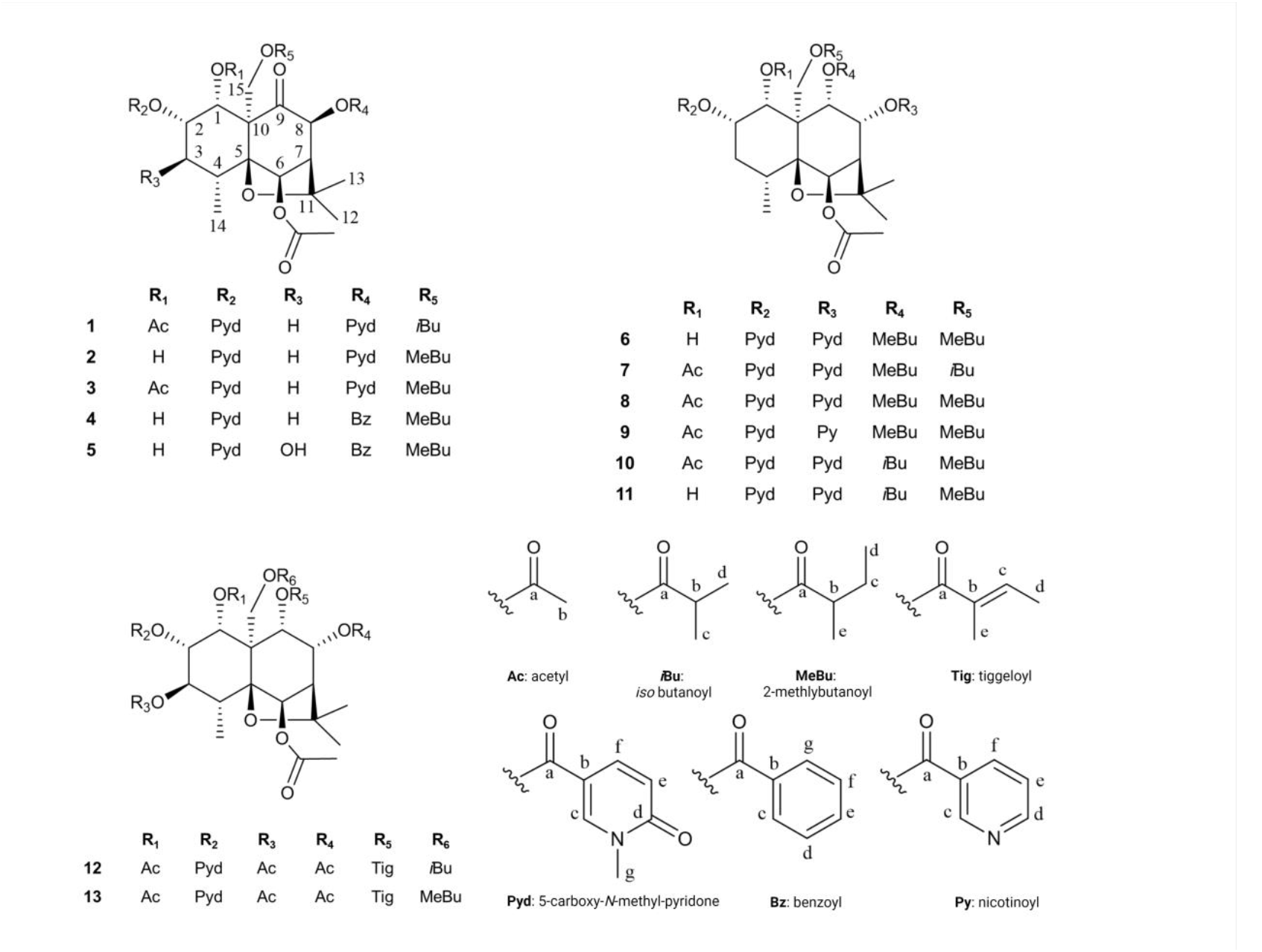
Original dihydro-*β*-agarofuran derivatives isolated from the *Pristimera indica* roots extract.

Based on Inventa’s score and the above considerations, the *Pristimera indic*a roots extract was prioritized and subjected to an in-depth phytochemical investigation for *de novo* structural identification of the potentially new NPs.

### 3.4 Considerations on intensity-based filters integration in *Inventa*

Based on the metabolite profiling results for the prioritized *Pristimera indic*a roots extract, shown in **Figure 2**, most of the unannotated specific ions corresponded to high-intensity features. To evaluate this aspect in the prioritization process of the extracts, two different filters have been implemented in *Inventa*. The aim of such filters is to enable the user to explore how filtering-out the least abundant features affects the *Inventa* scoring results. For this, the original aligned feature table is normalized sample-wise (each row corresponding to an extract). The filters are applied to each sample. These filtered data are then treated by Inventa as the input for all the computations, as described above. The first filter minimizes to zero all the features with a normalized area of less than 2% in each extract (user-defined value, see **Supplementary Figure S83**). For example, after the application of this filter, the number of features for the *Pristimera indic*a (Willd.) A.C.Sm roots was reduced by 85% (from 727 to 104). The second filter uses the quantile distribution for the features normalized area intensity. With this quantile filter only features with a normalized area intensity above the defined quantile value are considered ((default quantile value is 0.75); the features that have their normalized areas below this quantile value are minimized to zero (see **Supplementary Figure S83**). For the *Pristimera indic*a roots extract, the number of features varied from 727 to 182 with the default quantile value.

Both filters can be applied independently or sequentially according to the user’s preferences. **Table 2** shows the differences in the results obtained when both filters are used jointly for the set of Celastraceae plants. For the *Pristimera indic*a roots extract, the application of the quantile-based filter on the remaining 104 features after intensity-based filtering left a total of 26 features. This data reduction was found consistent with the visible BPI peaks after visual inspection of the chromatogram (see **Figure 2.A**). Furthermore, it was found to be in good agreement with all the prioritized NPs that could finally be isolated, as detailed below.

**Table 2.**
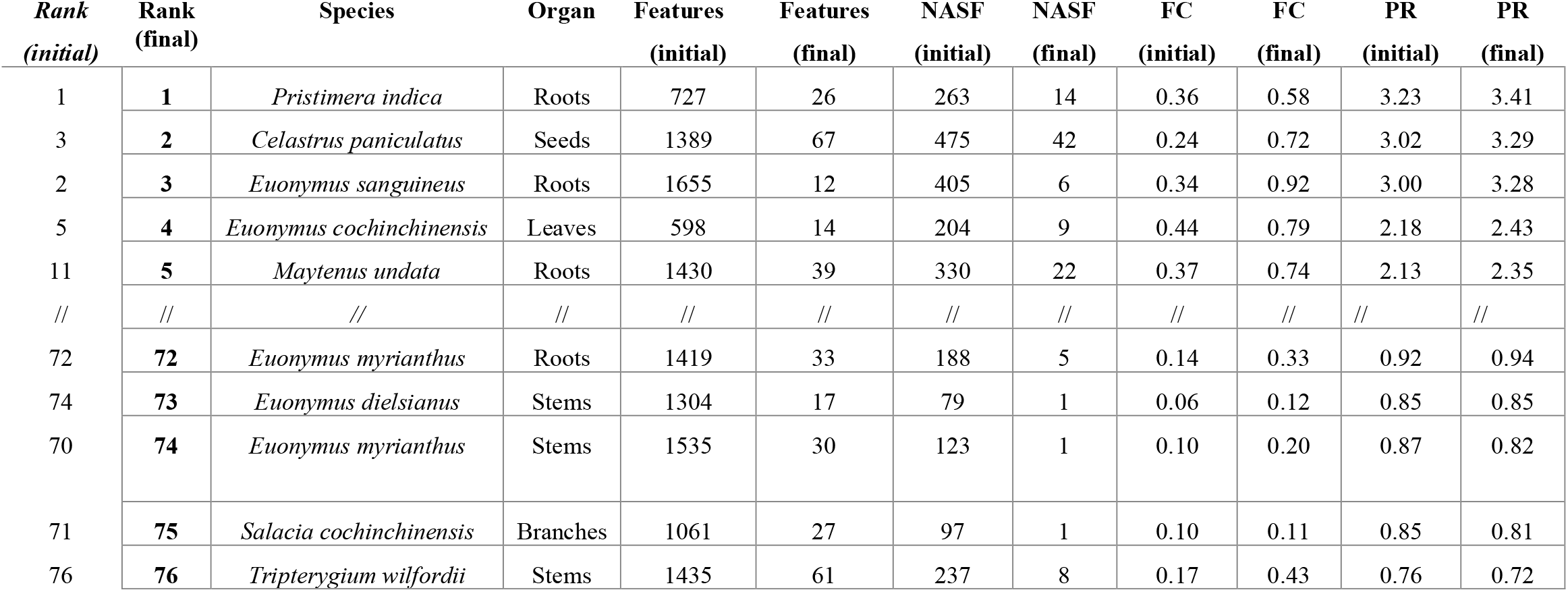
Top and lowest five extracts of the Celastraceae collection after application with and without filters of *Inventa. initial*: before filtering; *final*: after filtering; NASF: unannotated specific features; FC: Feature Component; PS: Priority Score.

| *Rank*<br>*(initial)* | *Rank*<br>*(final)* | *Species* | *Organ* | *Features*<br>*(initial)* | *Features*<br>*(final)* | *NASF*<br>*(initial)* | *NASF*<br>*(final)* | *FC*<br>*(initial)* | *FC*<br>*(final)* | *PR*<br>*(initial)* | *PR*<br>*(final)* |
| --- | --- | --- | --- | --- | --- | --- | --- | --- | --- | --- | --- |
| 1 | **1** | *Pristimera indica* | Roots | 727 | 26 | 263 | 14 | 0.36 | 0.58 | 3.23 | 3.41 |
| 3 | **2** | *Celastrus paniculatus* | Seeds | 1389 | 67 | 475 | 42 | 0.24 | 0.72 | 3.02 | 3.29 |
| 2 | **3** | *Euonymus sanguineus* | Roots | 1655 | 12 | 405 | 6 | 0.34 | 0.92 | 3.00 | 3.28 |
| 5 | **4** | *Euonymus cochinchinensis* | Leaves | 598 | 14 | 204 | 9 | 0.44 | 0.79 | 2.18 | 2.43 |
| 11 | **5** | *Maytenus undata* | Roots | 1430 | 39 | 330 | 22 | 0.37 | 0.74 | 2.13 | 2.35 |
| // | // | // | // | // | // | // | // | // | // | // | // |
| 72 | **72** | *Euonymus myrianthus* | Roots | 1419 | 33 | 188 | 5 | 0.14 | 0.33 | 0.92 | 0.94 |
| 74 | **73** | *Euonymus dielsianus* | Stems | 1304 | 17 | 79 | 1 | 0.06 | 0.12 | 0.85 | 0.85 |
| 70 | **74** | *Euonymus myrianthus* | Stems | 1535 | 30 | 123 | 1 | 0.10 | 0.20 | 0.87 | 0.82 |
| 71 | **75** | *Salacia cochinchinensis* | Branches | 1061 | 27 | 97 | 1 | 0.10 | 0.11 | 0.85 | 0.81 |
| 76 | **76** | *Tripterygium wilfordii* | Stems | 1435 | 61 | 237 | 8 | 0.17 | 0.43 | 0.76 | 0.72 |

The effect of filtering was evaluated for the complete set of Celastraceae, and it significantly lowered the number of features for all extracts. *Inventa*’s scores were not strongly affected but highlighted better the putative novel NPs. Depending on the set of extracts to be evaluated, comparing the results before and after filtering may help the selection process.

### 3.5 Isolation and *de novo* structural identification of thirteen new dihydro-*β*-agarofuran from the *Pristimera indica* roots extract

Inspection of the *Inventa*’s scores obtained before and after filtering proposed the *Pristimera indic*a roots extract as the best potential source of novel NPs. To verify if this plant contains potentially new *β*-agarofuran sesquiterpenoids, its roots material was extracted at a larger scale to generate enough extract for isolation. For this purpose, three successive extraction steps with solvents of increasing polarity (hexane, ethyl acetate, and methanol) were used. A comparison of their UHPLC-HRMS profiling with the original ethyl acetate extract showed that the main NPs of interest were present in the ethyl acetate and methanolic extracts.

The chromatographic optimization and isolation efforts were focused on the retention window from 3.0 to 5.0 min since this region contained most of the unannotated and specific compounds (see **Figure 2.A**). Before isolation, the UHPLC chromatographic conditions were optimized based on the original UHPLC-HRMS chromatogram. A geometric gradient transfer method (Guillarme et al., 2008) was used to scale up the conditions to an analytical HPLC level for evaluation and validation. The HPLC scale conditions were calculated at the semi-preparative HPLC scale for isolation. This process enabled the alignment of the analytical and semi-preparative HPLC scales with the UHPLC scale and localizing the NPs of interest. The isolation was done using a dry-load-based injection, keeping a high resolution, and maximizing the sample load (Queiroz et al., 2019) (**Supplementary Figure S84**).

This methodology efficiently allowed to yield thirteen compounds with enough material for *de novo* structural identification (see **Figure 5)**, only with three consecutive injections of 50 mg each (for each extract, ethyl acetate and methanol). From the ethyl acetate extract, ten pure compounds (**1**-**7, 9, 12**-**13**) and a mixture containing two compounds (**8, 10**) were obtained. The mixture was separated under optimized conditions to purify both compounds, giving twelve pure compounds in total. To obtain compound **11**, the methanolic extract was separated in the same conditions. **Figure 4** summarizes the position of the isolated compounds in both the chromatogram and the original molecular network. Their *de novo* structural elucidation and absolute configuration are described below.

**Figure 4.**
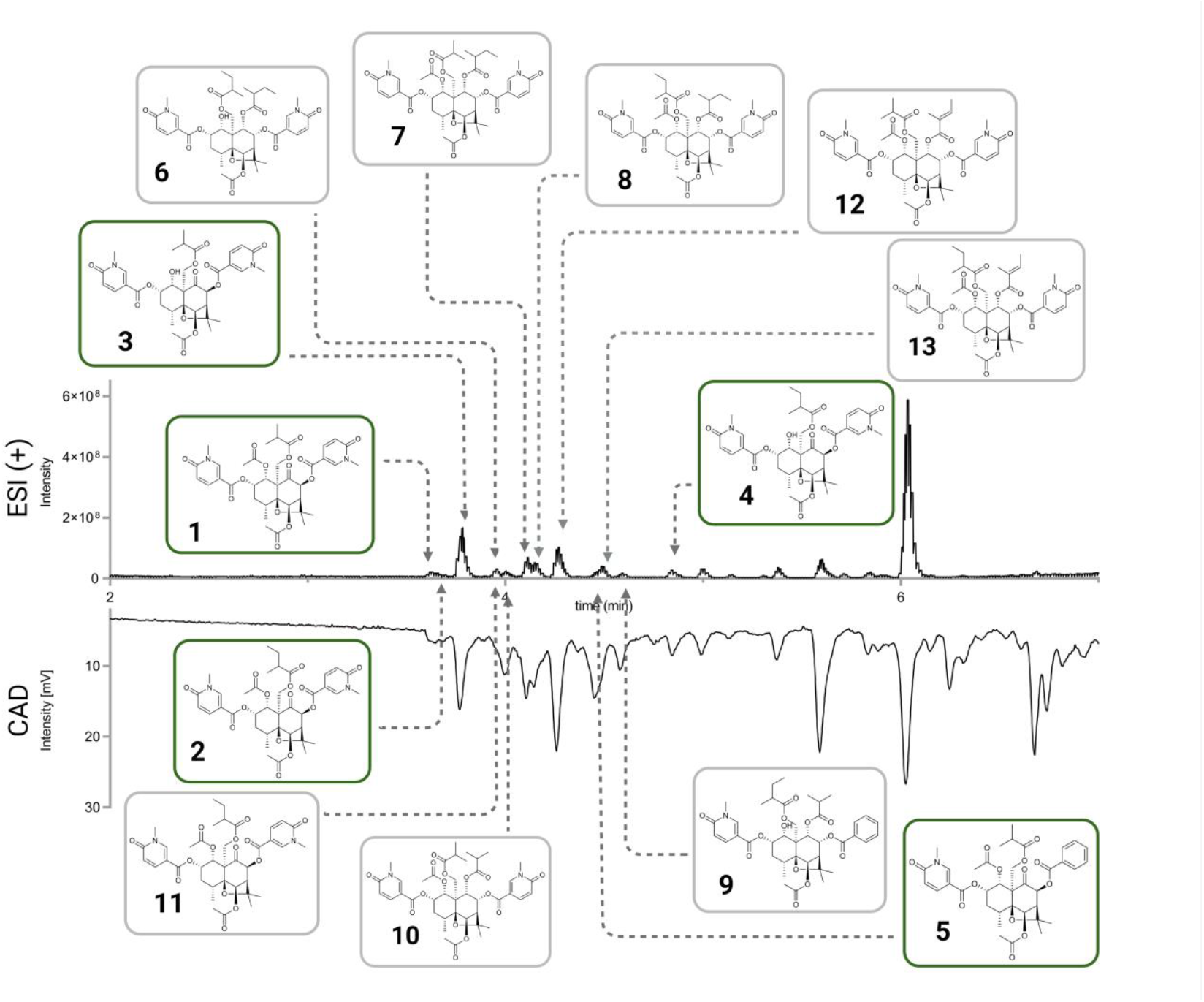
The relative position of the isolated compound (**1**-**13**) in the chromatogram for the ethyl acetate extract of *Pristimera indica* roots. The upper chromatographic trace corresponds to the ESI in positive ionization mode, while the lower trace corresponds to the Charged Aerosol Detector (CAD), a semi-quantitative trace. Compounds highlighted in green hold a new 9-oxodihydro-*β*-agarofuran base scaffold.

**Figure 5.**
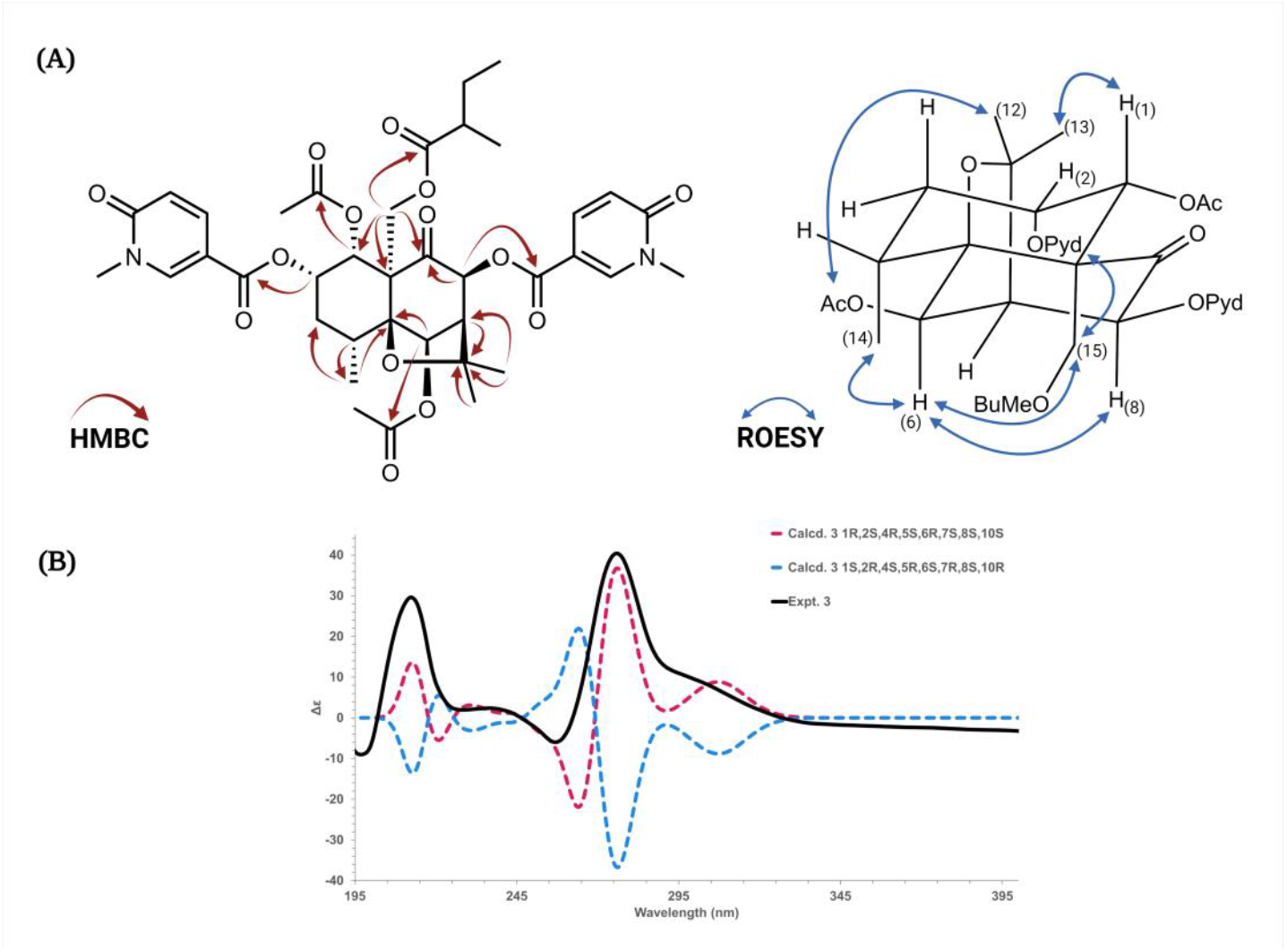
A) HMBC Key and ROESY correlations for the compounds isolated from *P. indica* roots extract. b) Experimental and B3LYP/def2svp//B3LYP/6-31G(d,p) calculated spectra in acetonitrile for compound **3**.

Analysis of the NMR data confirmed that all the isolated compounds were dihydro-*β*-agarofuran as proposed by the CC chemical classes of *Inventa*. They all presented the characteristic 2 methyl singlets at δ_H_ between 1.41 and 1.56 (H_3_-12 and H_3_-13), a methyl doublet at δ_H_ between 1.11 and 1.22 (H_3_-14), an oxymethylene at δ_H_ between 4.99-5.51 and 4.18-4.79 (H-15’ and H-15’’, respectively), an acetate in C-6 at δ_H_ between 2.12-2.21 (H_3_-6b), a particularly deshielded H-6 proton (δ_H_ between 6.24-6.78) and several oxygenated methines (δ_H_ between 3.92-6.18). These compounds could be divided into 3 series.

The first one (compounds **1**-**5**, see **Table 3**.) had a carbonyl in C-9 observed at δ_C_ between 203-207 on the ^13^C and HMBC spectra. The purest compound and the one isolated in the greatest quantity is compound **3**, it will be described first.

**Table 3.** ^1^H and ^13^C NMR spectroscopic data for compounds **1**-**5** (δ in ppm, *J* in Hz).

| Compound 1 |  |  | Compound 2 |  | Compound 3 |  | Compound 4 |  | Compound 5 |  |
| --- | --- | --- | --- | --- | --- | --- | --- | --- | --- | --- |
| No | $\delta_{\text{H}}$ (Multiplicity, $J$, nH) | $\delta_{\text{C}}$ | $\delta_{\text{H}}$ (Multiplicity, $J$, nH) | $\delta_{\text{C}}$ | $\delta_{\text{H}}$ (Multiplicity, $J$, nH) | $\delta_{\text{C}}$ | $\delta_{\text{H}}$ (Multiplicity, $J$, nH) | $\delta_{\text{C}}$ | $\delta_{\text{H}}$ (Multiplicity, $J$, nH) | $\delta_{\text{C}}$ |
| 1 | 5.74 (d, 3.9 Hz, 1H) | 71.1 | 4.61 (d, 3.8 Hz, 1H) | 70.0 | 5.75 (d, 3.9 Hz, 1H) | 71.1 | 4.62 (d, 3.8 Hz, 1H) | 70.4 | 4.88 (d, 4.0 Hz, 1H) | 67.0 |
| 2 | 5.54 (q, 3.9,3.0 Hz, 1H) | 72.5 | 5.44 (q, 3.8,3.5 Hz, 1H) | 74.7 | 5.55 (q, 3.9,2.8 Hz, 1H) | 72.5 | 5.45 (q, 3.8,2.7 Hz, 1H) | 75.1 | 5.42 (ddd, 4.0, 3.3, 1.1 Hz, 1H) | 77.6 |
| 3 | $\alpha$ 1.97 (dd, 14.3,3.0 Hz, 1H)<br>$\beta$ 2.40 (m, 1H) | 31.6 | $\alpha$ 1.98 (dd, 15.2,3.5 Hz, 1H)<br>$\beta$ 2.31 (ddd, 15.2,6.4,3.8 Hz, 1H) | 31.3 | $\alpha$ 1.97 (dd, 14.0,2.8 Hz, 1H)<br>$\beta$ 2.41 (m, 1H) | 31.6 | $\alpha$ 1.99 (dt, 15.4,2.7,0.9 Hz, 1H)<br>$\beta$ 2.32 (ddd, 15.4,6.4,3.6 Hz, 1H) | 31.7 | 3.92 (dd, 3.3,1.8 Hz, 1H) | 73.1 |
| 4 | 2.40 (m, 1H) | 34.1 | 2.36 (m, 1H) | 34.1 | 2.41 (m, 1H) | 34.1 | 2.37 (m, 1H) | 34.4 | 2.51 (qdd, 7.9,1.8,1.1 Hz, 1H) | 40.9 |
| 5 | - | 91.9 | - | 91.3 | - | 91.9 | - | 91.6 | - | 92.9 |
| 6 | 6.31 (s, 1H) | 77.6 | 6.24 (s, 1H) | 77.4 | 6.30 (d, 0.8 Hz, 1H) | 77.7 | 6.28 (s, 1H) | 77.8 | 6.33 (s, 1H) | 78.3 |
| 7 | 2.97 (d, 3.4 Hz, 1H) | 54.1 | 2.97 (d, 3.4 Hz, 1H) | 54.0 | 2.97 (dd, 3.4,0.8 Hz, 1H) | 54.1 | 3.00 (d, 3.5 Hz, 1H) | 54.4 | 3.00 (d, 3.5 Hz, 1H) | 53.2 |
| 8 | 6.03 (d, 3.4 Hz, 1H) | 78.8 | 6.10 (d, 3.4 Hz, 1H) | 78.4 | 6.01 (d, 3.4 Hz, 1H) | 78.8 | 6.17 (d, 3.5 Hz, 1H) | 78.8 | 6.18 (d, 3.5 Hz, 1H) | 78.7 |
| 9 | - | 203.1 | - | 206.8 | - | 203.1 | - | 207.0 | - | 206.3 |
| 10 | - | 60.2 | - | 61.4 | - | 60.1 | - | 61.7 | - | 61.6 |
| 11 | - | 85.7 | - | 85.0 | - | 85.6 | - | 85.2 | - | 87.0 |
| 12 | 1.49 (s, 3H) | 30.2 | 1.48 (s, 3H) | 29.9 | 1.49 (s, 3H) | 30.2 | 1.48 (s, 3H) | 27.5 | 1.54 (s, 3H) | 27.4 |
| 13 | 1.42 (s, 3H) | 27.2 | 1.43 (s, 3H) | 27.1 | 1.41 (s, 3H) | 27.2 | 1.49 (s, 3H) | 30.2 | 1.53 (s, 3H) | 30.1 |
| 14 | 1.22 (d, 7.6 Hz, 3H) | 17.5 | 1.15 (d, 7.7 Hz, 3H) | 17.4 | 1.22 (d, 7.7 Hz, 3H) | 17.5 | 1.15 (d, 7.6 Hz, 3H) | 17.7 | 1.16 (d, 7.9 Hz, 3H) | 15.4 |
| 15 | 4.70 (d, 12.2 Hz, 1H)<br>5.10 (d, 12.2 Hz, 1H) | 63.9 | 4.77 (d, 12.2 Hz, 1H)<br>5.05 (d, 12.2 Hz, 1H) | 63.6 | 4.65 (d, 12.3 Hz, 1H)<br>5.12 (d, 12.3 Hz, 1H) | 63.9 | 4.79 (d, 12.3 Hz, 1H)<br>5.06 (d, 12.3 Hz, 1H) | 64.0 | 4.75 (d, 12.3 Hz, 1H)<br>4.99 (d, 12.3 Hz, 1H) | 63.8 |
| **R1** |  |  |  |  |  |  |  |  |  |  |
| 1a | - | 171.4 |  |  | - | 171.2 |  |  |  |  |
| 1b | 1.95 (s, 3H) | 21.0 |  |  | 1.95 (s, 3H) | 21.0 |  |  |  |  |
| **R2** |  |  |  |  |  |  |  |  |  |  |
| 2a | - | no | - | 164.1 | - | 164.8 | - | no | - | 164.6 |
| 2b | - | 111.1 | - | 111.3 | - | 111.1 | - | 111.7 | - | 111.2 |
| 2c | 8.48 (d, 2.6 Hz, 1H) | 145.7 | 8.48 (d, 2.6 Hz, 1H) | 145.1 | 8.47 (d, 2.5 Hz, 1H) | 145.6 | 8.49 (d, 2.5 Hz, 1H) | 145.4 | 8.48 (d, 2.6 Hz, 1H) | 145.6 |
| 2d | - | 165.2 | - | 165.0 | - | 165.2 | - | 165.2 | - | 165.3 |
| 2e | 6.58 (d, 9.5 Hz, 1H) | 119.9 | 6.57 (d, 9.5 Hz, 1H) | 119.4 | 6.58 (d, 9.5 Hz, 1H) | 119.9 | 6.58 (d, 9.4 Hz, 1H) | 119.8 | 6.58 (d, 9.5 Hz, 1H) | 119.8 |
| 2f | 8.01 (dd, 9.5,2.6 Hz, 1H) | 140.6 | 8.04 (dd, 9.5,2.6 Hz, 1H) | 140.5 | 8.01 (dd, 9.5,2.5 Hz, 1H) | 140.6 | 8.04 (dd, 9.4,2.5 Hz, 1H) | 140.8 | 8.04 (dd, 9.5,2.6 Hz, 1H) | 140.8 |
| 2g | 3.70 (s, 3H) | 38.7 | 3.71 (s, 3H) | 38.3 | 3.70 (s, 3H) | 38.7 | 3.71 (s, 3H) | 38.7 | 3.70 (s, 3H) | 38.7 |
| **6-Ac** |  |  |  |  |  |  |  |  |  |  |
| 6a | - | 171.4 | - | 171.1 | - | 171.3 | - | 171.3 | - | 171.2 |
| 6b | 2.19 (s, 3H) | 20.9 | 2.18 (s, 3H) | 20.6 | 2.19 (s, 3H) | 21.0 | 2.18 (s, 3H) | 21.0 | 2.20 (s, 3H) | 21.0 |
| **R4** |  |  |  |  |  |  |  |  |  |  |
| 8a | - | no | - | 163.6 | - | 163.9 | - | 166.3 | - | 166.3 |
| 8b | - | 110.7 | - | 110.5 | - | 110.7 | - | 130.6 | - | 130.6 |
| 8c | 8.44 (d, 2.5 Hz, 1H) | 146.3 | 8.47 (d, 2.5 Hz, 1H) | 145.9 | 8.44 (d, 2.5 Hz, 1H) | 146.3 | 8.07 (dd, 8.0,1.3 Hz, 1H) | 130.8 | 8.07 (dd, 8.0,1.2 Hz, 1H) | 130.8 |
| 8d | - | 165.2 | - | 164.8 | - | 165.2 | 7.53 (tt, 8.0,1.3 Hz, 1H) | 129.8 | 7.53 (t, 8.0 Hz, 1H) | 129.8 |
| 8e | 6.56 (d, 9.5 Hz, 1H) | 119.6 | 6.57 (d, 9.5 Hz, 1H) | 119.4 | 6.56 (d, 9.5 Hz, 1H) | 119.6 | 7.66 (tt, 8.0,1.3 Hz, 1H) | 134.7 | 7.66 (tt, 8.0,1.2 Hz, 1H) | 134.8 |
| 8f | 7.95 (dd, 9.5,2.5 Hz, 1H) | 140.5 | 7.97 (dd, 9.5,2.5 Hz, 1H) | 140.1 | 7.95 (dd, 9.5,2.5 Hz, 1H) | 140.5 | 7.53 (tt, 8.0,1.3 Hz, 1H) | 129.8 | 7.53 (t, 8.0 Hz, 1H) | 129.8 |
| 8g | 3.62 (s, 3H) | 38.7 | 3.63 (s, 3H) | 38.4 | 3.62 (s, 3H) | 38.7 | 8.07 (dd, 8.0,1.3 Hz, 1H) | 130.8 | 8.07 (dd, 8.0,1.2 Hz, 1H) | 130.8 |
| **R5** |  |  |  |  |  |  |  |  |  |  |
| 15a | - | 177.8 | - | 177.8 | - | 177.6 | - | 178.1 | - | 178.0 |
| 15b | 2.80 (hept, 7.0 Hz, 1H) | 35.2 | 2.67 (h, 7.0 Hz, 1H) | 42.0 | 2.65 (h, 7.0 Hz, 1H) | 42.3 | 2.70 (h, 7.0 Hz, 1H) | 42.4 | 2.71 (h, 7.0 Hz, 1H) | 42.4 |
| 15c | 1.27 (d, 7.0 Hz, 3H) | 19.2 | 1.58 (m, 1H)<br>1.77 (m, 1H) | 27.4 | 1.57 (m, 1H)<br>1.76 (m, 1H) | 27.8 | 1.60 (m, 1H)<br>1.80 (m, 1H) | 27.8 | 1.61 (m, 1H)<br>1.81 (m, 1H) | 27.8 |
| 15d | 1.24 (d, 7.0 Hz, 3H) | 19.3 | 0.93 (t, 7.5 Hz, 3H) | 11.6 | 0.92 (t, 7.4 Hz, 3H) | 11.9 | 0.95 (t, 7.5 Hz, 3H) | 11.9 | 0.96 (t, 7.5 Hz, 3H) | 11.9 |
| 15e |  |  | 1.27 (d, 7.0 Hz, 3H) | 16.5 | 1.25 (d, 7.0 Hz, 3H) | 16.7 | 1.30 (d, 7.0 Hz, 3H) | 16.8 | 1.30 (d, 7.0 Hz, 3H) | 16.8 |

Compound **3** was isolated as an amorphous powder with a [M+H]^+^ of *m/z* 755.3017 and a molecular formula of C_38_H_47_N_2_O_14_. The ^1^H-NMR and HSQC spectra indicated the presence of 3 oxymethine (in addition to H-6) at δ_H_/δ_C_ 5.75/71.1 (H/C-1), 5.55/72.5 (H/C-2), and 6.01/78.8 (H/C-8). These methines were substituted by an acetate at δ_H_ 1.95, and two 5-carboxy-*N*-methyl-pyridone at δ_H_ 3.70 (3H, s, H_3_-2g), 6.58 (1H, d, *J* = 9.5 Hz, H-2e), 8.01 (1H, dd, *J* = 9.5, 2.5 Hz, H-2f), and 8.47 (1H, d, *J* = 2.5 Hz, H-2c) for the first one and 3.62 (3H, s, H_3_-8g), 6.56 (1H, d, *J* = 9.5 Hz, H-8e), 7.95 (1H, dd, *J* = 9.5, 2.5 Hz, H-8f), and 8.44 (1H, d, *J* = 2.5 Hz, H-8c) for the second one. The acetate at δ_H_ 1.95 was positioned thanks to its HMBC correlation with the carbonyl C-1a at δ_C_ 171.2 and this latter correlating with H-1. The position of both 5-carboxy-*N*-methyl-pyridones was confirmed by the HMBC correlations from H-8, H-8c, and H-8f to C-8a, from H-2c and H-2f to C-2a, and the weak correlation from H-2 to C-2a. The ROESY correlations from the aromatic protons of the pyridone in C-2 with H-15 agreed with that. The C-3 position was not substituted as indicated by the presence of two methylene protons at δ_H_ 1.97 and 2.41. Finally a 2-methyl-butanoyl group at δ_H_ 0.92 (3H, t, *J* = 7.4 Hz, H_3_-15d), 1.25 (3H, d, *J* = 7.0 Hz, H_3_-15e), 1.57 (1H, m, H-15c’’), 1.76 (1H, m, H-15c’), and 2.65 (1H, h, *J* = 7.0 Hz, H-15b) was linked to C-15 due to the HMBC correlations from H_2_-15, H-15b, H_2_-15c and H_3_-15e to the ester carbonyl C-15a at δ_C_ 177.6. The HMBC correlations from H-1, H-8, and H_2_-15 to the ketone C-9 at δ_C_ 203.1 confirmed the presence of the carbonyl in C-9 (Figure 7.A). Oxidations in this position have never been reported before for the dihydro-*β*-agarofuran-type compounds; usually, the oxo group is in C-8 (Gao et al., 2007). All the other COSY and HMBC correlations confirmed this flat structure. The <u>MS^2^ spectrum</u> for this compound shows fragments associated with the 5-carboxy-*N*-methyl-pyridone (*m/z* 136), and losses of 2-methyl-butanoyl (*m/z* 85) and acetyl groups (*m/z* 59) in agreement with the literature (Kuo et al., 1995). This trend is observed throughout the entire series of compounds.

The ROESY correlations from H_2_-15 to the aromatic protons of the 5-carboxy-*N*-methyl-pyridone in C-2 (H-2c/H-2f) and H-6, from H-6 to H-8 and H_3_-14 indicated that these protons were on the same side of the molecule. The correlation from H_3_-12 to the acetate in C-6 confirmed that the C5-O-C11-C7 bridge is on the opposite side. The weak correlation between H_3_-13 and H-1 indicated that H-1 is in the same orientation as the bridge and that H-1 should be axial (Figure 7.A). Thus, the relative configuration of the substituents in **3** was proposed as 1*α*, 2*α, 4α, 5β*, 6*β, 7β, 8β* and 10*α*.

To establish the absolute configuration of compound **3**, the ECD spectrum was calculated based on the relative configuration proposed by NMR and compared to the experimental data (See **Figure 5.B**). The absolute configuration of the agarofuran moiety (4*R*,5*S*,6*R*,7*S*,10*S*) agrees with the reports in the literature for the type of chemical structure proposed.

After the comparison, compound **3** was assigned as (1*R*,2*S*,4*R*,5*S*,6*R*,7*S*,8*S*,10*S*)-1α,6β-diacetoxy-2α,8β-di-(5-carboxy-*N*-methyl-3-pyridoxy)-15-(2-methylbutanoyloxy)-9-oxodihydro-*β*-agarofuran and named Silviatine C.

Compound **1** was isolated as a white amorphous powder. The molecular formula, C_37_H_45_N_2_O_14_, was calculated based on the HRMS-ESI-MS for [M+H]^+^ of *m/z* 741.2864. The NMR data of **1** were very similar to that of **3**, except that an *iso*-butanoyl group at C-15 replaced the 2-methylbutanoyl group present at the same position in **3**. The HMBC correlations from H-15’ at δ_H_ 5.10 (1H, d, *J* = 12.2 Hz), H-15b at δ_H_ 2.80 (1H, hept, *J* = 7.0 Hz), H_3_-15c at δ_H_ 1.27 (3H, d, *J* = 7.0 Hz) and H_3_-15d at δ_H_ 1.24 (3H, d, *J* = 7.0 Hz) to the ester group C-15a at δ_C_ 177.8 confirmed the position of the *iso*-butanoyl group. Same cross-peaks in the ROESY spectrum as **3** were observed, suggesting the same relative configuration as **3**. After calculation and comparison of the ECD spectra compound **3** was assigned as (1*R*,2*S*,4*R*,5*S*,6*R*,7*S*,8*S*,10*S*)-1*α*,6*β*-diacetoxy-15-*iso*-butanoyloxy-2*α*,8*β*-di-(5-carboxy-*N*-methyl-3-pyridoxy)-9-oxodihydro-*β*-agarofuran, and named Silviatine A.

Compound **2** was a white amorphous powder with a molecular formula of C_36_H_45_N_2_O_13_, calculated for [M+H]^+^ of *m/z* 713.2323. The ^1^H-NMR signals were closely related to those of **3**, indicating that they shared the same core. The major differences were observed for the substituents. Only one acetyl group was found and positioned at C-6 due to the HMBC correlation fromH-6 (δ_H_ 6.24) and the acetate at δ_H_ 2.18 to the carbonyl at δ_c_ 171.1. Position one was suggested to bear a hydroxyl group due to the higher field signal of H-1 (δ_H_ 4.61) and no other HMBC correlations. Two 5-carboxy-*N*-methyl-3-pyrodone substituents were found in positions C-2 (δ_c_ 74.7) and C-8 (δ_c_ 78.4). The substituent in position C-15 (δ_C_ 63.6) was as in **3** a 2-methyl-butanoate moiety consisting of a carbonyl (δ_C_ 177.8), one methine (δ_C_ 42.0, δ_H_ 2.67), one methylene as a diastereotopic system (δ_C_ 27.4, δ_H_ 1.58 and 1.77), two methyl groups (δ_C_ 11.6, δ_H_ 0.93 and δ_C_ 16.5, δ_H_ 1.27). Analysis of the ROESY correlations indicates that the relative configuration is the same as those of compounds **1** and **3**. After comparison of the experimental and calculated ECD spectra, compound **2** was assigned as (1*R*,2*S*,4*R*,5*S*,6*R*,7*S*,8*S*,10*S*)-6*β*-acetoxy-2*α*,8*β*-di-(5-carboxy-*N*-methyl-3-pyridoxy)-1*α*-hydroxy-15-(2-methylbutanoyloxy)-9-oxodihydro-*β*-agarofuran, and named Silviatine B.

Compound **4** had a molecular formula of C_36_H_44_NO_12_ for a [M+H]^+^ of *m/z* 682.2850. The core structure agrees with the one proposed for **2**, due to the same patterns and correlations observed in the NMR data. The molecular formula suggested the presence of just one nitrogen. According to the ^1^H-NMR signals, two different aromatic groups were identified: a 5-carboxy-*N*-methyl-3-pyridone, as in the previous compounds, at δ_H_ 6.58 (1H, d, *J* = 9.4 Hz, H-2e), 8.04 (1H, dd, *J* = 9.4, 2.5 Hz, H-2f), and 8.49 (1H, d, *J* = 2.5 Hz, H-2c), and a benzene at δ_H_ 7.53 (2H, tt, *J* = 8.0, 1.3 Hz, H-8d, H-8f), 7.66 (1H, tt, *J* = 8.0, 1.3 Hz, H-8e), and 8.07 (2H, dd, *J* = 8.0, 1.3 Hz, H-8c, H-8g). The 5-carboxy-*N*-methyl-3-pyridone was positioned in C-2 thanks to the ROESY correlation from H_2_-15 to H-2c and H-2f.. As for compounds **2** and **3**, the position C-15 was functionalized with a 2-methylbutanoate moiety according to the HMBC correlations from H-15a/H-15b (δ_H_ 4.79 and 5.06), H-15b (δ_H_ 2.70), H_2_-15c (δ_H_ 1.60 and 1.80) and H_3_-15e (δ_H_ 1.30) to the carbonyl at δ_C_ 178.1. As in **2**, H-2 (δ_H_ 5.45) is in a higher field, suggesting this position carries an OH group.

As explained above, the relative configuration of **4** was assigned based on the ROESY data as 1*α*, 2*α*, 6*β*, and 8*β*. After comparison of the experimental and calculated ECD spectra, compound **4** was assigned as (1*R*,2*S*,4*R*,5*S*,6*R*,7*S*,8*S*,10*S*)-6*β*-acetoxy-8*β*-benzoyloxy-2*α*-(5-carboxy-*N*-methyl-3002Dpyridoxy)-1*α*-hydroxy-15-(2-methylbutanoyloxy)-9-oxodihydro-*β*-agarofuran, and named Silviatine D.

Compound **5** was obtained as an amorphous white powder, giving a [M+H]^+^ of *m/z* 698.2802 with a molecular formula of C_36_H_44_NO_13_, one oxygen more than compound **4**. NMR data was closely related to **4**; the major difference was the absence of the diastereotopic methylene in C-3; instead, a signal in a lower field was found at δ_H_ 3.92, integrating for one proton as a doublet of doublets (*J* = 3.3, 1.8 Hz). The chemical shift for C-3 (δ_C_ 73.1) suggested the presence of an OH group, and the COSY correlations fromH-2 (δ_H_ 5.42) and H-4 (δ_H_ 2.51) to H-3 corroborated its position. **5** was thus a hydroxyl-derivative of compound **4**. The key ROESY correlations were the same as for previous compounds: from H-15 to H-6, H_3_-14, H-2c, and H-2f, from H-6 to H-8, and from H_3_-12/13 to H-4 and H-1. The ROESY correlation from H-3 to H_3_-14 indicated their *trans* configuration. The absolute configuration of compound **5** was assigned as (1*R*,2*S*,3*S*,4*R*,5*S*,6*R*,7*S*,8*S*,10*S*)-6*β*-acetoxy-8*β*-benzoyloxy-2*α*-(5-carboxy-*N*-methyl-3-pyridoxy)-1*α*,3*β*-dihydroxy-15-(2-methylbutanoyloxy)-9-oxodihydro-*β*-agarofuran, after comparison with the calculated ECD spectra, and named Silviatine E. The second group of dihydro-*β*-agarofuran structures was composed of 6 new alatol-type structures (**6**-**11**, see **Table 4**)., They were oxygenated in almost all positions except C-3.

**Table 4.**
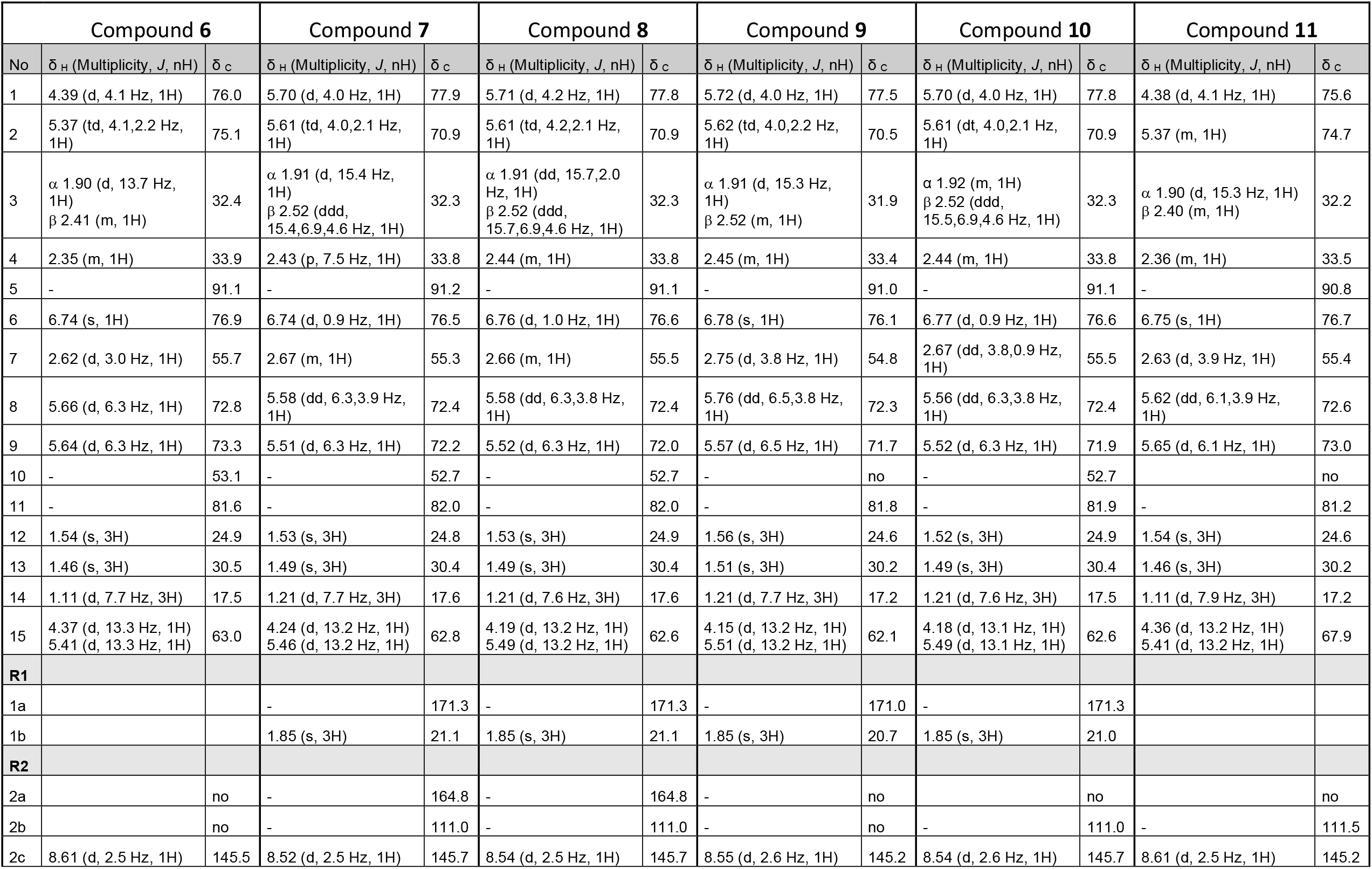
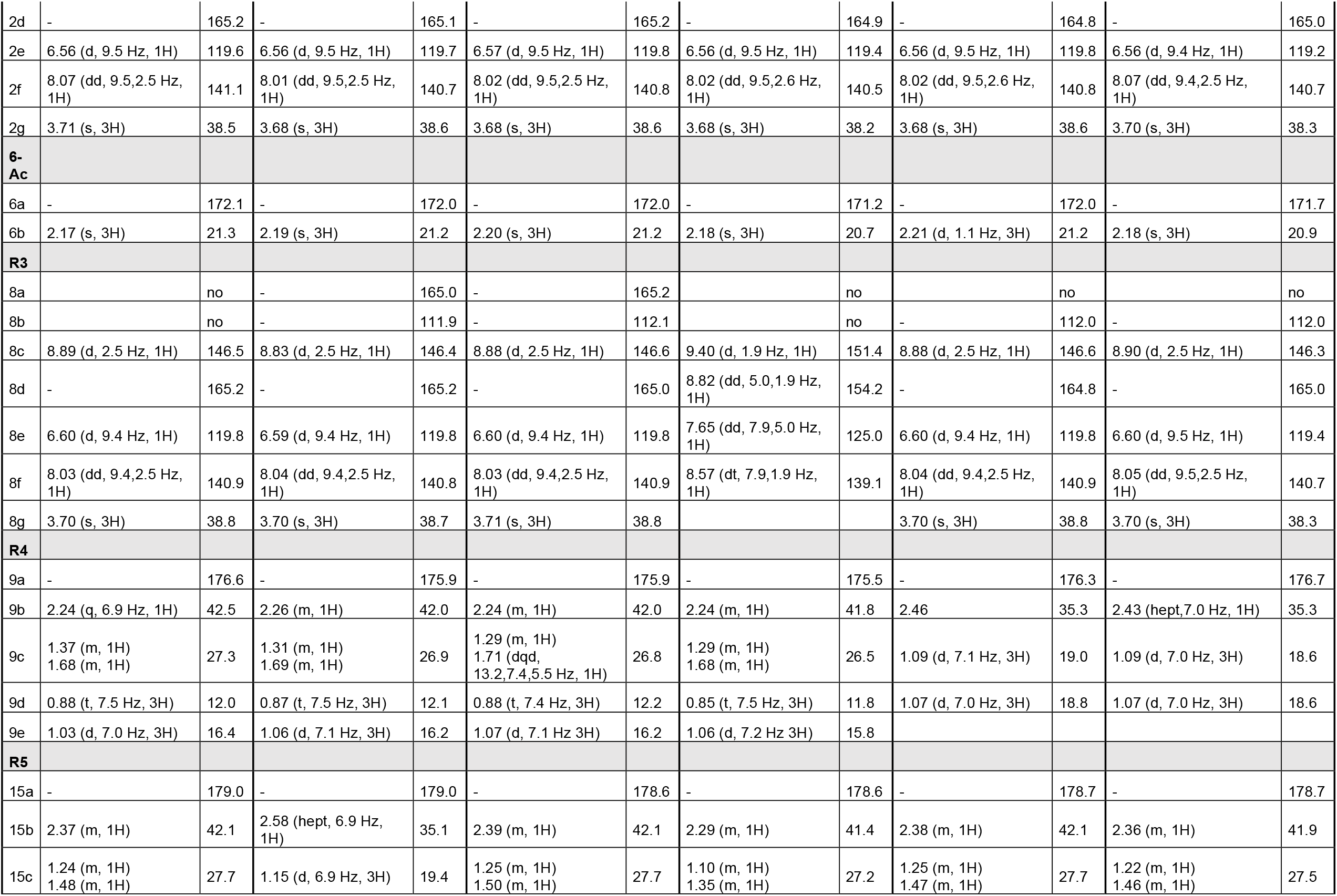
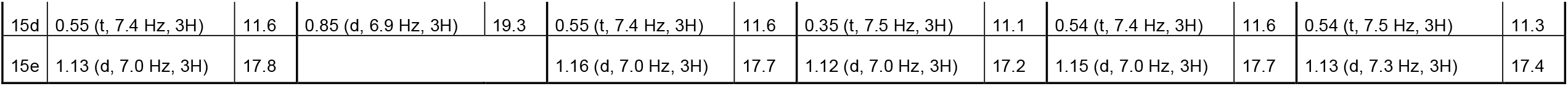
^1^H and ^13^C NMR spectroscopic data for compounds **6**-**11** (δ in ppm, *J* in Hz).

| Compound 6 |  |  | Compound 7 |  | Compound 8 |  | Compound 9 |  | Compound 10 |  | Compound 11 |  |
| --- | --- | --- | --- | --- | --- | --- | --- | --- | --- | --- | --- | --- |
| No | $\delta_{\text{H}}$ (Multiplicity, $J$, nH) | $\delta_{\text{C}}$ | $\delta_{\text{H}}$ (Multiplicity, $J$, nH) | $\delta_{\text{C}}$ | $\delta_{\text{H}}$ (Multiplicity, $J$, nH) | $\delta_{\text{C}}$ | $\delta_{\text{H}}$ (Multiplicity, $J$, nH) | $\delta_{\text{C}}$ | $\delta_{\text{H}}$ (Multiplicity, $J$, nH) | $\delta_{\text{C}}$ | $\delta_{\text{H}}$ (Multiplicity, $J$, nH) | $\delta_{\text{C}}$ |
| 1 | 4.39 (d, 4.1 Hz, 1H) | 76.0 | 5.70 (d, 4.0 Hz, 1H) | 77.9 | 5.71 (d, 4.2 Hz, 1H) | 77.8 | 5.72 (d, 4.0 Hz, 1H) | 77.5 | 5.70 (d, 4.0 Hz, 1H) | 77.8 | 4.38 (d, 4.1 Hz, 1H) | 75.6 |
| 2 | 5.37 (td, 4.1,2.2 Hz, 1H) | 75.1 | 5.61 (td, 4.0,2.1 Hz, 1H) | 70.9 | 5.61 (td, 4.2,2.1 Hz, 1H) | 70.9 | 5.62 (td, 4.0,2.2 Hz, 1H) | 70.5 | 5.61 (td, 4.0,2.1 Hz, 1H) | 70.9 | 5.37 (m, 1H) | 74.7 |
| 3 | $\alpha$ 1.90 (d, 13.7 Hz, 1H)<br>$\beta$ 2.41 (m, 1H) | 32.4 | $\alpha$ 1.91 (d, 15.4 Hz, 1H)<br>$\beta$ 2.52 (ddd, 15.4,6.9,4.6 Hz, 1H) | 32.3 | $\alpha$ 1.91 (dd, 15.7,2.0 Hz, 1H)<br>$\beta$ 2.52 (ddd, 15.7,6.9,4.6 Hz, 1H) | 32.3 | $\alpha$ 1.91 (d, 15.3 Hz, 1H)<br>$\beta$ 2.52 (m, 1H) | 31.9 | $\alpha$ 1.92 (m, 1H)<br>$\beta$ 2.52 (ddd, 15.5,6.9,4.6 Hz, 1H) | 32.3 | $\alpha$ 1.90 (d, 15.3 Hz, 1H)<br>$\beta$ 2.40 (m, 1H) | 32.2 |
| 4 | 2.35 (m, 1H) | 33.9 | 2.43 (p, 7.5 Hz, 1H) | 33.8 | 2.44 (m, 1H) | 33.8 | 2.45 (m, 1H) | 33.4 | 2.44 (m, 1H) | 33.8 | 2.36 (m, 1H) | 33.5 |
| 5 | - | 91.1 | - | 91.2 | - | 91.1 | - | 91.0 | - | 91.1 | - | 90.8 |
| 6 | 6.74 (s, 1H) | 76.9 | 6.74 (d, 0.9 Hz, 1H) | 76.5 | 6.76 (d, 1.0 Hz, 1H) | 76.6 | 6.78 (s, 1H) | 76.1 | 6.77 (d, 0.9 Hz, 1H) | 76.6 | 6.75 (s, 1H) | 76.7 |
| 7 | 2.62 (d, 3.0 Hz, 1H) | 55.7 | 2.67 (m, 1H) | 55.3 | 2.66 (m, 1H) | 55.5 | 2.75 (d, 3.8 Hz, 1H) | 54.8 | 2.67 (dd, 3.8,0.9 Hz, 1H) | 55.5 | 2.63 (d, 3.9 Hz, 1H) | 55.4 |
| 8 | 5.66 (d, 6.3 Hz, 1H) | 72.8 | 5.58 (dd, 6.3,3.9 Hz, 1H) | 72.4 | 5.58 (dd, 6.3,3.8 Hz, 1H) | 72.4 | 5.76 (dd, 6.5,3.8 Hz, 1H) | 72.3 | 5.56 (dd, 6.3,3.8 Hz, 1H) | 72.4 | 5.62 (dd, 6.1,3.9 Hz, 1H) | 72.6 |
| 9 | 5.64 (d, 6.3 Hz, 1H) | 73.3 | 5.51 (d, 6.3 Hz, 1H) | 72.2 | 5.52 (d, 6.3 Hz, 1H) | 72.0 | 5.57 (d, 6.5 Hz, 1H) | 71.7 | 5.52 (d, 6.3 Hz, 1H) | 71.9 | 5.65 (d, 6.1 Hz, 1H) | 73.0 |
| 10 | - | 53.1 | - | 52.7 | - | 52.7 | - | no | - | 52.7 | - | no |
| 11 | - | 81.6 | - | 82.0 | - | 82.0 | - | 81.8 | - | 81.9 | - | 81.2 |
| 12 | 1.54 (s, 3H) | 24.9 | 1.53 (s, 3H) | 24.8 | 1.53 (s, 3H) | 24.9 | 1.56 (s, 3H) | 24.6 | 1.52 (s, 3H) | 24.9 | 1.54 (s, 3H) | 24.6 |
| 13 | 1.46 (s, 3H) | 30.5 | 1.49 (s, 3H) | 30.4 | 1.49 (s, 3H) | 30.4 | 1.51 (s, 3H) | 30.2 | 1.49 (s, 3H) | 30.4 | 1.46 (s, 3H) | 30.2 |
| 14 | 1.11 (d, 7.7 Hz, 3H) | 17.5 | 1.21 (d, 7.7 Hz, 3H) | 17.6 | 1.21 (d, 7.6 Hz, 3H) | 17.6 | 1.21 (d, 7.7 Hz, 3H) | 17.2 | 1.21 (d, 7.6 Hz, 3H) | 17.5 | 1.11 (d, 7.9 Hz, 3H) | 17.2 |
| 15 | 4.37 (d, 13.3 Hz, 1H)<br>5.41 (d, 13.3 Hz, 1H) | 63.0 | 4.24 (d, 13.2 Hz, 1H)<br>5.46 (d, 13.2 Hz, 1H) | 62.8 | 4.19 (d, 13.2 Hz, 1H)<br>5.49 (d, 13.2 Hz, 1H) | 62.6 | 4.15 (d, 13.2 Hz, 1H)<br>5.51 (d, 13.2 Hz, 1H) | 62.1 | 4.18 (d, 13.1 Hz, 1H)<br>5.49 (d, 13.1 Hz, 1H) | 62.6 | 4.36 (d, 13.2 Hz, 1H)<br>5.41 (d, 13.2 Hz, 1H) | 67.9 |
| **R1** |  |  |  |  |  |  |  |  |  |  |  |  |
| 1a |  |  | - | 171.3 | - | 171.3 | - | 171.0 | - | 171.3 |  |  |
| 1b |  |  | 1.85 (s, 3H) | 21.1 | 1.85 (s, 3H) | 21.1 | 1.85 (s, 3H) | 20.7 | 1.85 (s, 3H) | 21.0 |  |  |
| **R2** |  |  |  |  |  |  |  |  |  |  |  |  |
| 2a |  | no | - | 164.8 | - | 164.8 | - | no |  | no |  | no |
| 2b |  | no | - | 111.0 | - | 111.0 | - | no | - | 111.0 | - | 111.5 |
| 2c | 8.61 (d, 2.5 Hz, 1H) | 145.5 | 8.52 (d, 2.5 Hz, 1H) | 145.7 | 8.54 (d, 2.5 Hz, 1H) | 145.7 | 8.55 (d, 2.6 Hz, 1H) | 145.2 | 8.54 (d, 2.6 Hz, 1H) | 145.7 | 8.61 (d, 2.5 Hz, 1H) | 145.2 |
| 2d | - | 165.2 | - | 165.1 | - | 165.2 | - | 164.9 | - | 164.8 | - | 165.0 |
| 2e | 6.56 (d, 9.5 Hz, 1H) | 119.6 | 6.56 (d, 9.5 Hz, 1H) | 119.7 | 6.57 (d, 9.5 Hz, 1H) | 119.8 | 6.56 (d, 9.5 Hz, 1H) | 119.4 | 6.56 (d, 9.5 Hz, 1H) | 119.8 | 6.56 (d, 9.4 Hz, 1H) | 119.2 |
| 2f | 8.07 (dd, 9.5,2.5 Hz, 1H) | 141.1 | 8.01 (dd, 9.5,2.5 Hz, 1H) | 140.7 | 8.02 (dd, 9.5,2.5 Hz, 1H) | 140.8 | 8.02 (dd, 9.5,2.6 Hz, 1H) | 140.5 | 8.02 (dd, 9.5,2.6 Hz, 1H) | 140.8 | 8.07 (dd, 9.4,2.5 Hz, 1H) | 140.7 |
| 2g | 3.71 (s, 3H) | 38.5 | 3.68 (s, 3H) | 38.6 | 3.68 (s, 3H) | 38.6 | 3.68 (s, 3H) | 38.2 | 3.68 (s, 3H) | 38.6 | 3.70 (s, 3H) | 38.3 |
| **6-Ac** |  |  |  |  |  |  |  |  |  |  |  |  |
| 6a | - | 172.1 | - | 172.0 | - | 172.0 | - | 171.2 | - | 172.0 | - | 171.7 |
| 6b | 2.17 (s, 3H) | 21.3 | 2.19 (s, 3H) | 21.2 | 2.20 (s, 3H) | 21.2 | 2.18 (s, 3H) | 20.7 | 2.21 (d, 1.1 Hz, 3H) | 21.2 | 2.18 (s, 3H) | 20.9 |
| **R3** |  |  |  |  |  |  |  |  |  |  |  |  |
| 8a |  | no | - | 165.0 | - | 165.2 |  | no |  | no |  | no |
| 8b |  | no | - | 111.9 | - | 112.1 |  | no | - | 112.0 | - | 112.0 |
| 8c | 8.89 (d, 2.5 Hz, 1H) | 146.5 | 8.83 (d, 2.5 Hz, 1H) | 146.4 | 8.88 (d, 2.5 Hz, 1H) | 146.6 | 9.40 (d, 1.9 Hz, 1H) | 151.4 | 8.88 (d, 2.5 Hz, 1H) | 146.6 | 8.90 (d, 2.5 Hz, 1H) | 146.3 |
| 8d | - | 165.2 | - | 165.2 | - | 165.0 | 8.82 (dd, 5.0,1.9 Hz, 1H) | 154.2 | - | 164.8 | - | 165.0 |
| 8e | 6.60 (d, 9.4 Hz, 1H) | 119.8 | 6.59 (d, 9.4 Hz, 1H) | 119.8 | 6.60 (d, 9.4 Hz, 1H) | 119.8 | 7.65 (dd, 7.9,5.0 Hz, 1H) | 125.0 | 6.60 (d, 9.4 Hz, 1H) | 119.8 | 6.60 (d, 9.5 Hz, 1H) | 119.4 |
| 8f | 8.03 (dd, 9.4,2.5 Hz, 1H) | 140.9 | 8.04 (dd, 9.4,2.5 Hz, 1H) | 140.8 | 8.03 (dd, 9.4,2.5 Hz, 1H) | 140.9 | 8.57 (dt, 7.9,1.9 Hz, 1H) | 139.1 | 8.04 (dd, 9.4,2.5 Hz, 1H) | 140.9 | 8.05 (dd, 9.5,2.5 Hz, 1H) | 140.7 |
| 8g | 3.70 (s, 3H) | 38.8 | 3.70 (s, 3H) | 38.7 | 3.71 (s, 3H) | 38.8 |  |  | 3.70 (s, 3H) | 38.8 | 3.70 (s, 3H) | 38.3 |
| **R4** |  |  |  |  |  |  |  |  |  |  |  |  |
| 9a | - | 176.6 | - | 175.9 | - | 175.9 | - | 175.5 | - | 176.3 | - | 176.7 |
| 9b | 2.24 (q, 6.9 Hz, 1H) | 42.5 | 2.26 (m, 1H) | 42.0 | 2.24 (m, 1H) | 42.0 | 2.24 (m, 1H) | 41.8 | 2.46 | 35.3 | 2.43 (hept,7.0 Hz, 1H) | 35.3 |
| 9c | 1.37 (m, 1H)<br>1.68 (m, 1H) | 27.3 | 1.31 (m, 1H)<br>1.69 (m, 1H) | 26.9 | 1.29 (m, 1H)<br>1.71 (dq, 13.2,7.4,5.5 Hz, 1H) | 26.8 | 1.29 (m, 1H)<br>1.68 (m, 1H) | 26.5 | 1.09 (d, 7.1 Hz, 3H) | 19.0 | 1.09 (d, 7.0 Hz, 3H) | 18.6 |
| 9d | 0.88 (t, 7.5 Hz, 3H) | 12.0 | 0.87 (t, 7.5 Hz, 3H) | 12.1 | 0.88 (t, 7.4 Hz, 3H) | 12.2 | 0.85 (t, 7.5 Hz, 3H) | 11.8 | 1.07 (d, 7.0 Hz, 3H) | 18.8 | 1.07 (d, 7.0 Hz, 3H) | 18.6 |
| 9e | 1.03 (d, 7.0 Hz, 3H) | 16.4 | 1.06 (d, 7.1 Hz, 3H) | 16.2 | 1.07 (d, 7.1 Hz 3H) | 16.2 | 1.06 (d, 7.2 Hz 3H) | 15.8 |  |  |  |  |
| **R5** |  |  |  |  |  |  |  |  |  |  |  |  |
| 15a | - | 179.0 | - | 179.0 | - | 178.6 | - | 178.6 | - | 178.7 | - | 178.7 |
| 15b | 2.37 (m, 1H) | 42.1 | 2.58 (hept, 6.9 Hz, 1H) | 35.1 | 2.39 (m, 1H) | 42.1 | 2.29 (m, 1H) | 41.4 | 2.38 (m, 1H) | 42.1 | 2.36 (m, 1H) | 41.9 |
| 15c | 1.24 (m, 1H)<br>1.48 (m, 1H) | 27.7 | 1.15 (d, 6.9 Hz, 3H) | 19.4 | 1.25 (m, 1H)<br>1.50 (m, 1H) | 27.7 | 1.10 (m, 1H)<br>1.35 (m, 1H) | 27.2 | 1.25 (m, 1H)<br>1.47 (m, 1H) | 27.7 | 1.22 (m, 1H)<br>1.46 (m, 1H) | 27.5 |
| 15d | 0.55 (t, 7.4 Hz, 3H) | 11.6 | 0.85 (d, 6.9 Hz, 3H) | 19.3 | 0.55 (t, 7.4 Hz, 3H) | 11.6 | 0.35 (t, 7.5 Hz, 3H) | 11.1 | 0.54 (t, 7.4 Hz, 3H) | 11.6 | 0.54 (t, 7.5 Hz, 3H) | 11.3 |
| 15e | 1.13 (d, 7.0 Hz, 3H) | 17.8 |  |  | 1.16 (d, 7.0 Hz, 3H) | 17.7 | 1.12 (d, 7.0 Hz, 3H) | 17.2 | 1.15 (d, 7.0 Hz, 3H) | 17.7 | 1.13 (d, 7.3 Hz, 3H) | 17.4 |

Compound **6** was assigned as (1*R*,2*S*,4*R*,5*S*,6*R*,7*S*,8*R*,9*S*,10*S*)-6*β*-acetoxy-2*α*,8*α*-di-(5-carboxy-*N*-methyl-3-pyridoxy)-9*α*,15-di-(2-methylbutanoyloxy)-dihydro-*β*-agarofuran. It presented typical ^1^H-NMR signals of a dihydro-*β*-agarofuran scaffold, with a molecular formula of C_41_H_55_N_2_O_14_ for [M+H]^+^ of *m/z* 799.3649. Position one carried a hydroxyl group as indicated by the chemical shift of H-1 at δ_H_ 4.39. The other positions (2, 8, 9, and 15) were esterified by two 5-carboxy-*N*-methyl-3-pyridone and two 2-methylbutanoate. These latter were positioned in C-9 and C-15 due to the HMBC correlations from H-9b at δ_H_ 2.24, H_2_-9c at δ_H_ 1.37 and 1.68, H_3_-9e at δ_H_ 1.03, and H-9 at δ_H_ 5.64 to C-9a δ_C_ 176.6 and from H-15b at δ_H_ 2.37, H_2_-15c at δ_H_ 1.24 and 1.48, H_3_-15e at δ_H_ 1.13, and H-15’ at δ_H_ 5.41 to C-15a δ_C_ 179.0. The 5-carboxy-*N*-methyl-3-pyridones were thus in C-2 and C-8. The ROESY correlation from the aromatic protons H-2c (δ_H_ 8.61) and H-2f (δ_H_ 8.07) to H_2_-15 (δ_H_ 5.41 and 4.37) placed this 5-carboxy-*N*-methyl-3-pyridone in C-2 while the correlations from H-8c (δ_H_ 8.89) to H-6 (δ_H_ 6.74) placed the second one in C-8. The ROESY correlation from H-15 to H-6 and H_3_-14 indicated that H_2_-15, the two 5-carboxy-*N*-methyl-3-pyridone in C-2 and C-8, H_3_-14, and H-6 were on the same side of the molecule. On the other side, the acetate in C-6 (H_3_-6b) correlated with H_3_-13, H_3_-12 with H-1, and H-1 with H-9. Altogether these data indicated that the relative configuration should be 1*α*, 2*α*, 4*α*, 5*β*, 6*β*, 7*β*, 8*α*, 9*α*, 15*α*.

Compound **7** was obtained as an amorphous white powder with a molecular formula of C_42_H_55_N_2_O_15_ for [M+H]^+^ of *m/z* 827.3598. The 1D and 2D NMR data displayed a significant resemblance with **6**. One extra quaternary carbon at δ_C_ 171.4 was observed, belonging to an acetyl group fixed in position C-1 (δ_C_ 77.9), corroborated by the HMBC correlation with H-1 (δ_H_ 5.70). Positions C-2, C-8, and C-9 were substituted with the same groups as **6**. However, protons in C-15 (δ_H_ 4.24 and 5.46) correlated in the HMBC spectrum with a carbonyl group at δ_C_ 179.0, coupled to a methine (δ_H_ 2.58), and two methyl doublets (δ_H_ 0.85 and 1.15), corresponding to a methylpropanoate system. The ROESY spectrum presented the same correlations as **6**. After calculation of the ECD spectrum and comparison with the experimental, the compound **7** was assigned as (1*R*,2*S*,4*R*,5*S*,6*R*,7*S*,8*R*,9*S*,10*S*)-1*α*,6*β*-diacetoxy-2*α*,8*α*-di-(5-carboxy-*N*-methyl-3-pyridoxy)-15-*iso*-butanoyloxy-9*α*-(2-methylbutanoyloxy)-dihydro-*β*-agarofuran.

Compound **8** was obtained as a white amorphous powder with a molecular formula of C_43_H_57_N_2_O_15_ for [M+H]^+^ of *m/z* 841.3736. The NMR data showed to be closely related to **7**. However, in position C-15 (δ_C_ 62.6), the substituent corresponds to a 2-methylbutanoate as in **6**. The absolute configuration was assigned by comparison of the calculated ECD based on the relative configuration proposed as 1*β*, 2*β*, 8*β*, and 9*β* due to the observed ROESY correlations. Compound **8** was assigned as (1*R*,2*S*,4*R*,5*S*,6*R*,7*S*,8*R*,9*S*,10*S*)-1*α*,6*β*-diacetoxy-2*α*,8*α*-di-(5-carboxy-*N*-methyl-3-pyridoxy)-9*α*,15-di-(2-methylbutanoyloxy)-dihydro-*β*-agarofuran.

Compound **9** was assigned as (1*R*,2*S*,4*R*,5*S*,6*R*,7*S*,8*R*,9*S*,10*S*)-1*α*,6*β*-diacetoxy-2*α*-(5-carboxy-*N*-methyl-3-pyridoxy)-9*α*,15-di-(2-methylbutanoyloxy)-8*α*-nicotinoyloxydihydro-*β*-agarofuran, with a molecular formula C_42_H_53_N_2_O_11_ for [M+H]^+^ of *m/z* 811.3668. The major difference with **8** was the presence of a nicotinate moiety at δ_H_ 9.40 (1H, d, *J* = 1.9 Hz, H-8c), 8.82 (1H, dd, *J* = 5.0, 1.9 Hz, H-8d), 8.57 (1H, dt, J = 7.9, 1.9 Hz, H-8f), and 7.65 (1H, dd, *J* = 7.9, 5.0 Hz, H-8e) instead of one of the 5-carboxy-*N*-methyl-3-pyridone. The 5-carboxy-*N*-methyl-3-pyridone was positioned in C-2 due to the ROESY correlation of H-15’ with H-2c. The nicotinate was thus placed in C-8. The ROESY correlations remained the same as other alatol-type compounds. Calculations of the ECD spectrum were done to define the absolute configuration.

Compound **10**, C_42_H_55_N_2_O_15_, calculated for [M+H]^+^ of *m/z* 827.3595, presented the same formula and mass as **7**. The same core structure was proposed, but substituents in C-9 (δ_C_ 71.9) and C-15 (δ_C_ 62.6) were inverted. H-9 (δ_H_ 5.52) had an HMBC correlation with carbon at δ_C_ 176.3 which was connected to an *iso*-propyl system [δ_H_ 2.45 (1H, m, H-9b); 1.07 (3H, d, *J* = 7.0 Hz, H_3_-9d); 1.09 (3H, d, *J* = 7.1 Hz, H_3_-9c). H-15’/H-15’’ correlated with a carbon at δ_C_ 178.7, which was connected to an *iso*-butyl system [δ_H_ 2.38 (1H, m, H-15b); 1.25 (1H, m, H-15c’’), 1.47 (1H, m, H-15c’); 1.15 (3H, d, *J* = 7.0 Hz, H3-15e); 0.54 (3H, t, *J* = 7.4 Hz, H_3_-15d)]). The relative configuration was the same as **7** (1*α*, 2*α*, 8*α*,9*α*). Compound **10** was assigned as (1*R*,2*S*,4*R*,5*S*,6*R*,7*S*,8*R*,9*S*,10*S*)-1*α*,6*β*-diacetoxy-9*α*-*iso*-butanoyloxy-2*α*,8*α*-di-(5-carboxy-*N*-methyl-3-pyridoxy)-15-methylbutanoyloxydihydro-*β*-agarofuran.

Compound **11** was assigned as (1*R*,2*S*,4*R*,5*S*,6*R*,7*S*,8*R*,9*S*,10*S*)-6*β*-diacetoxy-9*α*-*iso*-butanoyloxy-2*α*,8*α*-di-(5-carboxy-*N*-methyl-3-pyridoxy)-1*α*-hydroxy-15-methylbutanoyloxydihydro-*β*-agarofuran, with a molecular formula of C_40_H_53_N_2_O_14_, calculated for [M+H]^+^ of *m/z* 785.3511. It presented the same substitution pattern as **10**, but position C-1 (δ_C_ 75.6) had a proton signal at a higher field (δ_H_ 4.38), suggesting the presence of a free hydroxyl group. The relative configuration was the same as the rest of the molecules from this group and the absolute assigned configuration was checked by ECD comparison between the calculated and experimental spectra.

The third group of dihydro-*β*-agarofuran structures was composed of 2 new euonymol-type structures (1**2** and **13**, see **Table 5**). They were oxygenated in all 7 possible positions.

**Table 5.** ^1^H and ^13^C NMR spectroscopic data for compounds **12**-**13** (δ in ppm, *J* in Hz).

| Compound **12** |  |  | Compound **13** |  |
| --- | --- | --- | --- | --- |
| No | $\delta_{\text{H}}$ (Multiplicity, $J$, nH) | $\delta_{\text{C}}$ | $\delta_{\text{H}}$ (Multiplicity, $J$, nH) | $\delta_{\text{C}}$ |
| 1 | 5.91 (d, 4.2 Hz, 1H) | 74.9 | 5.92 (d, 4.3 Hz, 1H) | 75.2 |
| 2 | 5.52 (m, 1H) | 71.3 | 5.53 (m, 1H) | 71.6 |
| 3 | 4.87 (m, 1H) | 75.8 | 4.87 (m, 1H) | 76.1 |
| 4 | 2.56 (m, 1H) | 38.0 | 2.56 (qt, 8.0, 1.0 Hz, 1H) | 38.2 |
| 5 | - | 90.3 | - | 90.6 |
| 6 | 6.58 (d, 1.1 Hz, 1H) | 76.5 | 6.55 (d, 1.0 Hz, 1H) | 76.9 |
| 7 | 2.50 (dd, 3.8, 1.0 Hz, 1H) | 53.3 | 2.50 (dd, 3.7, 1.0 Hz, 1H) | 53.6 |
| 8 | 5.52 (m, 1H) | 70.6 | 5.53 (m, 1H) | 70.8 |
| 9 | 5.49 (d, 6.5 Hz, 1H) | 71.9 | 5.49 (d, 6.6 Hz, 1H) | 72.1 |
| 10 | - | 52.0 | - | 52.1 |
| 11 | - | 82.5 | - | 82.7 |
| 12 | 1.53 (s, 3H) | 24.6 | 1.53 (s, 3H) | 24.9 |
| 13 | 1.44 (s, 3H) | 30.2 | 1.43 (s, 3H) | 30.6 |
| 14 | 1.21 (d, 7.9 Hz, 3H) | 14.9 | 1.20 (d, 7.9 Hz, 3H) | 15.2 |
| 15 | 4.28 (d, 13.2 Hz, 1H)<br>5.36 (d, 13.2 Hz, 1H) | 61.9 | 4.22 (d, 13.3 Hz, 1H)<br>5.44 (d, 13.3 Hz, 1H) | 62.1 |
| **R1** |  |  |  |  |
| 1a | - | 170.8 | - | 171.1 |
| 1b | 1.75 (s, 3H) | 20.2 | 1.75 (s, 3H) | 20.5 |
| **R2** |  |  |  |  |
| 2a | - | 163.9 | - | 164.1 |
| 2b | - | 109.9 | - | 110.3 |
| 2c | 8.55 (d, 2.6 Hz, 1H) | 145.8 | 8.58 (d, 2.5 Hz, 1H) | 146.1 |
| 2d | - | 165.0 | - | 165.3 |
| 2e | 6.58 (d, 9.5 Hz, 1H) | 119.6 | 6.58 (d, 9.5 Hz, 1H) | 119.9 |
| 2f | 8.00 (dd, 9.5, 2.6 Hz, 1H) | 140.4 | 8.02 (dd, 9.5, 2.5 Hz, 1H) | 140.7 |
| 2g | 3.71 (d, 1.7 Hz, 3H) | 38.3 | 3.71 (s, 3H) | 38.6 |
| **R3** |  |  |  |  |
| 3a | - | 171.3 | - | 171.5 |
| 3b | 2.12 (s, 3H) | 20.7 | 2.12 (s, 3H) | 21.2 |
| **6-Ac** |  |  |  |  |
| 6a | - | 171.1 | - | 171.2 |
| 6b | 2.12 (s, 3H) | 20.7 | 2.12 (s, 3H) | 21.1 |
| **R4** |  |  |  |  |
| 8a | - | 171.5 | - | 171.7 |
| 8b | 2.11 (s, 3H) | 20.7 | 2.13 (s, 3H) | 21.4 |
| **R5** |  |  |  |  |
| 9a | - | 167.0 | - | 167.3 |
| 9b | - | 128.9 | - | 129.2 |
| 9c | 6.83 (qq, 6.9, 1.3 Hz, 1H) | 140.0 | 6.83 (qq, 7.3, 1.6 Hz, 1H) | 140.2 |
| 9d | 1.78 (dq, 6.9, 1.3 Hz, 3H) | 14.1 | 1.78 (m, 3H) | 14.4 |
| 9e | 1.77 (p, 1.3 Hz, 3H) | 11.6 | 1.78 (m, 3H) | 12.1 |
| **R6** |  |  |  |  |
| 15a | - | 178.9 | - | 178.9 |
| 15b | 2.90 (hept, 6.9 Hz, 1H) | 34.8 | 2.74 (h, 7.0 Hz, 1H) | 42.3 |
| 15c | 1.26 (d, 6.9 Hz, 3H) | 19.3 | 1.58 (ddd, 13.8, 7.5, 6.4 Hz, 1H)<br>1.80 (m, 1H) | 27.9 |
| 15d | 1.24 (d, 6.9 Hz, 3H) | 19.3 | 0.97 (t, 7.4 Hz, 3H) | 12.1 |
|  |  |  | 1.25 (d, 7.0 Hz, 3H) | 17.4 |

Compound **12** was assigned as (1*R*,2*S*,3*S*,4*R*,5*S*,6*R*,7*S*,8*R*,9*S*,10*S*)-1*α*,3*β*,6*β*,8*α*-tetraacetoxy-15-*iso*-butanoyloxy-2*α*-(5-carboxy-*N*-methyl-3-pyridoxy)-9*α*-tigloyloxydihydro-*β*-agarofuran, based on the NMR data. It presented a molecular formula of C_39_H_52_NO_16_, calculated for [M+H]^+^ of *m/z* 790.3289. In the HMBC spectrum, six carbonyls, presumably esters, were observed at δ_C_ 178.9, 171.5, 171.3, 170.8, 167.0, and 163.9, in addition to the acetyl fixed in C-6 (δ_C_ 171.1). The HMBC correlations from H-1 (δ_H_ 5.91) and H_3_-1b (δ_H_ 1.75) to C-1a (δ_C_ 170.8) positioned an acetate in C-1, from H-2 (δ_H_ 5.52), H-2c (δ_H_ 8.55), and H-2f (δ_H_ 8.00) to C-2a (δ_C_ 163.9) positioned a 5-carboxy-*N*-methyl-pyridone in C-2, from H-3 (δ_H_ 4.87) and H_3_-3b (δ_H_ 2.12) to C-3a (δ_C_ 171.3) positioned an acetate in C-3, from H-8 (δ_H_ 5.52) and H_3_-8b (δ_H_ 2.11) to C-8a (δ_C_ 171.5) positioned an acetate in C-8, from H-9 (δ_H_ 5.49), H-9c (δ_H_ 6.83) and H_3_-9e (δ_H_ 1.77) to C-9a (δ_C_ 167.0) positioned a tiggeloyl in C-1, and from H_2_-15 (δ_H_ 4.28 and 5.36), H-15b (δ_H_ 2.90), H_3_-15c (δ_H_ 1.26) and H_3_-15d (δ_H_ 1.24) to C-15a (δ_C_ 178.9) positioned an *iso*-butanoyl in C-15. The ROESY correlations showed that the configuration of the ester groups in C-1, C-2, C-8, and C-9 was the same as the alatol-type structures (**6**-**11**). The ROESY between H-3 and H_3_-14 indicated that the acetate in C-3 and methyl 14 were in a *trans* configuration. This relative configuration was corroborated after a comparison of the experimental and calculated ECD spectra.

Compound **13**, was obtained as an amorphous powder, giving a [M+H]^+^ of *m/z* 804.3445 with a molecular formula of C_40_H_54_NO_16_. The mass difference of 14 observed between itself and **12**, suggested the presence of an extra CH_2_. This was corroborated due to the close resemblance of all the 1D and 2D NMR, except for the substituent in position C-15 (δ_C_ 62.1), which fitted with a 2-methylbutanoate moiety. The absolute configuration was corroborated by ECD calculation, using the relative configuration proposed by the ROESY spectrum. Thus, compound **13** was assigned as (1*R*,2*S*,3*S*,4*R*,5*S*,6*R*,7*S*,8*R*,9*S*,10*S*)-1*α*,3*β*,6*β*,8*α*-tetraacetoxy-2*α*-(5-carboxy-*N*-methyl-3-pyridoxy)-15-(2-methylbutanoyloxy)-9*α*-tigloyloxydihydro-*β*-agarofuran.

Based on the FC values and highlighted ions, the filtering results, and the *de novo* structural identifications, the chemical class proposed by Sirius-Canopus was confirmed, as well as the potential that *Inventa* holds to speed the discovery of novel NPs.

## 4 Conclusion

As explained throughout the article, prioritization of library extracts is difficult, multifactorial and time consuming. For this reason, the development of comprehensive prioritization pipelines combining the results of several bioinformatics tools is necessary to speed up and streamline extract selection for further in-depth phytochemical study. In this context, we propose Inventa, an innovative computational tool capable of combining various level of information (specificity, originality, annotations) from state-of-the-art bioinformatics programs, to highlight and prioritize extracts based on the possibility of finding structurally novel NPs. *Inventa* can be modulated according to the study parameters, and run locally or remotely via a web-based <u>*Binder*</u> notebook. The application of *Inventa* on a set of plant extracts showed how it can identify extracts where new compounds have high probability to be discovered. As a proof of concept, Inventa succeeded in the prioritization of the *Pristimera indic*a roots extract among a set of seventy-six extracts from the Celastraceae family. An *in-depth* phytochemical investigation of this extract led to the isolation and *de novo* structural identification of thirteen new *β*-agarofuran sesquiterpene compounds. Five of them presented a new 9-oxodihydro-*β*-agarofuran base scaffold. This example illustrates how *Inventa* can speed up the discovery of original NPs.

It is expected that in a near future Inventa, which allows prioritization of extract from large collections can be complemented by other tools, such as <u>FERMO</u>, under development (Zdouc. M, Medema. M, van der Hooft. JJ), which will allow in-detail analysis and visualization for a particular extract. Collaboration efforts are in place to make them compatible and enhance their applicability.

## Data and software availability

*Inventa* can be found on https://github.com/luigiquiros/inventa (https://luigiquiros.github.io/inventa/). All .RAW (Thermo), .mzML datafiles (positive ionization mode) and metadata are available on the Massive MSV000087970, [<u>doi:10.25345/C5PJ9N</u>]. An interactive visualization can be displayed using the <u>GNPS Dashboard</u>.

## Author’s contributions

LMQG and P-MA conceptualized the study. LMQG performed the data acquisition, analysis, and visualization. LMQG, L-FN, and AG developed the python scripts for the package. LMQG, EFQ, and LM performed the isolation and structural characterization. LMQG wrote the original manuscript. LMQG, P-MA, L-FN, BD, AG, AR, EFQ, LM, and J-LW revised the manuscript. All authors read, reviewed, and approved the paper.

## Acknowledgments

The authors extend a special thanks to Bruno David and Green Mission Pierre Fabre, Pierre Fabre Research Institute, Toulouse, France for the establishment of the collaboration and for sharing the great extract collection. The School of Pharmaceutical Sciences of the University of Geneva (J-LW) is thankful to the Swiss National Science Foundation for the support in the acquisition of the NMR 600 MHz (SNF: Equipment grant 316030_164095). LMQG is thankful for the scholarship (N 214171025) from Ministerio de Ciencia, Tecnología y Telecomunicaciones, MICITT, Costa Rica.

## Supplementary Material

The Supplementary Material for this article can be found online at:

