## Supplementary information for "*Inventa*: a computational tool to discover chemical novelty in natural extracts libraries"

### **Supplementary Table S**1. Glossary of terms and default values

| **Parameter** | **Default value** | **Description** |
|---|---|---|
| annot_gnps_df | | raw annotations attributes file downloaded directly from GNPS |
| annot_is_df | | annotation status table from in silico ISDB dereplication |
| annot_sirius_df | | annotation status table from SIRIUS dereplication |
| annotation_df | | combined annotation status table from all the dereplication results used |
| annotation_preference | 0 | Only Annotated features: '1' or Only Not annotated features: '0' #userdefined |
| ATTRIBUTE_Sppart' | | if needed this column is generated by merging the species and organe columns #userdefined |
| canopus_npc_df | | chemical classes retrieved from the Sirius dereplication results |
| canopus_npc_summary_filename | ../data/yourfilenamegoeshere.tsv' | path where the SIRIUS-CANOPUS results file is placed |
| CC | | Class component results table |
| CC_component | True (False) | CC will be calculated |
| col_id_unique | | unique identifier for the samples (filename, part, etc) #userdefined |
| correlation_groups_df | | ion identity annotation network number attributes |
| data_process_origin | MZmine3' / 'MZmine2' | version of MZmine used to treat the data #userdefined |
| df_annotations | | raw annotations file downloaded directly from GNPS |

| | | |
|---|---|---|
| family_colum | ATTRIBUTE_Family' (yourfamilycolumgoeshere) | column header for the family name #userdefined |
| FC | | Feature component results table |
| filename_header | filename' (yourfilenamecolumgoeshere) | column header for the species name #userdefined |
| filtered_F | | the residual number of features different from zero in the quant_df after filtering |
| filtered_full_df | | filtered (without QC and blanks) full table |
| filtered_metadata_df | | filtered (without QC and blanks) metadata table |
| FS | | Feature Specificity, ratio of Total_SF/filtered_F |
| full_df | | quantitative and metadata combined table |
| genus_colum | ATTRIBUTE_Genus' (yourgenuscolumgoeshere) | column header for the genus name #userdefined |
| gnps_annotations_consolidated | | processed annotations file from GNPS |
| gnps_annotations_filtered | | processed annotations file from GNPS after chemical descriptors consolidation |
| IF | | Isolation Forest |
| initial_F | | the initial number of features different from zero in the quant_df |
| ionisation_mode | pos' | ionisation mode according to experimental conditions 'pos' or 'neg' #userdefined |
| isdb_annotations | True (False) | the tima_results_filename will be considered in the calculations #userdefined |
| job_id | yourjobidgoeshere | GNPS Job ID #userdefined |
| LC | | Literature component results table |
| LC_component | True (False) | LC will be calculated |
| LOF | | Local Outlier Factor |
| max_comp_reported_f | 500 | max number of compounds reported at genus level,more than this value, the plant is considered less interesting #userdefined |
| max_comp_reported_g | 50 | max number of compounds reported at genus level,more than this value, the plant is considered less interesting #userdefined |
| max_comp_reported_sp | 20 | max number of compounds reported at species level, more than this value, the plant is considered less interesting #userdefined |

| | | |
|---|---|---|
| max_parts_per_organism | 4 (your max occurrence goes here) | max recurrence of the same organism species (for example: 5 samples, same species but different plant part) #userdefined |
| max_ppm_error | 5 | min error in ppm to consider an annotation valable #userdefined |
| max_spec_charge | 2 | maximum charge allowed #userdefined |
| metadata_df | | formated metadata table |
| metadata_filename | ../data/yourfilenamegoeshere.tsv' | path where the metadata is placed #userdefined |
| metric_df | | memo matrix table |
| MF_prediction_ratio | | Ratio of SNAGQMFF/filtered_F |
| min_class_confidence | 0.8 | cut-off filter for considering a sirius class valable. It is used in combination with min_recurrence #userdefined |
| min_ConfidenceScore | 0.25 | cut-off filter for considering a sirius annotation valable. '0.0' as default. #userdefined |
| min_cosine | 0.6 | min cosine score to consider an annotation valable #userdefined |
| min_recurrence | 5 | minimum recurrence of a chemical class to consider it acceptable #userdefined |
| min_score_final | 0.3 | cut-off filter for considering an isdb annotation valable. You must be extremenly carefull with this parameter, '0.0' as default. #userdefined |
| min_specificity | 0.9 | minimum value of relative area (0 to 1) to consider a feature specific or not #userdefined |
| min_ZodiacScore | 0.9 | cut-off filter for considering a sirius annotation valable. It is used in combination with min_ConfidenceScore. #userdefined |
| multiple_organism_parts | True (False) | True: the specificity is going to be considered as the sum of the 'max_parts_per_organism' shared in the samples. #userdefined |
| OCSVM | | One Class Support Vector Machine |
| organe_colum | ATTRIBUTE_Organ' (yourorgancolumgoeshere) | column header for the organe name (this column could be used to indicate different culture media, extraction solvents, etc.) #userdefined |
| PR | | Priority Rank results table |
| quant_df | | sample-wise normalized quantitative table |
| quantitative_data_filename | ../data/yourfilenamegoeshere.tsv' | path where the quantitative table is placed #userdefined |
| reduced_df | | col_id_unique restricted full df |
| Reported_comp_Family | | Number of compounds reported in the family in Lotus |

| | | |
|---|---|---|
| Reported_comp_Genus | | Number of compounds reported in the genus in Lotus |
| Reported_comp_Species | | Number of compounds reported in the species in Lotus |
| SC | | Similarity component results table |
| SC_component | True (False) | SC will be calculated |
| shared_peaks | 4 | min number of shared peaks between the MS2 experimental and MS2 from the database, to consider an annotation valable #userdefined |
| sirius_annotations | True (False) | the sirius_annotations_filename will be considered in the calculations |
| sirius_annotations_filename | ../data/yourfilenamegoeshere.tsv' | path where the SIRIUS results file is placed #userdefined |
| species_colum | ATTRIBUTE_Species' (yourspeciescolumgoeshere) | column header for the species name #userdefined |
| tima_results_filename | ../data/yourfilenamegoeshere.tsv' | path where the ISDB reponderated file is placed #userdefined |
| Total_SF | | the total number of Specific (S) features (F) |
| Total_SNA_GQMFF | | the total number of Specific (S) non-annotated (NA) features (F) with a molecular formula of good quality (GQMF) |
| Total_SNAF | | the total number of Specific (S) non-annotated (NA) features (F) |
| use_ion_identity | True (or False) | if True, the ion identity results will be used #userdefined |
| vectorized_data_filename | ../data/yourfilenamegoeshere.tsv' | path where the MEMO matrix is placed #userdefined |
| w1 | 1 | weight for the FC in the final PR result #userdefined |
| w2 | 1 | weight for the LC in the final PR result #userdefined |
| w3 | 1 | weight for the SC in the final PR result #userdefined |
| w4 | 1 | weight for the CC in the final PR result #userdefined |
| wf | 1 | weight for the family level in the LC component #userdefined |
| wg | 1 | weight for the genus level in the LC component #userdefined |
| ws | 1 | weight for the species level in the LC component #userdefined |

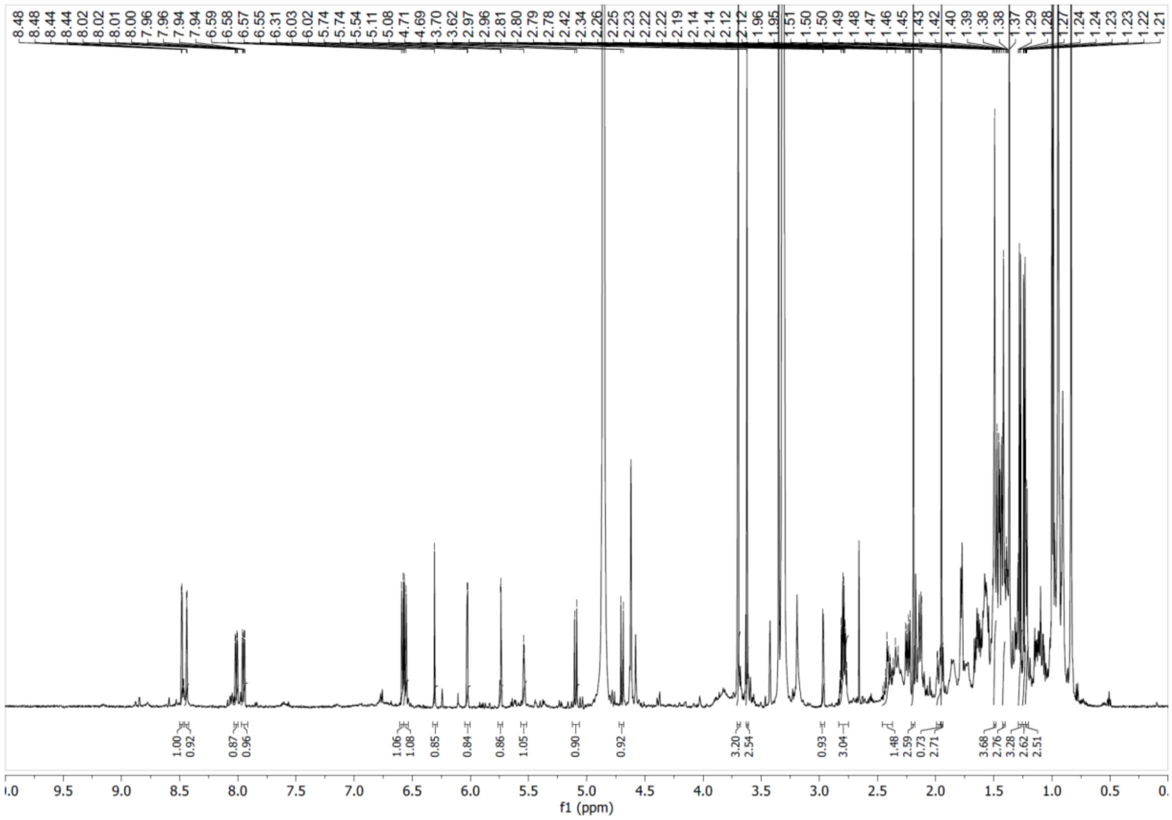


**Supplementary Figure S1**. $^{1}H$ NMR spectrum of compound **1** in $CD_3OD$ at 600 MHz.

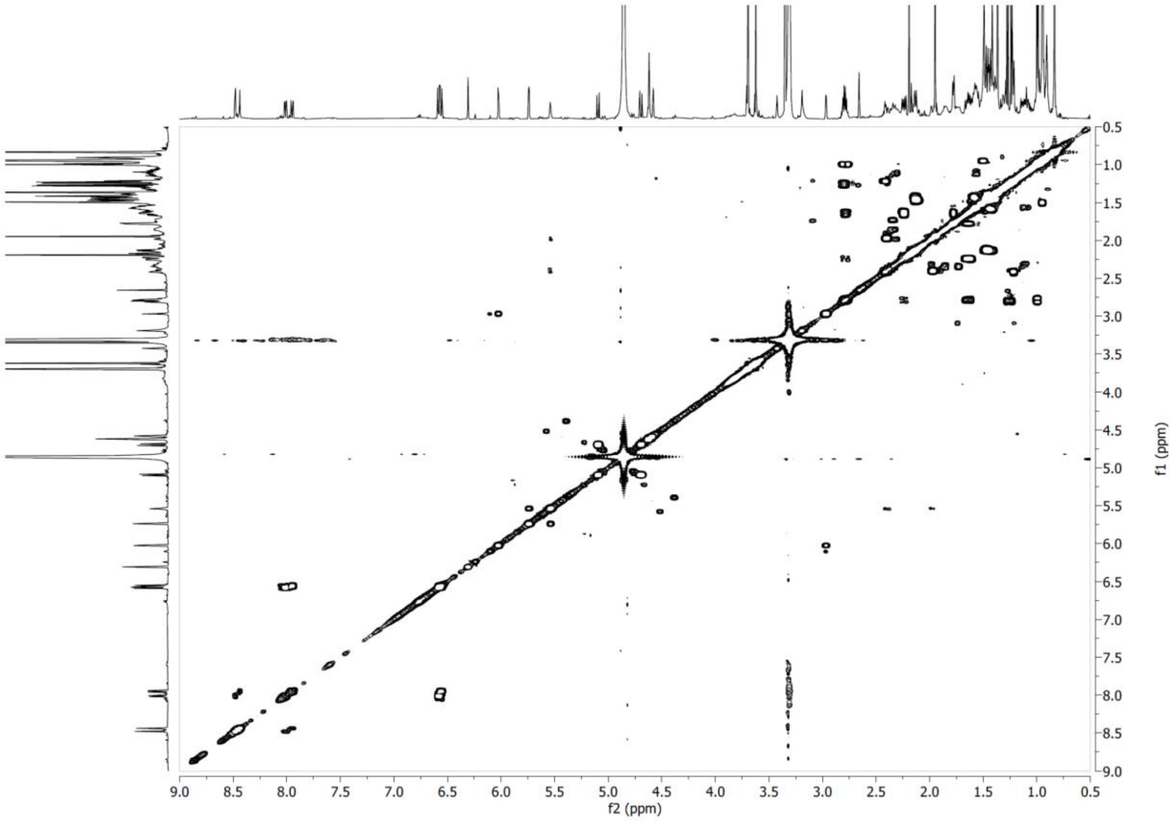


**Supplementary Figure S2**. COSY NMR spectrum of compound **1** in $CD_3OD$ at 600 MHz.

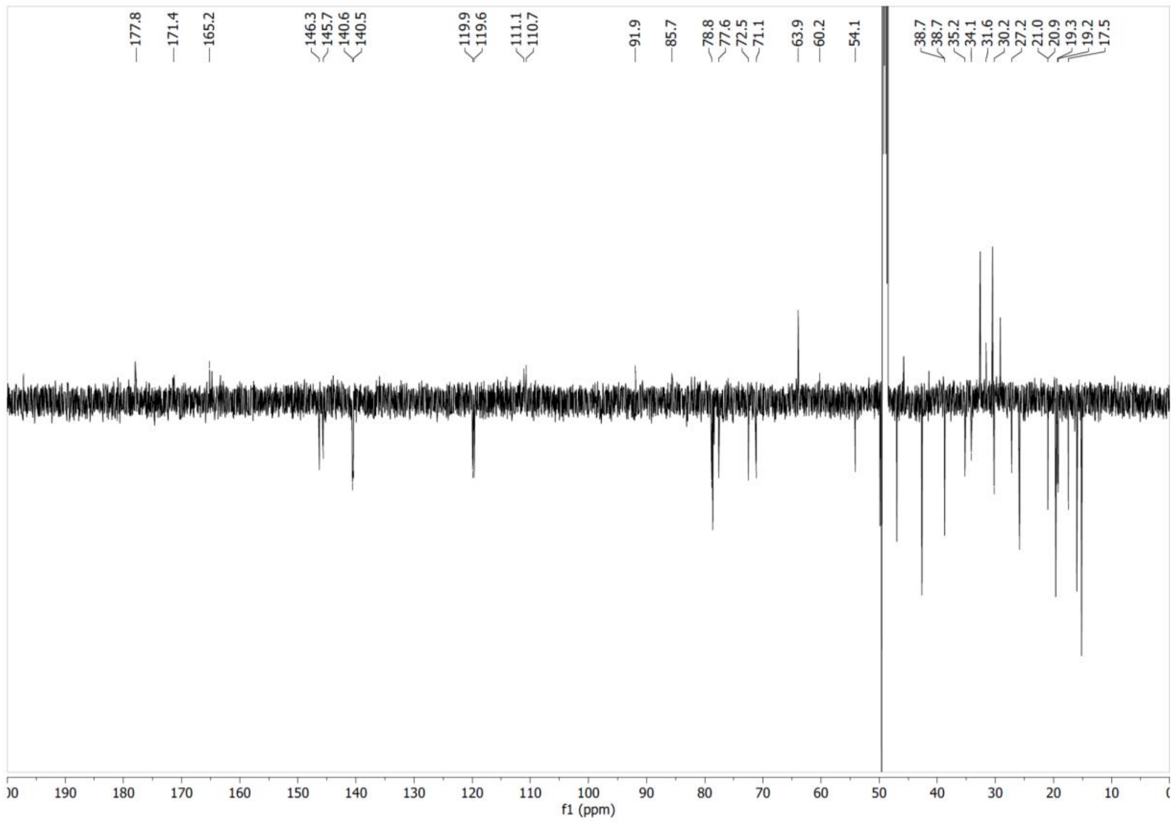


**Supplementary Figure S3**. $^{13}C$ NMR spectrum of compound **1** in $CD_3OD$ at 151 MHz.

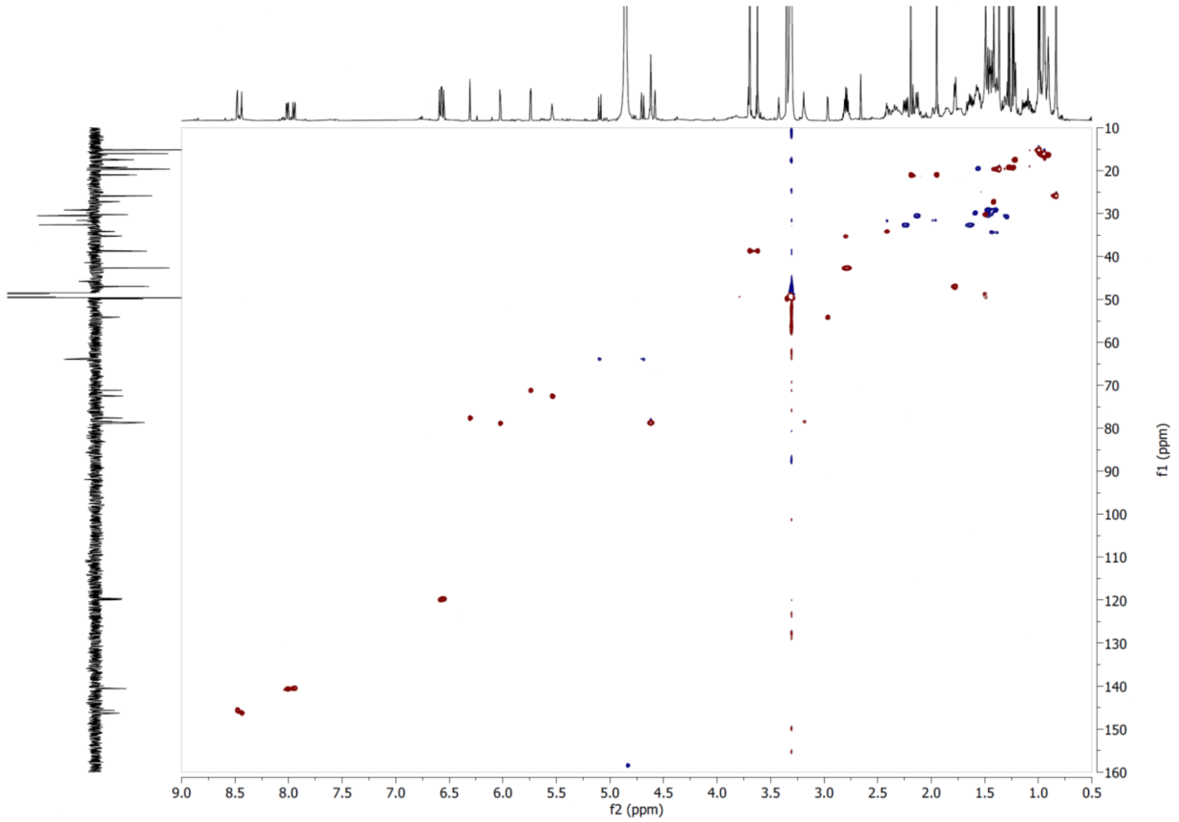


**Supplementary Figure S4**. HSQC NMR spectrum of compound **1** in $CD_3OD$.

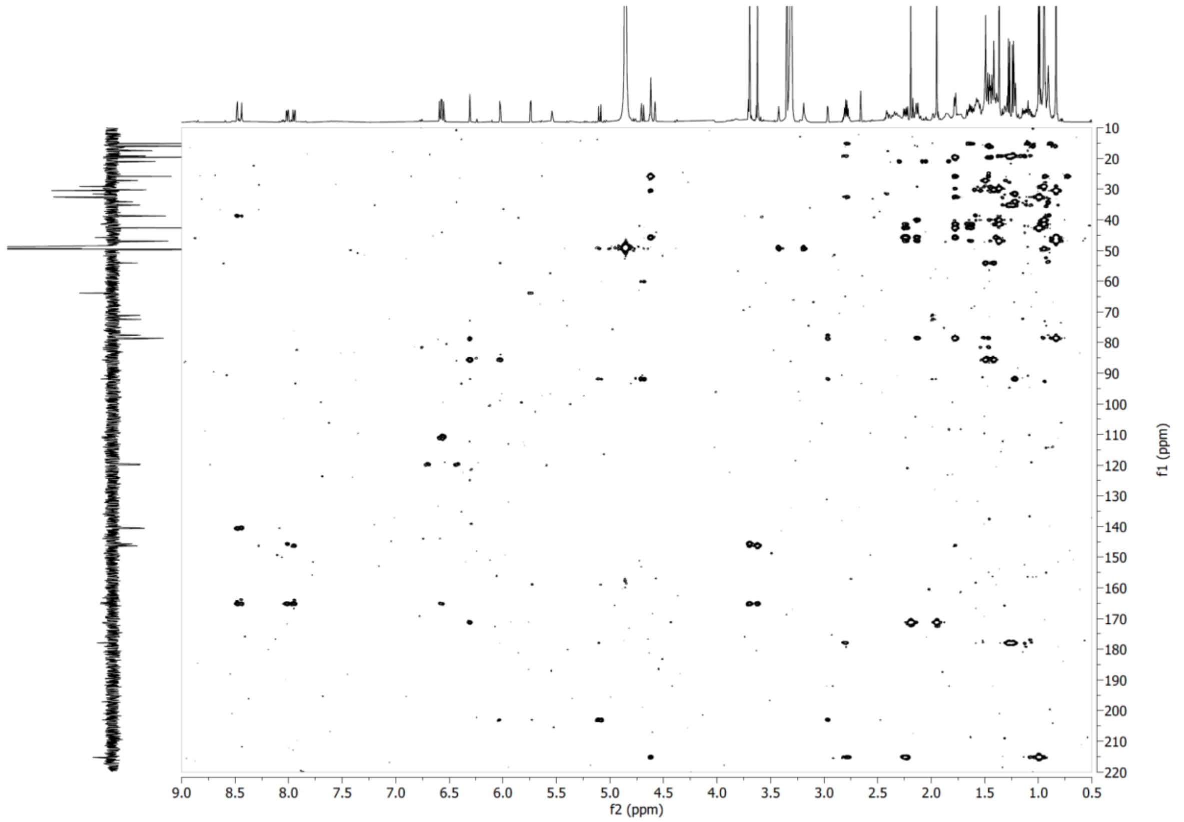


**Supplementary Figure S5**. HMBC NMR spectrum of compound **1** in $CD_3OD$.

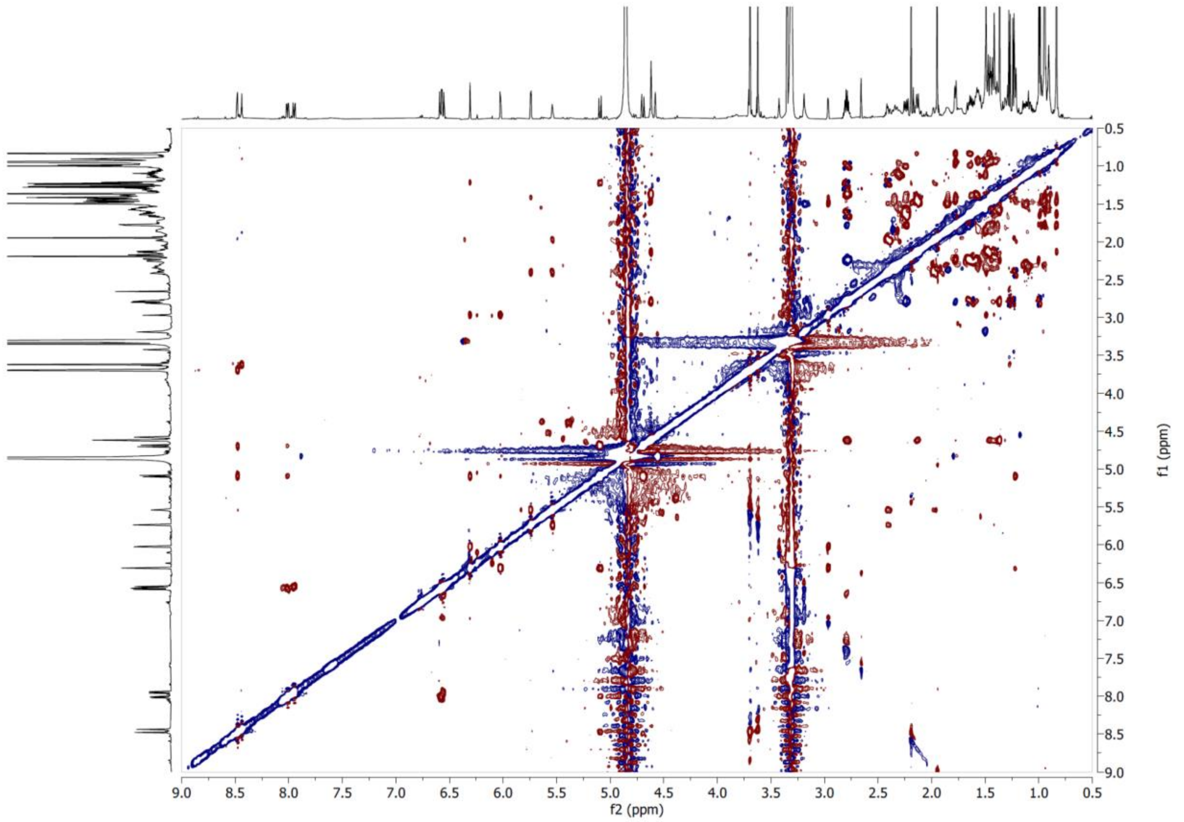


**Supplementary Figure S6**. ROESY NMR spectrum of compound **1** in $CD_3OD$ at 600 MHz.

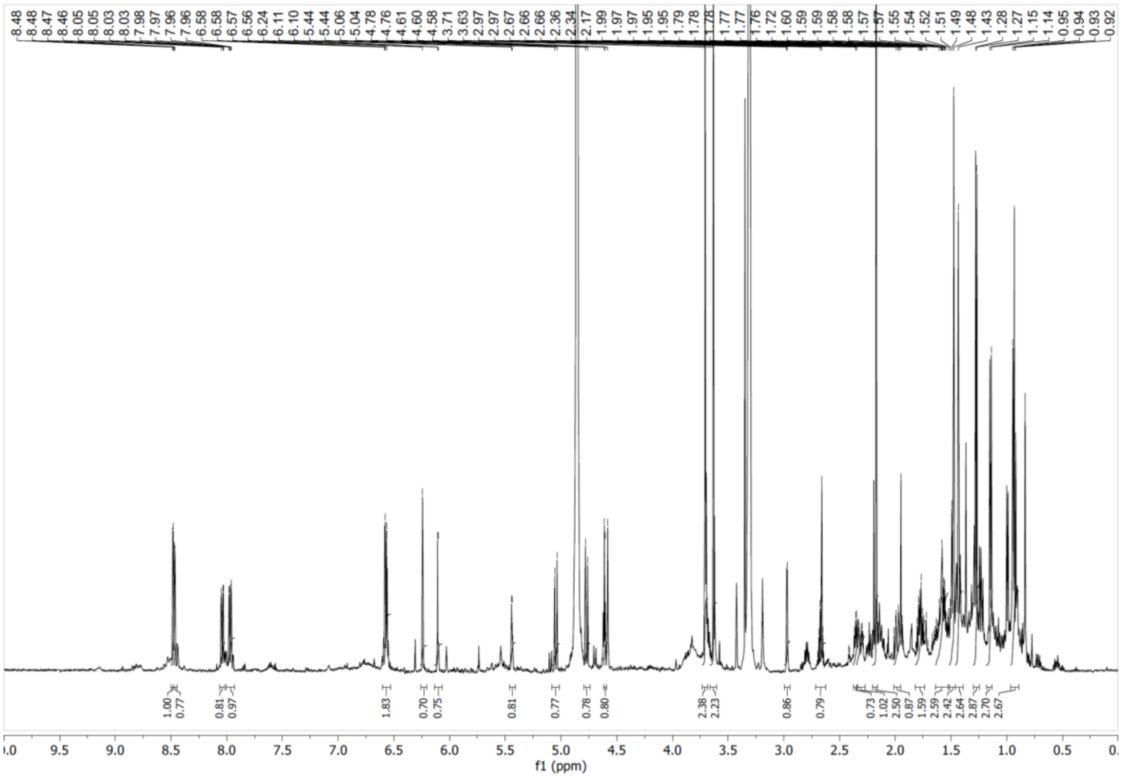

**Supplementary Figure S7**. [1]H NMR spectrum of compound **2** in $CD_3OD$ at 600 MHz.

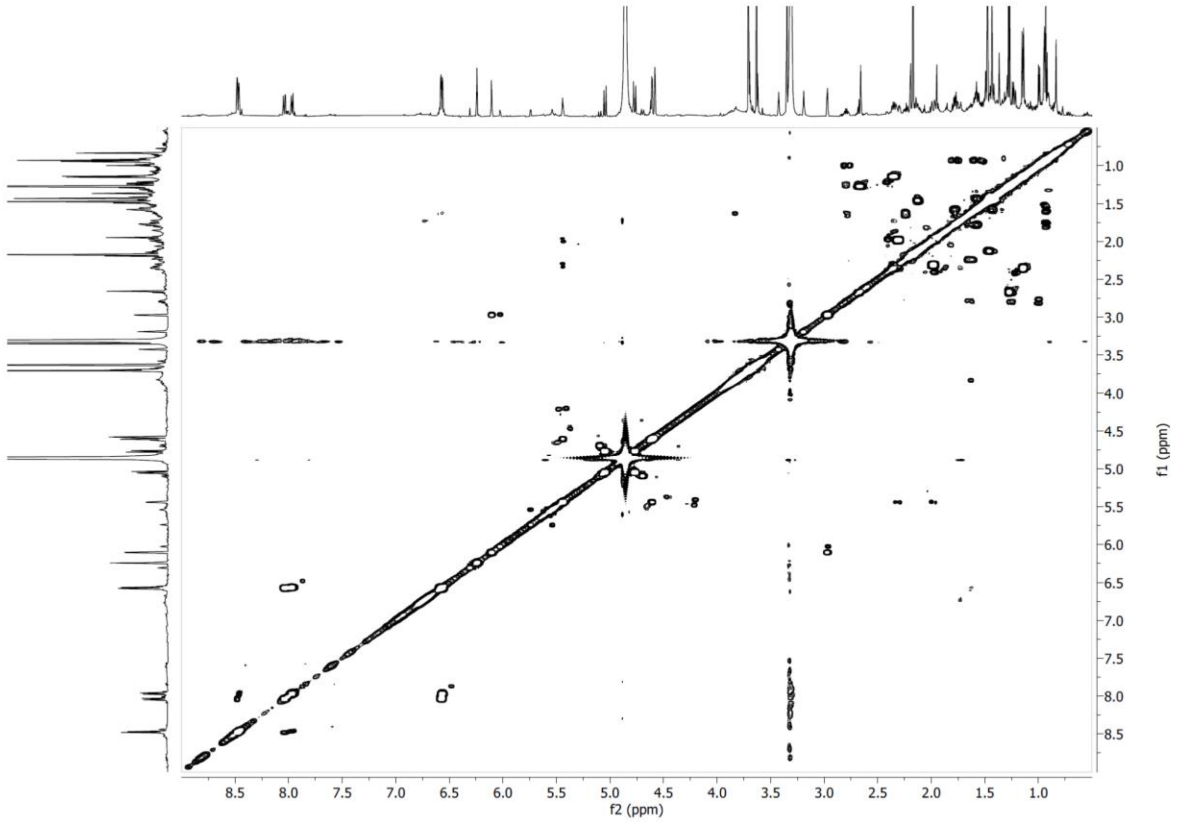


**Supplementary Figure S8**. COSY NMR spectrum of compound **2** in $CD_3OD$ at 600 MHz.

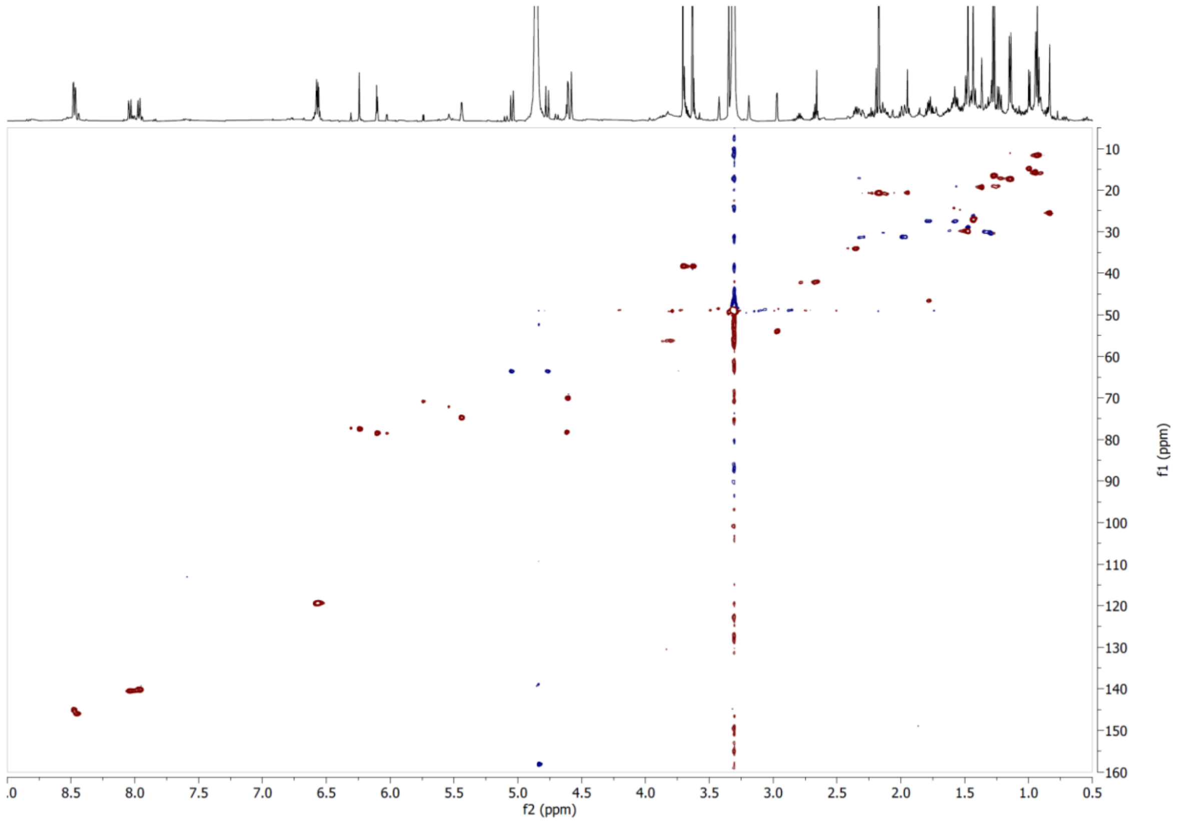


**Supplementary Figure S9**. HSQC NMR spectrum of compound **2** in $CD_3OD$.

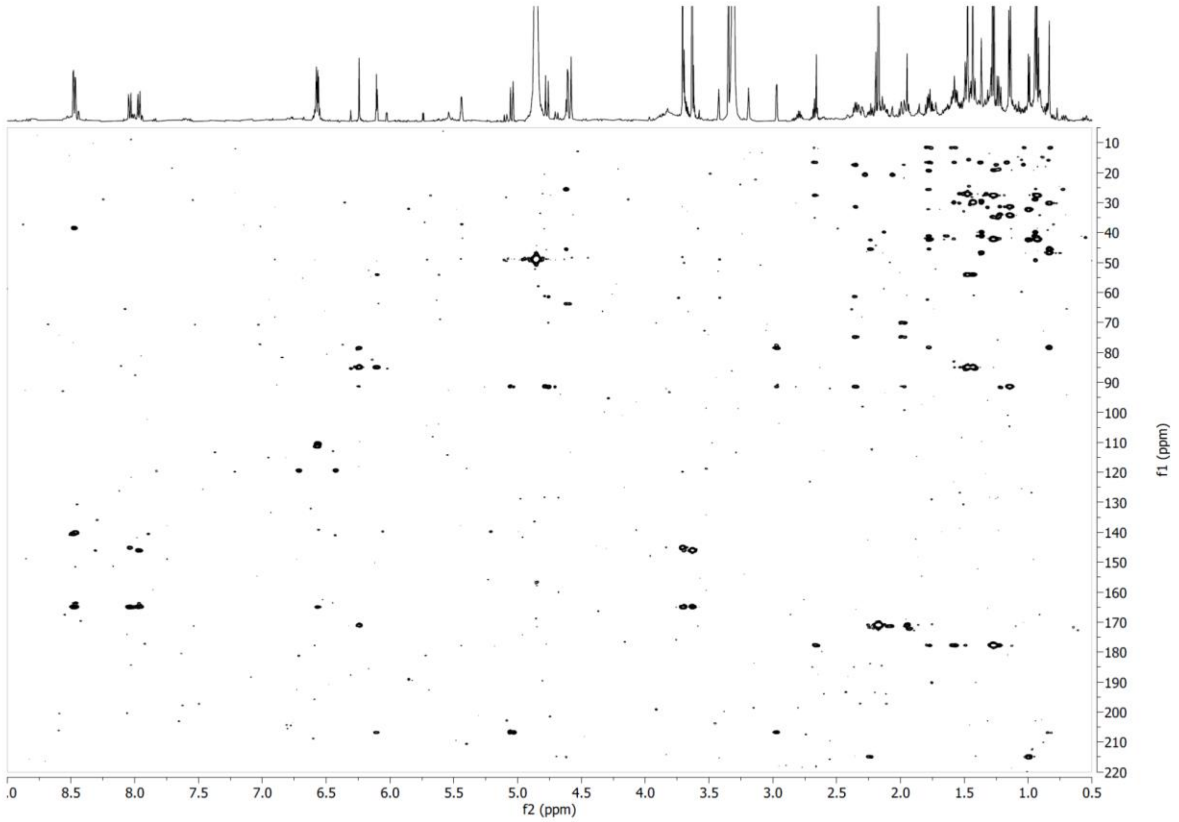


**Supplementary Figure S10**. HMBC NMR spectrum of compound **2** in $CD_3OD$.

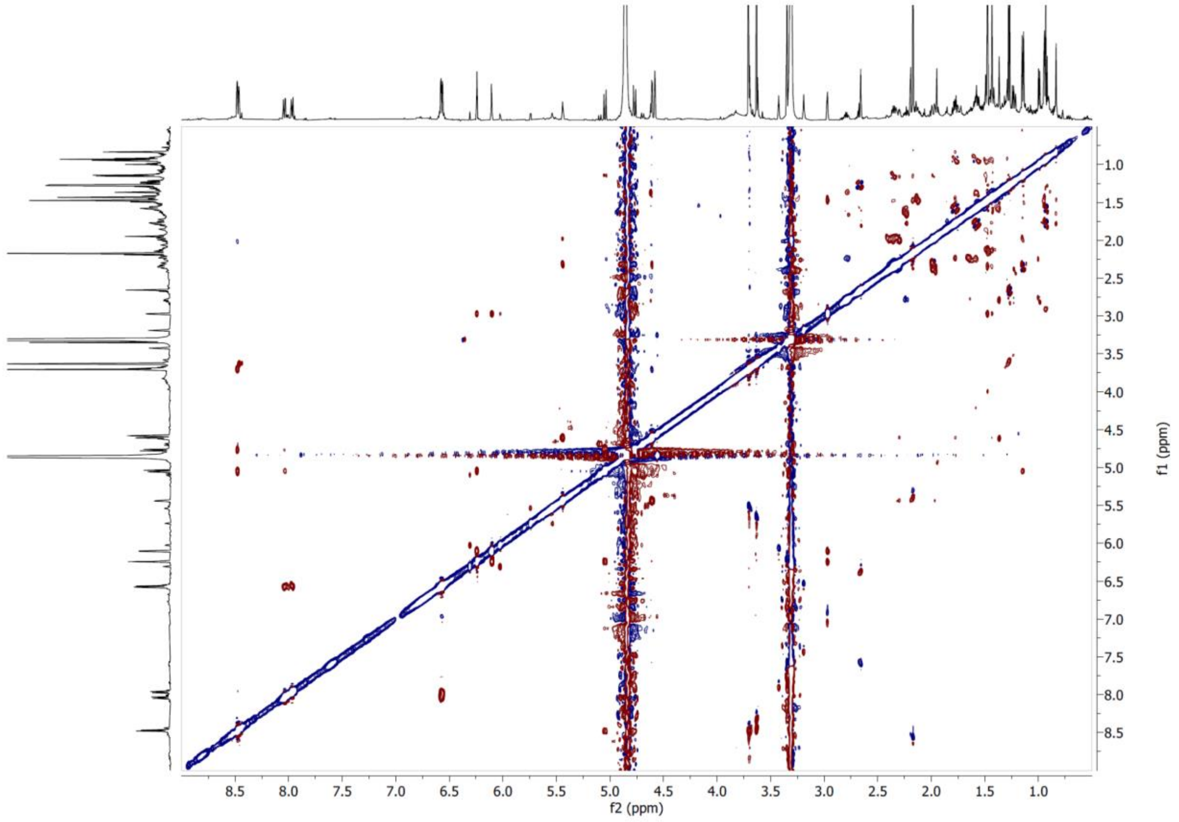


**Supplementary Figure S11**. ROESY NMR spectrum of compound **2** in $CD_3OD$ at 600 MHz.

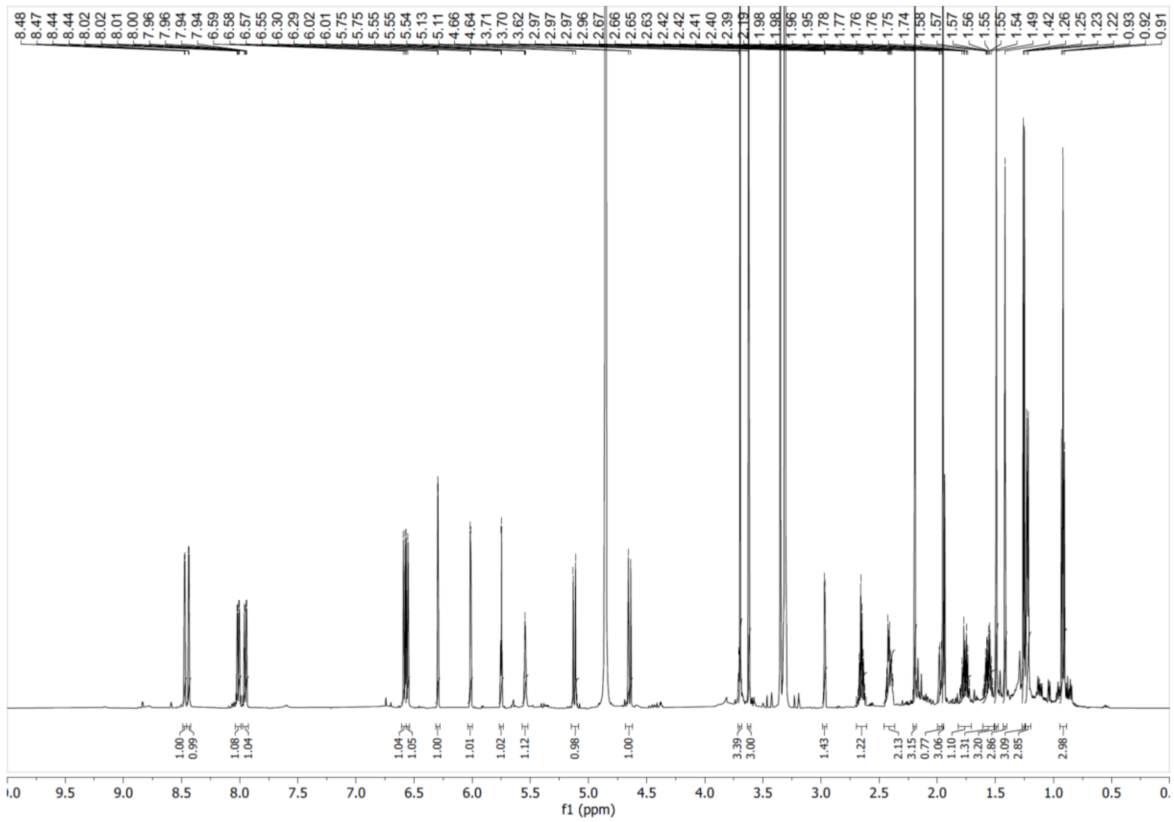

**Supplementary Figure S12**. $^{1}H$ NMR spectrum of compound **3** in $CD_3OD$ at 600 MHz.

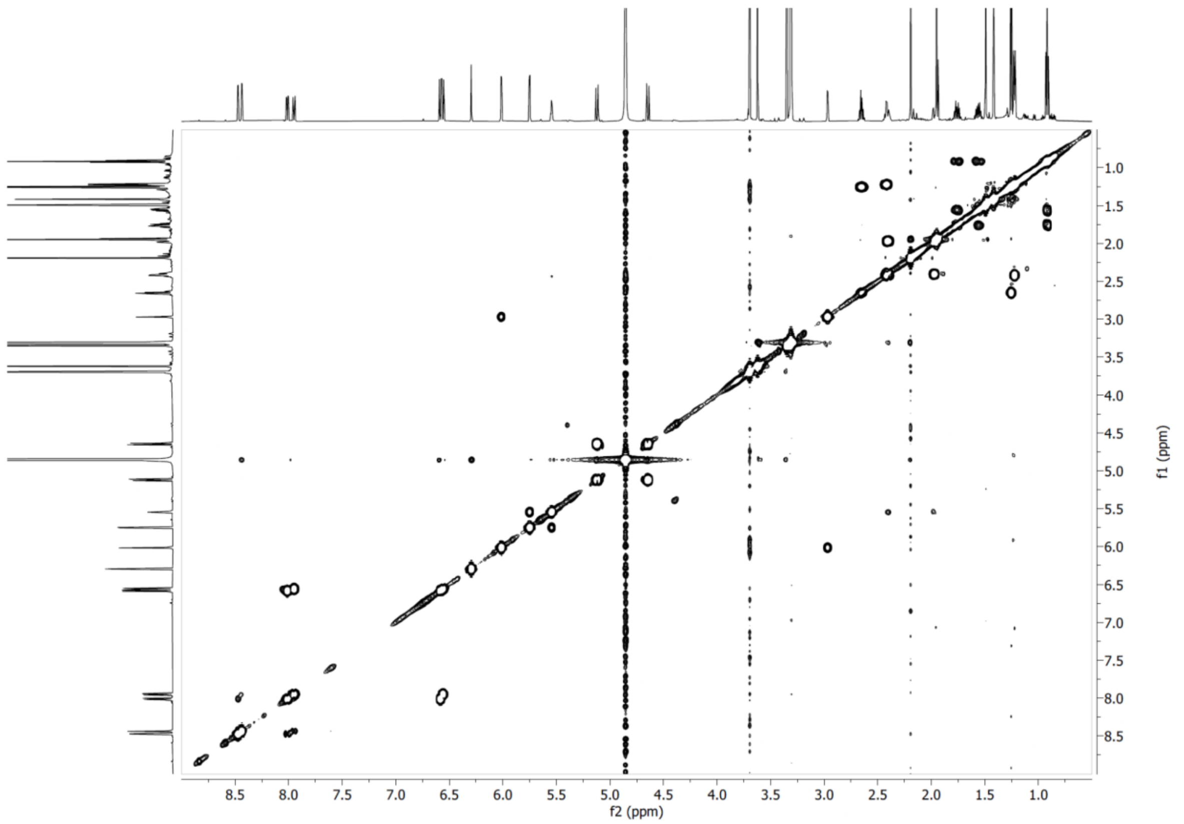

**Supplementary Figure S13**. COSY NMR spectrum of compound **3** in $CD_3OD$ at 600 MHz.

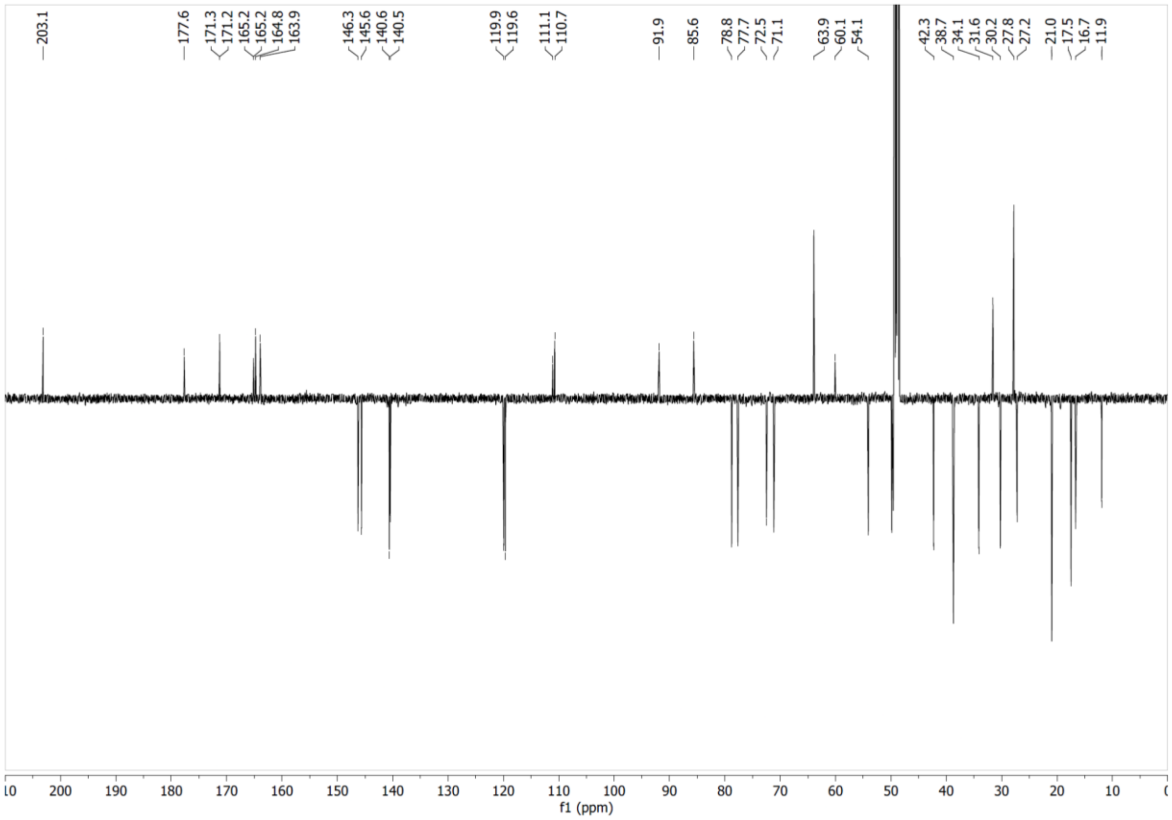


**Supplementary Figure S14**. $^{13}C$ NMR spectrum of compound **3** in $CD_3OD$ at 151 MHz.

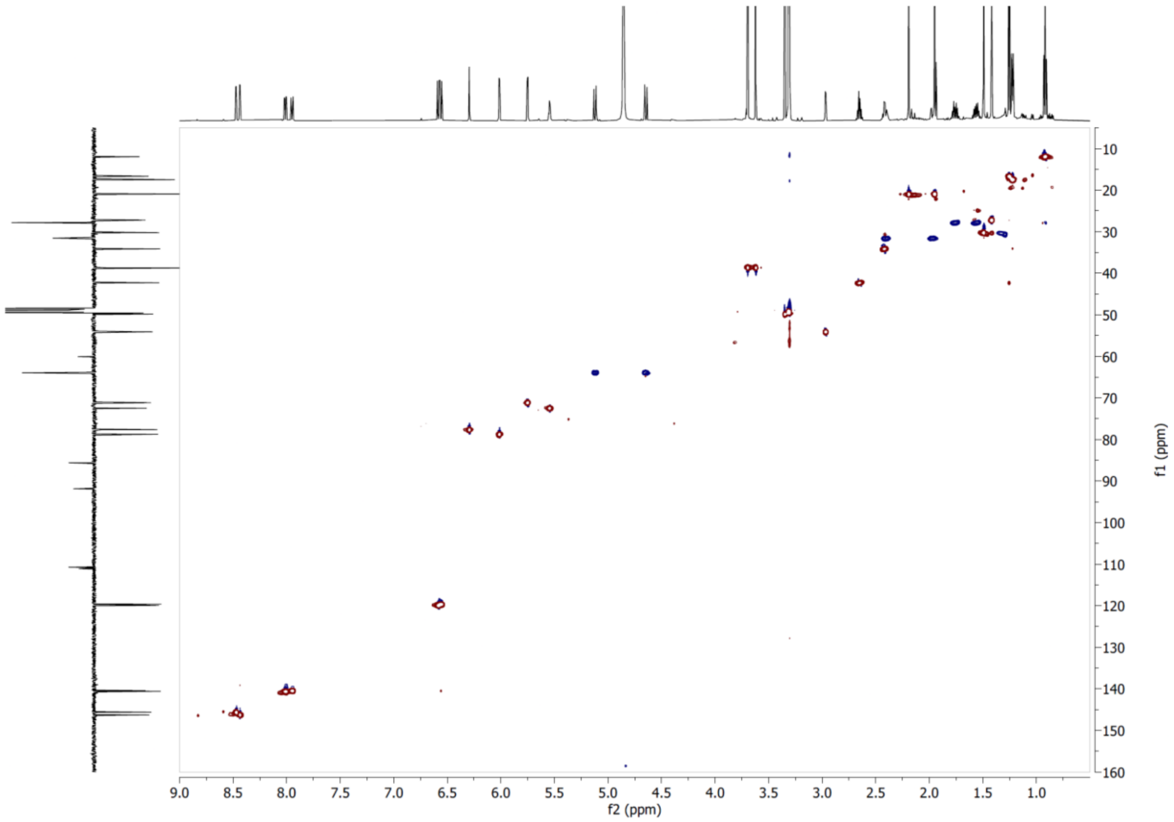


**Supplementary Figure S15**. HSQC NMR spectrum of compound **3** in $CD_3OD$.

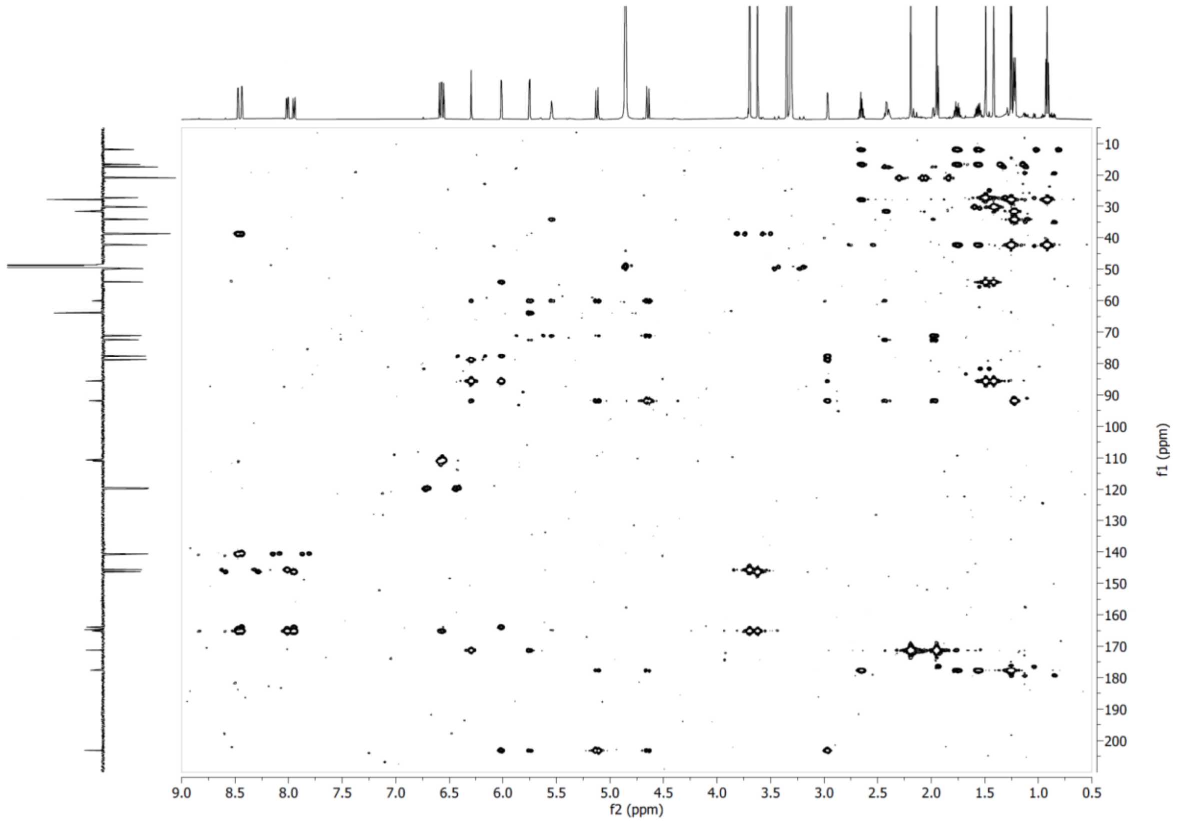


**Supplementary Figure S16**. HMBC NMR spectrum of compound **3** in $CD_3OD$.

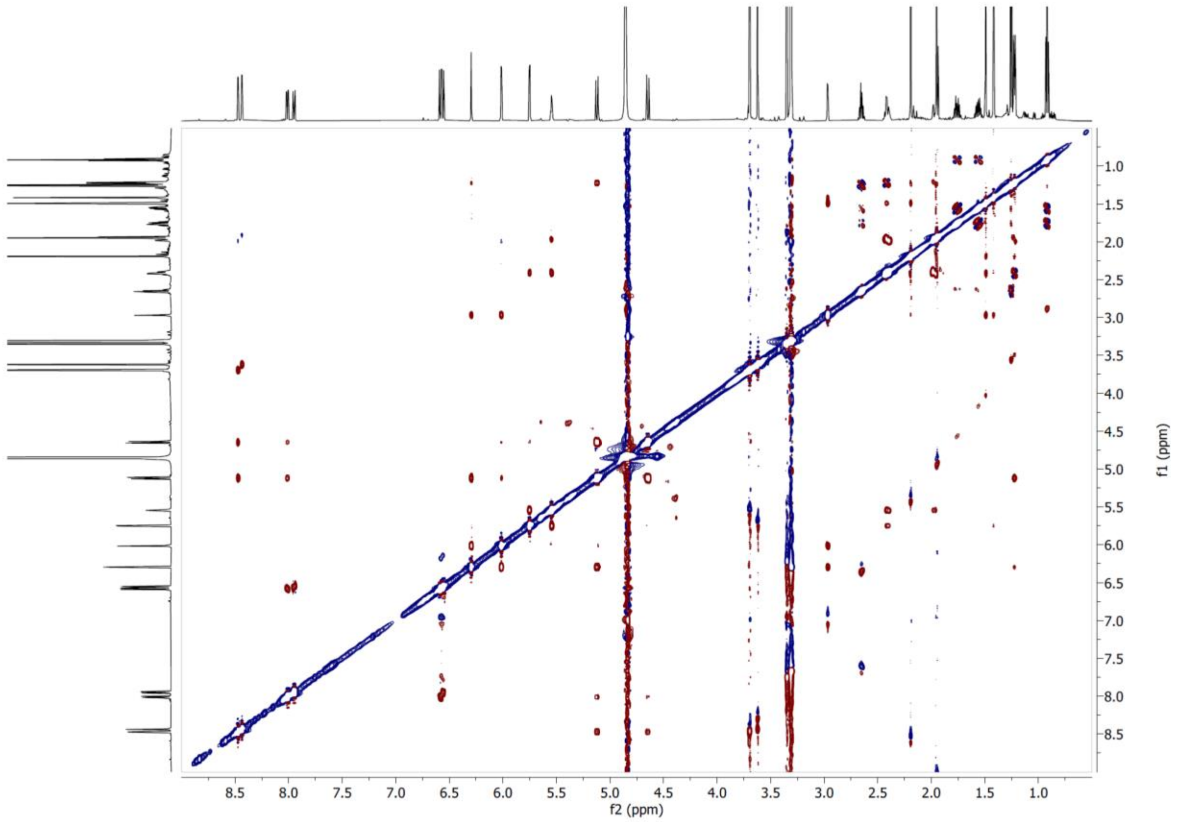


**Supplementary Figure S17**. ROESY NMR spectrum of compound **3** in $CD_3OD$ at 600 MHz.

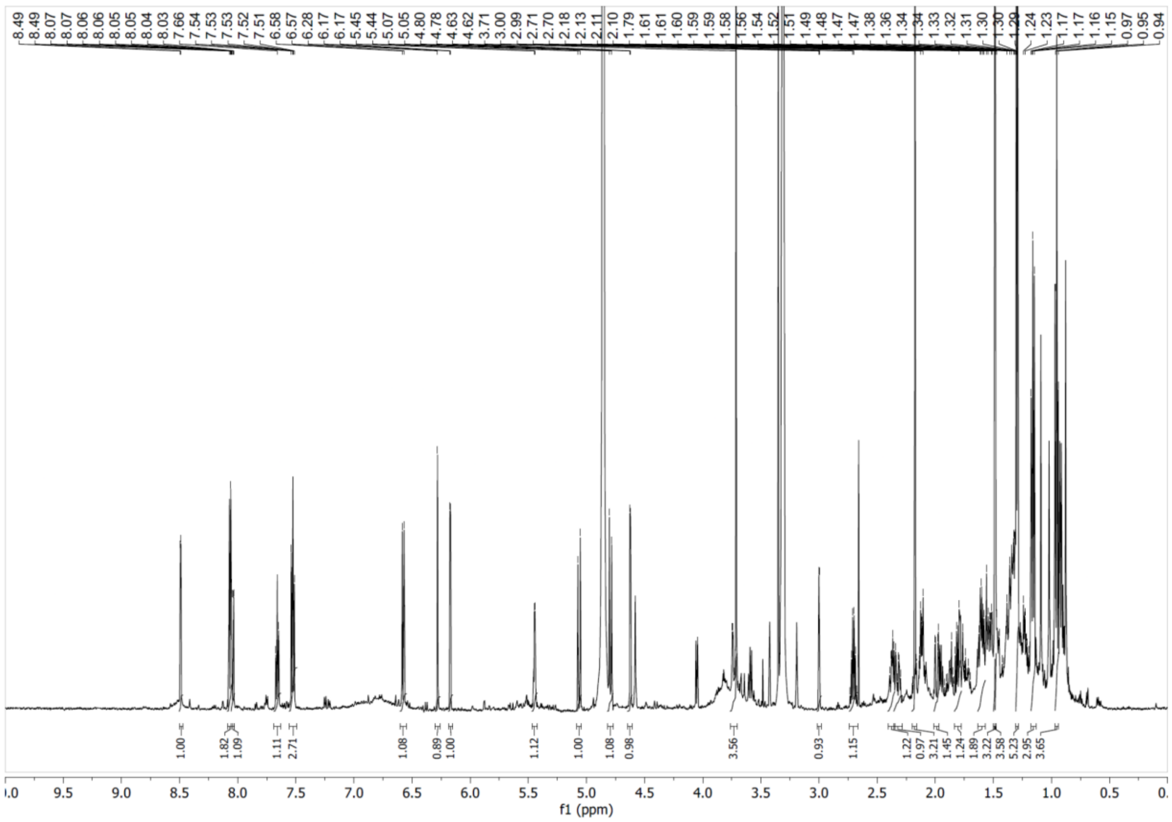

**Supplementary Figure S18**. $^1$H NMR spectrum of compound **4** in $CD_3OD$ at 600 MHz.

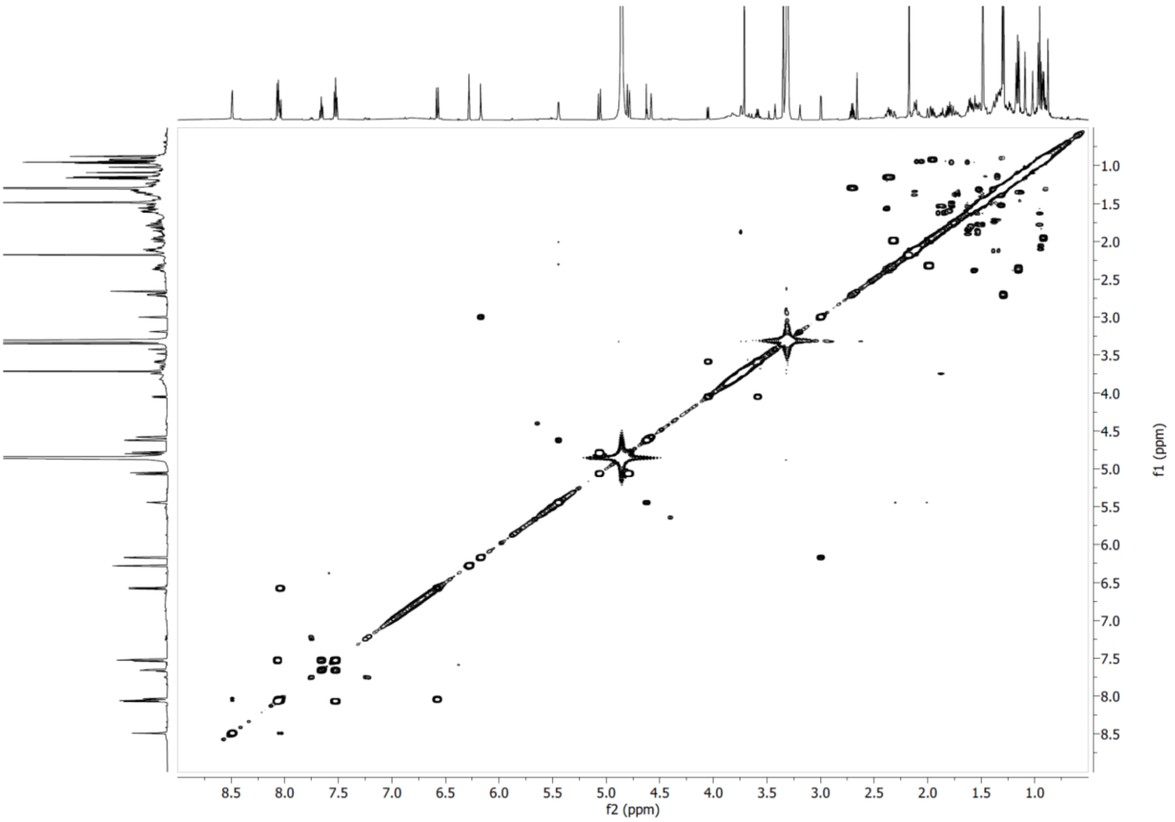


**Supplementary Figure S19**. COSY NMR spectrum of compound **4** in $CD_3OD$ at 600 MHz.

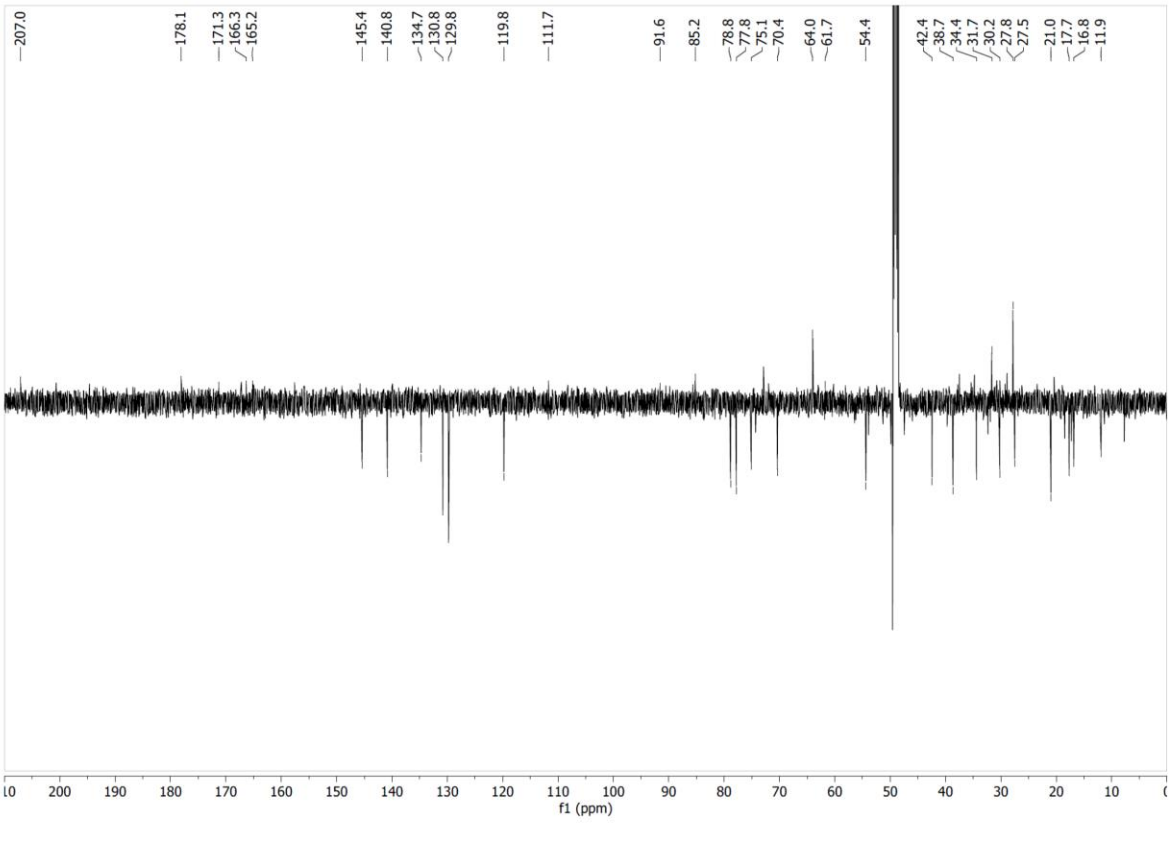


**Supplementary Figure S20**. $^{13}C$ NMR spectrum of compound **4** in $CD_3OD$ at 151MHz.

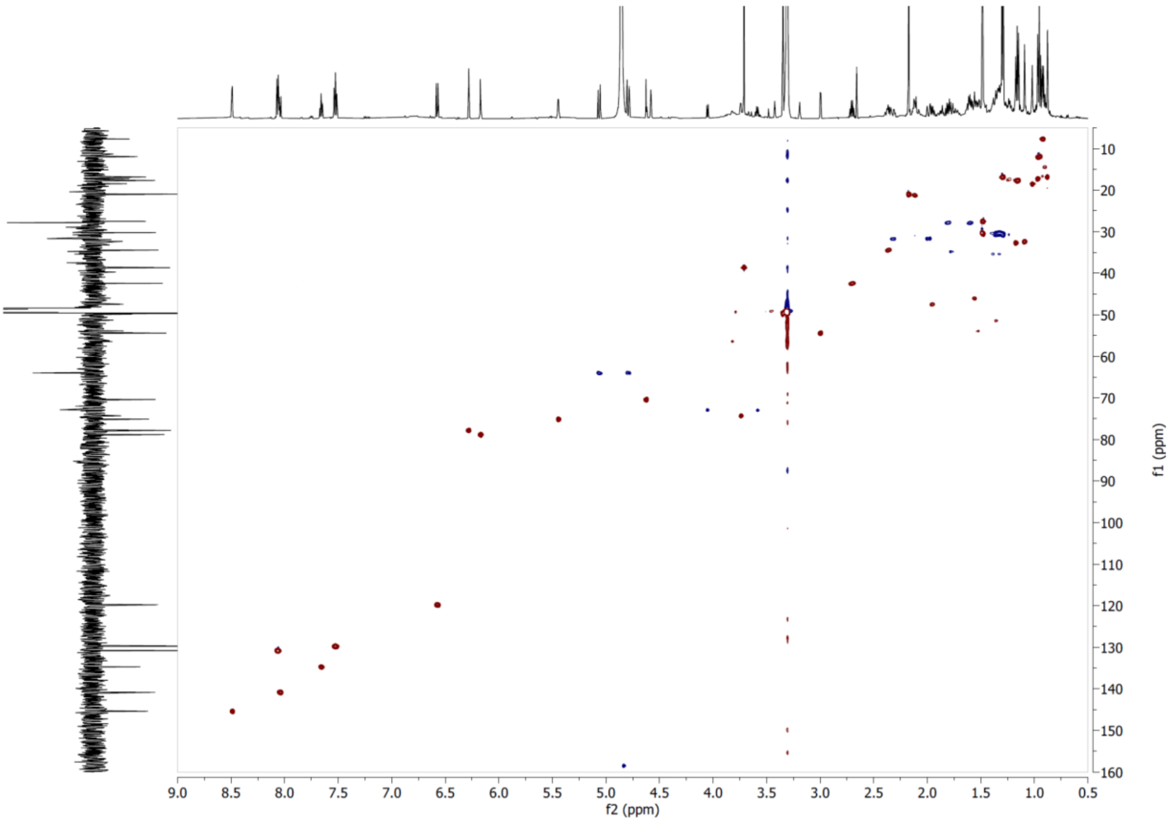


**Supplementary Figure S21**. HSQC NMR spectrum of compound **4** in $CD_3OD$.

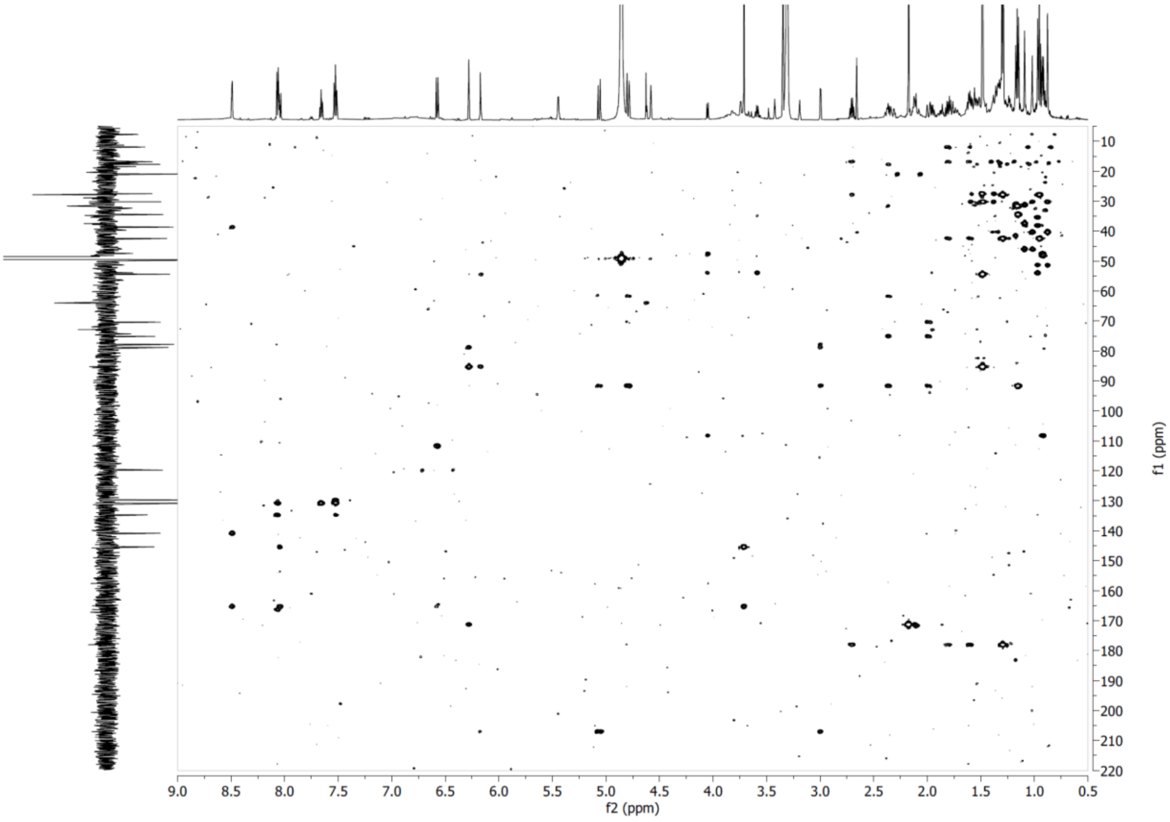


**Supplementary Figure S22**. HMBC NMR spectrum of compound **4** in $CD_3OD$.

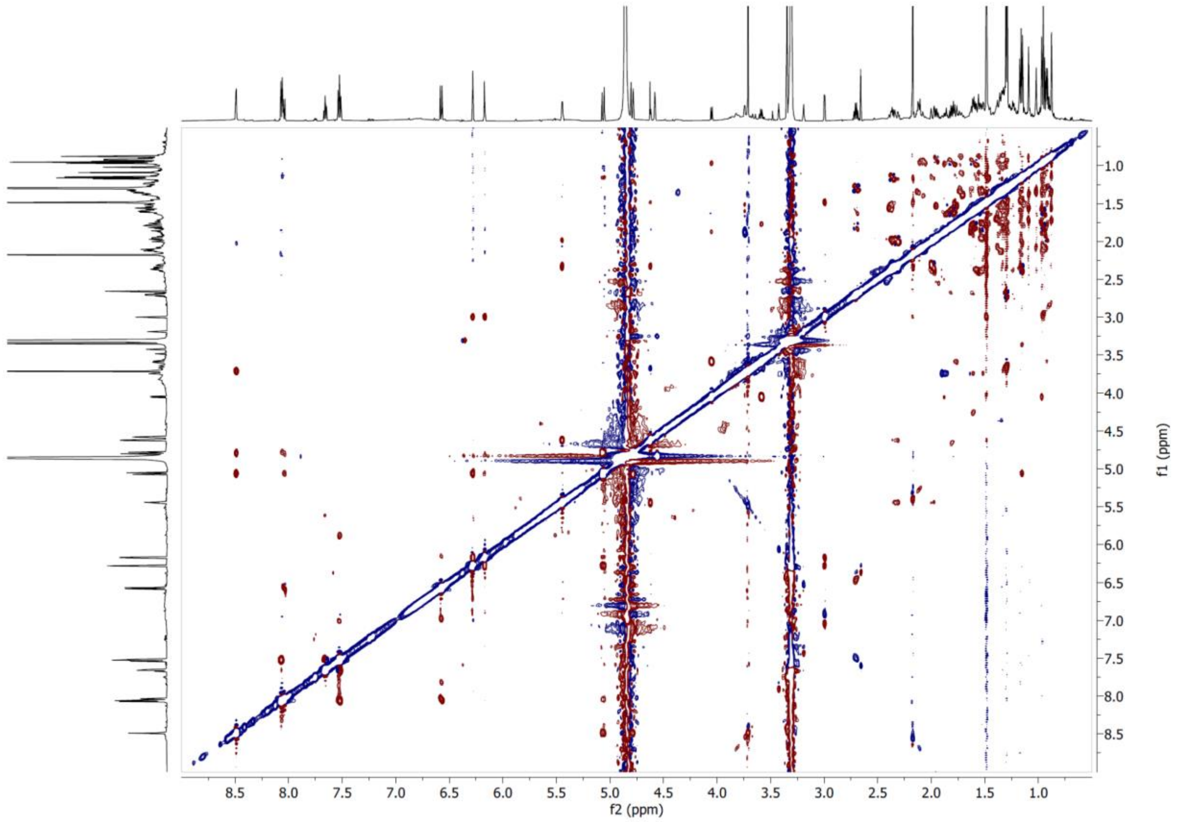


**Supplementary Figure S23**. ROESY NMR spectrum of compound **4** in $CD_3OD$ at 600 MHz.

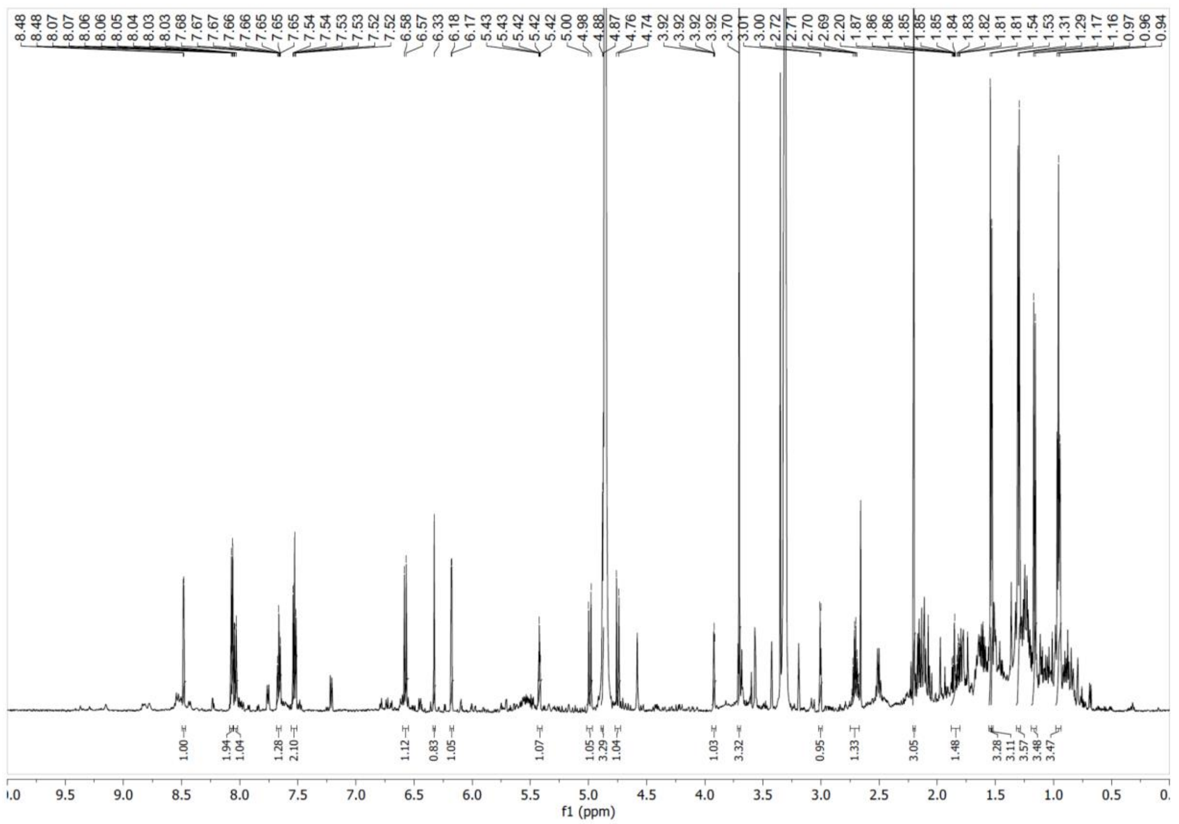


**Supplementary Figure S24**. $^1$H NMR spectrum of compound **5** in $CD_3OD$ at 600 MHz.

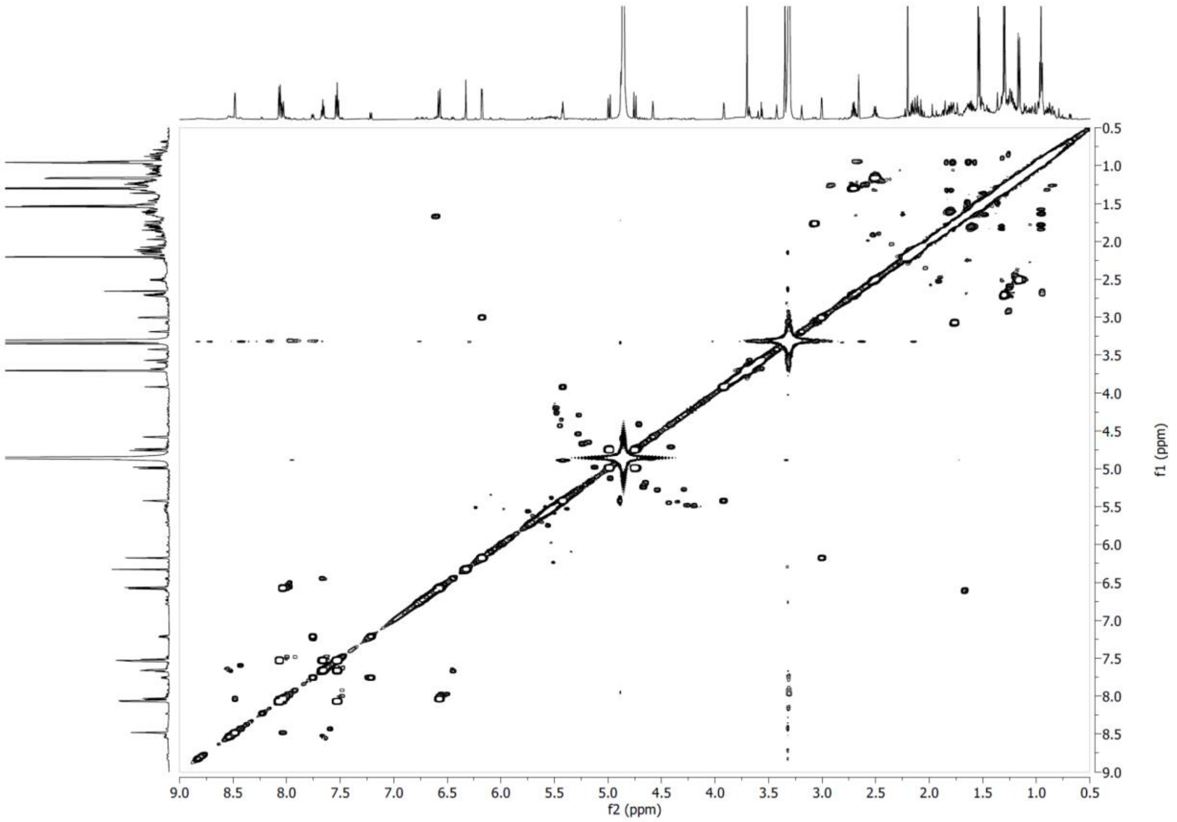


**Supplementary Figure S25**. COSY NMR spectrum of compound **5** in $CD_3OD$ at 600 MHz.

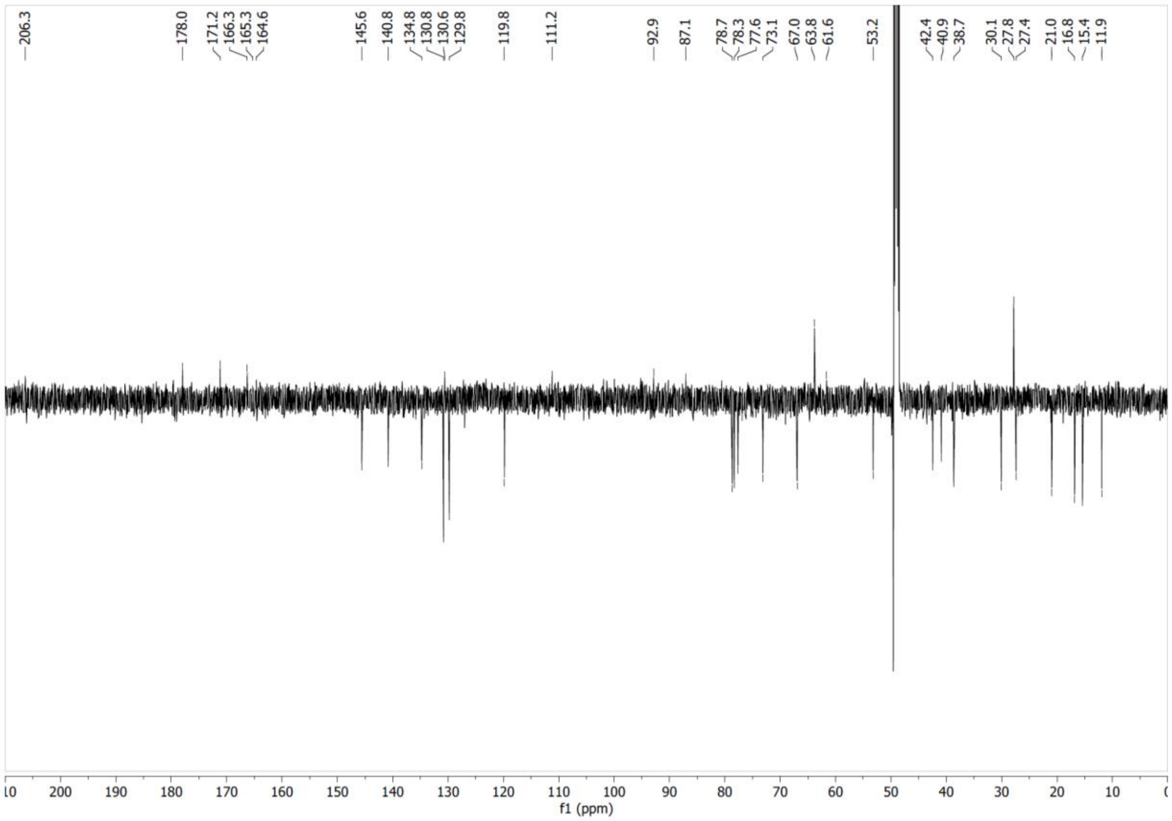


**Supplementary Figure S26**. $^{13}C$ NMR spectrum of compound **5** in $CD_3OD$ at 151 MHz.

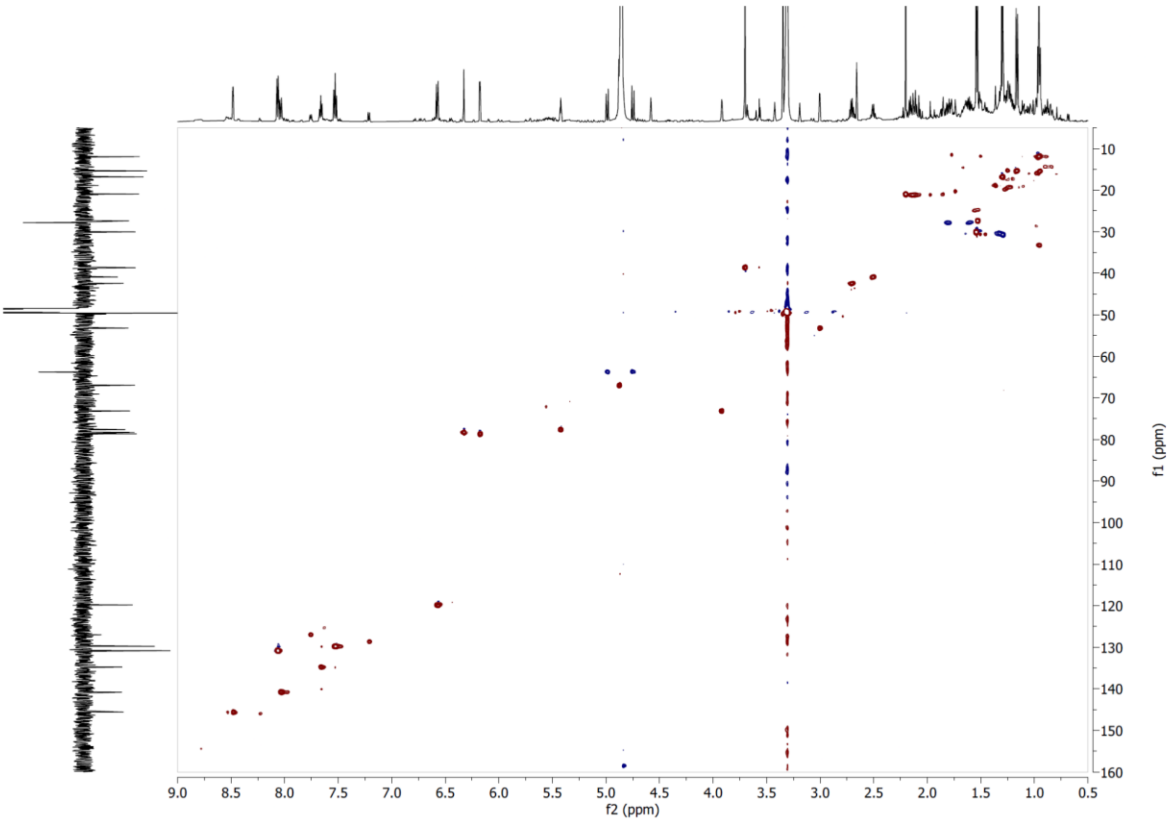


**Supplementary Figure S27**. HSQC NMR spectrum of compound **5** in $CD_3OD$.

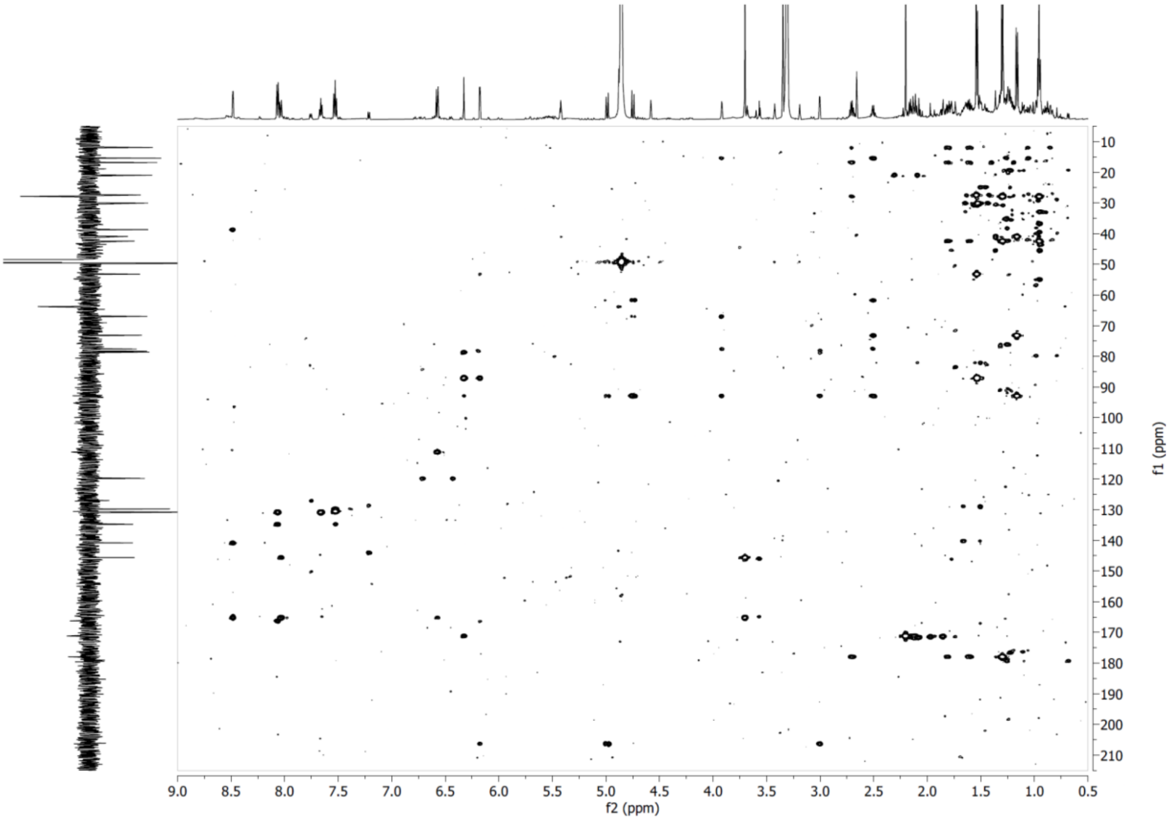


**Supplementary Figure S28**. HMBC NMR spectrum of compound **5** in $CD_3OD$.

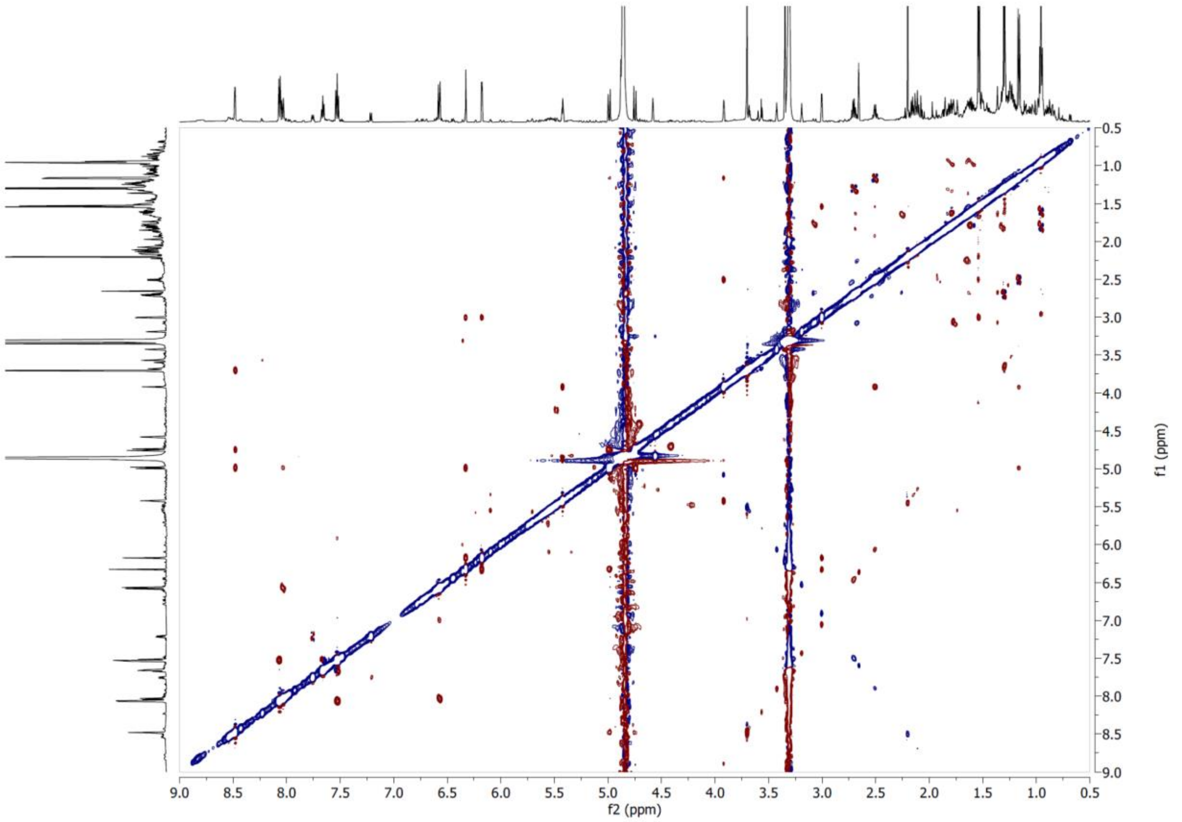

**Supplementary Figure S29**. ROESY NMR spectrum of compound **5** in $CD_3OD$ at 600 MHz.

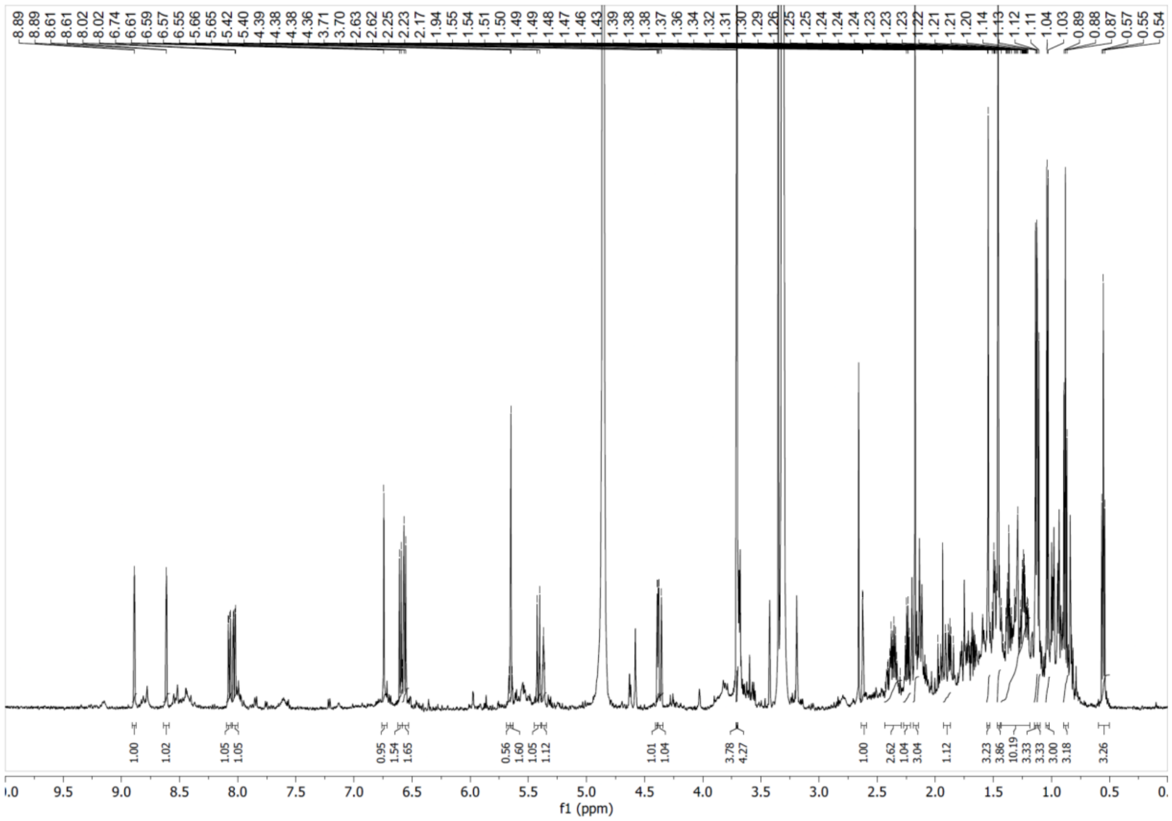

**Supplementary Figure S30**. $^1H$ NMR spectrum of compound **6** in $CD_3OD$ at 600 MHz.

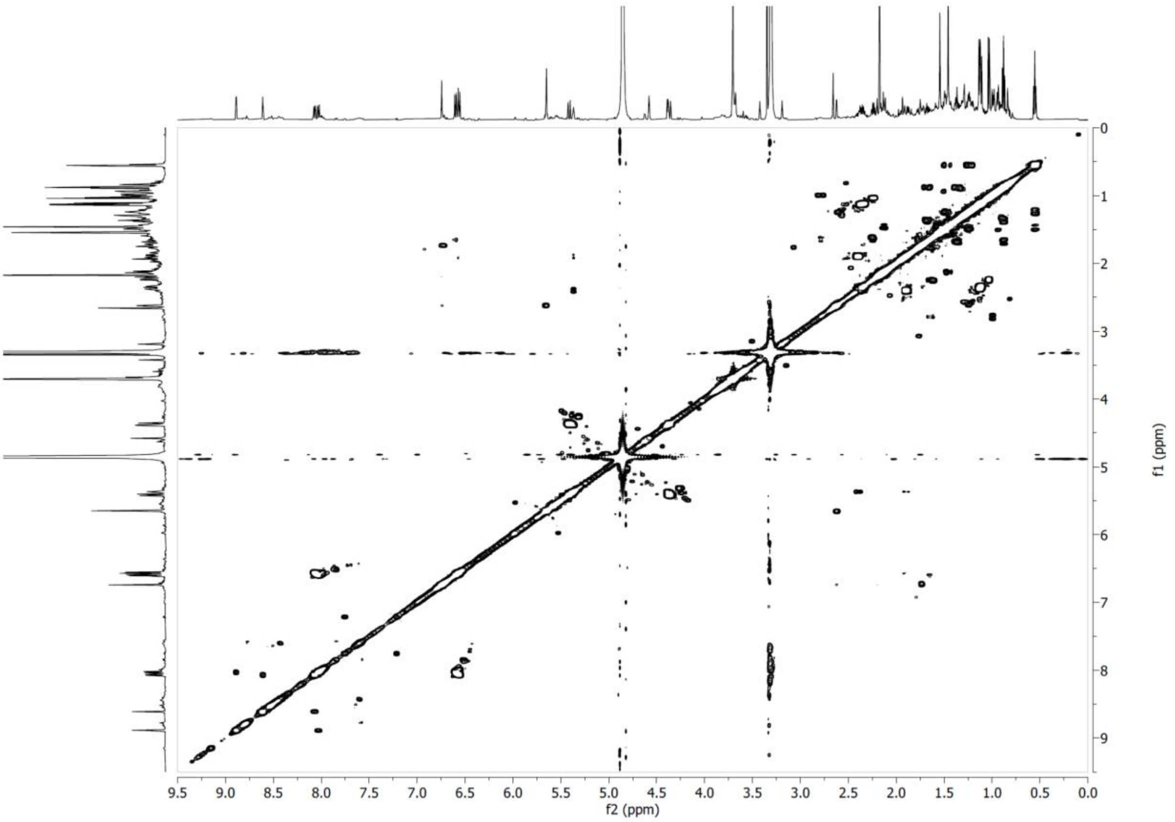


**Supplementary Figure S31**. COSY NMR spectrum of compound **6** in $CD_3OD$ at 600 MHz.

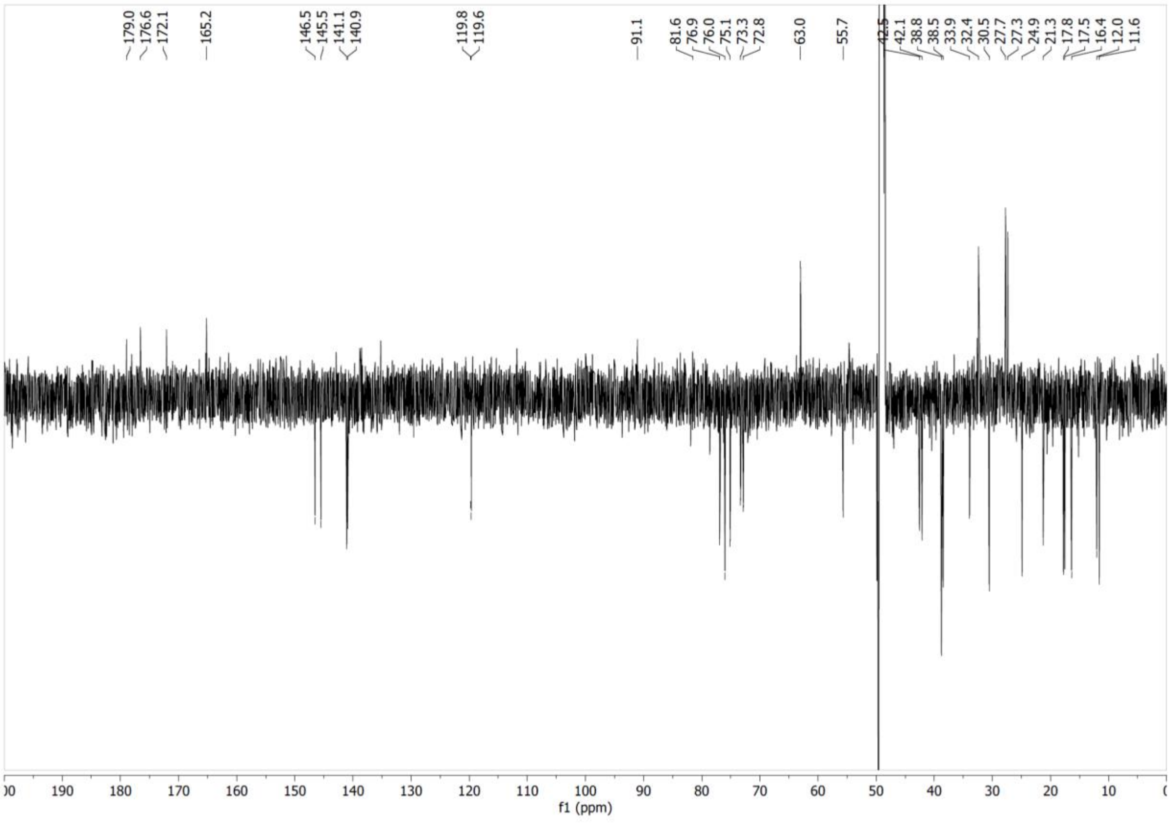


**Supplementary Figure S32**. $^{13}C$ NMR spectrum of compound **6** in $CD_3OD$ at 151 MHz.

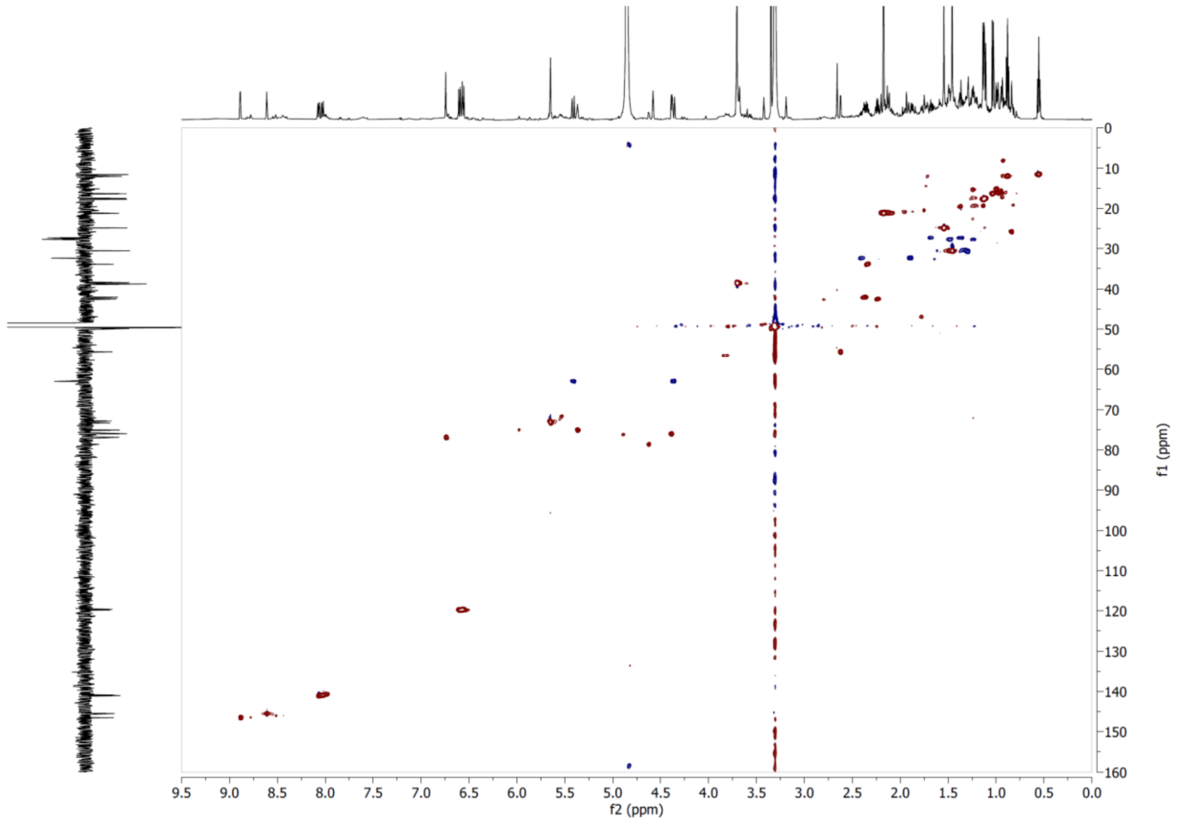


**Supplementary Figure S33**. HSQC NMR spectrum of compound **6** in $CD_3OD$.

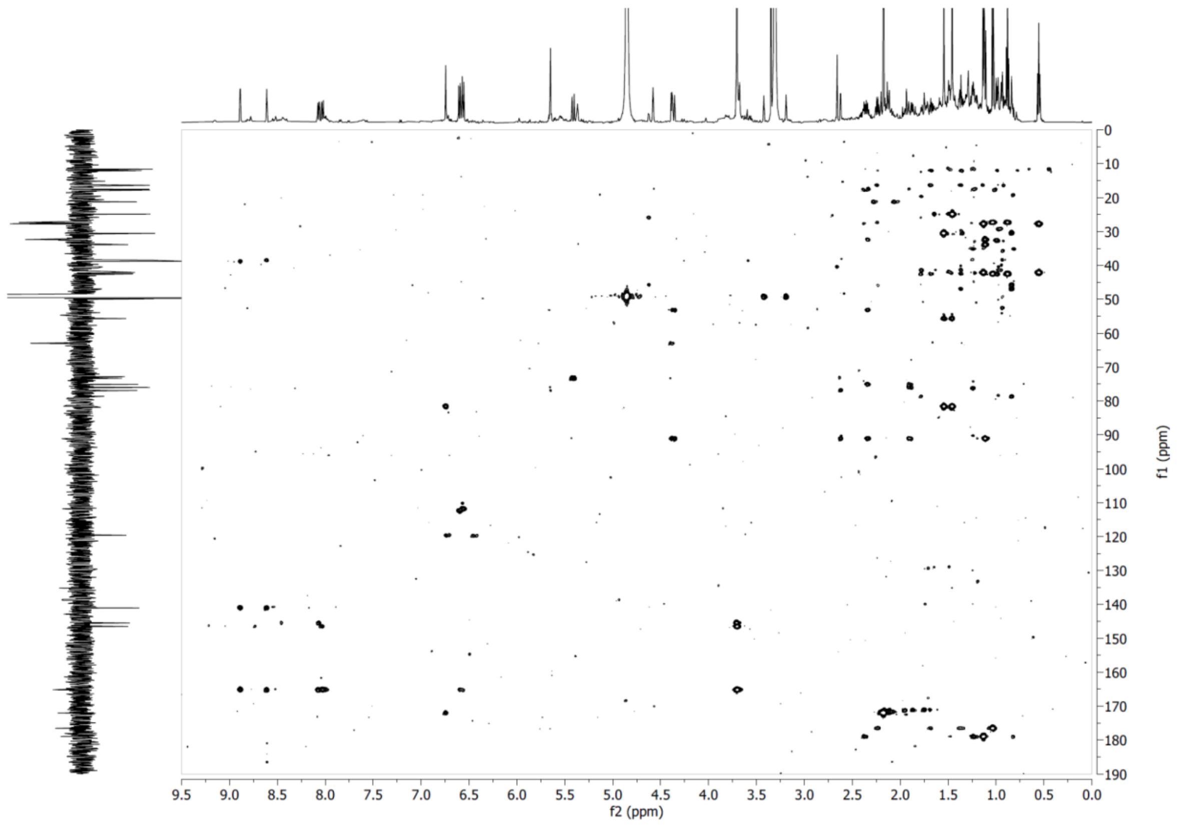


**Supplementary Figure S34**. HMBC NMR spectrum of compound **6** in $CD_3OD$.

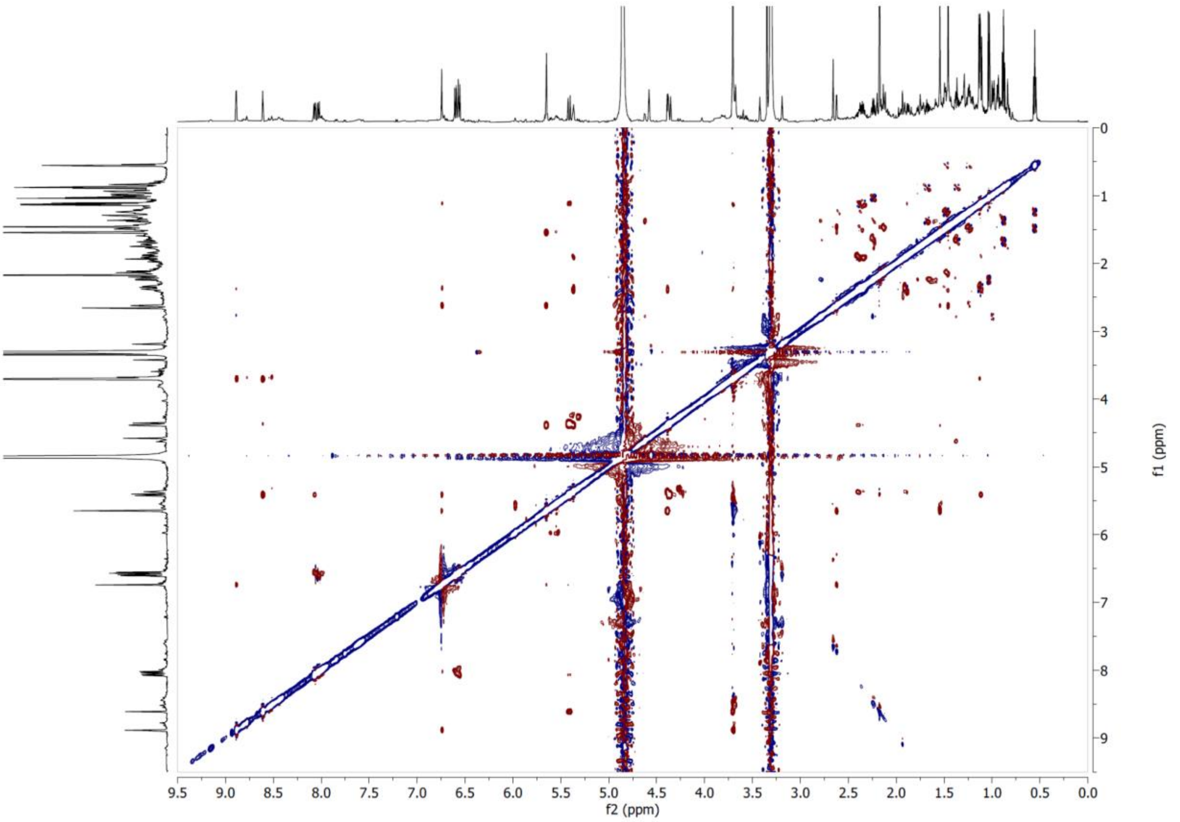


**Supplementary Figure S35**. ROESY NMR spectrum of compound **6** in $CD_3OD$ at 600 MHz.

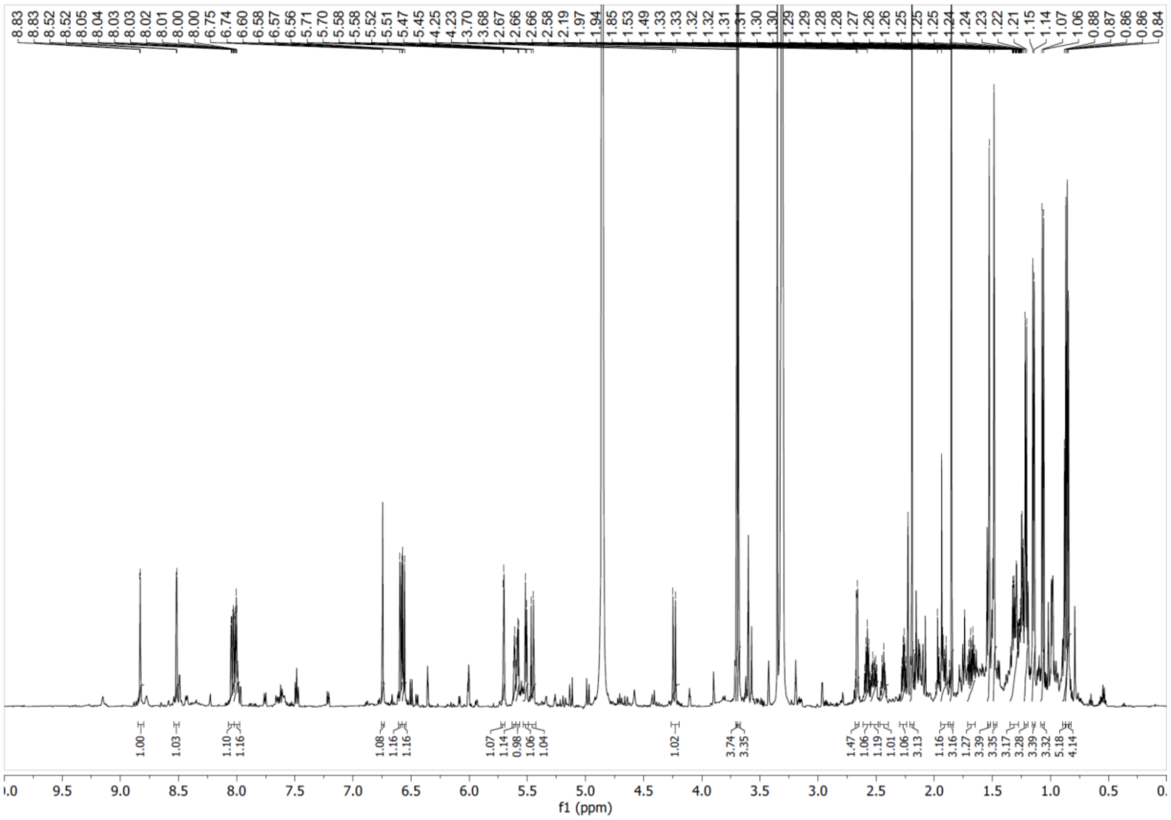


**Supplementary Figure S36**. $^1H$ NMR spectrum of compound **7** in $CD_3OD$ at 600 MHz.

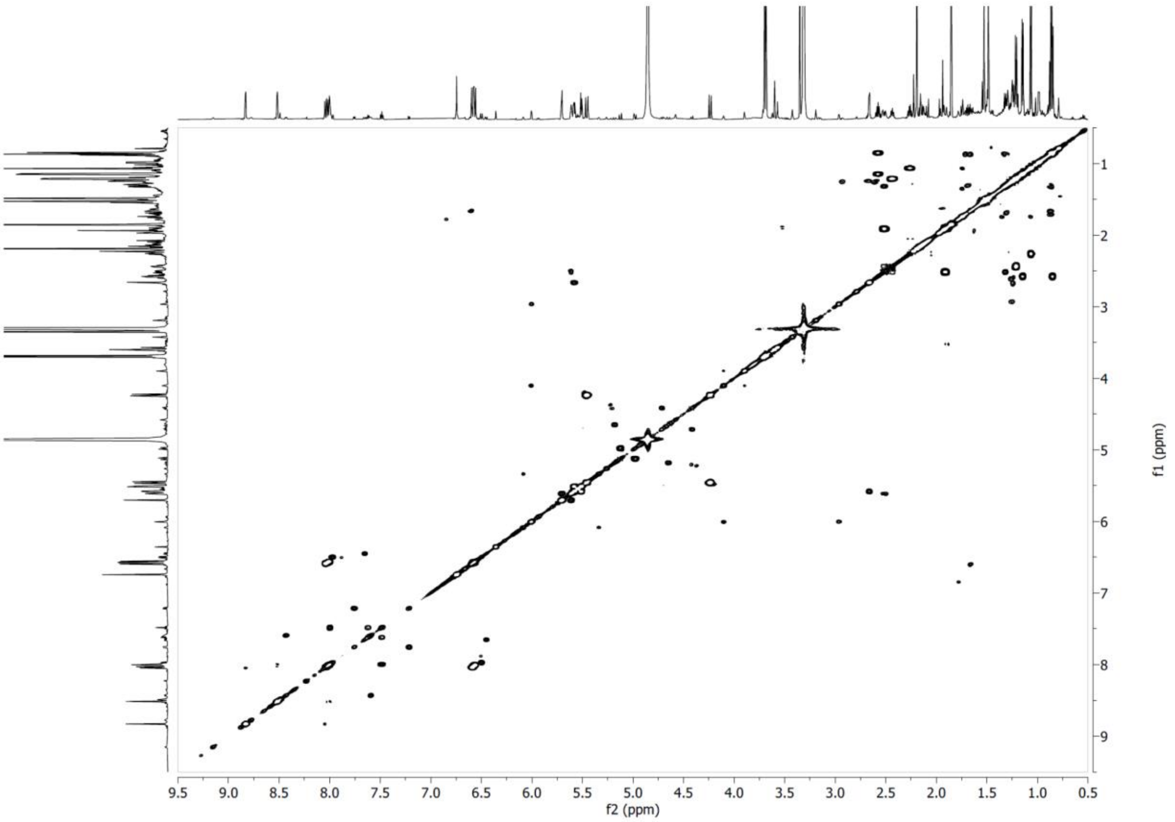


**Supplementary Figure S37**. COSY NMR spectrum of compound **7** in $CD_3OD$ at 600 MHz.

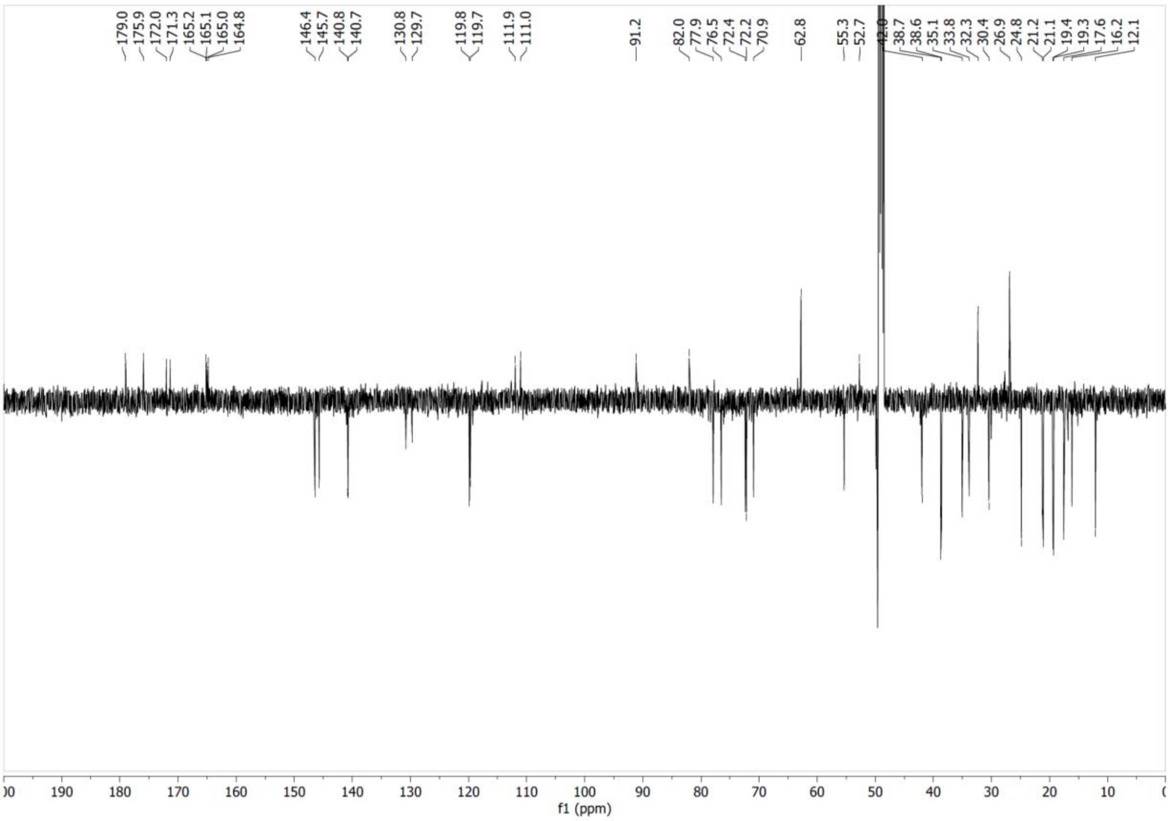


**Supplementary Figure S38**. $^{13}C$ NMR spectrum of compound **7** in $CD_3OD$ at 151 MHz.

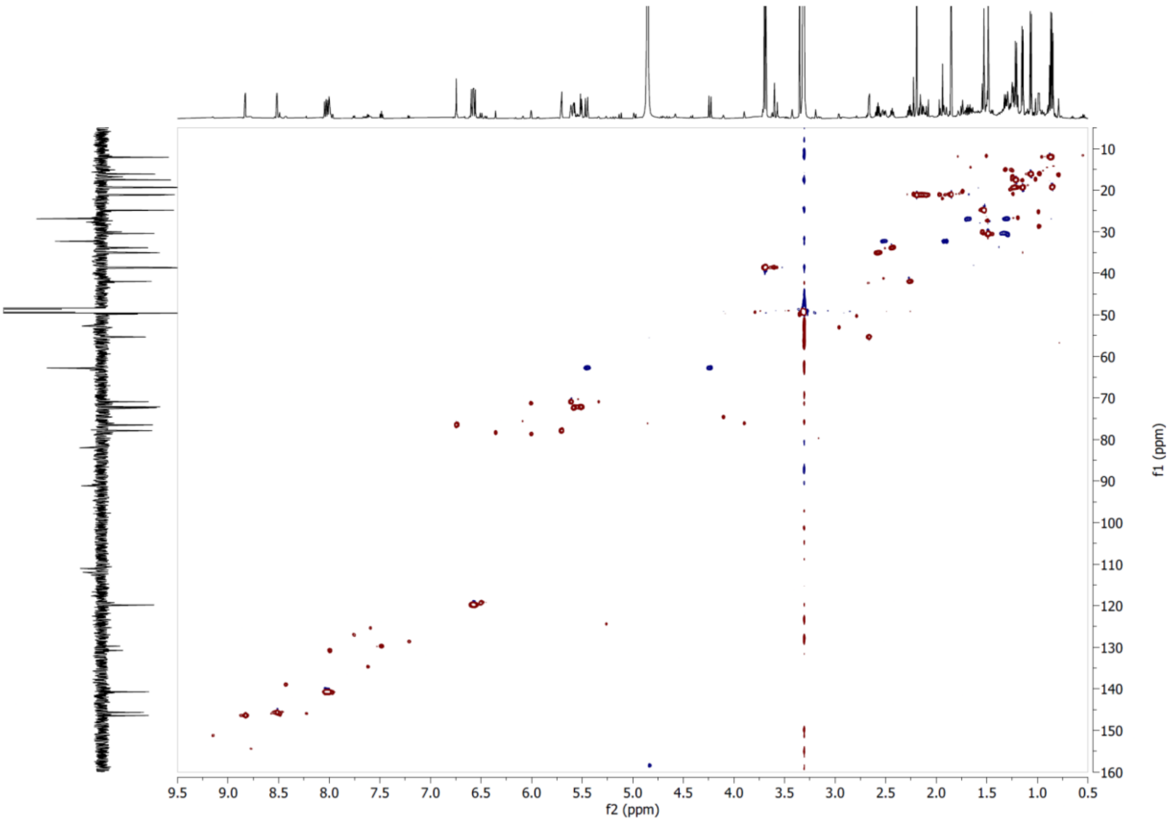


**Supplementary Figure S39**. HSQC NMR spectrum of compound **7** in $CD_3OD$.

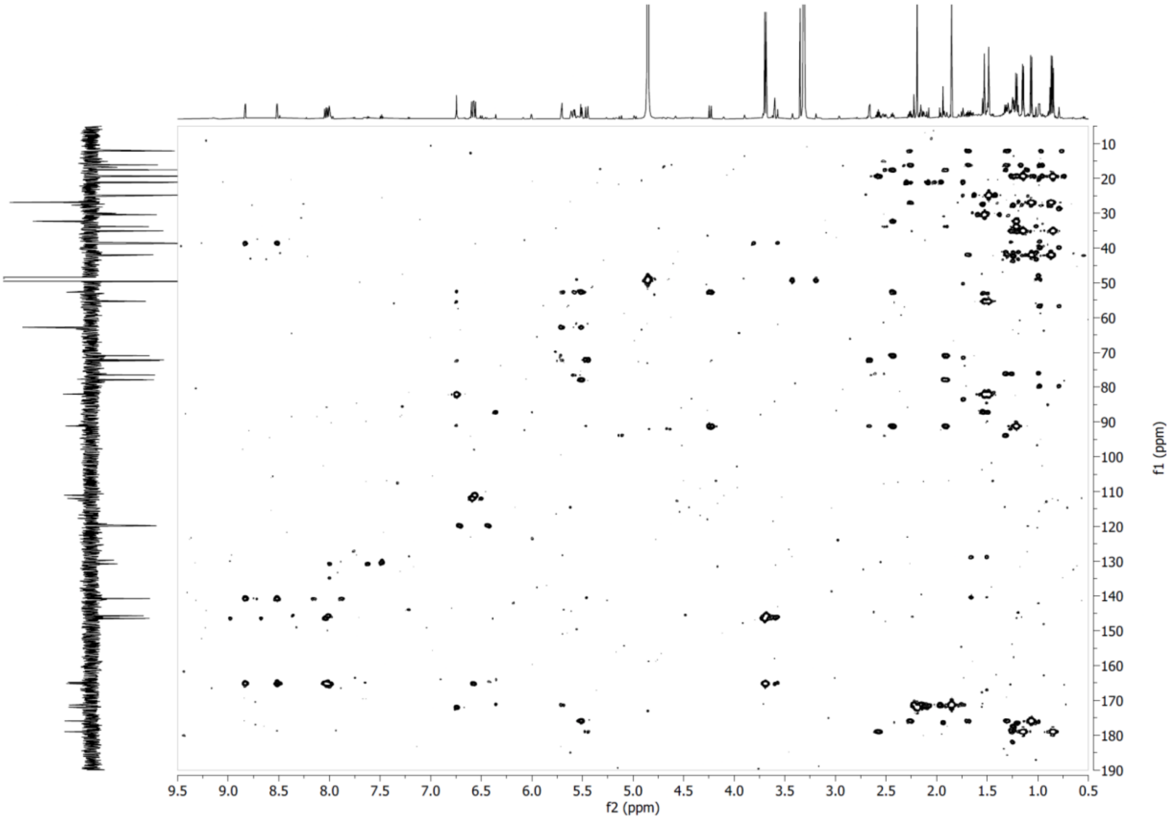


**Supplementary Figure S40**. HMBC NMR spectrum of compound **7** in $CD_3OD$.

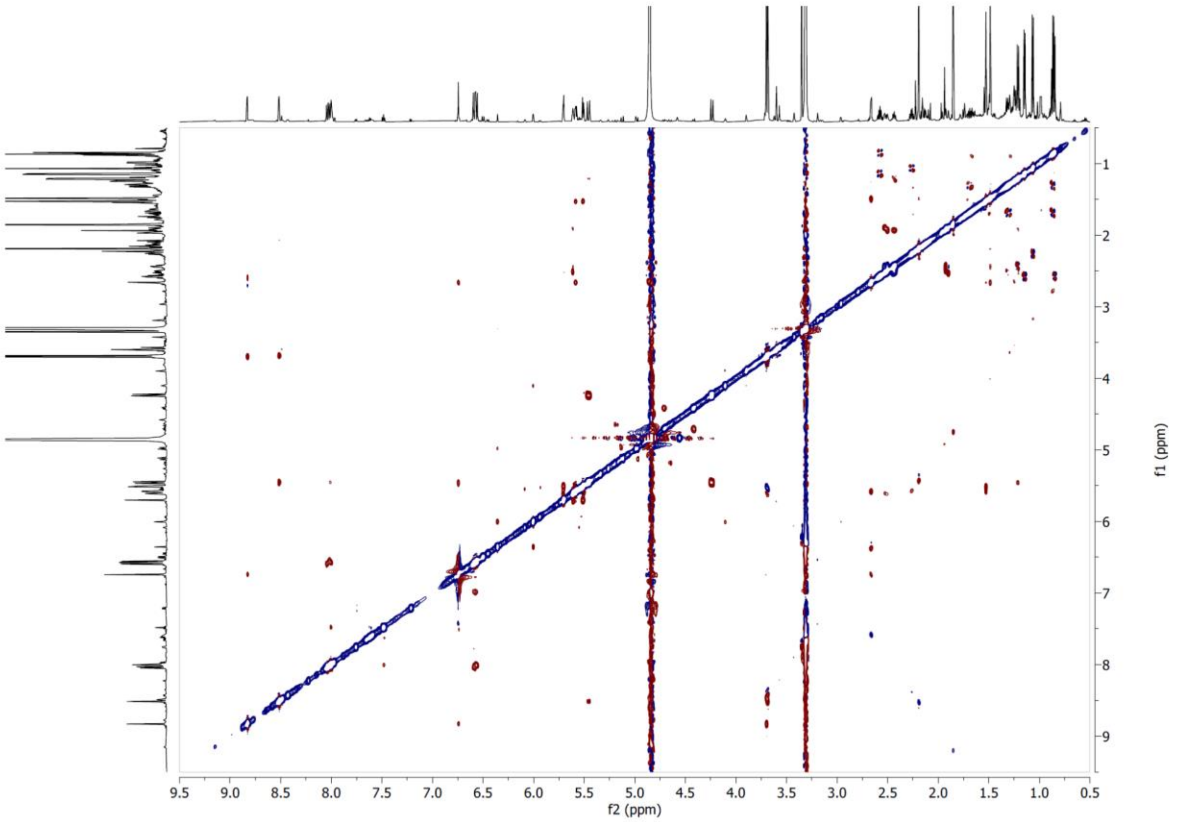


**Supplementary Figure S41**. ROESY NMR spectrum of compound **7** in $CD_3OD$ at 600 MHz.

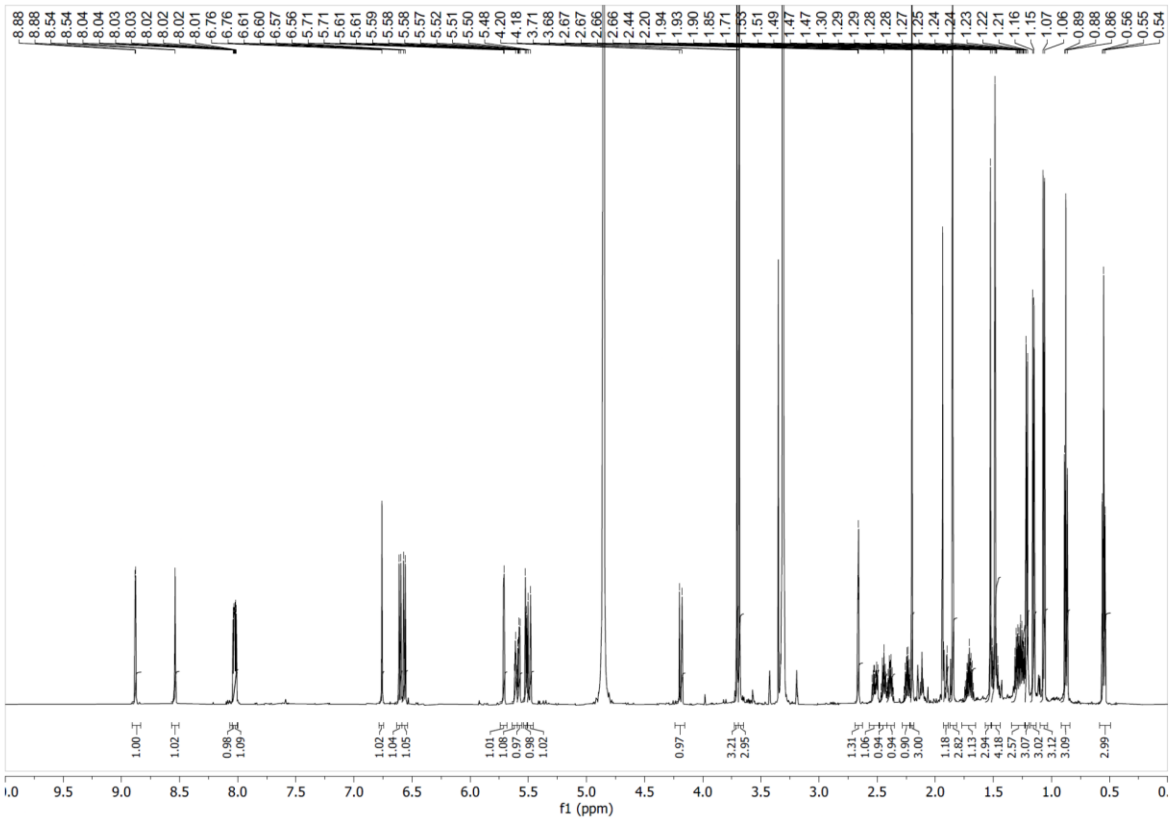


**Supplementary Figure S42**. $^1H$ NMR spectrum of compound **8** in $CD_3OD$ at 600 MHz.

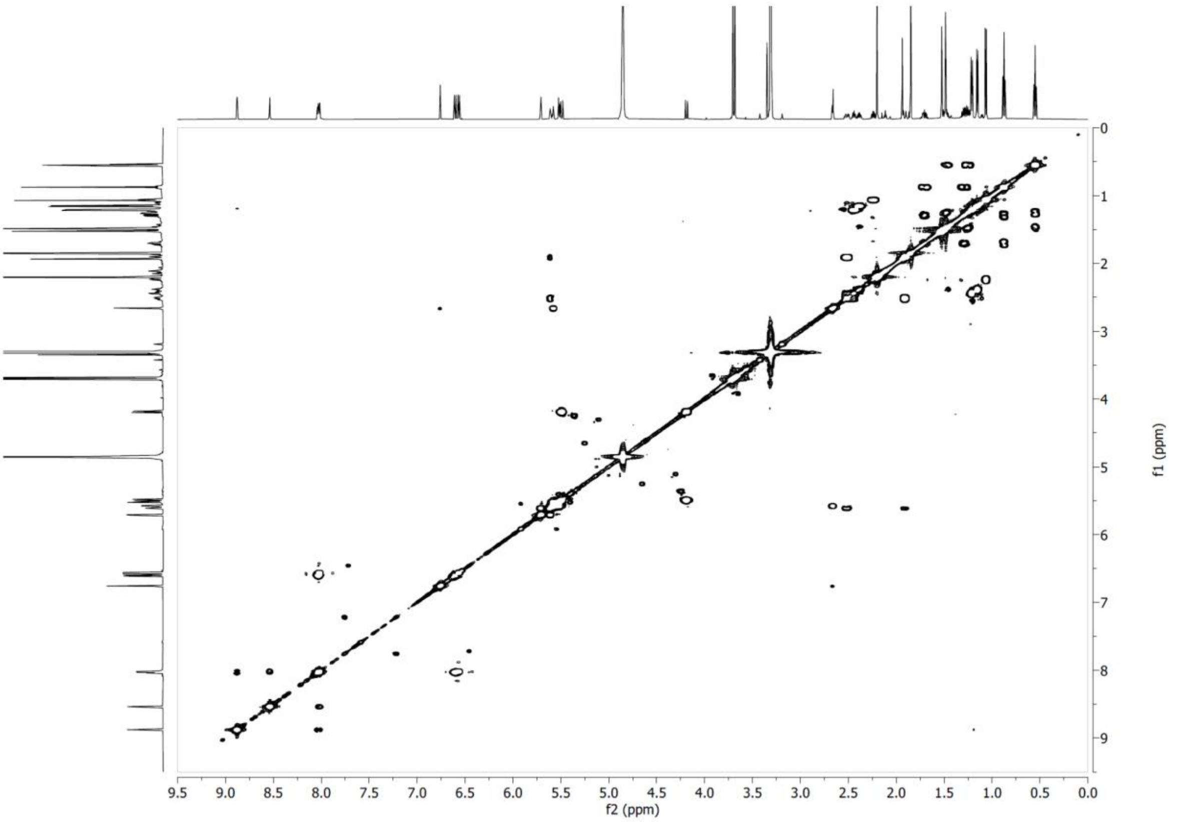


**Supplementary Figure S43**. COSY NMR spectrum of compound **8** in $CD_3OD$ at 600 MHz.

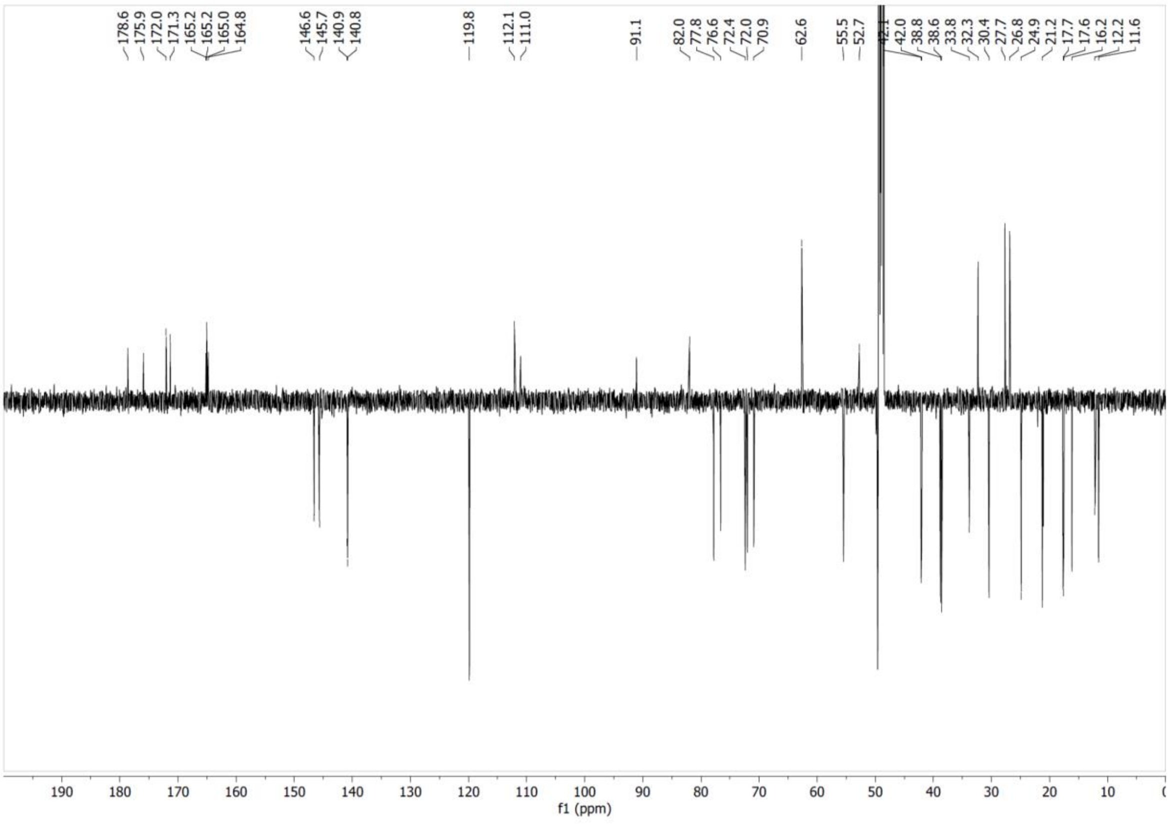


**Supplementary Figure S44**. $^{13}C$ NMR spectrum of compound **8** in $CD_3OD$ at 151 MHz.

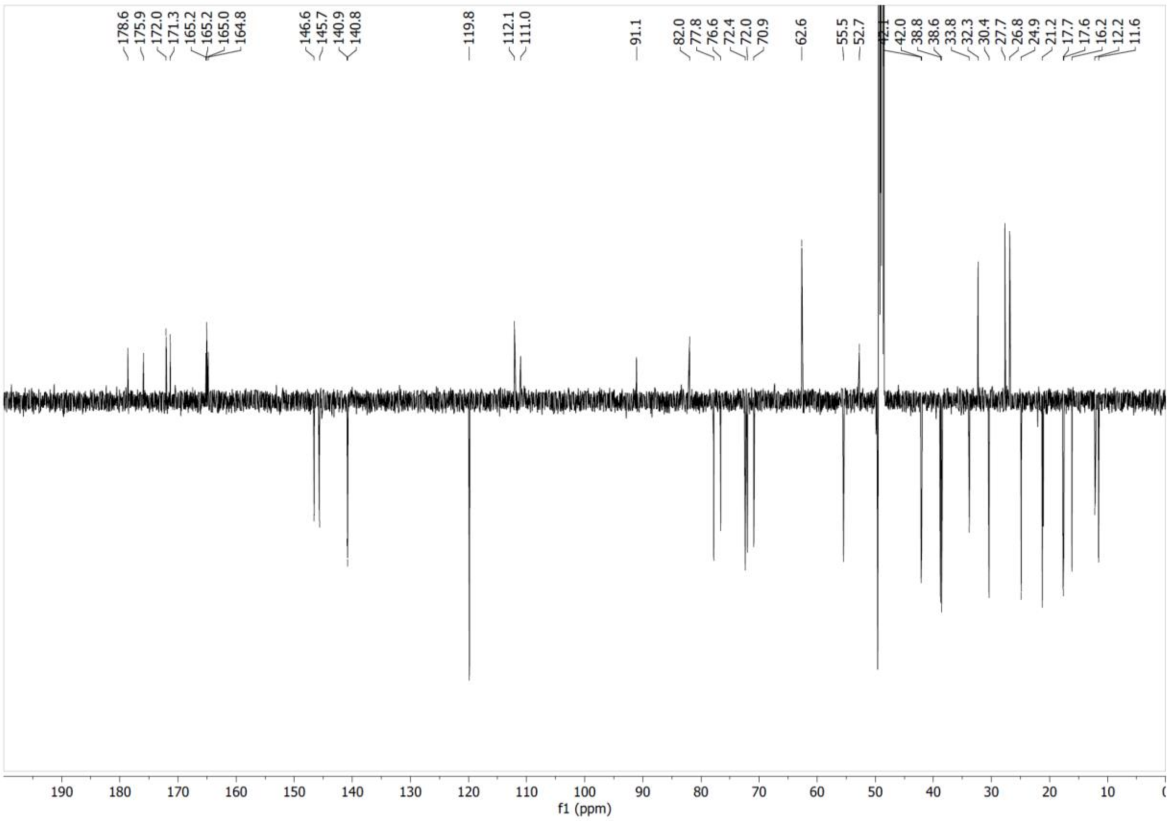


**Supplementary Figure S45**. HSQC NMR spectrum of compound **8** in $CD_3OD$.

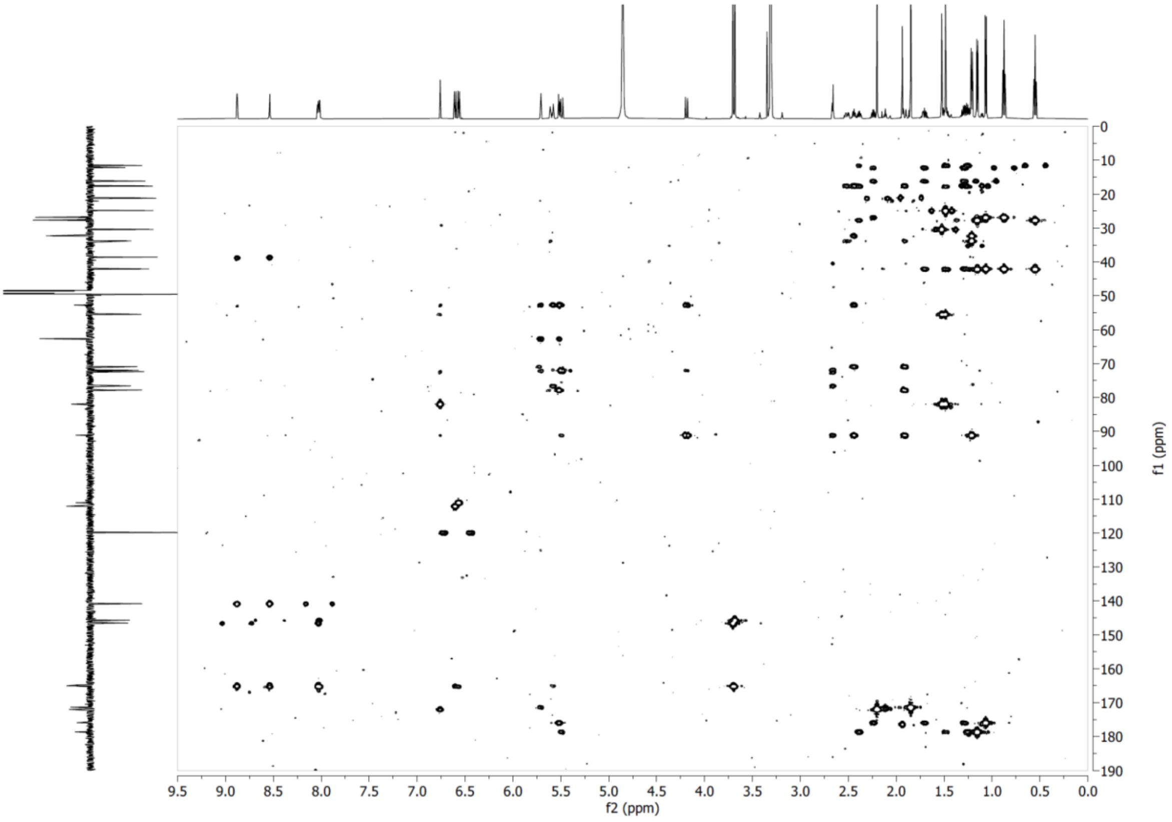


**Supplementary Figure S46**. HMBC NMR spectrum of compound **8** in $CD_3OD$.

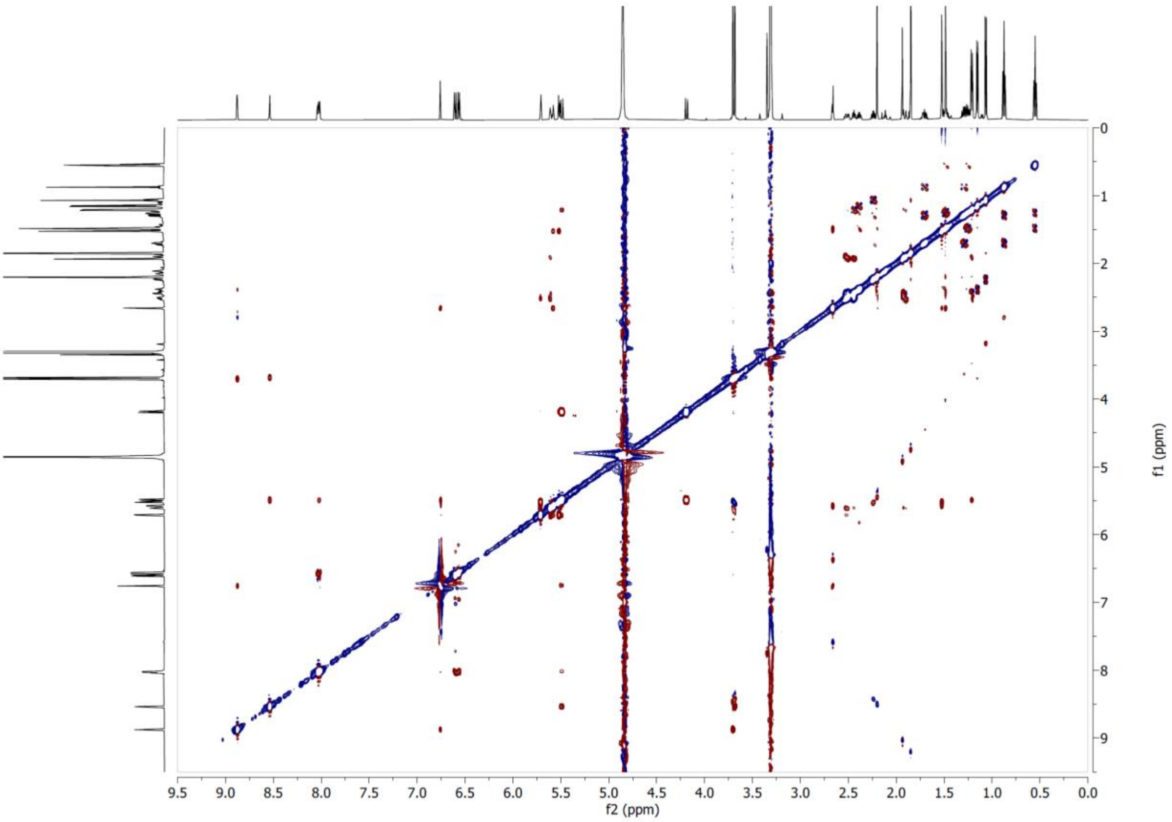

**Supplementary Figure S47**. ROESY NMR spectrum of compound **8** in $CD_3OD$ at 600 MHz.

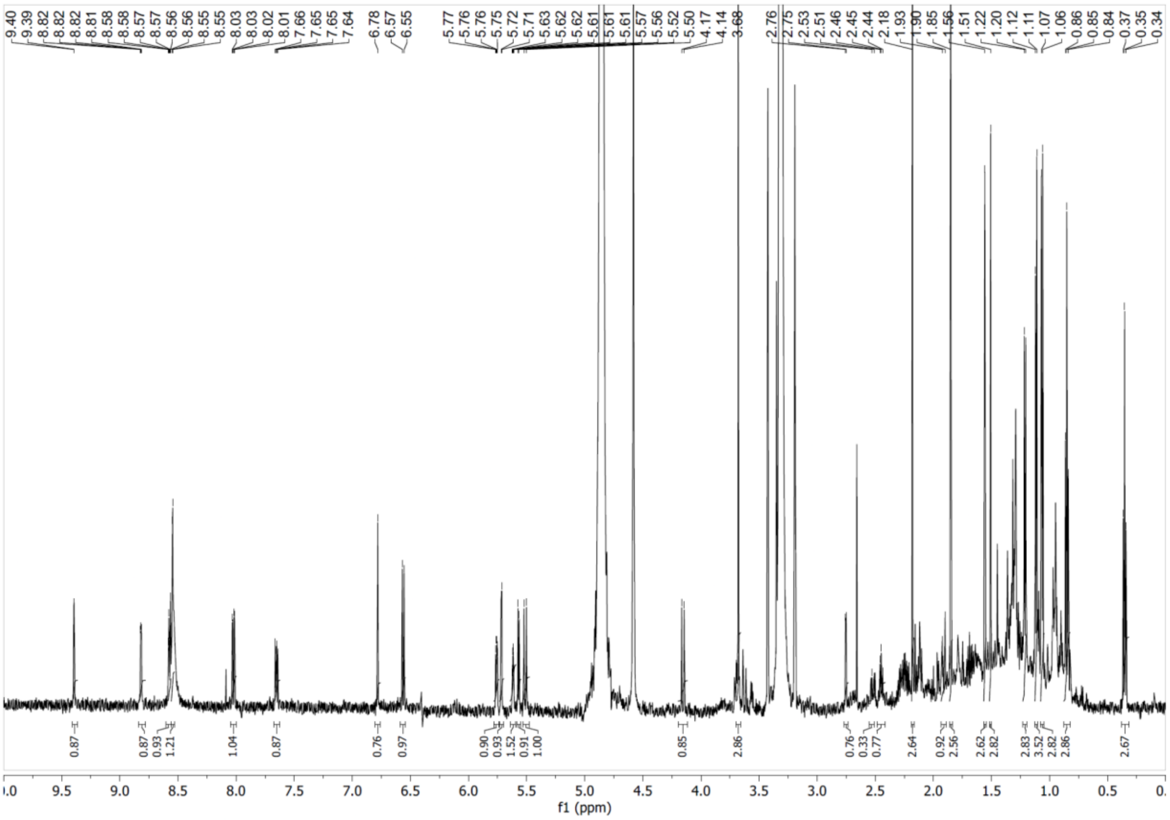

**Supplementary Figure S48**. $^{1}$H NMR spectrum of compound **9** in $CD_3OD$ at 600 MHz.

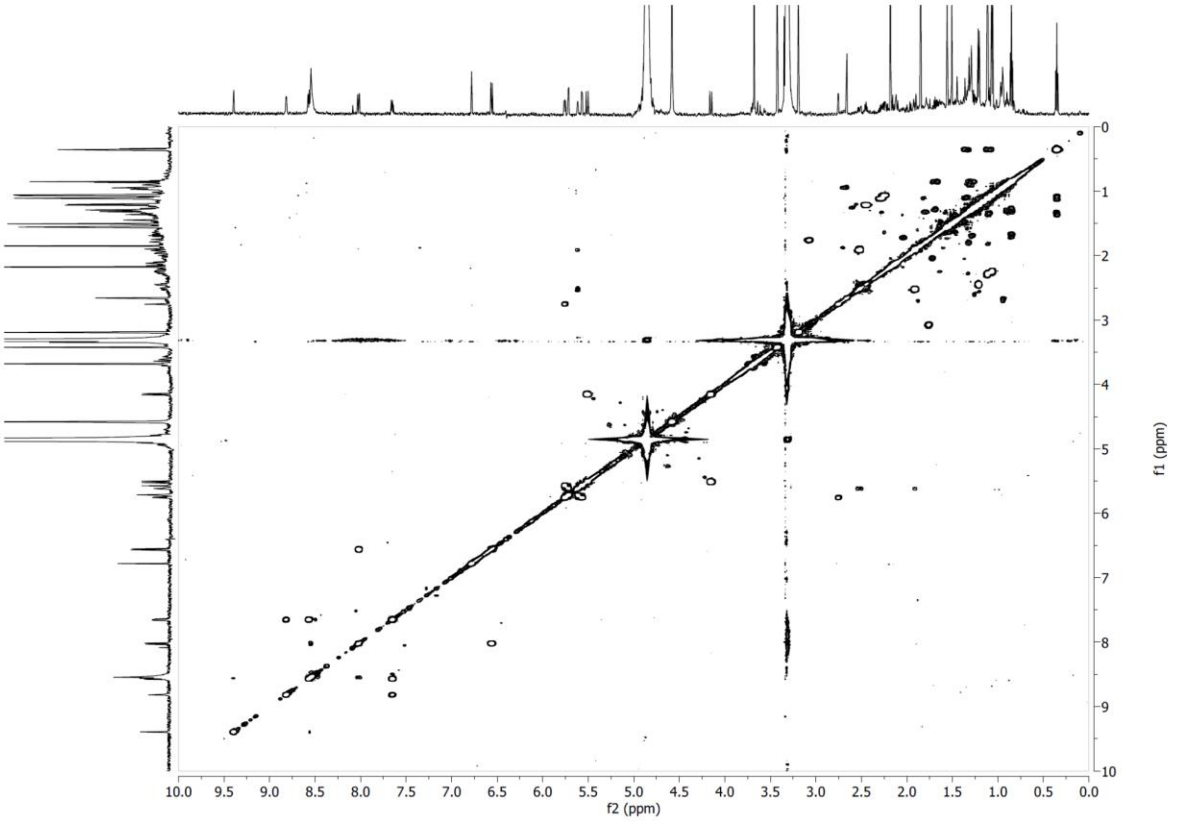


**Supplementary Figure S49**. COSY NMR spectrum of compound **9** in $CD_3OD$ at 600 MHz.

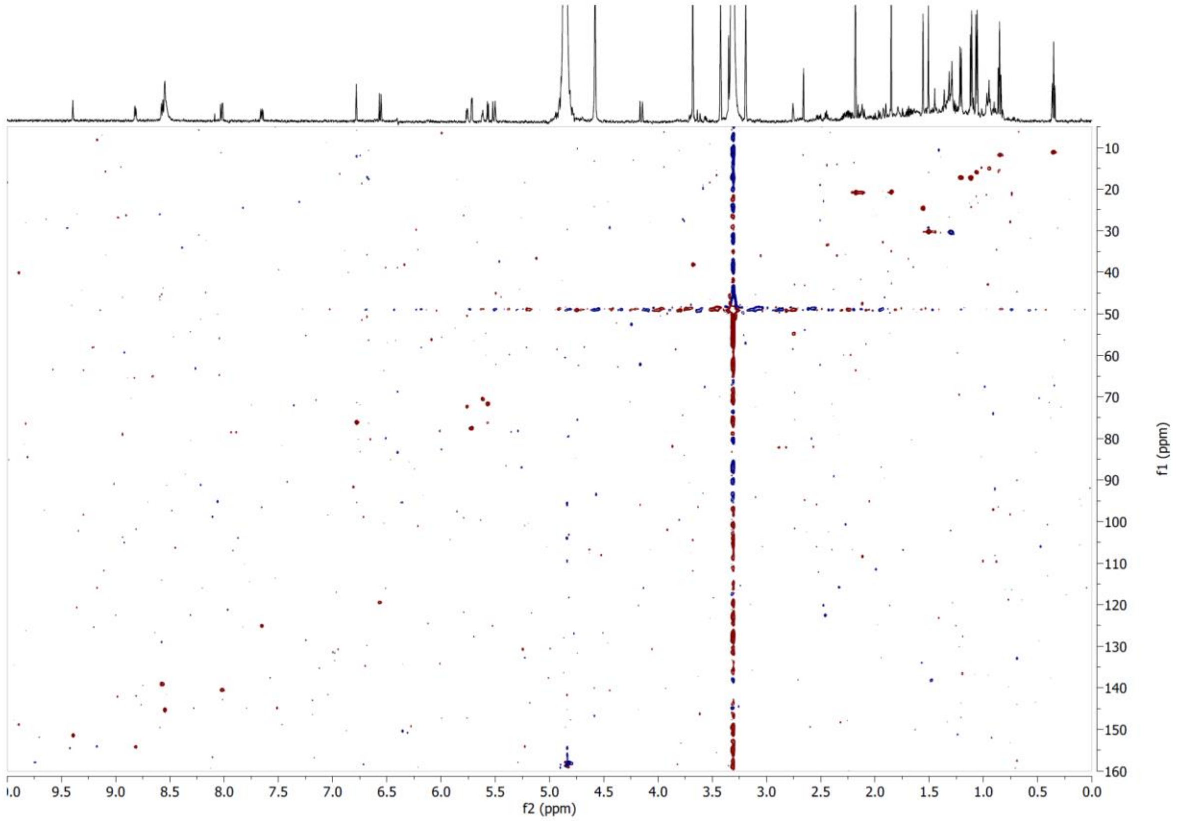


**Supplementary Figure S50**. HSQC NMR spectrum of compound **9** in $CD_3OD$.

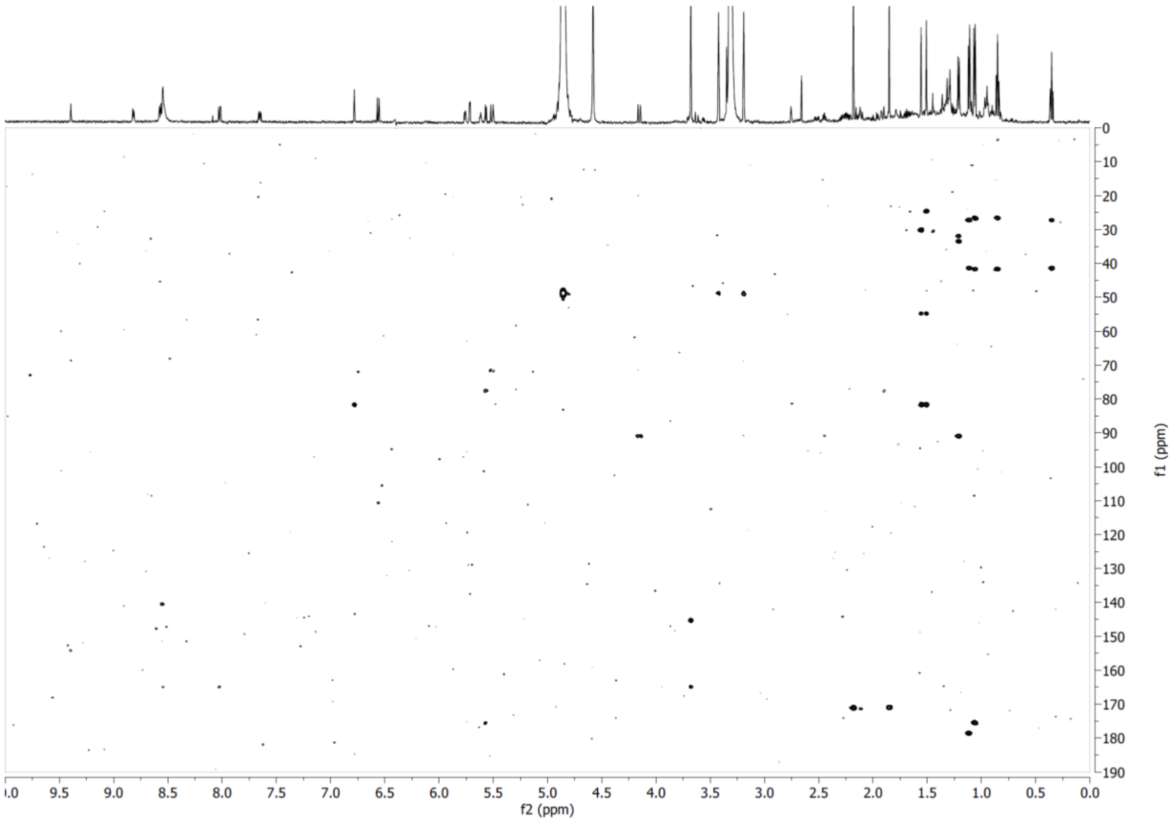


**Supplementary Figure S51**. HMBC NMR spectrum of compound **9** in $CD_3OD$.

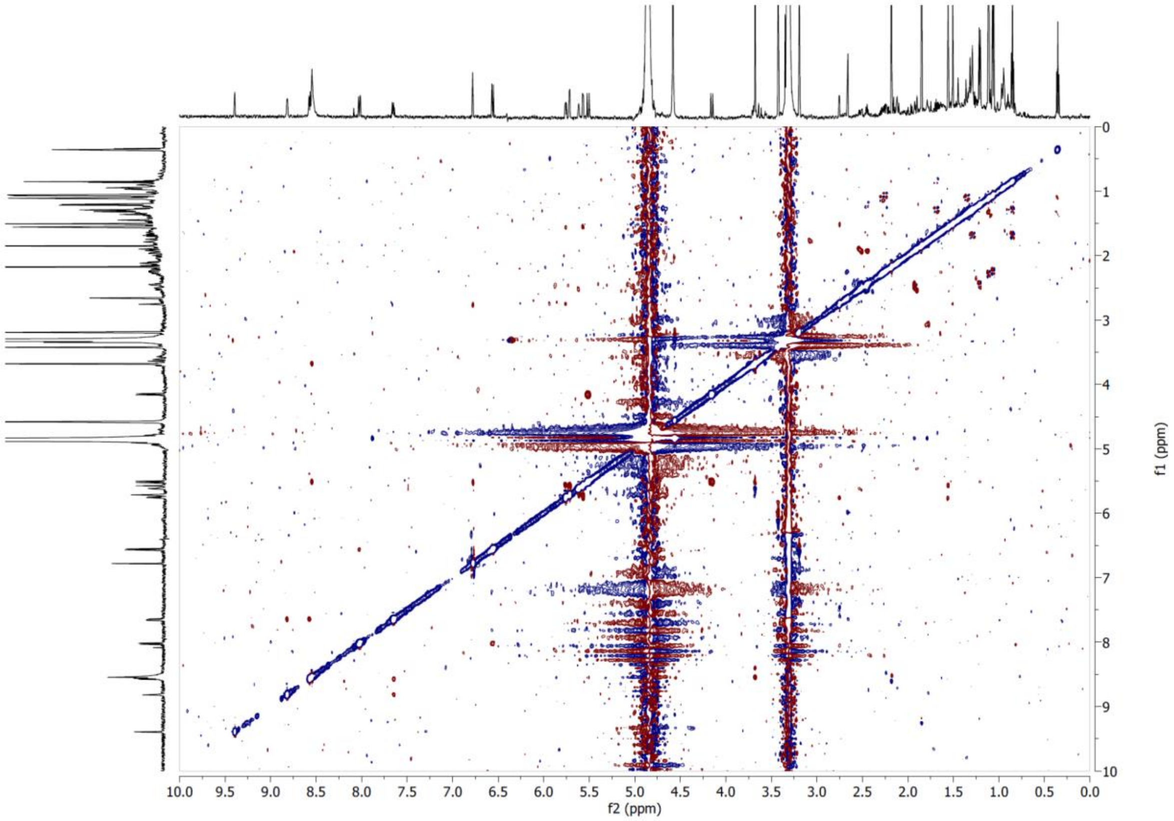


**Supplementary Figure S52**. ROESY NMR spectrum of compound **9** in $CD_3OD$ at 600 MHz.

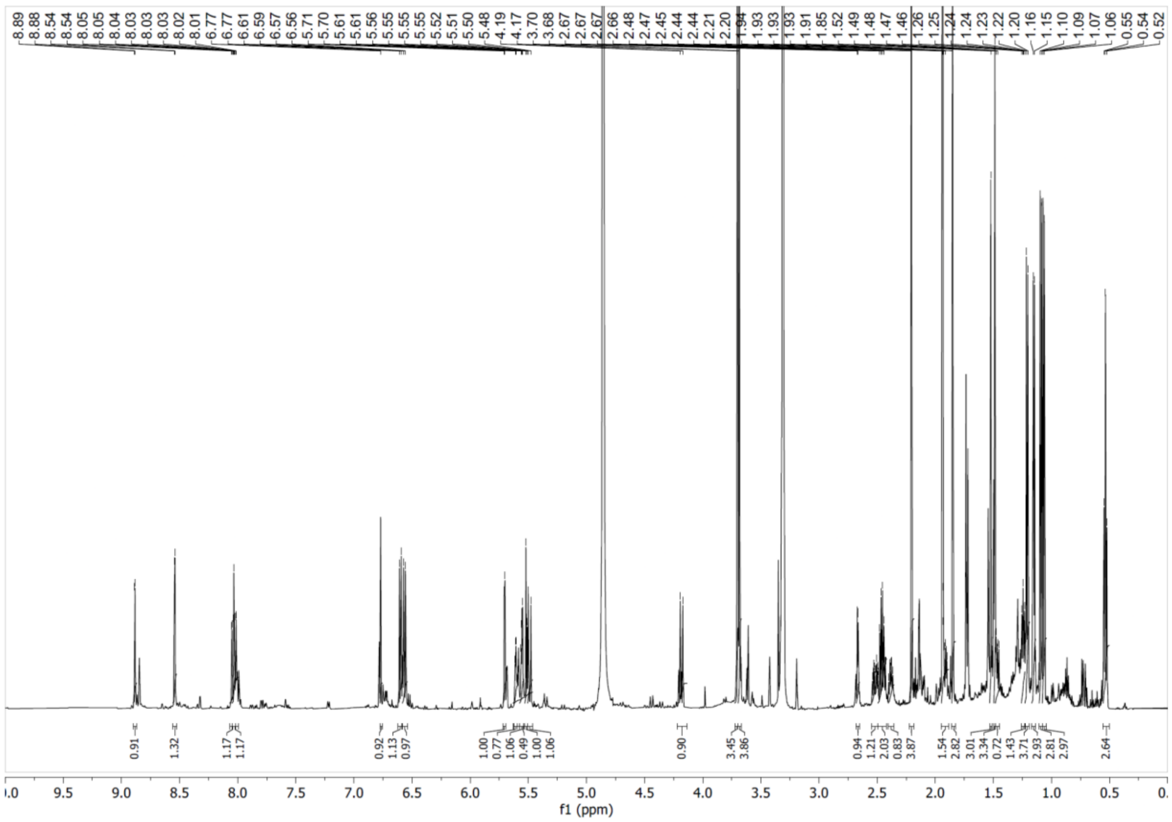

**Supplementary Figure S53**. [1]H NMR spectrum of compound **10** in $CD_3OD$ at 600 MHz.

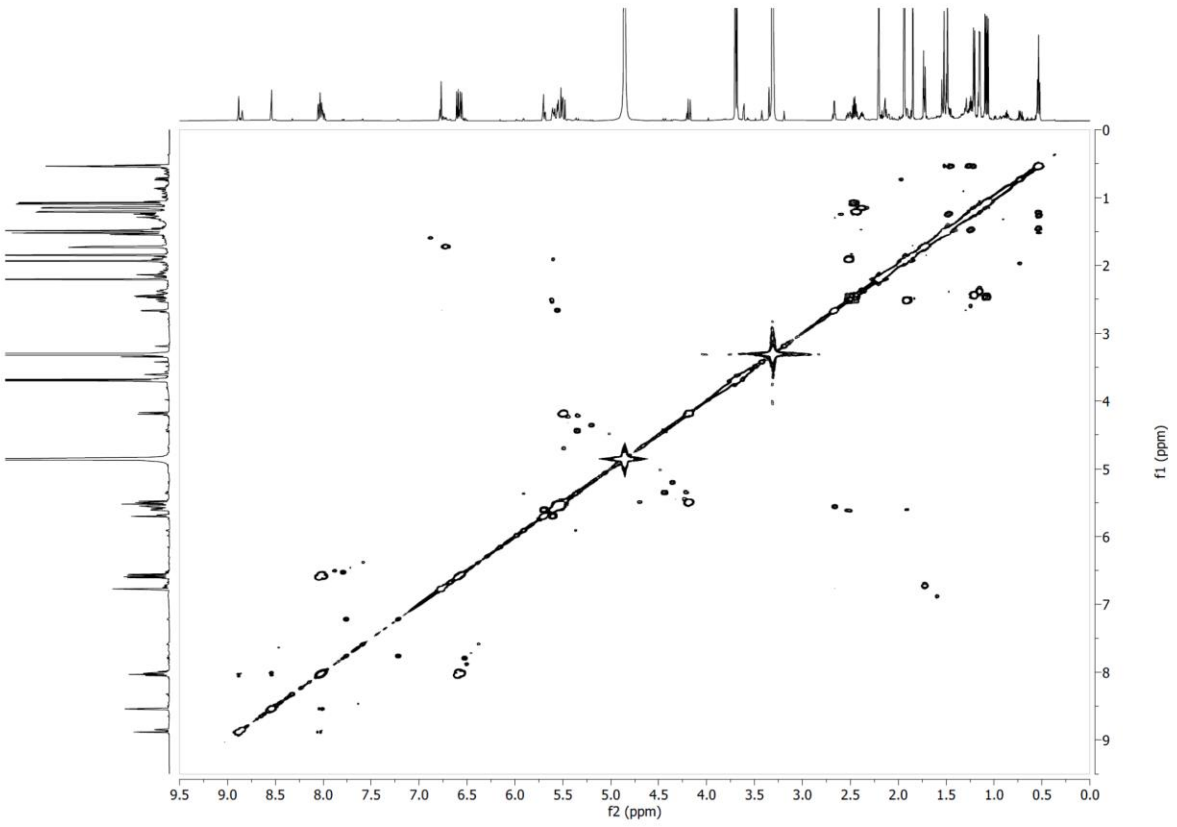

**Supplementary Figure S54**. COSY NMR spectrum of compound **10** in $CD_3OD$ at 600 MHz.

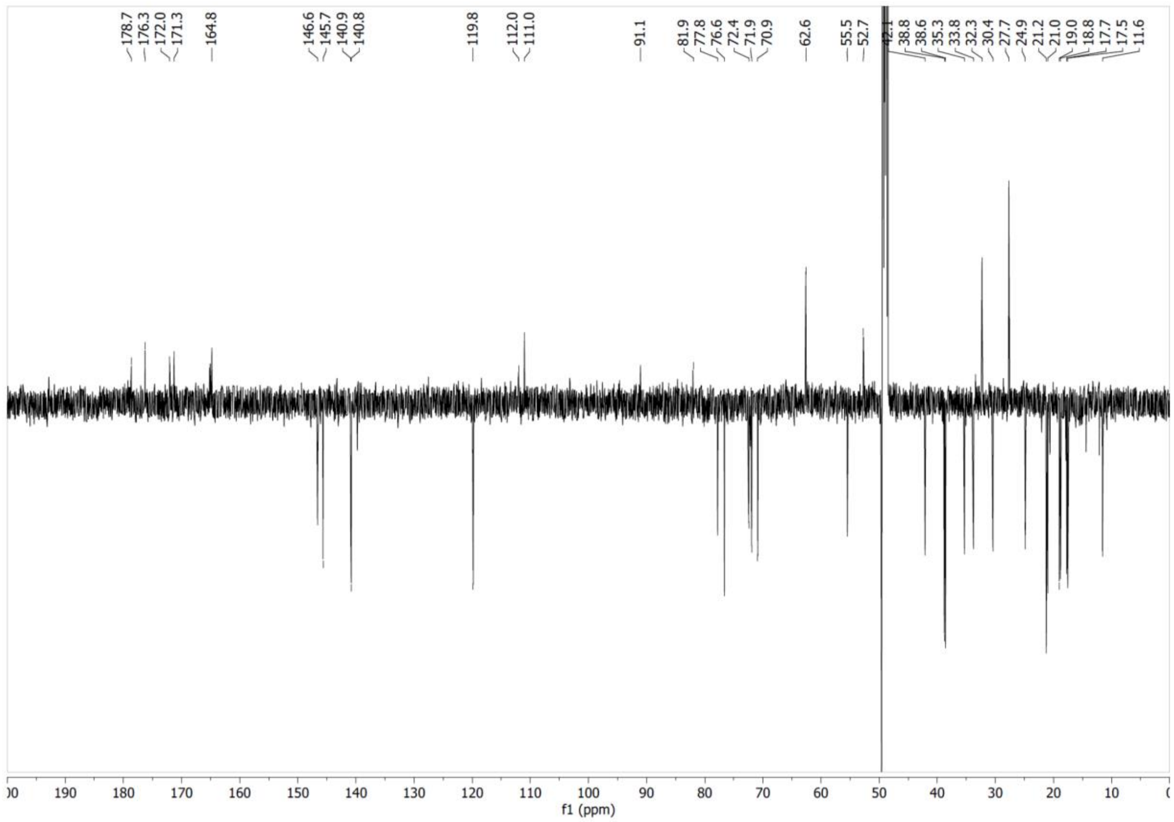


**Supplementary Figure S55**. $^{13}C$ NMR spectrum of compound **10** in $CD_3OD$ at 151 MHz.

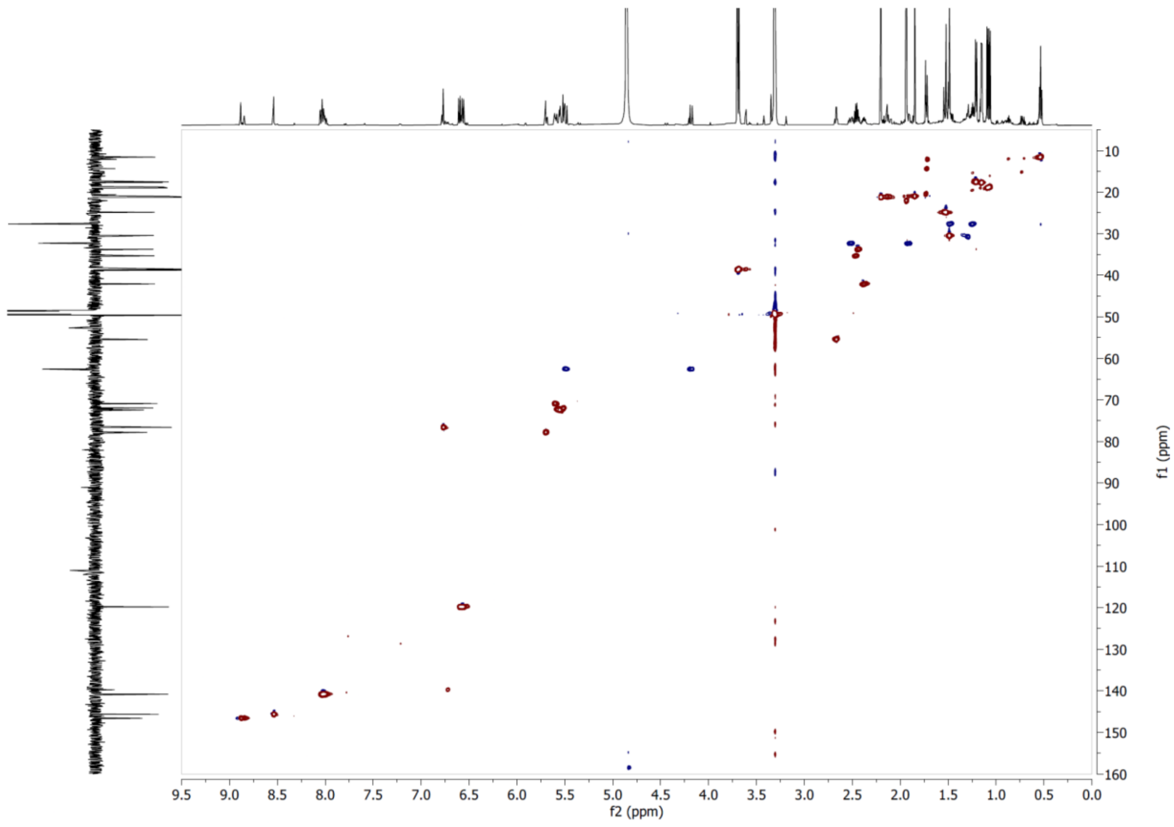


**Supplementary Figure S56**. HSQC NMR spectrum of compound **10** in $CD_3OD$.

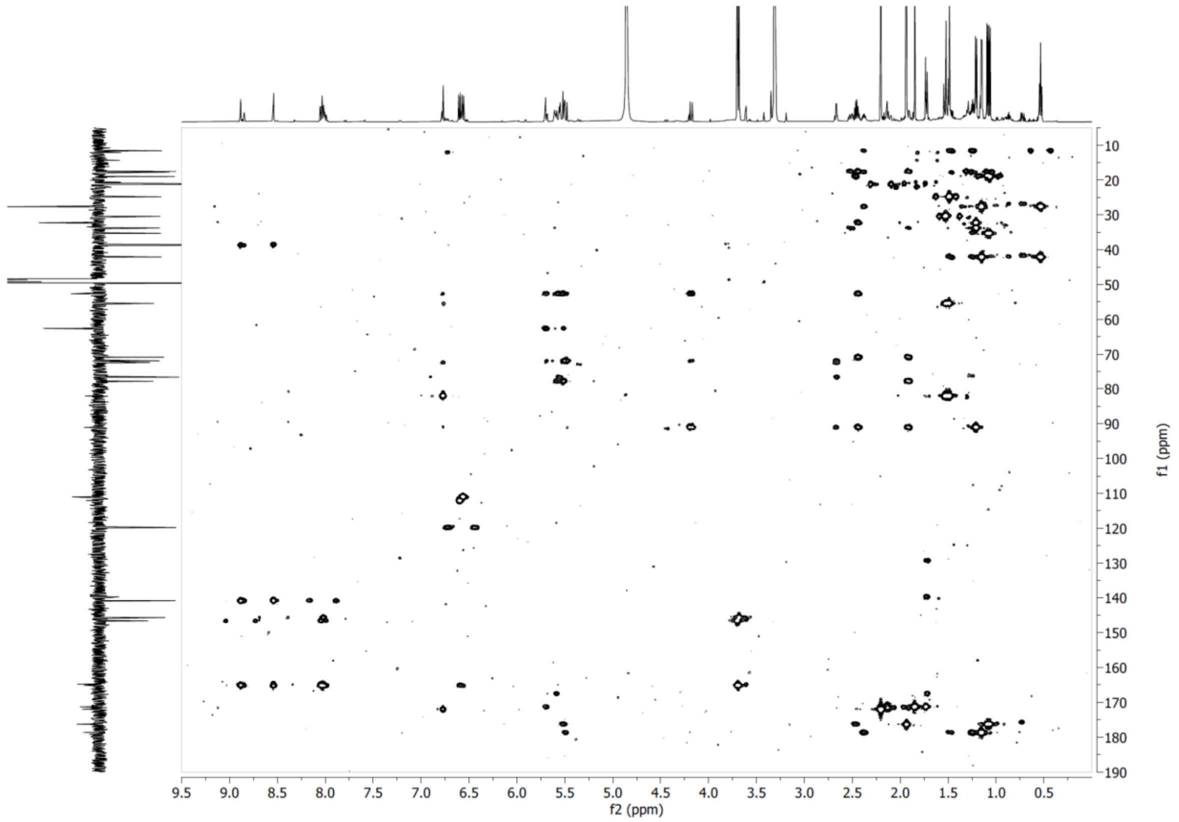


**Supplementary Figure S57**. HMBC NMR spectrum of compound **10** in $CD_3OD$.

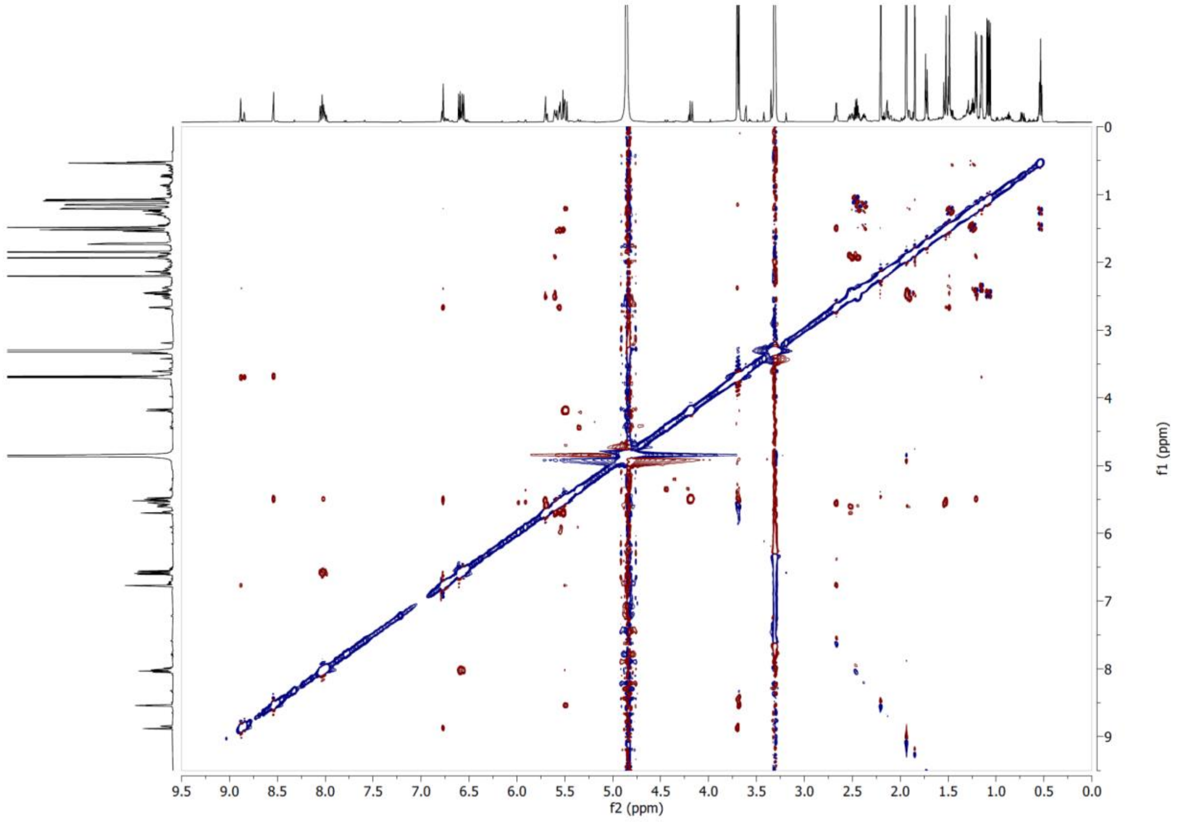


**Supplementary Figure S58**. ROESY NMR spectrum of compound **10** in $CD_3OD$ at 600 MHz.

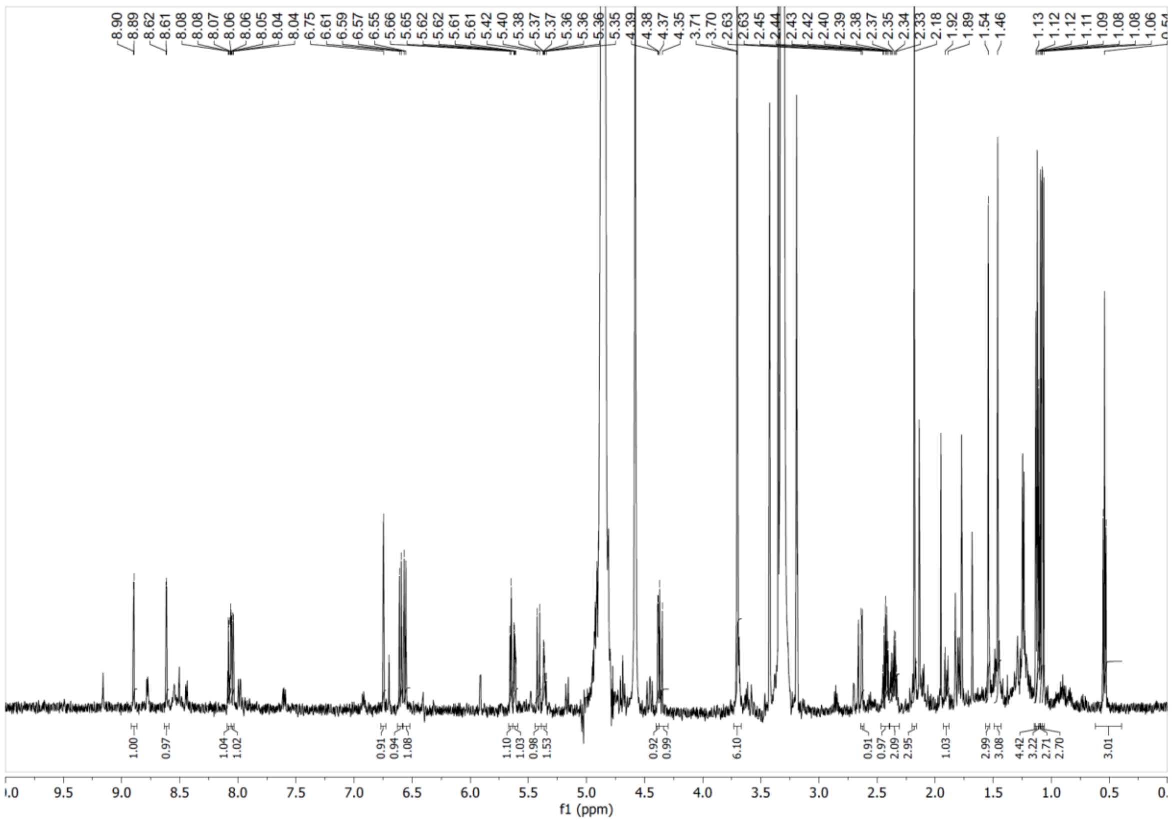


**Supplementary Figure S59**. [1]H NMR spectrum of compound **11** in $CD_3OD$ at 600 MHz.

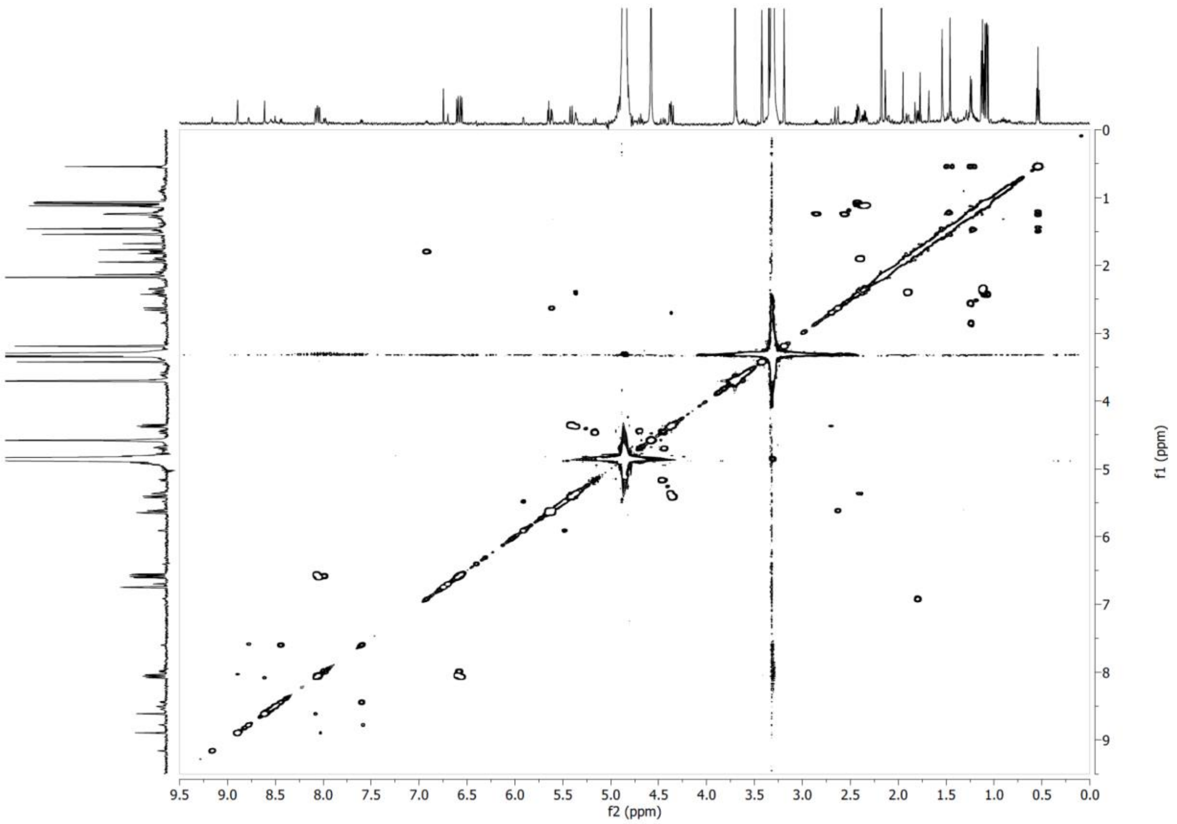


**Supplementary Figure S60**. COSY NMR spectrum of compound **11** in $CD_3OD$ at 600 MHz.

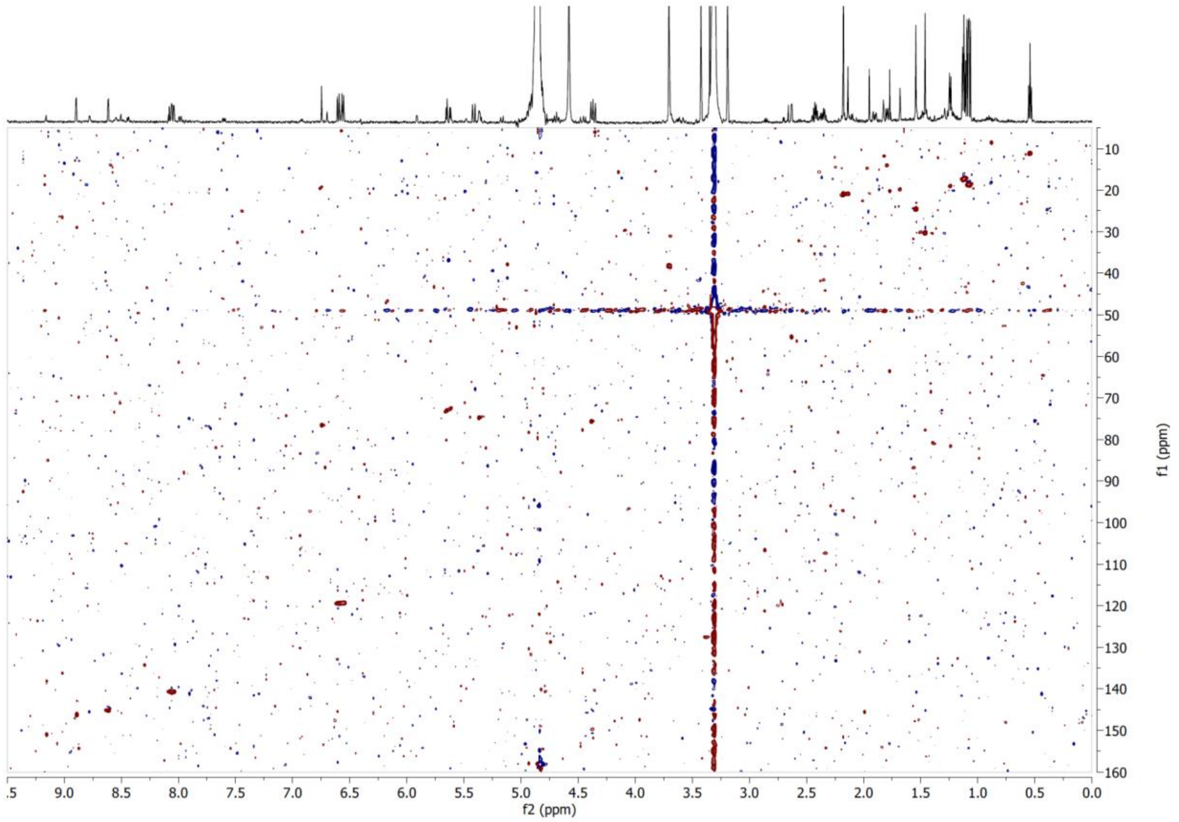


**Supplementary Figure S61**. HSQC NMR spectrum of compound **11** in $CD_3OD$.

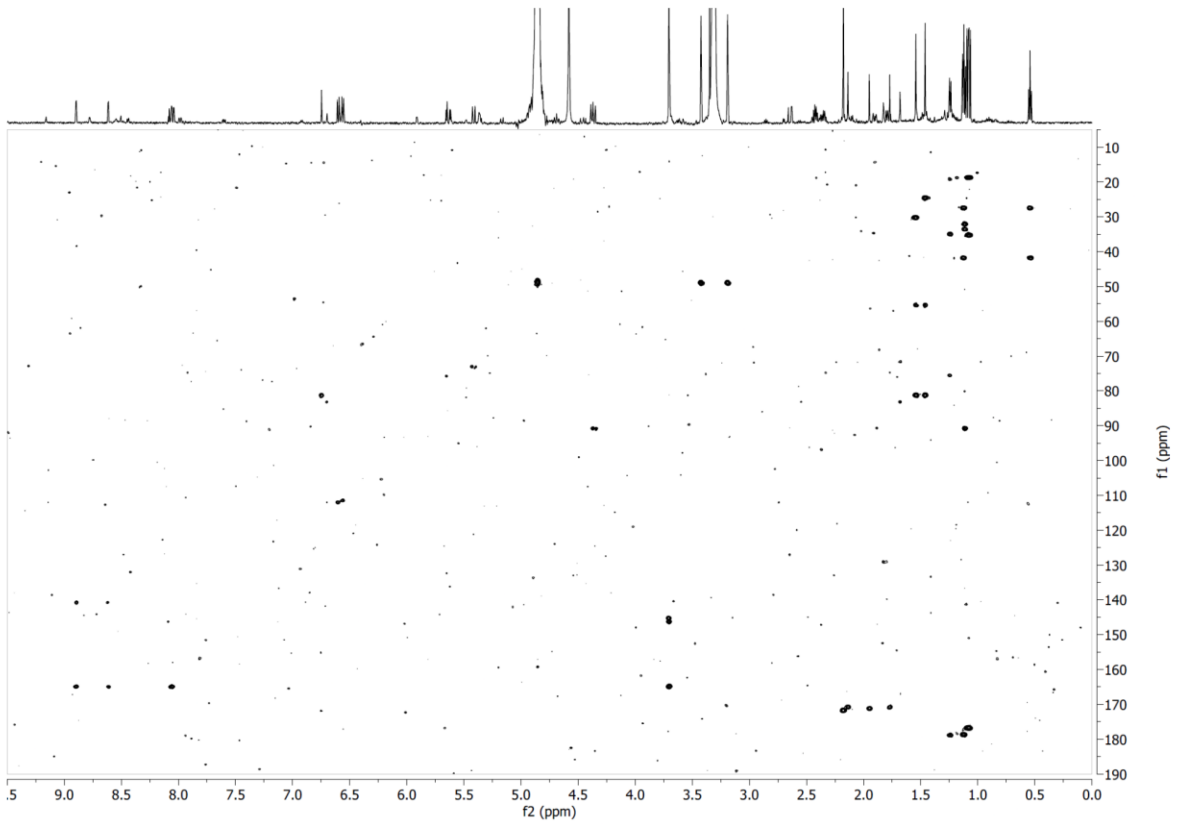


**Supplementary Figure S62**. HMBC NMR spectrum of compound **11** in $CD_3OD$.

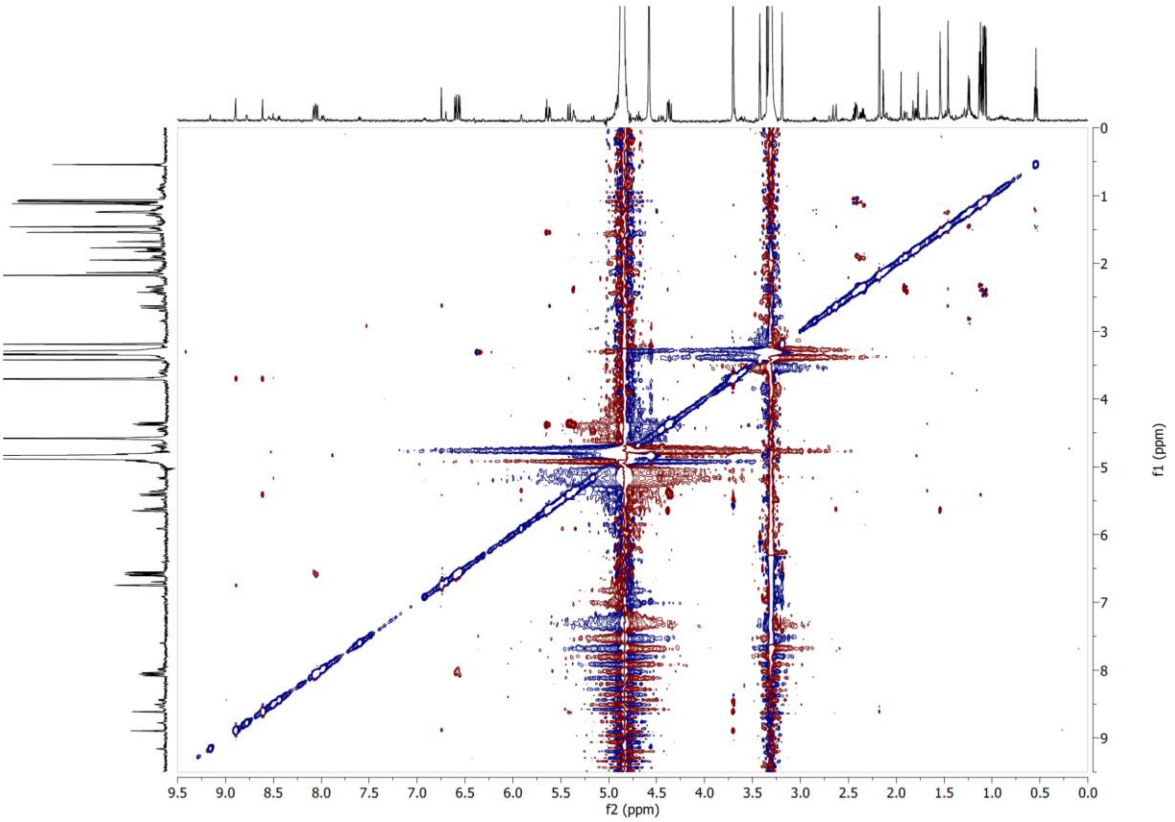

**Supplementary Figure S63**. ROESY NMR spectrum of compound **11** in $CD_3OD$ at 600 MHz.

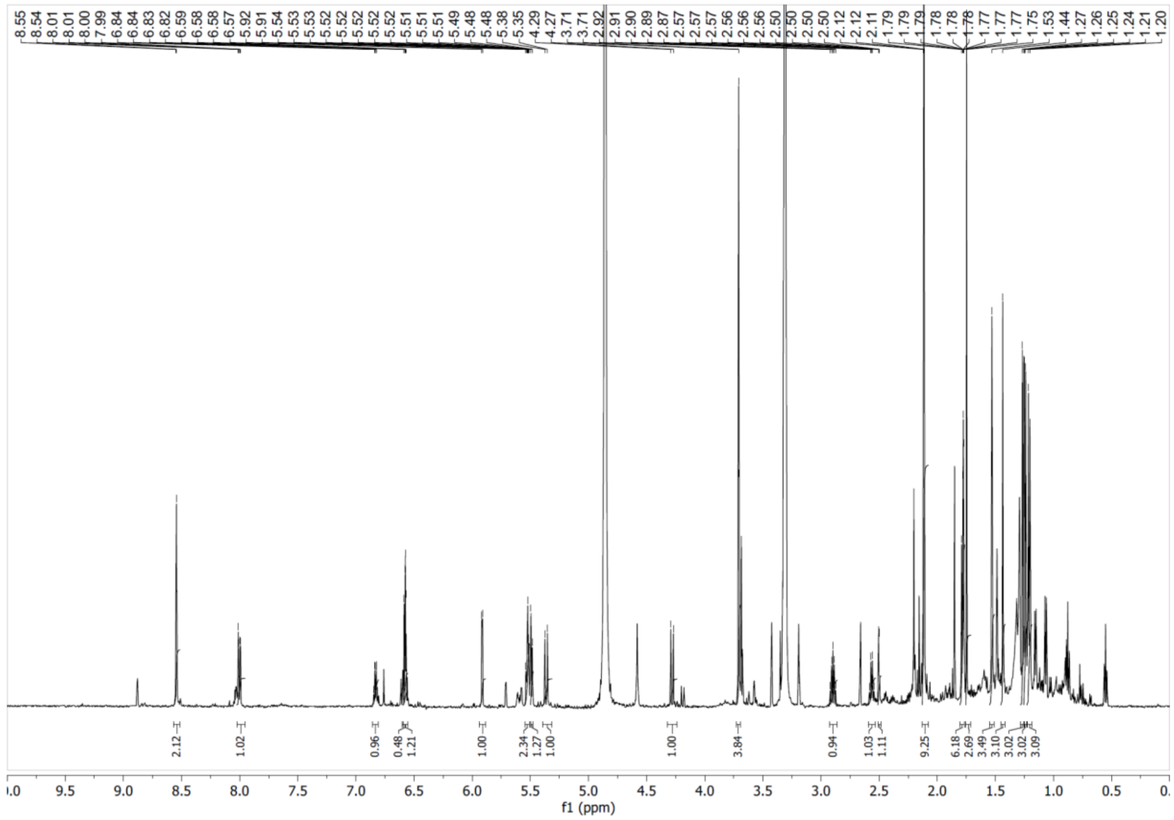

**Supplementary Figure S64**. $^1H$ NMR spectrum of compound **12** in $CD_3OD$ at 600 MHz.

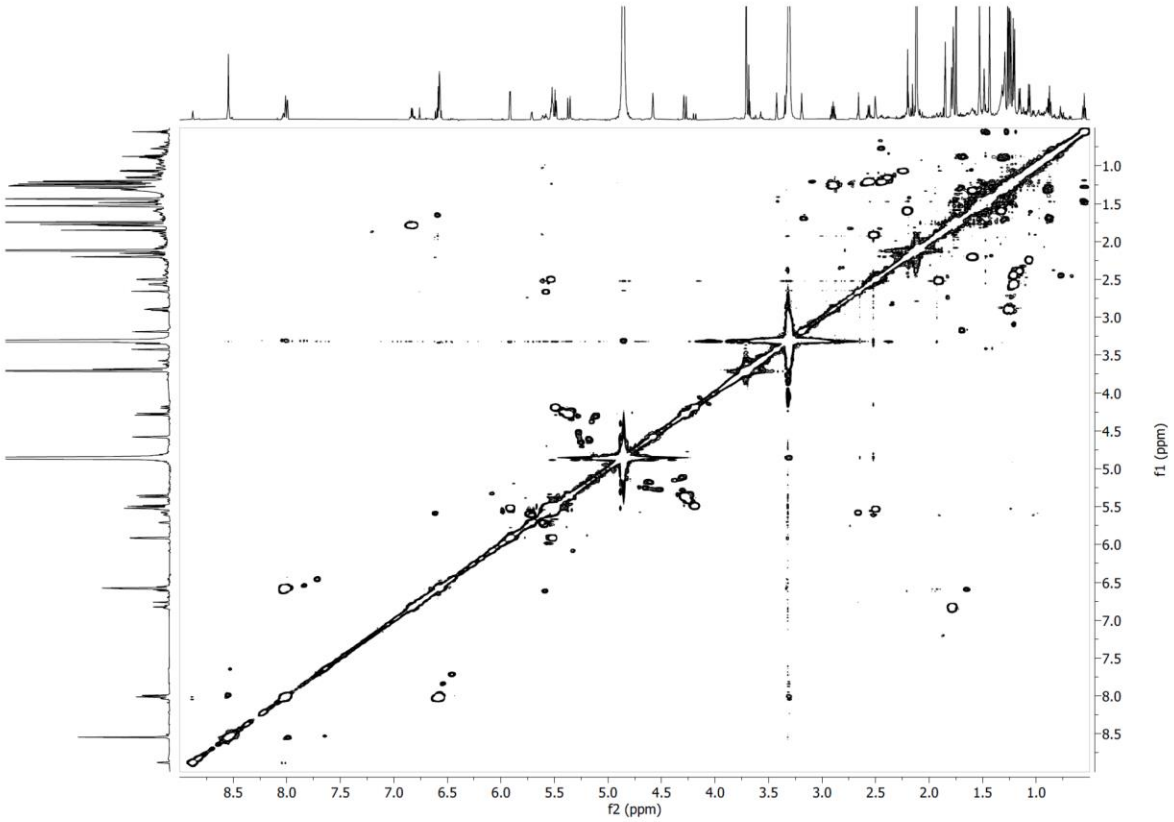


**Supplementary Figure S65**. COSY NMR spectrum of compound **12** in $CD_3OD$ at 600 MHz.

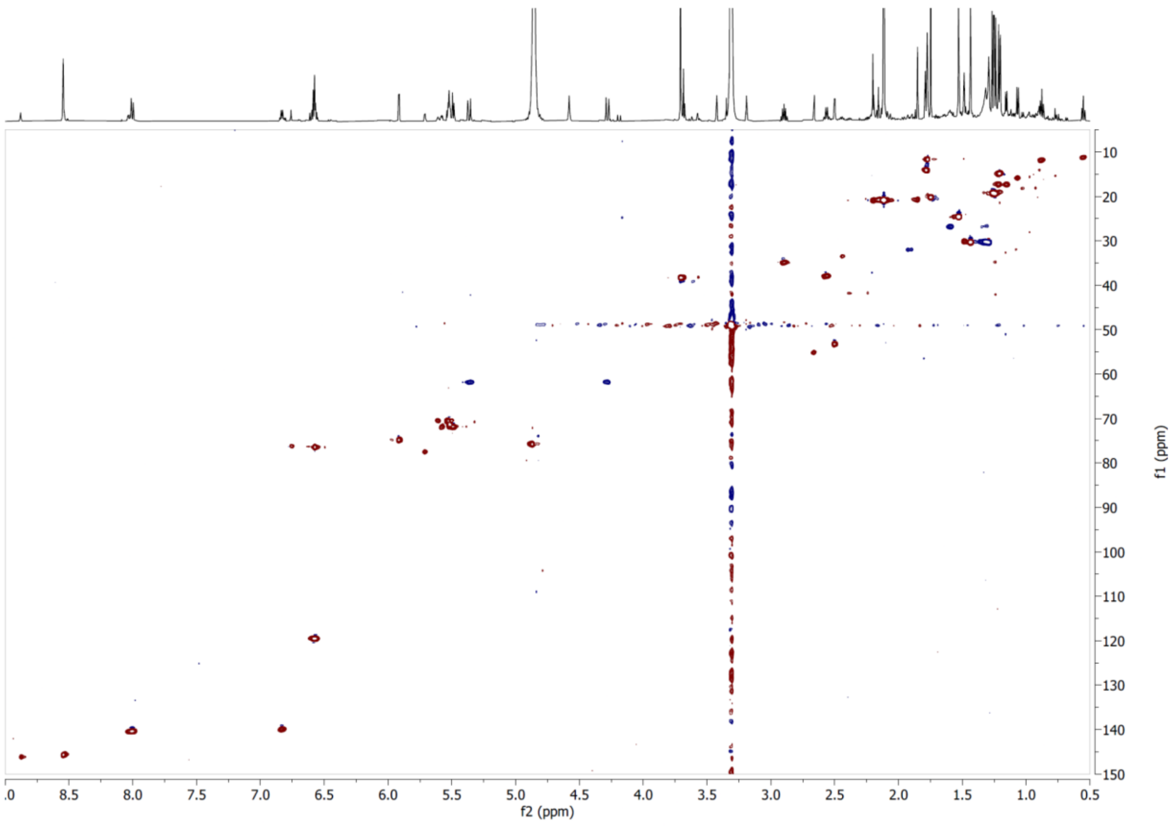


**Supplementary Figure S66**. HSQC NMR spectrum of compound **12** in $CD_3OD$.

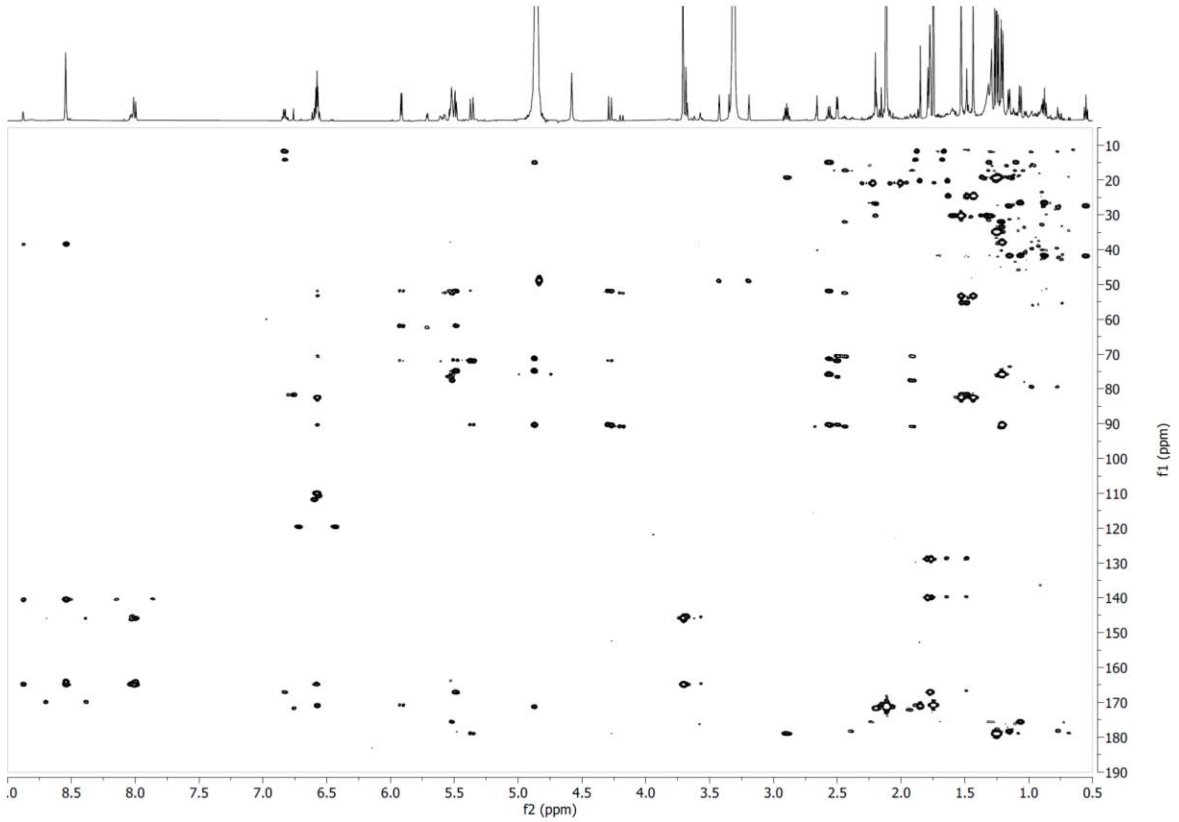


**Supplementary Figure S67**. HMBC NMR spectrum of compound **12** in $CD_3OD$.

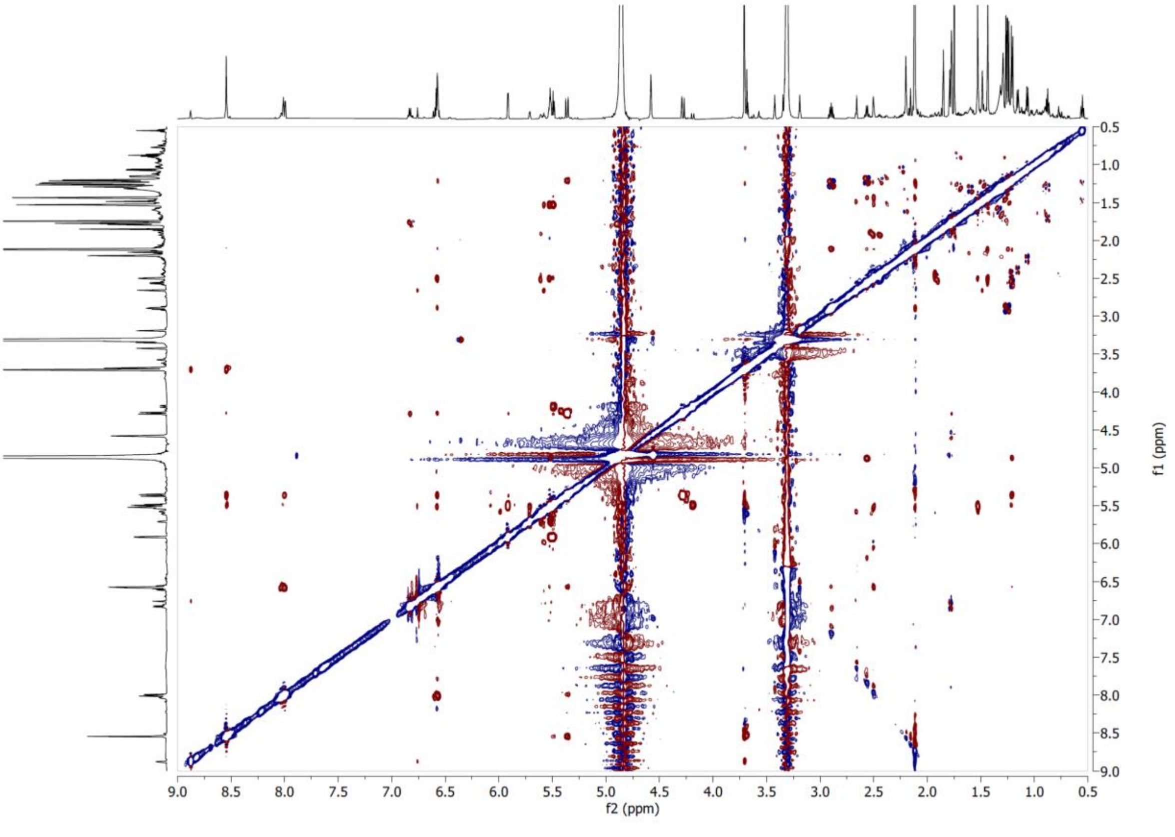


**Supplementary Figure S68**. ROESY NMR spectrum of compound **12** in $CD_3OD$ at 600 MHz.

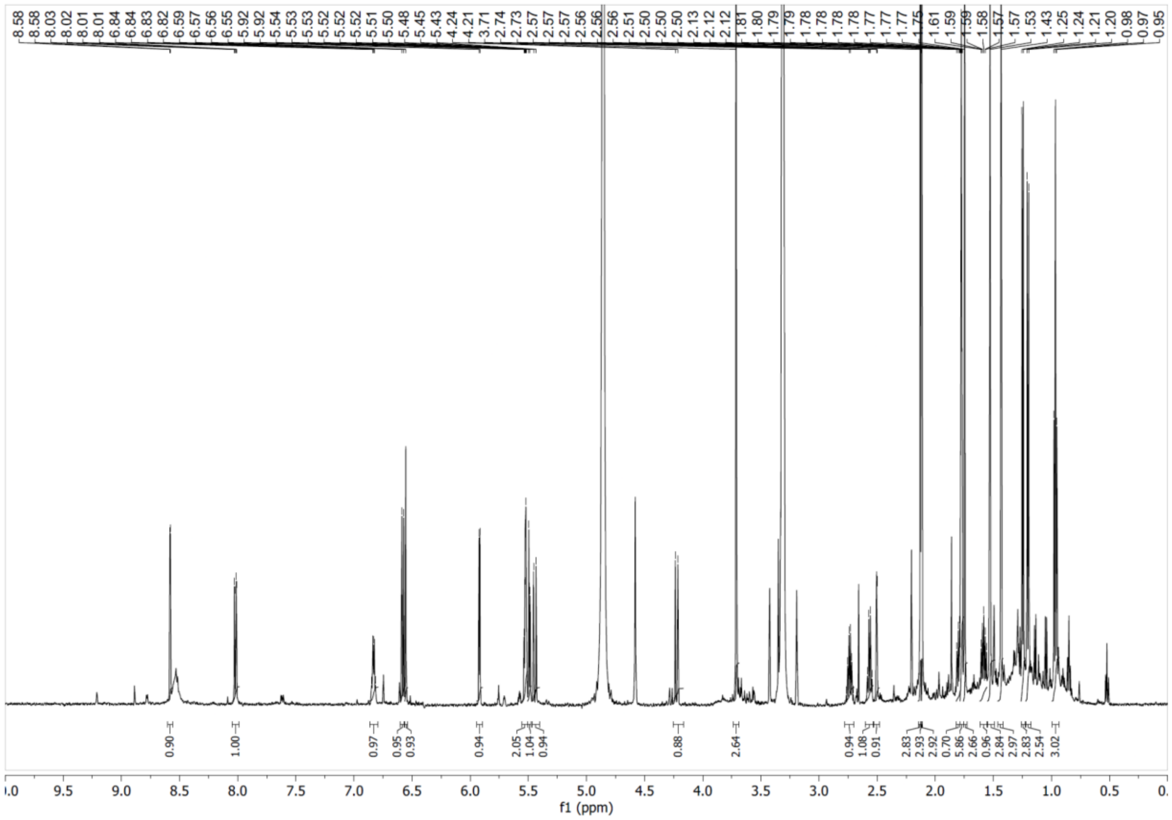


**Supplementary Figure S69**. $^{1}H$ NMR spectrum of compound **13** in $CD_3OD$ at 600 MHz.

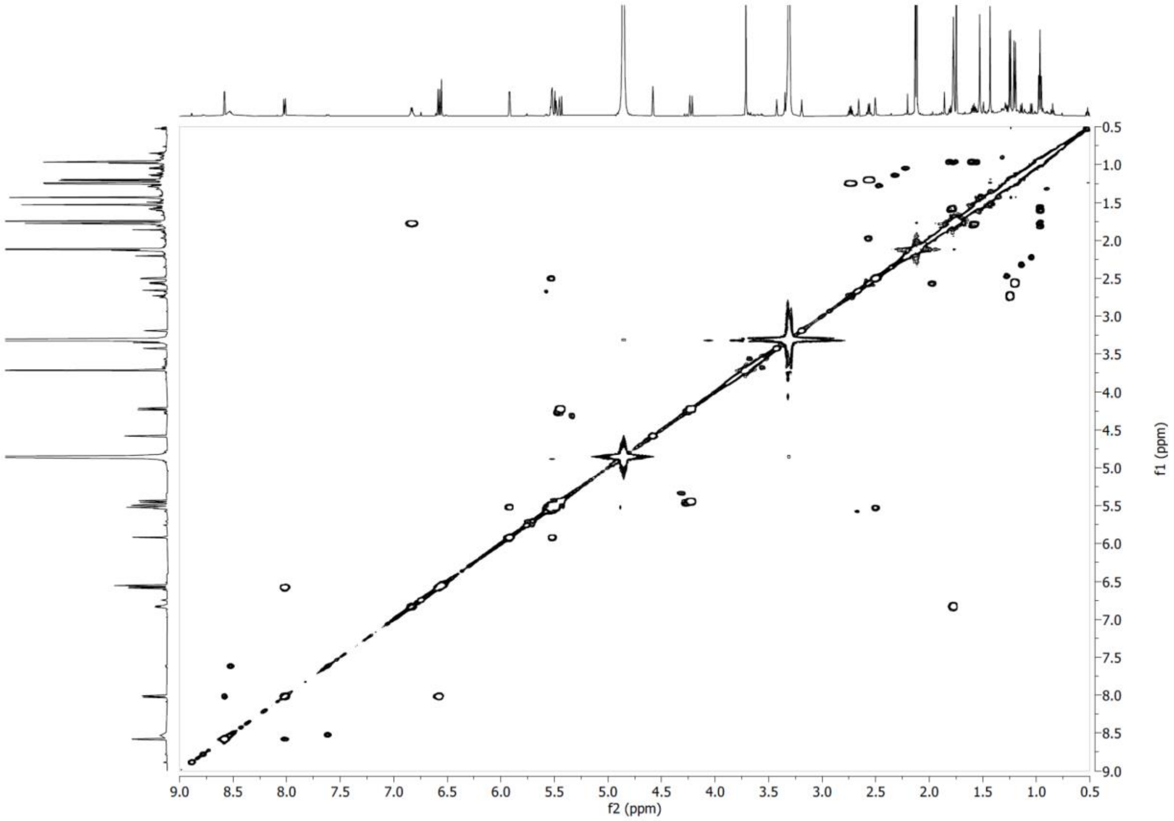


**Supplementary Figure S70**. COSY NMR spectrum of compound **13** in $CD_3OD$ at 600 MHz.

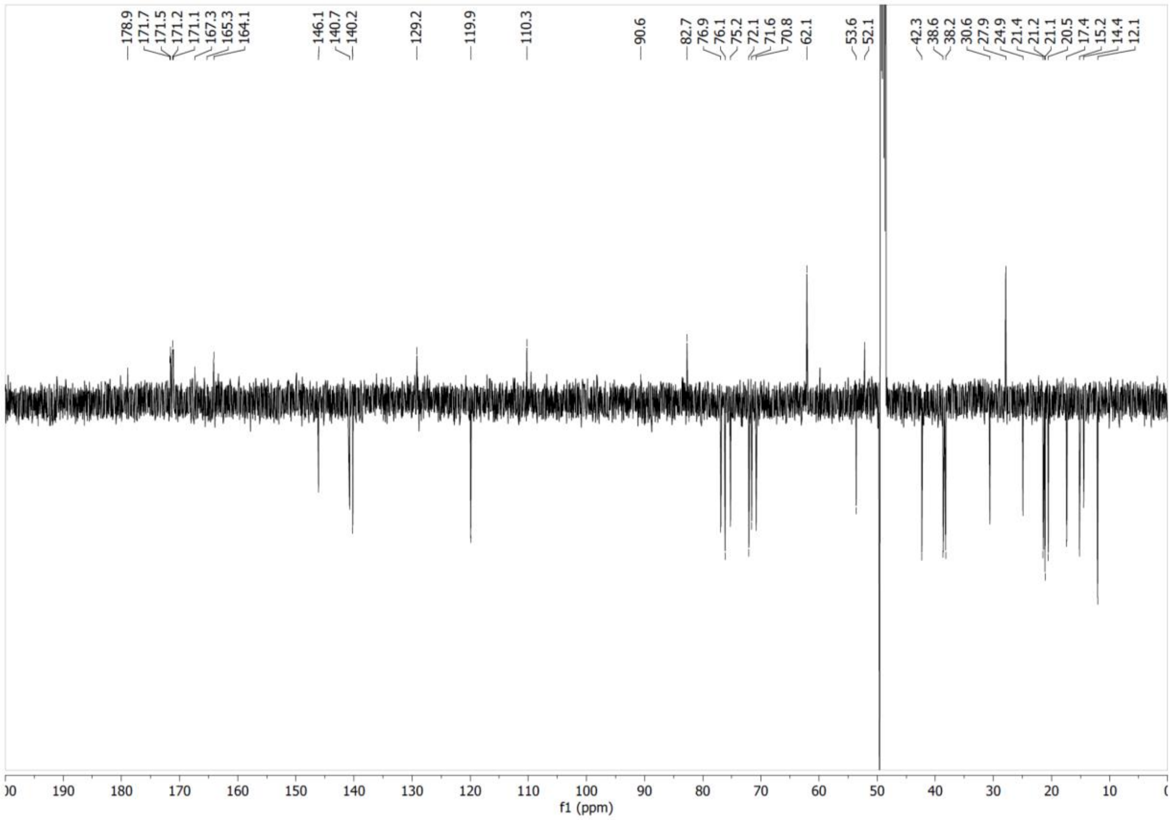


**Supplementary Figure S71**. $^{13}$C NMR spectrum of compound **13** in $CD_3OD$ at 151 MHz.

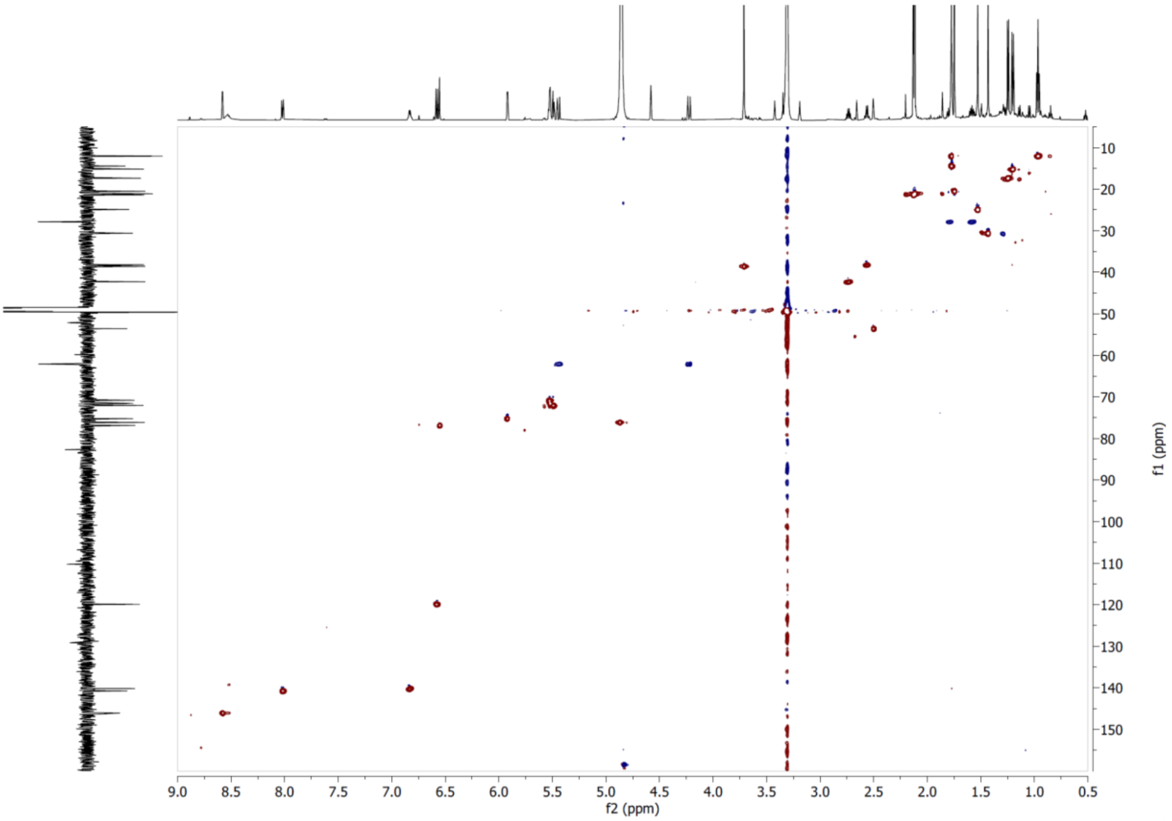


**Supplementary Figure S72**. HSQC NMR spectrum of compound **13** in $CD_3OD$.

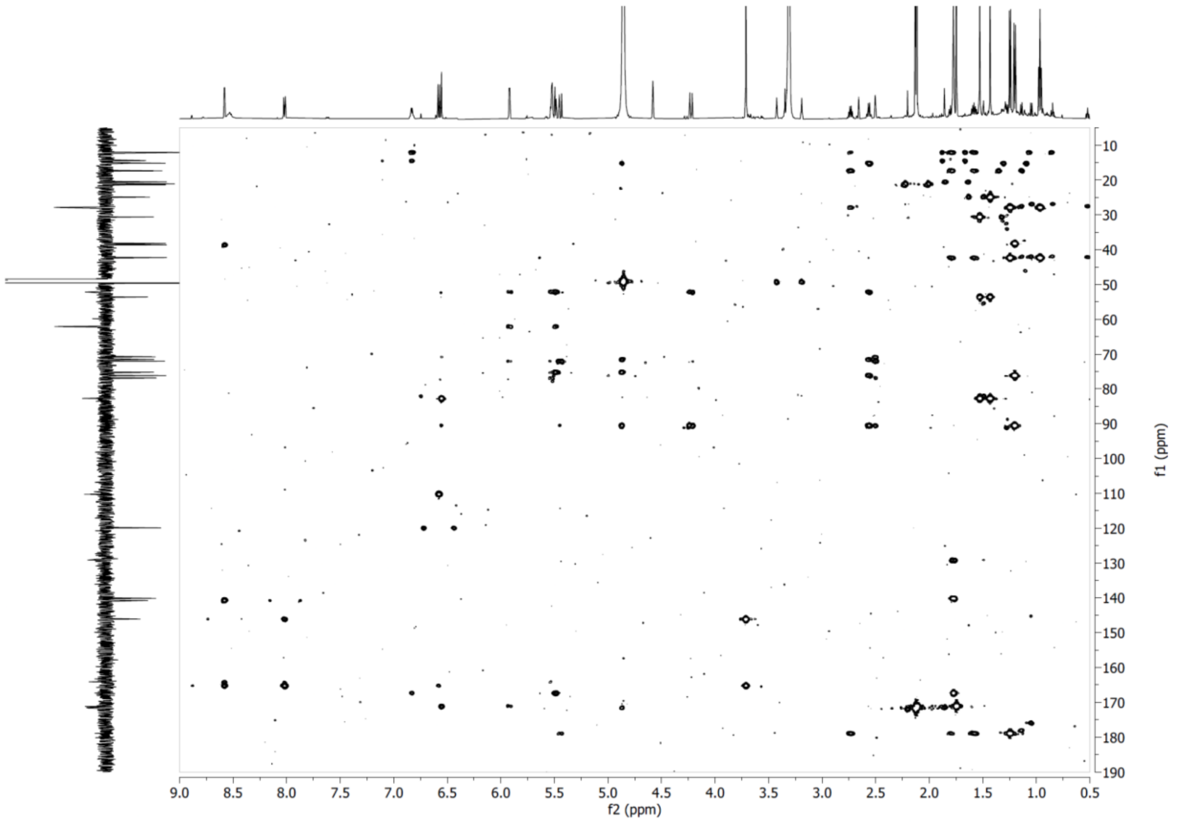


**Supplementary Figure S73**. HMBC NMR spectrum of compound **13** in $CD_3OD$.

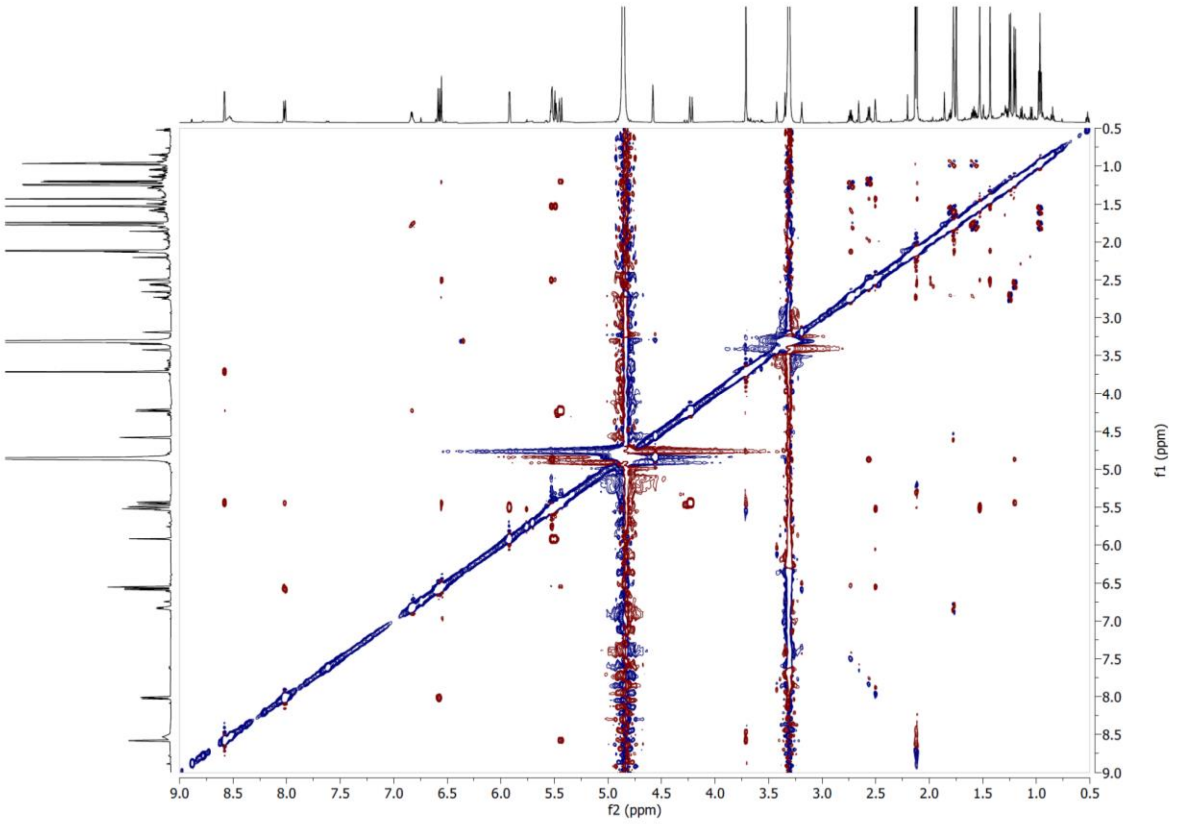


**Supplementary Figure S74**. ROESY NMR spectrum of compound **13** in $CD_3OD$ at 600 MHz.

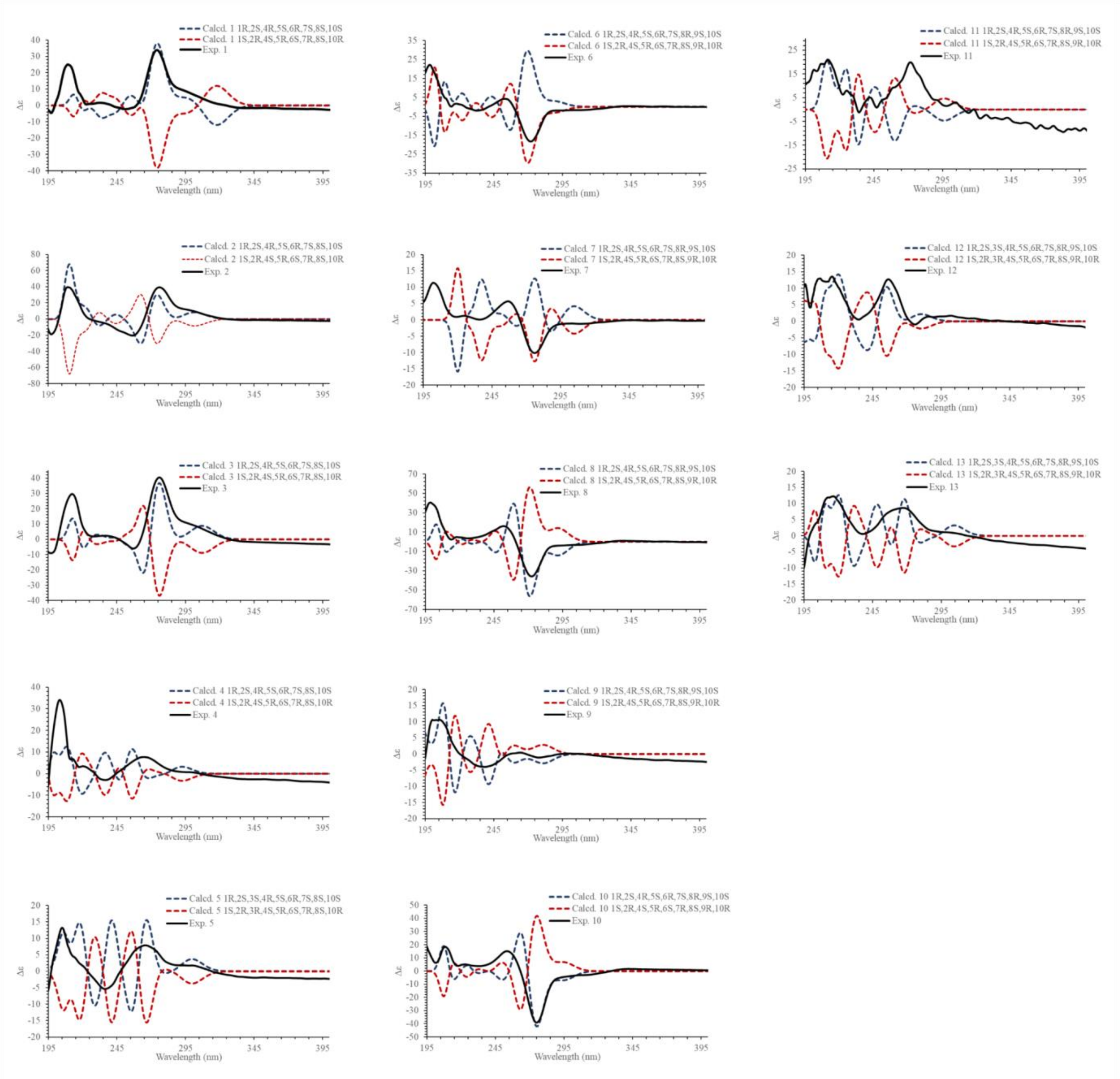


**Supplementary Figure S75**. Experimental and B3LYP/def2svp//B3LYP/6-31G(d,p) calculated spectra from compounds **1-13** in acetonitrile.

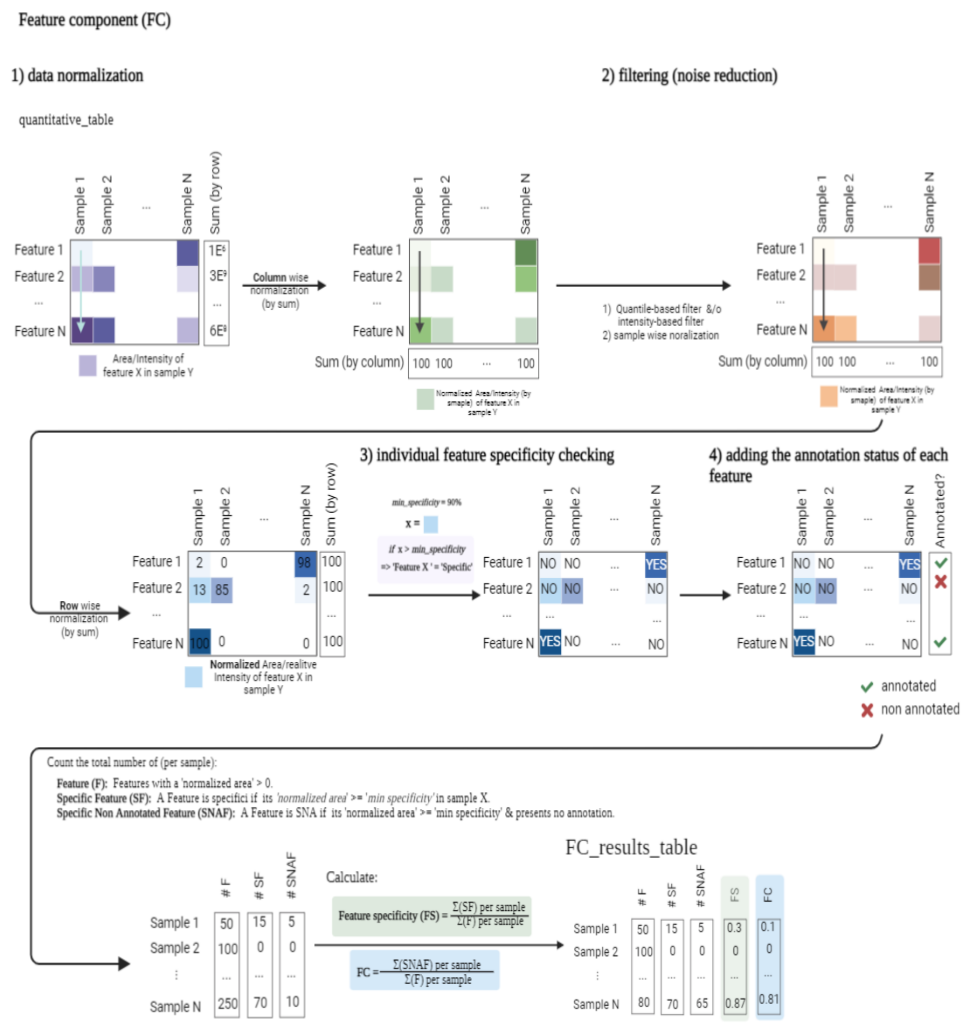


**Supplementary Figure S76.** Graphical representation of the steps followed for calculating the Feature Component (FC). The FC takes the feature-aligned quantification table obtained from the treatment of the raw data and the identification results from GNPS and other in silico tools as input. 1) The relative areas, based on the Total Ion Chromatogram, are normalized row-wise by total sum, to obtain relative percentages. 2) the resulting data frame is used to get the filename (or identifier preferred) of the sample where each ion is more concentrated (higher percentage). 3) The annotation status of each feature is linked to the results from point 2. For this particular step, several inputs from different annotation bioinformatics tools can be considered. Results from the GNPS platform are higher in relevance since they are based on an experimental database. *In silico* dereplication results from ISDB or SIRIUS can be included. If so, the script will consider the annotation status of a particular feature like 'annotated' if it presents identification in any of the inputs used. 4) The specificity status of each feature is checked according to the *min_specificity* threshold fixed by the user. If the area of a particular feature is equal to or higher than *min_specficity* then the feature is consider as 'specific'. 5) once both conditions are established for each feature (annotations and specificity status), the feature component ratio is calculated. The ratio is the division of the total of specific-non annotated features in a sample by the total of features present in it. Additionally, the Feature specificity is calculated in a similar way, but without taking into account the annotation status, only the specificity status. This means that the features specificity corresponds to the division of the total of specific features in a sample by the total of features present in the sample. The difference between both values indicates the percentage of specific ions in the sample presenting an annotation. The molecular formula prediction ratio (MF_prediction_ratio) is calculated in the same way, but adding a particular condition, ‘the feature presents a good quality predicted molecular formula’. This molecular formula comes from the dereplication process using the Sirius software. It proposes a molecular formula using the module ZODIAC. So, MF prediction ratio corresponds to the division of the total of specific-non annotated-’has a good quality MF’ features in a sample by the total of features present in the sample. Refer to **Supplementary Table 1** for actual results.

<>

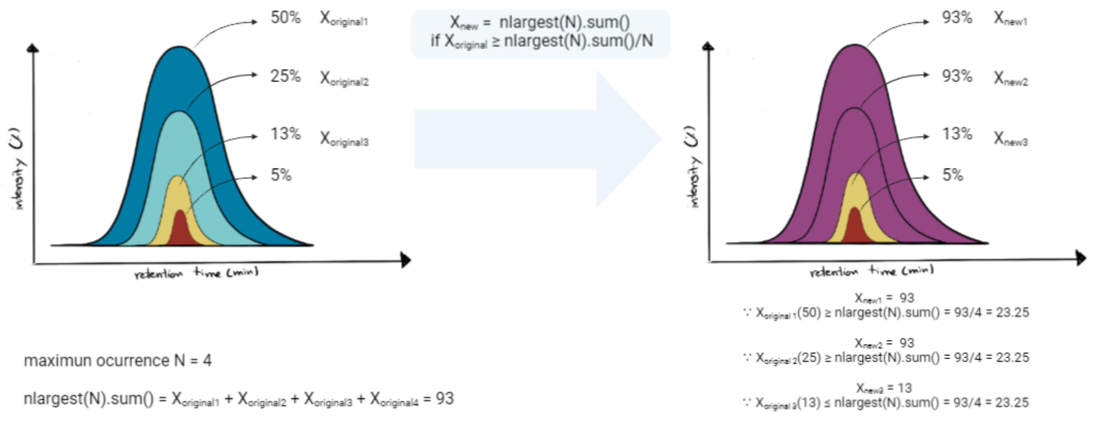


**Supplementary Figure S77**. Graphical representation of the data handling is performed when multiple samples with the same species are present in the dataset. The user will define the maximum occurrence of the species (N), this value will be used to get the sum of the N largest values for each feature in the table [nlargest(N).sum]. As shown in the figure, as an example for N = 4, the nlargest(N).sum = 93%, which means that this feature is potentially specific to the species. Because the selection and counting of specific features are done sample-wise, the original values of the feature in the N largest contributing samples are temporarily replaced by the ($X_{new}$ = nlargest(N).sum). This, is only done if the original value ($X_{original}$) is higher than nlargest(N).sum/N. In the example, only the 2 maximum values were replaced accordingly. This restriction avoids introducing the presence of a specific feature in a sample where the feature is in a low proportion (way more difficult to isolate). After the replacement of the values, the calculations for FS and FC are done as previously described.

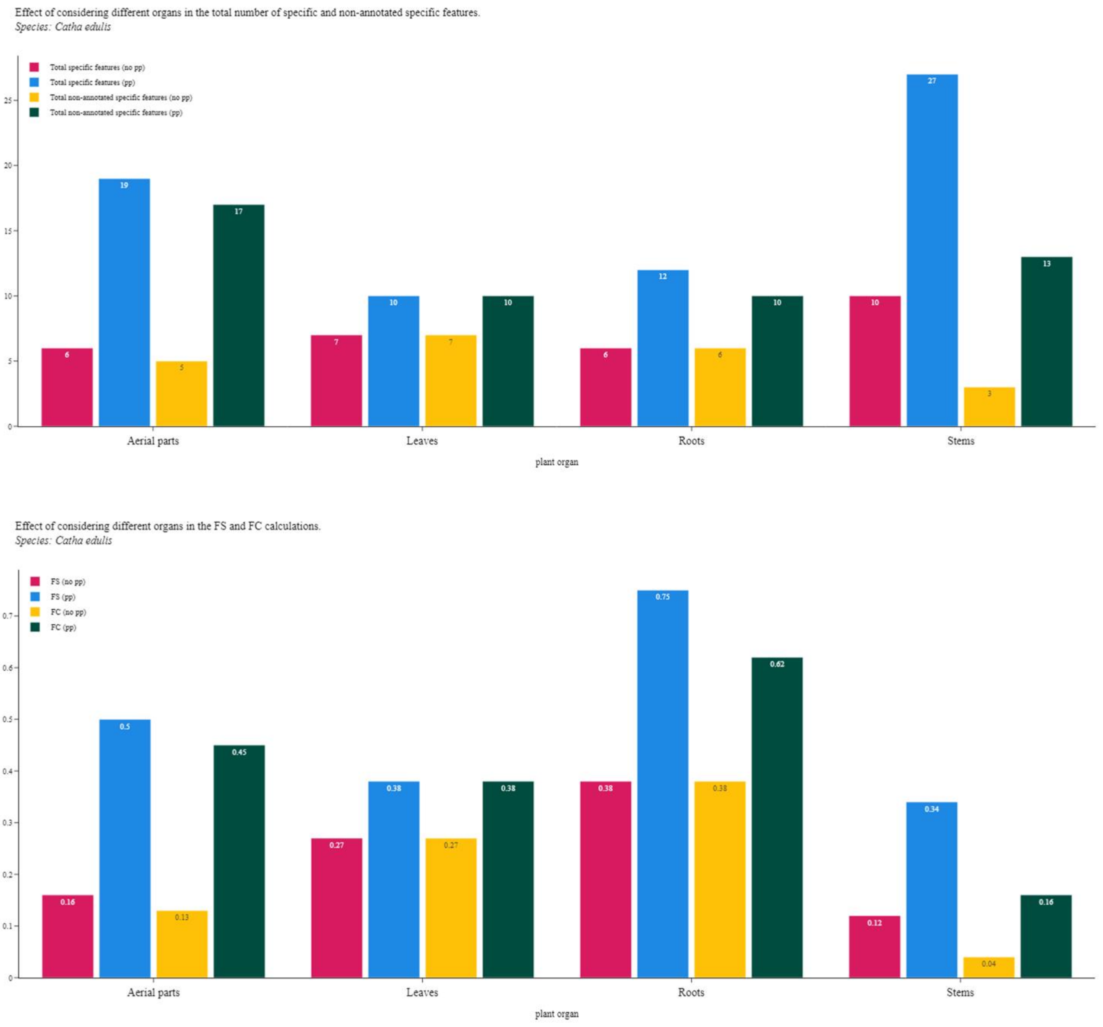


**Supplementary Figure S78**. Upper trace: Effect of considering the presence of different organs for the same species on the total number of specific and non-annotated specific features\. Lower trace: Effect of considering the presence of different organs for the same species on the FS and FC results.

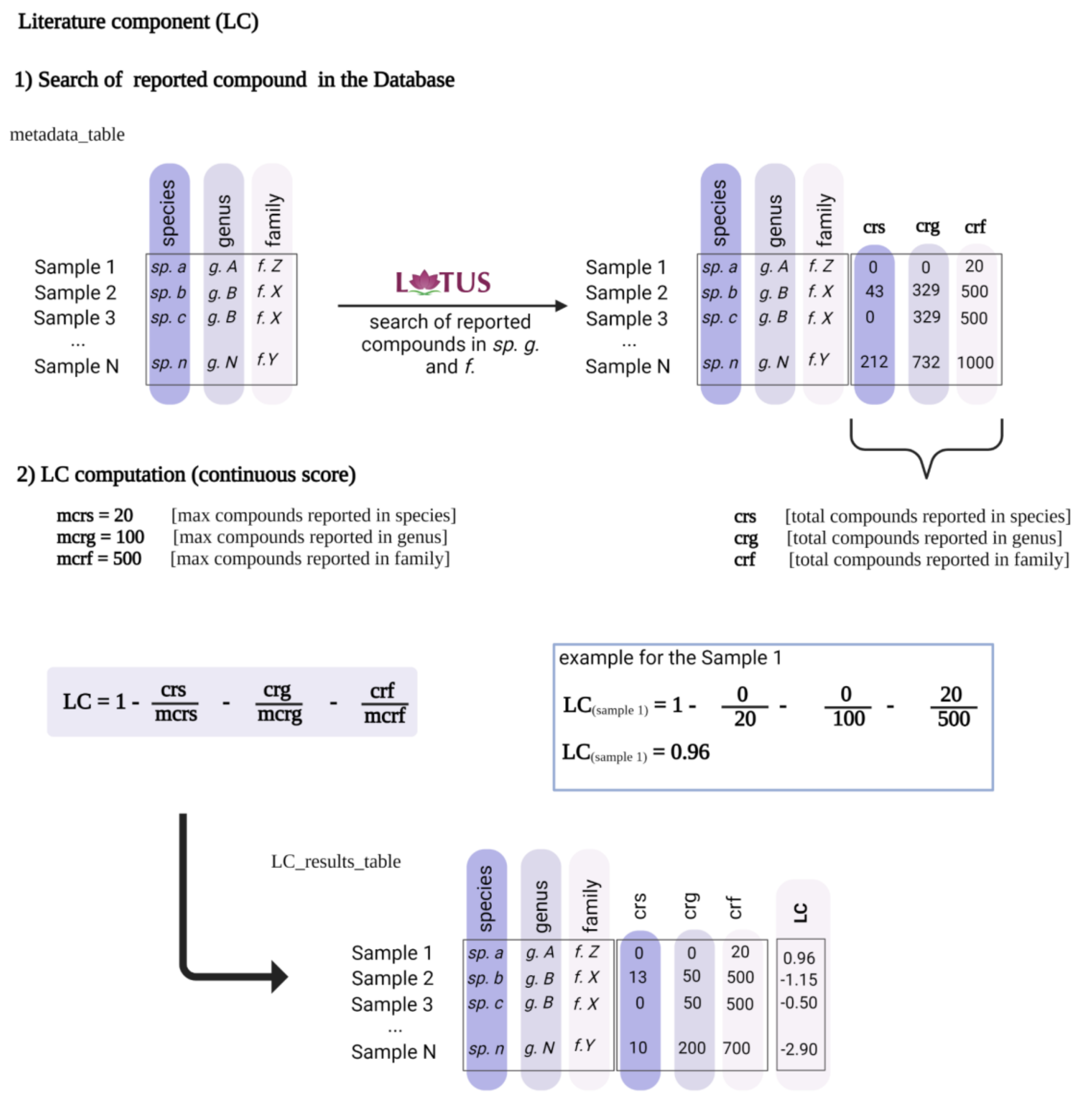


**Supplementary Figure S79.** Graphical representation of the steps followed for the calculation of the Literature Component (LC). The LC takes the taxonomical information for each sample as input. The species should be cleaned-up to have up to date recognized names, the Open Tree of Life (OTL), including the genus and family for each species. 1) The script will create a set{} with the species, genus and families found in the metadata to recover the total number of compounds reported in the database LOTUS for each level, and link the results directly to the samples. 2) The LC score is calculated with the formula shown above. The score is going to be modulated according to the maximum number of reported compounds in species (mrcs), genus (mrcg) and family (mcrf) defined by the user. The larger the number of reported compounds at each level, the lower the LC value.

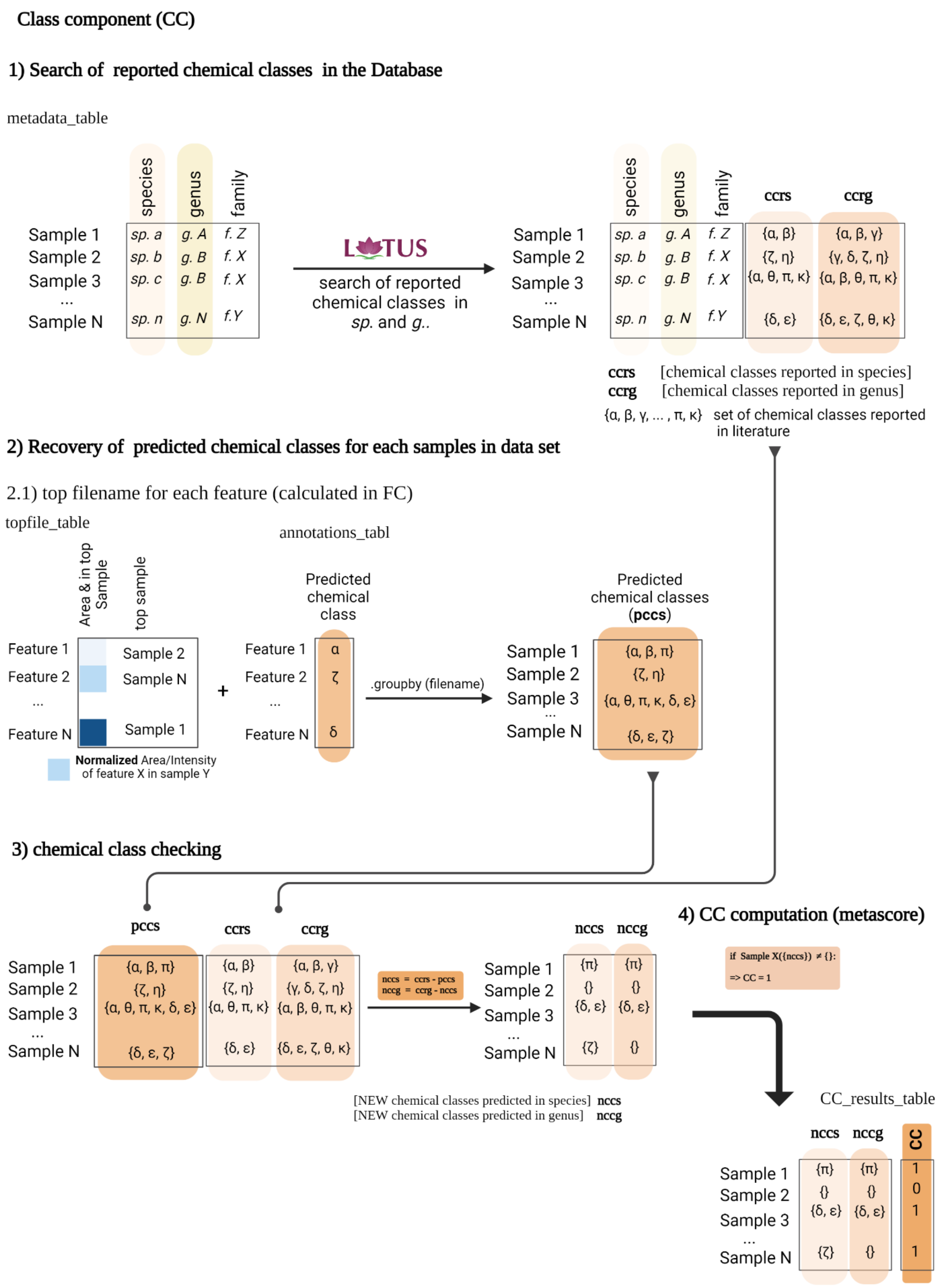
Class component (CC)
1) Search of reported chemical classes in the Database
metadata_table
species
genus
family
Sample 1
Sample 2
Sample 3
...
Sample N
sp. a g. A f. Z
sp. b g. B f. X
sp. c g. B f. X
sp. n g. N f.Y
LOTUS
search of reported chemical classes in sp. and g..
ccrs
ccrg
{α, β} {α, β, γ}
{ζ, η} {γ, δ, ζ, η}
{α, θ, π, κ} {α, β, θ, π, κ}
{δ, ε} {δ, ε, ζ, θ, κ}
ccrs [chemical classes reported in species]
ccrg [chemical classes reported in genus]
{α, β, γ, ... , π, κ} set of chemical classes reported in literature
2) Recovery of predicted chemical classes for each samples in data set
2.1) top filename for each feature (calculated in FC)
topfile_table
Area & in top Sample
top sample
Feature 1 Sample 2
Feature 2 Sample N
...
Feature N Sample 1
Normalized Area/Intensity of feature X in sample Y
+
annotations_tabl
Predicted chemical class
Feature 1 α
Feature 2 ζ
...
Feature N δ
.groupby (filename)
Predicted chemical classes (pccs)
Sample 1 {α, β, π}
Sample 2 {ζ, η}
Sample 3 {α, θ, π, κ, δ, ε}
...
Sample N {δ, ε, ζ}
3) chemical class checking
pccs
ccrs
ccrg
Sample 1 {α, β, π} {α, β} {α, β, γ}
Sample 2 {ζ, η} {ζ, η} {γ, δ, ζ, η}
Sample 3 {α, θ, π, κ, δ, ε} {α, θ, π, κ} {α, β, θ, π, κ}
...
Sample N {δ, ε, ζ} {δ, ε} {δ, ε, ζ, θ, κ}
nccs = ccrs - pccs
nccg = ccrg - nccs
nccs
nccg
Sample 1 {π} {π}
Sample 2 {} {}
Sample 3 {δ, ε} {δ, ε}
...
Sample N {ζ} {}
[NEW chemical classes predicted in species] nccs
[NEW chemical classes predicted in genus] nccg
4) CC computation (metascore)
if Sample X({nccs}) ≠ {}:
=> CC = 1
CC_results_table
nccs
nccg
CC
Sample 1 {π} {π} 1
Sample 2 {} {} 0
Sample 3 {δ, ε} {δ, ε} 1
...
Sample N {ζ} {} 1

**Supplementary Figure S80.** Graphical representation of the steps followed for calculating the Class Component (CC). The CC takes the taxonomical information for each sample, the quantitative table, and the dereplication results from SIRIUS-CANOPUS as input. The species should be cleaned-up to have up to date recognized names, the Open Tree of Life (OTL), including the genus and family for each species, this information is the same used for computing the LC. 1) The script will create a set{} with the species, genus, and families found in the metadata to recover the total of chemical classes for all the compounds reported in LOTUS for each level, and link the results directly to the samples. 2) Because the dereplication information comes from an aligned dataset, it is necessary to find to which sample a particular chemical class belongs, to simplify this, the table calculated in the second step of the FC (**Supplementary Figure 76**) is used, it contains the filename (sample) where each feature is present at the highest percentage. The CANOPUS results are linked to this table and the results are grouped by sample, generating a set of chemical classes predicted for each sample (pccs). 3) The new chemical classes predicted in species (nccs) and in genus (nccg) are obtained by simple subtraction (nccs = ccrs - pccs; nccg = ccrg - nccs). 4) The script will look into the nccs column to check if it is empty or contains any value (string), if there is some chemical class the Metascore CC for that particular sample will be assing as ‘1’, otherwise is ‘0’.

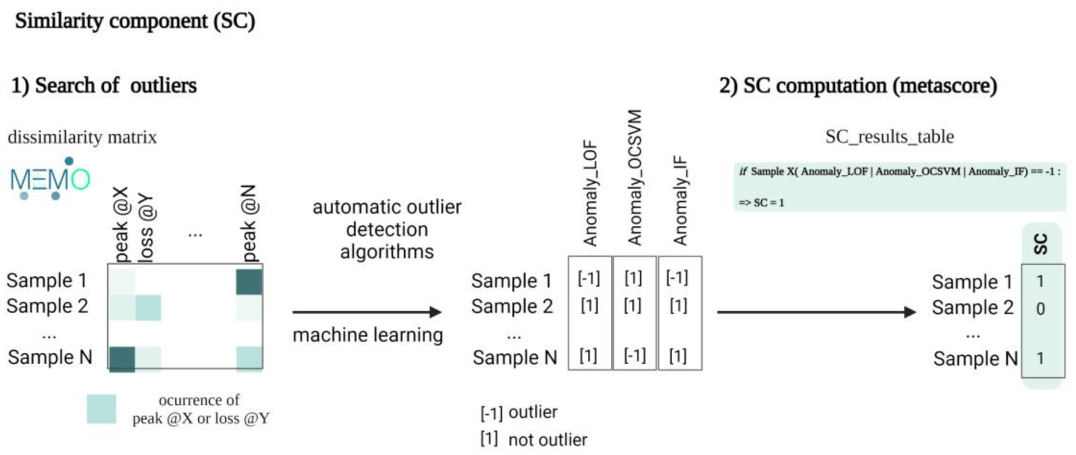

**Supplementary Figure S81.** Graphical representation of the steps followed for calculating the Similarity Component (SC). The SC takes the dissimilarity matrix generated from memo ms as input. 1) Three automatic outlier detectors are run over the set without further parameters to fix. The general parameters were fixed to 'auto'. However, depending on the user and the particularities of the set itself multiple parameters could be tunned. The script will add a column for the results of each of the three algorithms indicating the status, [-1] = outlier, [1] normal point. 2) Once the samples are tagged, the script will evaluate the status before adding the Metascore value '1' for outliers and '0' for normally distributed points accordingly. As explained in the main text, we used three different algorithms to cover a wider range of outliers predictions, the score will be '1' for the samples being 'outlier, [-1]' at least in one of the three algorithms.

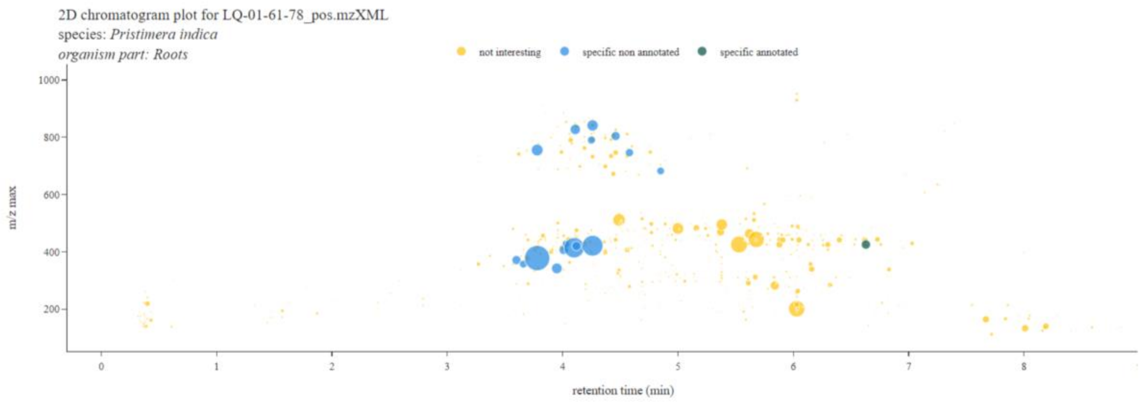

**Supplementary Figure S82.** Feature-based interactive Ion map plots showing the combined results of the FC and CC (if calculated) for the features of a selected sample (user-defined). The features are displayed according to their status (specific non-annotated (blue), specific annotated (green), and non-specific non-annotated -not interesting- (yellow)). Complementary information (adducts, row id, chemical class, etc) are displayed interactively for each feature if available. The intensities in both cases (bar's height and bubble's size) are proportional to the original quantification table. The scatter plot shows the *m/z* ratio of each feature (or ion network identity) on the y-axis.

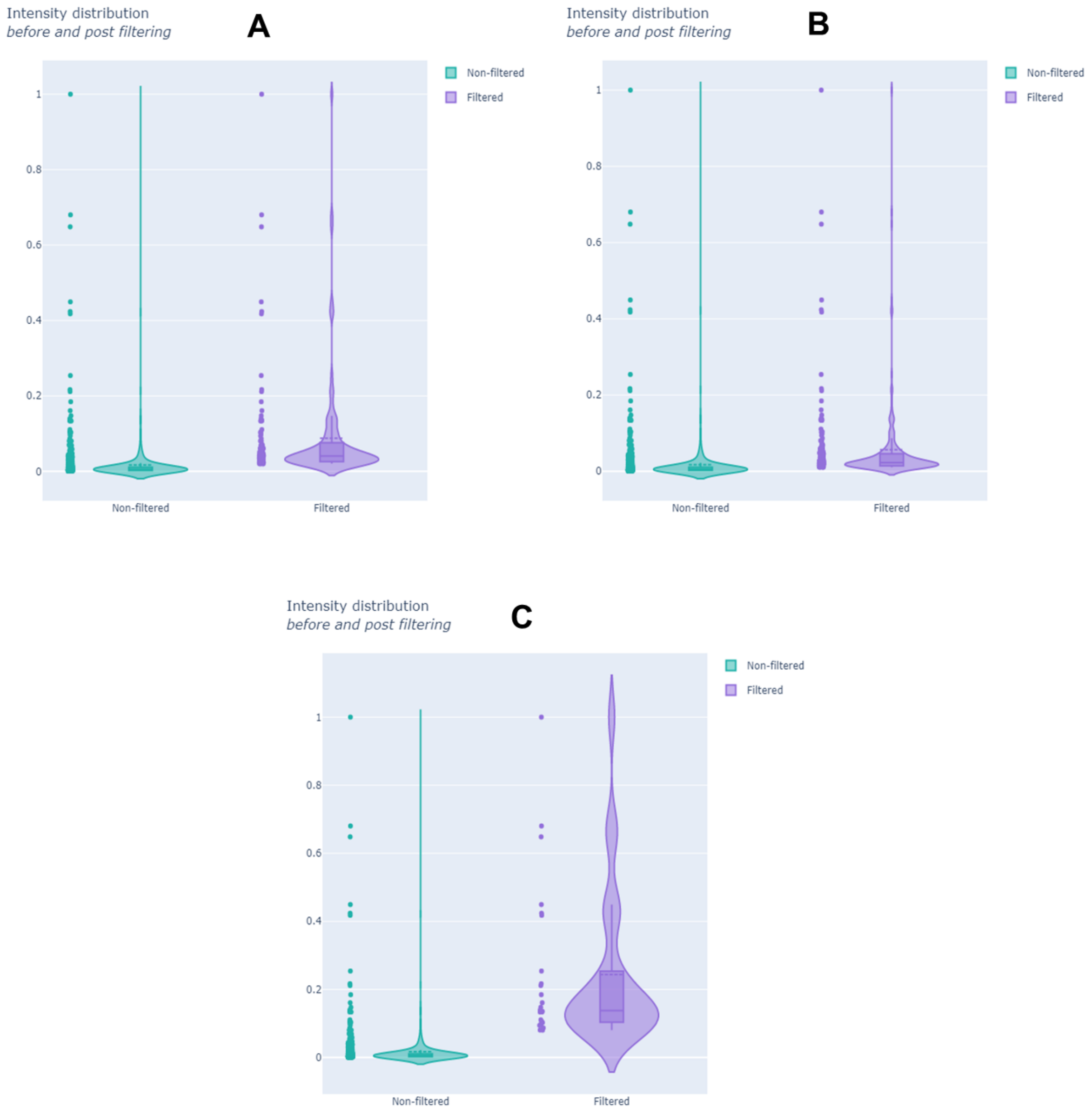


**Supplementary Figure S83**. Visualization of the effects of the filtering steps in a particular sample. A) Only the intensity-based filter was applied, all the values lower than 2% of relative intensity were minimized to zero [interactive figure here]. B) Only the quantile-based filter was applied, all the values lower than the third quantile (0.75) were minimized to zero [interactive figure here]. C) Effects of the sequential application of the intensity filter followed by the quantile-based filter [interactive figure here].

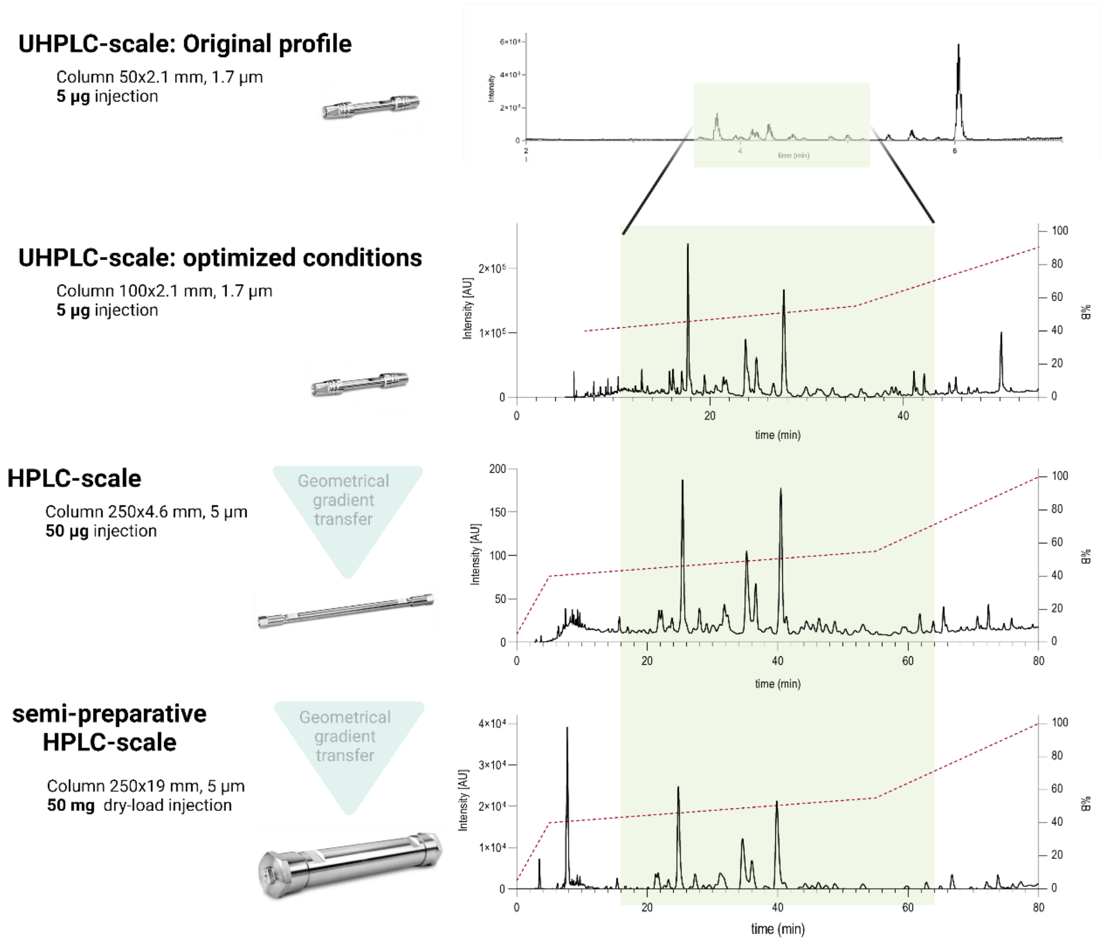


**Supplementary Figure S84**. Chromatograms at different analytical scales. A) UHPLC-254 nm, column Acquity BEH $C_{18}$, 100x2.1 mm, 1.7 $\mu$m. B) HPLC-254 nm, column Xbridge 250x4.6 mm, 5$\mu$m. C) Semi Preparative HPLC-254 nm, column Xbridge 250x19 mm, 5$\mu$m. The dashed line represents the %B [ACN + 0.1% Formic acid]. The region highlighted in light green indicates the region where the features spotted in the network are found.

**Supplementary Table 2**. Inventa Results for the Celastraceae Collection

| Sample ID | FC | Feature specificity | MF prediction ratio | LC | Reported compounds in Species | Reported compounds in Genus | SC | CC | New CC in sp | New CC in genus | PR |
|---|---|---|---|---|---|---|---|---|---|---|---|
| LQ-01-61-78_pos.mzXML | 0.82 | 0.84 | 0.32 | 1.00 | 2 | 8 | 1 | 1 | {'Agarofuran sesquiterpenoids', 'Primary amides', 'Pyridine alkaloids'} | {'Agarofuran sesquiterpenoids', 'Primary amides', 'Pyridine alkaloids'} | 3.82 |
| LQ-01-61-37_pos.mzXML | 0.81 | 0.84 | 0.54 | 1.00 | 1 | 440 | 1 | 1 | {'Chromones', 'Simple coumarins', 'Isoquinoline alkaloids', 'Agarofuran sesquiterpenoids', 'Pyranocoumarins', 'Simple phenolic acids', 'Isocoumarins', 'Furocoumarins'} | {'Chromones', 'Simple coumarins', 'Isoquinoline alkaloids', 'Pyranocoumarins', 'Simple phenolic acids', 'Isocoumarins', 'Furocoumarins'} | 3.81 |
| LQ-01-61-06_pos.mzXML | 0.84 | 0.86 | 0.48 | 0.89 | 212 | 732 | 1 | 1 | {'Tetracyclic diterpenoids', 'Paraliane diterpenoids', 'Open-chain polyketides'} | {'Tetracyclic diterpenoids', 'Paraliane diterpenoids', 'Open-chain polyketides'} | 3.73 |
| LQ-01-61-07_pos.mzXML | 0.70 | 0.71 | 0.50 | 0.96 | 71 | 732 | 1 | 1 | {'Paraliane diterpenoids', 'Labdane diterpenoids', 'Neutral glycosphingolipids', 'Daucane sesquiterpenoids', 'Tetracyclic diterpenoids', 'Dicarboxylic acids', 'Trichothecane sesquiterpenoids', 'Monoacylglycerols', 'Diacylglycerols'} | {'Paraliane diterpenoids', 'Neutral glycosphingolipids', 'Daucane sesquiterpenoids', 'Tetracyclic diterpenoids', 'Dicarboxylic acids', 'Trichothecane sesquiterpenoids', 'Monoacylglycerols', 'Diacylglycerols'} | 3.66 |
| LQ-01-61-05_pos.mzXML | 0.74 | 0.76 | 0.39 | 0.89 | 212 | 732 | 1 | 1 | {'Pyridine alkaloids'} | {'Pyridine alkaloids'} | 3.63 |
| LQ-01-61-56_pos.mzXML | 0.64 | 0.66 | 0.37 | 0.99 | 14 | 338 | 1 | 1 | {'Paraliane diterpenoids', 'Agarofuran sesquiterpenoids', 'Drimane sesquiterpenoids', 'Daucane sesquiterpenoids'} | {'Paraliane diterpenoids', 'Agarofuran sesquiterpenoids', 'Drimane sesquiterpenoids', 'Daucane sesquiterpenoids'} | 3.63 |
| LQ-01-61-54_pos.mzXML | 0.51 | 0.57 | 0.29 | 0.99 | 14 | 338 | 1 | 1 | {'Agarofuran sesquiterpenoids', 'Trichothecane sesquiterpenoids', 'Lanostane, Tirucallane and Euphane triterpenoids'} | {'Agarofuran sesquiterpenoids', 'Trichothecane sesquiterpenoids', 'Lanostane, Tirucallane and Euphane triterpenoids'} | 3.50 |
| LQ-01-61-55_pos.mzXML | 0.50 | 0.52 | 0.33 | 0.99 | 14 | 338 | 1 | 1 | {'Agarofuran sesquiterpenoids', 'Trichothecane sesquiterpenoids', 'Drimane sesquiterpenoids', 'Daucane sesquiterpenoids'} | {'Agarofuran sesquiterpenoids', 'Trichothecane sesquiterpenoids', 'Drimane sesquiterpenoids', 'Daucane sesquiterpenoids'} | 3.50 |
| LQ-01-61-03_pos.mzXML | 0.50 | 0.51 | 0.27 | 0.94 | 126 | 126 | 1 | 1 | {'Ursane and Taraxastane triterpenoids', 'Tetraketide meroterpenoids', 'Triketide meroterpenoids', 'Daucane sesquiterpenoids'} | {'Ursane and Taraxastane triterpenoids', 'Daucane sesquiterpenoids', 'Tetraketide meroterpenoids', 'Triketide meroterpenoids'} | 3.44 |
| LQ-01-61-02_pos.mzXML | 0.49 | 0.50 | 0.29 | 0.94 | 126 | 126 | 1 | 1 | {'Oleanane triterpenoids'} | {'Oleanane triterpenoids'} | 3.42 |
| LQ-01-61-75_pos.mzXML | 0.38 | 0.41 | 0.23 | 0.49 | 1011 | 1353 | 1 | 1 | {'Vitamin D2 and derivatives'} | {'Vitamin D2 and derivatives'} | 2.87 |
| LQ-01-61-60_pos.mzXML | 0.81 | 0.82 | 0.53 | 1.00 | 0 | 0 | 1 | 0 | | | 2.81 |

| | | | | | | | | | | | |
|---|---|---|---|---|---|---|---|---|---|---|---|
| LQ-01-61-47_pos.mzXML | 0.78 | 0.81 | 0.57 | 1.00 | 0 | 0 | 1 | 0 | | | 2.78 |
| LQ-01-61-23_pos.mzXML | 0.78 | 0.81 | 0.50 | 1.00 | 0 | 0 | 1 | 0 | | | 2.78 |
| LQ-01-61-62_pos.mzXML | 0.71 | 0.74 | 0.44 | 1.00 | 3 | 50 | 0 | 1 | {'Dipeptides'} | {'Dipeptides'} | 2.71 |
| LQ-01-61-41_pos.mzXML | 0.68 | 0.78 | 0.39 | 1.00 | 0 | 0 | 1 | 0 | | | 2.68 |
| LQ-01-61-19_pos.mzXML | 0.64 | 0.65 | 0.40 | 1.00 | 0 | 0 | 1 | 0 | | | 2.64 |
| LQ-01-61-58_pos.mzXML | 0.61 | 0.65 | 0.36 | 0.99 | 14 | 211 | 0 | 1 | {'Ursane and Taraxastane triterpenoids', 'Agarofuran sesquiterpenoids'} | {'Ursane and Taraxastane triterpenoids', 'Agarofuran sesquiterpenoids'} | 2.61 |
| LQ-01-61-31_pos.mzXML | 0.58 | 0.60 | 0.39 | 1.00 | 0 | 0 | 1 | 0 | | | 2.58 |
| LQ-01-61-43_pos.mzXML | 0.58 | 0.63 | 0.33 | 0.99 | 15 | 329 | 0 | 1 | {'Flavones', 'Flavonols'} | {'Flavones', 'Flavonols'} | 2.57 |
| LQ-01-61-22_pos.mzXML | 0.55 | 0.58 | 0.29 | 1.00 | 0 | 0 | 1 | 0 | | | 2.55 |
| LQ-01-61-44_pos.mzXML | 0.56 | 0.59 | 0.23 | 0.99 | 15 | 329 | 0 | 1 | {'Lanostane, Tirucallane and Euphane triterpenoids', 'Pregnane steroids', 'Lupane triterpenoids'} | {'Lupane triterpenoids', 'Pregnane steroids', 'Lanostane, Tirucallane and Euphane triterpenoids'} | 2.55 |
| LQ-01-61-73_pos.mzXML | 0.61 | 0.61 | 0.21 | 0.92 | 162 | 1353 | 1 | 0 | | | 2.53 |
| LQ-01-61-72_pos.mzXML | 0.49 | 0.50 | 0.41 | 1.00 | 0 | 0 | 1 | 0 | | | 2.49 |
| LQ-01-61-53_pos.mzXML | 0.45 | 0.49 | 0.28 | 0.98 | 43 | 329 | 0 | 1 | {'Cholestane steroids'} | {'Cholestane steroids'} | 2.43 |
| LQ-01-61-52_pos.mzXML | 0.43 | 0.45 | 0.32 | 0.98 | 43 | 329 | 0 | 1 | {'Cholestane steroids'} | {'Cholestane steroids'} | 2.41 |
| LQ-01-61-01_pos.mzXML | 0.46 | 0.49 | 0.27 | 0.94 | 126 | 126 | 1 | 0 | | | 2.40 |
| LQ-01-61-16_pos.mzXML | 0.44 | 0.50 | 0.29 | 0.95 | 92 | 440 | 0 | 1 | {'Open-chain polyketides'} | {'Open-chain polyketides'} | 2.39 |
| LQ-01-61-04_pos.mzXML | 0.45 | 0.46 | 0.31 | 0.94 | 126 | 126 | 1 | 0 | | | 2.38 |
| LQ-01-61-67_pos.mzXML | 0.51 | 0.57 | 0.37 | 0.84 | 313 | 514 | 0 | 1 | {'Straight chain fatty acids', 'Lanostane, Tirucallane and Euphane triterpenoids'} | {'Straight chain fatty acids', 'Lanostane, Tirucallane and Euphane triterpenoids'} | 2.36 |
| LQ-01-61-66_pos.mzXML | 0.50 | 0.52 | 0.33 | 0.84 | 313 | 514 | 0 | 1 | {'Cholestane steroids', 'Lanostane, Tirucallane and Euphane triterpenoids'} | {'Cholestane steroids', 'Lanostane, Tirucallane and Euphane triterpenoids'} | 2.34 |
| LQ-01-61-30_pos.mzXML | 0.35 | 0.37 | 0.25 | 0.95 | 92 | 440 | 0 | 1 | {'Zearalenones', 'Dicarboxylic acids'} | {'Zearalenones', 'Dicarboxylic acids'} | 2.30 |
| LQ-01-61-71_pos.mzXML | 0.42 | 0.49 | 0.25 | 0.84 | 313 | 514 | 0 | 1 | {'Lanostane, Tirucallane and Euphane triterpenoids'} | {'Lanostane, Tirucallane and Euphane triterpenoids'} | 2.26 |

| | | | | | | | | | |
|---|---|---|---|---|---|---|---|---|---|
| LQ-01-61-74_pos.mzXML | 0.51 | 0.53 | 0.29 | 0.49 | 1011 | 1353 | 1 | 0 | 2.01 |
| LQ-01-61-27_pos.mzXML | 0.83 | 0.90 | 0.51 | 0.99 | 11 | 440 | 0 | 0 | 1.83 |
| LQ-01-61-28_pos.mzXML | 0.76 | 0.81 | 0.63 | 1.00 | 1 | 440 | 0 | 0 | 1.76 |
| LQ-01-61-42_pos.mzXML | 0.71 | 0.71 | 0.41 | 1.00 | 0 | 0 | 0 | 0 | 1.71 |
| LQ-01-61-57_pos.mzXML | 0.71 | 0.82 | 0.38 | 0.99 | 14 | 211 | 0 | 0 | 1.70 |
| LQ-01-61-70_pos.mzXML | 0.67 | 0.68 | 0.28 | 1.00 | 0 | 0 | 0 | 0 | 1.67 |
| LQ-01-61-09_pos.mzXML | 0.70 | 0.70 | 0.19 | 0.96 | 71 | 732 | 0 | 0 | 1.66 |
| LQ-01-61-08_pos.mzXML | 0.70 | 0.70 | 0.48 | 0.96 | 71 | 732 | 0 | 0 | 1.66 |
| LQ-01-61-68_pos.mzXML | 0.66 | 0.66 | 0.34 | 1.00 | 0 | 0 | 0 | 0 | 1.66 |
| LQ-01-61-33_pos.mzXML | 0.65 | 0.68 | 0.50 | 1.00 | 0 | 0 | 0 | 0 | 1.65 |
| LQ-01-61-40_pos.mzXML | 0.64 | 0.65 | 0.42 | 1.00 | 0 | 0 | 0 | 0 | 1.64 |
| LQ-01-61-69_pos.mzXML | 0.63 | 0.74 | 0.37 | 1.00 | 0 | 0 | 0 | 0 | 1.63 |
| LQ-01-61-51_pos.mzXML | 0.62 | 0.65 | 0.50 | 1.00 | 0 | 0 | 0 | 0 | 1.62 |
| LQ-01-61-14_pos.mzXML | 0.61 | 0.62 | 0.35 | 1.00 | 0 | 0 | 0 | 0 | 1.61 |
| LQ-01-61-15_pos.mzXML | 0.65 | 0.71 | 0.32 | 0.95 | 92 | 440 | 0 | 0 | 1.60 |
| LQ-01-61-63_pos.mzXML | 0.59 | 0.63 | 0.43 | 1.00 | 0 | 0 | 0 | 0 | 1.59 |
| LQ-01-61-12_pos.mzXML | 0.57 | 0.61 | 0.40 | 1.00 | 0 | 0 | 0 | 0 | 1.57 |
| LQ-01-61-32_pos.mzXML | 0.56 | 0.59 | 0.37 | 1.00 | 0 | 0 | 0 | 0 | 1.56 |
| LQ-01-61-46_pos.mzXML | 0.55 | 0.57 | 0.43 | 1.00 | 0 | 0 | 0 | 0 | 1.55 |
| LQ-01-61-59_pos.mzXML | 0.55 | 0.56 | 0.37 | 0.99 | 14 | 211 | 0 | 0 | 1.54 |
| LQ-01-61-29_pos.mzXML | 0.58 | 0.62 | 0.33 | 0.95 | 92 | 440 | 0 | 0 | 1.54 |
| LQ-01-61-64_pos.mzXML | 0.53 | 0.57 | 0.43 | 1.00 | 0 | 0 | 0 | 0 | 1.53 |

| | | | | | | | | | |
|---|---|---|---|---|---|---|---|---|---|
| LQ-01-61-61_pos.mzXML | 0.48 | 0.52 | 0.33 | 1.00 | 0 | 0 | 0 | 0 | 1.48 |
| LQ-01-61-20_pos.mzXML | 0.46 | 0.46 | 0.28 | 1.00 | 2 | 440 | 0 | 0 | 1.46 |
| LQ-01-61-35_pos.mzXML | 0.45 | 0.51 | 0.29 | 1.00 | 0 | 0 | 0 | 0 | 1.45 |
| LQ-01-61-45_pos.mzXML | 0.45 | 0.49 | 0.32 | 1.00 | 0 | 0 | 0 | 0 | 1.45 |
| LQ-01-61-11_pos.mzXML | 0.45 | 0.49 | 0.29 | 1.00 | 0 | 0 | 0 | 0 | 1.45 |
| LQ-01-61-10_pos.mzXML | 0.44 | 0.50 | 0.35 | 1.00 | 0 | 0 | 0 | 0 | 1.44 |
| LQ-01-61-18_pos.mzXML | 0.44 | 0.44 | 0.28 | 1.00 | 0 | 0 | 0 | 0 | 1.44 |
| LQ-01-61-21_pos.mzXML | 0.43 | 0.46 | 0.24 | 1.00 | 0 | 0 | 0 | 0 | 1.43 |
| LQ-01-61-38_pos.mzXML | 0.42 | 0.43 | 0.28 | 1.00 | 1 | 440 | 0 | 0 | 1.42 |
| LQ-01-61-34_pos.mzXML | 0.42 | 0.42 | 0.22 | 1.00 | 0 | 0 | 0 | 0 | 1.42 |
| LQ-01-61-36_pos.mzXML | 0.40 | 0.41 | 0.26 | 1.00 | 0 | 0 | 0 | 0 | 1.40 |
| LQ-01-61-26_pos.mzXML | 0.39 | 0.42 | 0.24 | 1.00 | 0 | 0 | 0 | 0 | 1.39 |
| LQ-01-61-17_pos.mzXML | 0.38 | 0.40 | 0.15 | 1.00 | 0 | 0 | 0 | 0 | 1.38 |
| LQ-01-61-39_pos.mzXML | 0.38 | 0.38 | 0.28 | 1.00 | 1 | 440 | 0 | 0 | 1.38 |
| LQ-01-61-50_pos.mzXML | 0.38 | 0.41 | 0.22 | 1.00 | 0 | 0 | 0 | 0 | 1.38 |
| LQ-01-61-13_pos.mzXML | 0.37 | 0.42 | 0.29 | 1.00 | 0 | 0 | 0 | 0 | 1.37 |
| LQ-01-61-24_pos.mzXML | 0.37 | 0.40 | 0.27 | 1.00 | 0 | 0 | 0 | 0 | 1.37 |
| LQ-01-61-48_pos.mzXML | 0.36 | 0.44 | 0.18 | 1.00 | 0 | 0 | 0 | 0 | 1.36 |
| LQ-01-61-49_pos.mzXML | 0.33 | 0.33 | 0.15 | 1.00 | 0 | 0 | 0 | 0 | 1.33 |
| LQ-01-61-25_pos.mzXML | 0.30 | 0.31 | 0.22 | 1.00 | 0 | 0 | 0 | 0 | 1.30 |
| LQ-01-61-65_pos.mzXML | 0.29 | 0.30 | 0.24 | 0.84 | 313 | 514 | 0 | 0 | 1.13 |

**Supplementary Table S3**. Putative identities proposed for the extract of *Pristemira indica* roots.

| cluster index | Exact mass (*m/z*) | Retention time (min) | Molecular Formula | Libname | Organism | Structure InChI |
|---|---|---|---|---|---|---|
| 14749 | 768.3211 | 4.60 | $C_{40}H_{49}NO_{14}$ | ISDB (CAS 1403757-10-7) | Celastrus angulatus | InChI=1S/C40H49NO14/c1-21(2)33(44)49-20-39-30(53-36(47)26-17-13-14-18-41-26)27(50-23(5)42)19-38(9,48)40(39)31(51-24(6)43)28(37(7,8)55-40)29(52-34(45)22(3)4)32(39)54-35(46)25-15-11-10-12-16-25/h10-18,21-22,27-32,48H,19-20H2,1-9H3 |
| 8921 | 465.3001 | 6.04 | $C_{30}H_{46}O_7$ | CCMSLIB00004693704 | Tripterygium wilfordii | InChI=1S/C30H40O4/c1-18-19-8-9-22-28(4,20(19)16-21(31)24(18)32)13-15-30(6)23-17-27(3,25(33)34-7)11-10-26(23,2)12-14-29(22,30)5/h8-9,16,23,32H,10-15,17H2,1-7H3/t23-,26-,27-,28+,29-,30+/m1/s1 |
| 95 | 421.2739 | 4.77 | $C_{10}H_{14}N_4O_4$ | CCMSLIB00000078862 | Maytenus | InChI=1S/C28H36O3/c1-16-13-23-25(3,15-21(16)30)9-11-27(5)22-8-7-18-17(2)24(31)20(29)14-19(18)26(22,4)10-12-28(23,27)6/h7-8,14,16,23,31H,9-13,15H2,1-6H3/t16-,23?,25+,26+,27-,28+/m1/s1 |
